# Rice WRKY13 TF protein binds to motifs in the promoter region to regulate downstream disease resistance-related genes

**DOI:** 10.1101/2023.01.18.523945

**Authors:** John Lilly Jimmy, Rohit Karn, Sweta Kumari, Chitathoor Balasubramane Sruthilaxmi, Singh Pooja, Subramanian Babu

**Author notes:** Corresponding author: Jimmy John.

## Abstract

In plants, pathogen resistance is brought about due to the binding of certain transcription factors (TF) proteins to the cis-elements of certain target genes. These cis-elements are present up-stream in the motif of the promoters of each gene. This ensures the binding of a specific transcription factor to a specific promoter, therefore regulating the expression of that gene. Therefore, the study of each promoter sequence of all the rice genes would help identify the target genes of a specific transcription factor. Rice 1kb upstream promoter sequences of 55,986 annotated genes were analyzed using the Perl program algorithm to detect WRKY13 binding motifs (bm). The resulting genes were grouped using gene ontology and gene set enrichment analysis. Gene with more than 4 TFbm in their promoter was selected. Nine genes reported to have a role in rice disease resistance were selected for further analysis. *Cis*-acting regulatory element analysis was carried out to find the cis-elements and to confirm the presence of the corresponding motifs in the promoter sequences of these genes. The 3D structure of WRKY13 TF and the corresponding nine genes were built and the interacting residues were determined. The binding capacity of WRKY13 to the promoter of these selected genes was analyzed using docking studies. *WRKY*13 was also considered for docking analysis based on the prior reports of autoregulation. Molecular dynamic simulations provided more details regarding the interactions. Expression data revealed the expression of the genes that helped to provide the mechanism of interaction. Further co-expression network helped to characterize the interaction of these selected diseases resistance-related genes with WRKY13 TF protein. This study suggests the target downstream genes that are regulated by WRKY13 TF. The molecular mechanism involving the gene network regulated by WRKY13 TF in disease resistance against rice fungal pathogens is explored.

## Introduction

Transcription factors (TF) are regulatory proteins that modulate the expression of several other genes by binding to their promoter regions thereby leading to transcription. Apart from transcription modulation, TF responds to environmental signals by regulating gene expression. Stress adaptive regulatory mechanisms could be better understood if the functions of TF during stress conditions could be known. Target genes that have specific *cis*-acting regulatory elements corresponding to a particular TF will be regulated by that TF. A single gene may contain several types of *cis*-acting regulatory elements, thereby the target gene may be regulated by different families of TF (Wang *et al*., 2016). Co-expressing or cross-regulating TFs may be identified better by searching the molecular targets and can be used for crop improvement towards tolerance to different stresses. Multiple stress-responsive TF genes can be identified from their expression pattern and recognise the regulated genes that can be utilized for universal stress responses (Prasch and Sonnewald, 2015). TFs are the apparent candidates for genetic engineering to breed stress-tolerant crops due to their function of regulating various stress-responsive genes (Wang *et al*., 2016). According to the Plant Transcription Factor Database v4.0, there are 56 TF families in rice. There are around 101 WRKY TF in *japonica* and *indica* rice (Jimmy and Babu, 2019). WRKY TF is one of the largest TF families and is abbreviated by four amino acids tryptophan (W), arginine (R), lysine (K), and tyrosine (Y).

In rice, *WRKY*13 TF gene-mediated signalling in plant physiological pathways related to development and disease resistance. WRKY13 was described to act as a positive regulator of two cis-elements: PRE2 and PRE4 present in its promoter region. Several proteins were described to bind to these *cis-*elements that activated *WRKY*13 during a defense response (Cai *et al*., 2008). *WRKY*13 TF was reported by Qiu *et al*. (2007) to mediate other defense-related genes that are involved in salicylic acid (SA) and jasmonic acid (JA) dependent signalling (Qiu *et al*., 2007). JA, a cyclopentanone derived from linolenic acid has a regulatory role against fungal infection (He *et al*., 2017). Linolenic acid was found to be significantly enriched in rice leaves infected with *R. solani* indicating an active JA-signaling pathway during fungal infection (et al.,). Whereas, in *Arabidopsis thaliana*, WRKY13 TF was described to bind to the promoter of *NST*2, a gene related to secondary cell wall synthesis necessary for the development of sclerenchyma cells. *WRKY*13-knockout mutant plants had reduced stem diameter and a low number of sclerenchymatous cells (Li *et al*., 2015). *WRKY13* also promoted flowering under short-day conditions in *Arabidopsis* (Li *et al*., 2016).

In rice, *WRKY*13 act as a potential regulator of various physiological processes; also influence more than

500 genes involved in various plant functions: multiple signalling pathways belonging to disease resistance, redox homeostatic, abiotic stress responses and development. Qiu *et al*., (2009) stated that the gene upregulated by the expression of *WRKY*13 was monitored by other WRKY genes, whereas, the genes downregulated by the expression of *WRKY*13 were monitored by MYB and AP2/EREBP proteins (Qiu *et al*., 2009). Rice *WRKY*13 gene is involved in gene cascades and networks including other WRKYs and other proteins; also acts as a gene responsible for cross-talk between biotic and abiotic stresses. In a *WRKY*45-2-*WRKY*13-*WRKY*42 regulatory transcriptional cascade, *WRKY*13 expression was necessary to suppress the expression of *WRKY*42 against *Magnaporthe oryzae* infection that causes blast disease in rice (Cheng *et al*., 2015). *WRKY*13 up-regulated against two fungal diseases, sheath blight caused by *Rhizoctonia solani* and sheath rot caused by *Sarocladium oryzae. WRKY13* upregulation was associated with the expression of *TIFY*9, *WRKY*12 and *PR*2 (Jimmy and Babu, 2019).

*WRKY*13 activated rice resistance against both fungal and bacterial pathogens and acted as a mediator of cross-talk between the genes of the pathogen-induced salicylic-dependent defense pathway and five other physiological pathways (Qiu *et al*., 2008). A network where *WRKY*13 acted as a positive regulator, was traced out in rice against bacterial leaf blight (BLB). *WRKY*13-overexpressing rice revealed about 236 and 273 genes were enhanced and suppressed respectively when infected by BLB (Qiu *et al*., 2008). The *WRKY*13 gene regulates antagonistic crosstalk by suppressing genes responsible for drought resistance (*SNAC*1) and rice-bacterial interaction (*WRKY*45-1) (Xiao *et al*., 2013). The gene targets regulated by WRKY13 during fungal infections was not traced out yet. Therefore, a systematic study was designed that utilizes a Perl program algorithm to find the possible binding locations of the WRKY13 TF. Cis-acting regulatory element analysis was used to find the cis-elements and to validate the presence of WRKY13 TF binding motifs (bm) in the promoter regions of the genes involved in disease resistance selected from the genome-wide promoter analysis. A docking study between the modelled WRKY13 TF protein was carried out with these selected disease resistance-related genes.

## Materials and Methods

### Data Retrieval, Preprocessing and Genome-wide search of WRKY13-controlled genes

Target genes controlled by WRKY13 TF were identified using genome-wide search using computational analysis of 1 kb promoter sequences of all the genes from the rice genome. Perl program was used to carry out promoter analysis and an algorithm was designed to find each binding motif in the promoter regions of all rice genes. Promoter sequences (1 kb upstream) of all the 55,986 genes of rice were retrieved from the Rice Genome Annotation Project Database (RGAP; http://rice.plantbiology.msu.edu/) (Kawahara *et al*., 2013). Runtime error was avoided through data processing and fine-tuning to a uniform format. Any redundant sequences were removed. These sequences were saved in the FASTA format.

### Identification of WRKY13 TF Binding Sites

Binding sites specific to WRKY13 were found through available literature. Binding motifs were selected to analyze the WRKY13 TF-controlled genes and their network-associated genes. Therefore, 20 bm were selected for the study, out of theses, 14 were WRKY13 bm and 6 were WRKY general bm.

### Determination of WRKY13 TF Binding Sites in Rice Gene Promoters

WRKY13 TF bm distribution in rice is not been reported. Perl program algorithm was designed to find each bm in the promoter sequences of all the rice genes as per the protocol mentioned by Pooja *et al*. (2015) (Figure 1). The locations and the frequencies of occurrence of each of the bm on the sequence were noted. This data helped to create a small library of promoter sequences with the highest occurrence frequency of bm. Since the frequency of protein binding increases with the increase in the availability of the binding regions (Lifanov et al., 2003), the promoter sequences with a bm frequency greater than four were included in the study. The gene descriptions of these promoter sequences were then obtained from the RAP-DB (https://rapdb.dna.affrc.go.jp/ Sakai *et al*., 2013). The locus IDs (RAP IDs) of all these sequences were manually converted to RAGP rice gene locus IDs using the RAP-DB converter tool. These RAGP locus IDs were used for further analysis.

**Figure 1.**
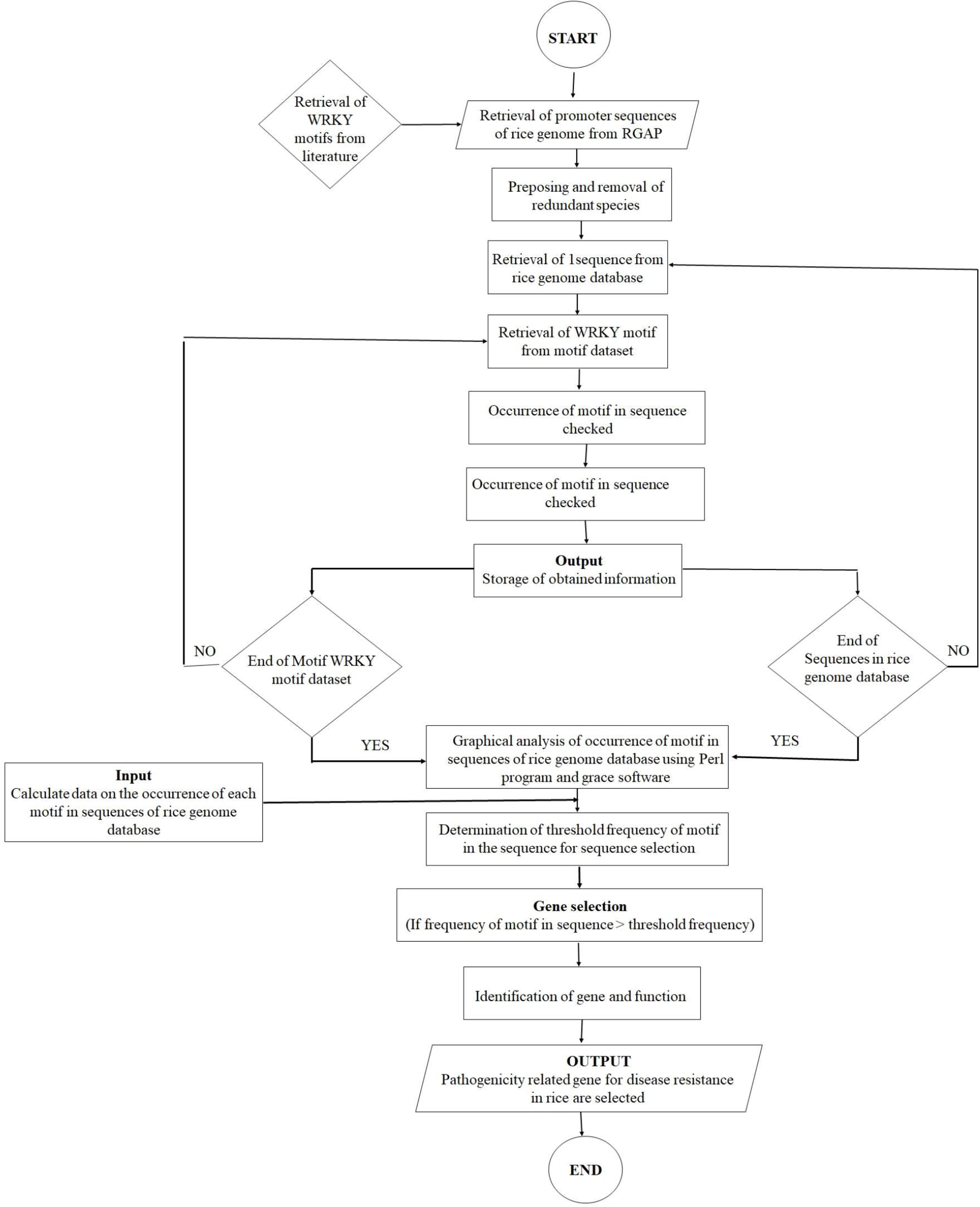
Flow Chart for Analysis of Promoter Sequences to search WRKY13 TFbm in the Rice Genome

### Gene Ontology (GO) and Gene Set Enrichment Analysis

The genes obtained were classified based on the function of their resulting proteins using gene ontology (GO) analysis of molecular processes, biological processes and cellular components performed with PANTHER v14.0 (Protein Analysis Through Evolutionary Relationships) classification system software (http://www.pantherdb.org, Mi *et al*., 2019). The converted IDs were used as input to carry out the classification using gene list analysis. The organism was selected as ‘*Oryza sativa’* and the type of analysis was ‘Functional classification viewed in graphic charts’. These genes were classified based on the function of their proteins using gene ontology (GO) analysis with PANTHER software. Gene set enrichment analysis of all the genes found from the promoter analysis was carried out using AgriGO v1.2 (http://bioinfo.cau.edu.cn/agriGO/ analysis.php, Du *et al*. 2010). The list of genes was used as an input to derive a list of the significant term that was used as an input for Singular Enrichment Analysis (SEA), the organism (*O. sativa*) was selected and default parameters were used. The GO enrichment data was transferred to REVIGO (http://revigo.irb.hr/, Supek *et al*., 2011), a tool designed to remove redundant terms, calculate, and summarize the GO terms appropriately into their enrichments: biological process, cellular components and molecular function. The allowed similarity value was set to 0.5 and visualization of these GO terms was based on their semantic similarities in scatterplots. Genes related to disease resistance were selected for further analysis.

### Target Gene selection based on the presence of WRKY13 TFbm

Promoter sequences of the selected target genes that revealed the presence of a high density of bm were classified for disease resistance and subjected to gene identification using the algorithm (Figure 2). Gene descriptions were identified using Rice Annotation Project Database (RAP-DB, https://rapdb.dna.affrc.go.jp/, Sakai *et al*., 2013). The IDs were converted and individually searched as described previously. The chromosome number, their location on the chromosome and the resulting gene descriptions were noted and recorded in a tabular form with their corresponding locus IDs. The target genes were *aminotransferase*, *ankyrin*, germin 8-7 (*ger*6), β-Glycosidase14, *PR*1b, *PR*2, *PR*5, shikimate biosynthesis protein (*aroDE*), *TIFY*9 and *WRKY*12 and were selected genes and were subjected to further analysis.

**Figure 2.**
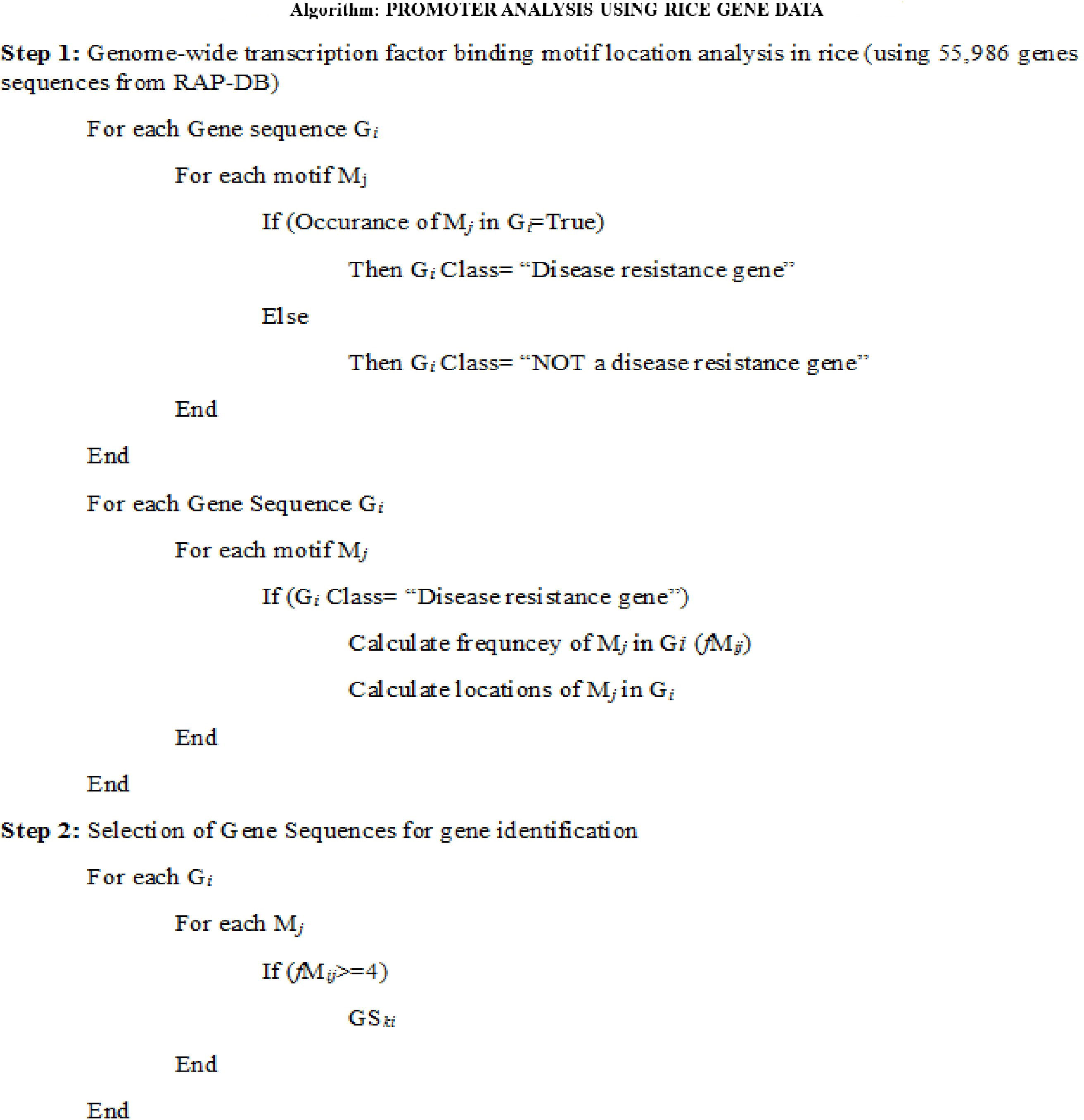
The algorithm of the Perl Program used for the Promoter Sequence Analysis of WRKY13 Gi refers to the individual gene sequence from RAP-DB, Mj represents the individual TF bm of WRKY13, fMij represents the frequency of Mj the TF bm in Gi the sequence, GSk refers to the individual sequence selected for further gene analysis.

### Cis-Acting Regulatory Elements Analysis

Cis-acting regulatory elements (bm) in the promoter sequences were identified in the selected target disease resistance genes. This was carried out using the Plant Cis-Acting DNA Elements (PLACE) database (Higo *et al*., 1999). Promoter sequences of the selected target genes were used as input. The output of the analysis – factor site name (bm name) and the strand (+ or -) were recorded. This was reconfirmed using plant Promoter Analysis Network (PAN) v3.0 (http://plantpan.itps.ncku.edu.tw/, Chow *et al*., 2019). A visualization of the position of these cis-acting regulatory elements in the promoter of the target disease resistance genes was obtained from plant PAN.

### Modelling of WRKY13 TF and Promoter Sequence

The protein sequence of the TF and the DNA sequence of the promoter sequences were retrieved from the GenBank database of the NCBI server. Since there were no experimentally reported structures or WRKY13 TF, an *in silico* approach for modeling the 3-dimensional structure of the protein was used. BlastP revealed no appropriate template hence a protein structure homology modeling workspace, Swiss-model (Arnold *et al*., 2006), was employed to model the structure of WRKY13 TF. The DNA double helical structure for the promoter sequences was modelled using the online DNA sequence to structure tool (Arnott *et al*., 1976). The energy minimization was done for the protein (taken as a ligand) using the Swiss-PDB viewer tool v4.1.0 (http://expasy.org/spdbv/, Guex and Peitsch, 1997). The protein structure was subjected to Structure Assessment, a tool provided by Swiss-model (Arnold *et al*., 2006). The protein structure was used as an input to inspect the Phi and Psi Ramachandran plot. The model was selected for further analysis with more than 90% residues in the favourable region was considered a good-quality protein structure and was selected for further analysis.

### Interaction analysis by docking of WRKY13 with Selected Promoters

The modeled structures were further used for docking studies. Autodock v4.2 software was used to carry out docking studies that employ the preparation of the receptor (DNA) by the addition of hydrogens and conversion of PDB to pdbqt file. The ligand (Protein) was assigned nonpolar hydrogens. Docking simulations were carried out using the Lamarckian Genetic algorithm (LGA) which was a reliable and efficient method of Autodock. The grid points for Autogrid calculations were set to 106 x 126 x 72 Å to consider the motif region at the centre of the grid box. The docking parameters were set to an LGA calculation of 10,000 runs. The energy evaluations were set to 1,500,000 and 27,000 generations. The population size was set to 150 and the rate of gene mutation and the rate of gene crossover were set to 0.02 and 0.8 respectively. The obtained confirmations were then collected and extracted by using the Autodock tool and summarized. Interaction analysis of DNA-protein complexes was carried out using a fully automated protein-ligand interaction profiler (PLIP) (http://projects.biotec.tu-dresden.de/plip-web, Salentin *et al.,* 2015). PLIP is used to identify residues involved in DNA-protein interaction. The PLIP web server helps to detect the residues involved in DNA-protein interactions through comprehensive detection and visualization. The input was the DNA-protein complexes obtained from the Autodock tool. The residues and the type of bonds were noted. Hydrogen bonding and interactions of WRKY13-DNA were recognized using HBPLUS and Nucplot programs (McDonald and Thornton, 1994; Luscombe et al., 1997).

### Molecular dynamic simulation of docked WRKY13 and Selected Promoters

The docked molecules of WRKY13-selected promoters were subjected to MD simulation using Gromacs 2018.4 software package (Abraham et al., 2015). This was carried out using the AMBER94 force field (Hamzeh-Mivehroud *et al*., 2015). The docked complex (WRKY13-selected promoters) was solvated in cubic water cox using Simple Point Charge (SPC) water model (Wu et al., 2006). The ions were added to the entire system to neutralize by substituting the water molecules that ensured the charge neutrality of the docked molecules. Steric clashes were removed by minimizing the energy for 50,000 cycles. The minimized system was then equilibrated into NPT and NVT for 5000 ps. The temperature and pressure of the system to 300K and 1 bar respectively were maintained using the Vrescale and Parrinello-Rahman pressure coupling method (5, 6). This equilibrated system was then subjected to the production run at 300 K and 1 bar pressure for 100000 ps. The dynamic stability and behaviours of the residues of all the docked complexes were analyzed using Gromacs inbuilt tools and VMD. Thus, the radius of gyration (Rg), root mean square fluctuation (RMSF), solvent accessible surface area (SASA), root mean square deviation (RMSD) and hydrogen bond were plotted using the MD trajectory on grace software.

### Gene network analysis

The locus ID for all the 11-selected genes was retrieved from the Rice annotation project database (RAP-DB; http://rapdb.dna.affrc.go.jp/gbl/) (Cao *et al*.,2012). The obtained gene locus IDs were used to download the expression profiles of WRKY genes using the Rice Oligonucleotide Array Database (ROAD) (http://www.ricearray.org/) (Cao *et al*.,2012). The genes were searched for their response against the two pathogens: *Xanthomonas oryzae* pv *oryzae* (*Xoo*) and *M. grisea*. Gene Expression Data (GED) datasets that had an expression for both or either of the pathogens were selected. The heatmap obtained for GED dataset accession ID was GSE9653 from the selected platform NSF20 K. The default value of the PCC cutoff in ROAD is 0.8, for biotic stresses. The expression values of all the 11-disease resistance-related genes were used as an input in Cytoscape 3.4.0 (Shannon *et al*., 2003) based for visualization of the network and to provide searching and analyzing information on the molecular interactions around a set of genes. The networking genes were classified according to their functions using the GO terms and assignments for rice genes mentioned in the Gramene database (http://www.gramene.org/) (Jaiswal et al., 2006) and the RAP rice gene locus IDs were converted to RAGP IDs using RAP-DB converter tool (http://rapdb.dna.affrc.go.jp/tools/converter) (Cao *et al*.,2012).

## Results

### Binding Sites of WRKY13 TF in Rice Promoters

Earlier literature reports led to 20 motifs as probable binding sites (Table 1). Occurrences for all these WRKY13 TFbm were searched in all the rice chromosomes. Promoter sequences of all the 55, 986 genes in the rice genome were obtained from RAP-DB. The occurrences of these bm are enlisted in Table 2 and represented using Figure 3. The maximum number of motifs (1,09,545) was observed in the promoter regions of the genes located in chromosome 1. This was followed by chromosome 3 (94,747) and chromosome 2 (87,603) and chromosome 4 (70196). The promoter regions of the genes located in chromosomes 9, 10, 11 and 12 contained comparatively a smaller number of WRKY13 TF bm. The motif GGTTAGTTA was observed to occur the least in the promoters of the genes in all the chromosomes, followed by the motifs TTGACCTC, TTTTCCAC, GTACGTAC, and GTTGACC.

**Figure 3.**
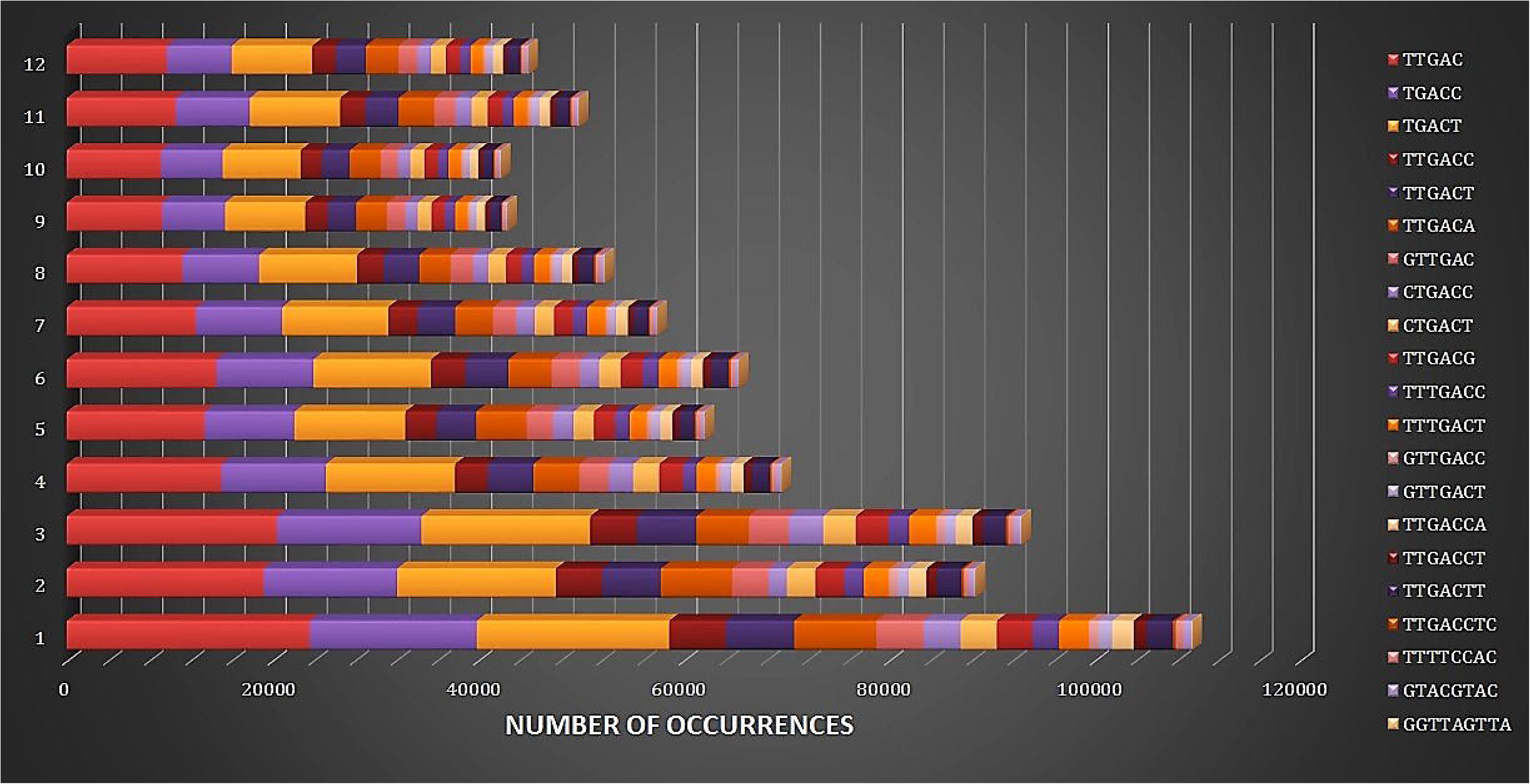
Occurrence and Distribution of WRKY13 bm in the Rice Genome X-axis— number of occurrences; Y-axis—rice chromosomes.

**Table 1.** Binding Motifs of WRKY13 TF

**Table 2.** Occurrences and Chromosome-Wise Distribution of WRKY13 bm in Rice Genome

The promoters of the genes of chromosome 1 contained multiple occurrences of all the motifs. Also, motifs TGACC and TGACT were observed to occur 6 times and 9 times within 90-600 sequence of 1 kb sequence of locus ID Os01t0104900-01 (Transferase family protein) and Os01t0202800-00 (peroxisomal membrane protein), respectively in chromosome 1. Followed by an occurrence of TGACT 7 times within 43-800 sequence of 1 kb sequence of locus IDs Os01t0788900-01 (Pentatricopeptide repeat domain.), Os01t0937300-01 and Os01t0937300-02 (NB-ARC domain). In chromosome 2, an occurrence of 7 times of motif CTGAC within 100-400 sequences of 1kb sequences of locus ID Os02t0741900-01 (T23E23.20) followed by motif TGACT that had an occurrence of 6 times in 150-900 kb sequences of 1 kb sequences of locus IDs Os02t0158100-01 (EF-hand domain) and Os02t0704800-01 (Ornithine carbamoyl transferase). TTGAC had an occurrence of 6 times in 17-900 kb sequences of 1 kb sequences of locus IDs Os02t0538000-01 (Threonyl-tRNA synthetase) and Os02t0704800-01 (Ornithine carbamoyl transferase). Whereas, chromosome 3 revealed an occurrence of 9 times of TTGAC for 190-1000 kb sequence of locus ID Os03t0822100-01 (Transposase). This was followed by an occurrence of 7 times of the motif CTGAC in Os03t0812800-00 (Calcium-binding allergen Olee8) and 7 times of motif TTGAC Os03t0159200-01 (Region of unknown function XS domain) in 190-1000 kb sequence and LOC_Os03g60620 (Non-cell-autonomous heat shock cognate protein 70) in 30-860 kb sequence. In chromosome 4, motif TGACC had 7 times occurrences in 300-650kb of sequences of locus ID Os04t0398900-01 (LEML3 - Anther-specific LEM1 family protein precursor), whereas, in chromosome 5, motif TGACC had 7 times occurrences in 17-750 kb of a sequence of locus ID Os05t0357600-01 (Developmentally regulated GTP binding protein). In chromosome 6, Os06t0208151-00 (Quinone oxidoreductase) revealed 8 times occurrences in 160-760 kb of promoter sequences of the motif TGACT and 7 times occurrences in 18-900 kb promoter sequences of Os05t0346901-00 (TATA-binding protein-associated factor), Os06t0308900-01 (DHHC zinc finger domain) and Os06t0313440-00 (SAM-dependent carboxyl methyltransferase domain). Chromosome 7 revealed two genes Os07t0290800-01 (Tic22-like family protein) and Os07t0623200-02 (heavy metal-associated domain) had occurrences of TTGAC in the promoter sequences of 15-560 kb. In chromosome 8, the motif TTGAC had occurrences of 7 times in the 220-750 kb sequence of the promoter of Os08t0526500-01 (SOX-1 protein). The gene Os10t0502500-04 (Cytochrome b5 domain) had TGACT occurrence of 9 times in the promoter sequence of 113-920 kb, whereas, motif TTGAC had an occurrence of 8 times in 530-720 kb in Os10t0502500-04 (Cytochrome b5 domain) in chromosome 10. Chromosome 11 had TGACC and TTGAC occurring 6 times in 5-720 kb promoter sequences of the genes Os11t0529900-00 (Chalcone/stilbene synthase), Os12t0117100-01 (Alpha/beta hydrolase fold-1 domain) and Os12t0630800-00 (NAC140). Whereas, chromosome 12 has an occurrence of 7 times TGACC in 510-950 kb promoter sequences of Os12t0506500-01 (Calmodulin-binding protein). The list of all the rice genes with WRKY13 bm in their promoters is provided in Supplementary table 1.

### Classification of Genes using Gene Ontology and Enrichment

The list of genes was subjected to classify them based on the function of their resulting proteins using gene ontology (GO) analysis of molecular processes, biological processes and cellular components performed with PANTHER classification system software. 1,012 genes found from promoter analysis were subjected to functional classification viewed in selected graphic charts. These genes were classified based on the function of their proteins using gene ontology (GO) analysis. The genes ontologies terms are descriptions of the gene products, and the genes were classified into ‘biological process’, ‘molecular function’, ‘cellular component’, and ‘protein class’. Based on their GO terms these genes were grouped and graphically represented using Figure 4(i). GO Biological process include response to stimulus (4%), developmental process (3%), cellular process (38%), multicellular organismal process (2%), metabolic process (34%), Biological regulation (6%), cellular components organization or biogenesis (6%), and localization (7%). Cellular component organization or biogenesis involves 50% Cellular component organization and Cellular component biogenesis. Localization involves protein localization (83%) and transport (17%). Response to stimulus involves response to stress (67%) and response to abiotic stress (33%). The metabolic process involves the nucleobase-containing compound metabolic process (52%), lipid metabolic process (5%), cellular amino acid metabolic process (14%), protein metabolic process (24%) and carbohydrate metabolic process (5%). The protein metabolic process involves proteins responsible for proteolysis (20%), protein folding (20%) and cellular protein modification process (60%). Nucleobase-containing compound metabolic process involves the regulation of the nucleobase-containing compound metabolic process (27%), DNA metabolic process (9%), RNA metabolic process (55%) and purine nucleobase metabolic process (9%). GO Molecular function includes binding (35%), structural molecular activity (3%), catalytic activity (46%) and transporter activity (16%). Binding includes nucleic acid binding (30%) and protein binding (70%). The catalytic activity involves ligase activity (11.1%), oxidoreductase activity (11.1%), transferase activity (33.3%), enzyme regulator activity (5.6%), hydrolase activity (33.3%) and lyase activity (1.6%). GO Cellular components include membrane (22%), macromolecular complex (12%), cell part (38%) and organelle (28%). The membrane constitutes 67 % plasma membrane, 27% integral to membrane and 6% mitochondrial inner membrane. The cell part constitutes 36% plasma membrane proteins and 64% intracellular proteins. The macromolecular complex constitutes 17% protein complex and 83% ribonucleoprotein complex. Organelle constitutes mitochondrion (19%), cytoskeleton (12%), endoplasmic reticulum (12%) and Golgi apparatus (14%). The GO terms under the protein class include proteins groups transcription factor (4%), ligase (4%), nucleic acid binding (4%), receptor (4%), cytoskeletal protein (14%), storage protein (4%), signalling protein (7%), transporter (14%), membrane traffic protein (3%), hydrolase (11%), oxidoreductase (3%), cell adhesion molecule (3%), enzyme modulator (7%), lyase (4%) and transferase (14%),

**Figure 4.**
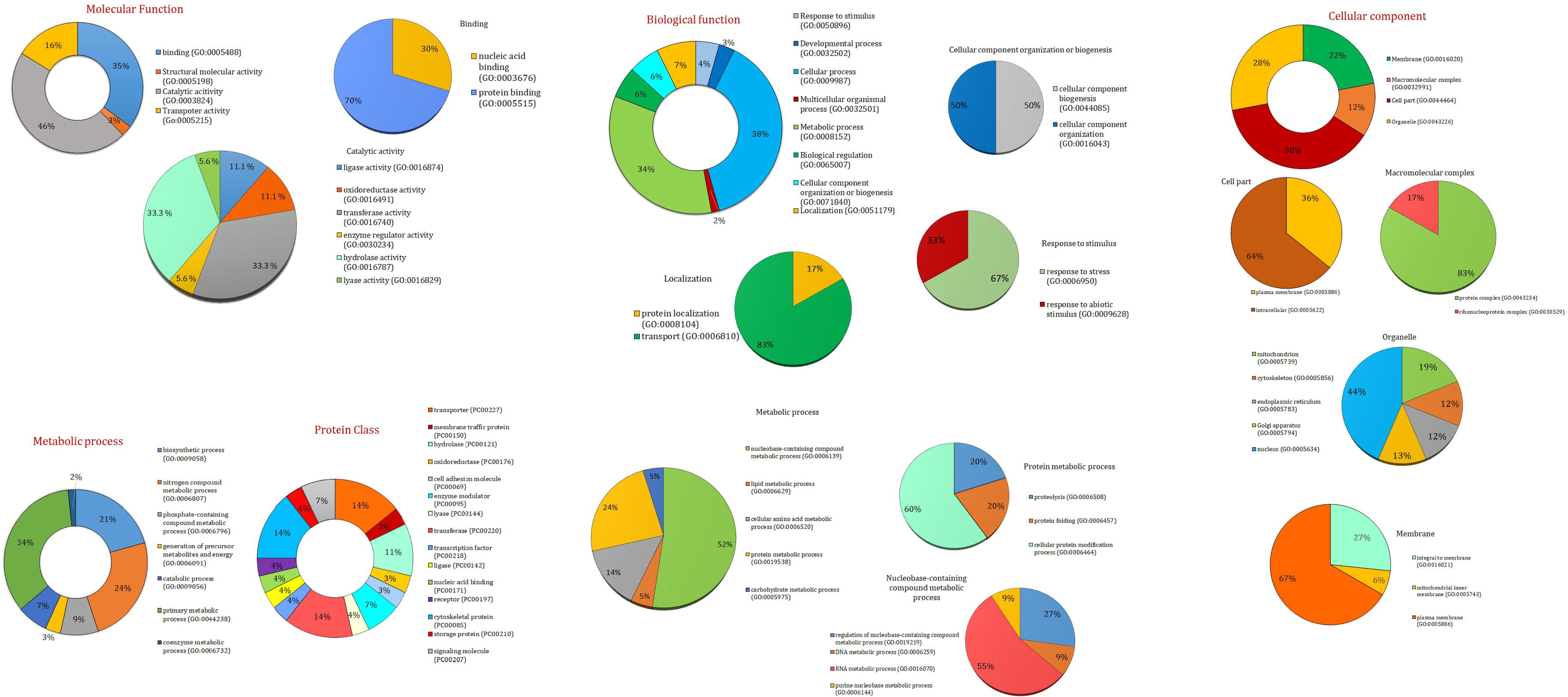
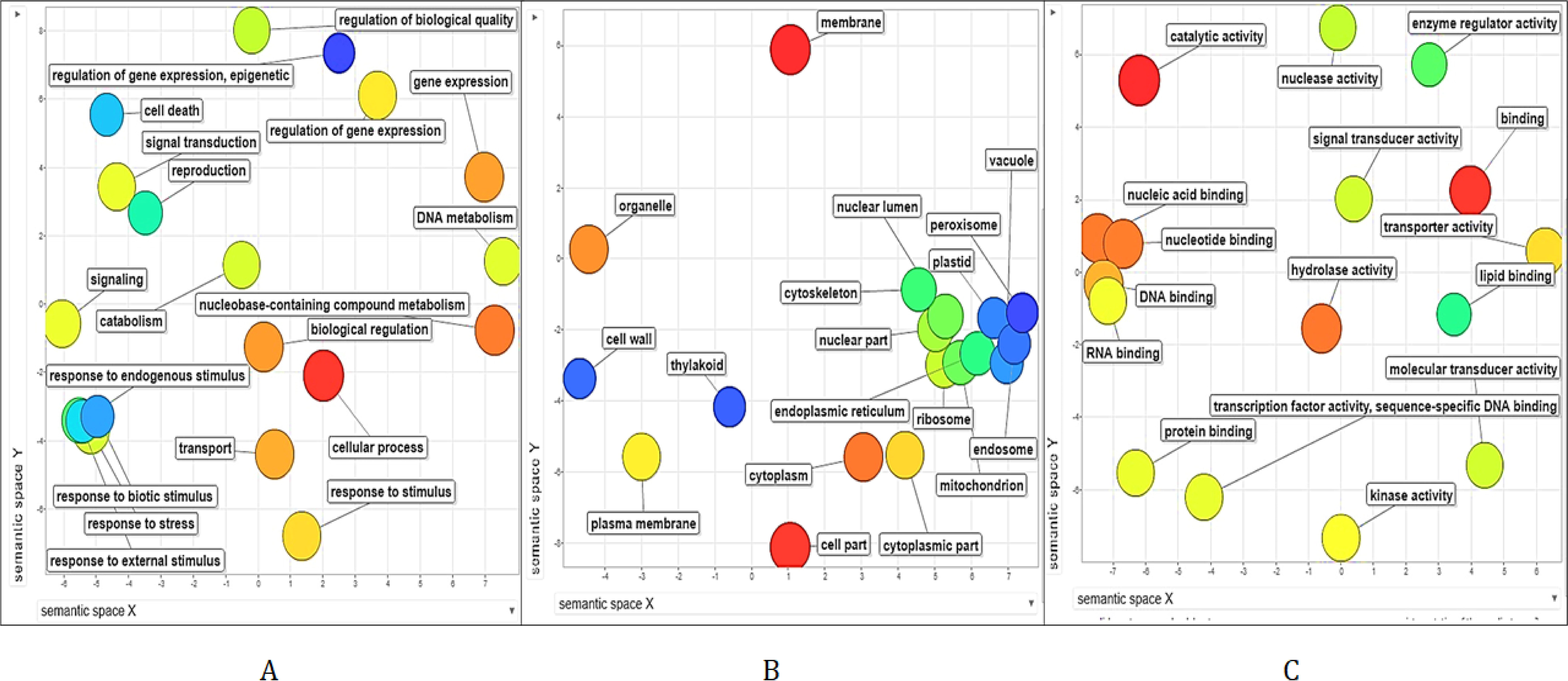
**(i)** Gene Classification using Gene Ontology Terms a. Molecular function, b. Catalytic activity, c. Binding, d. Biological function, e. Localization, f. Cellular component organization or biogenesis, g. Response to stimulus, h. metabolic process, i. Protein metabolic process, j. Nucleobase-containing compound metabolic process, k. Metabolic process, l. Protein class, m. Cellular component, n. Cell part, o. Macromolecular complex, p. Membrane, q. Organelle. (ii) Gene Ontology Enrichment Analysis exhibiting Functional and Significant GO Terms represented using Scattered Plot Diagram A. Biological process; B. Cellular processes; C. Molecular function

ReviGO analysis helped to summarize and remove redundant GO terms and uses statistical testing to determine enriched GO terms and assist in graph-based visualization and render semantic relationships in the input data. REVIGO also provides the same three separate aspects associated with the biological identity of the gene product like the PANTHER classification system software. While ReviGO provided classification and group them based on the evolutionary family dependent on their structural or functional similarities represented by Figure 4(ii). The input 1,012 genes reveal significant terms in the GO biological process that are a response to biotic stimulus (GO:0009607), response to stress (GO:0050896), response to external stimulus (GO:0009605), regulation of gene expression (GO:0010468) and signal transduction (GO:0007165). Whereas five significant terms in GO cellular location are nuclear part (GO:0044428), ribosome(GO:0005840), cytoplasm (GO:0005737), endoplasmic reticulum (GO:0005783) and thylakoid (GO:0009579). The five significant terms in GO molecular function are nucleic acid binding (GO:0003676), nucleotide binding(GO:0000166), DNA binding(GO:0003677), RNA binding(GO:0003723), transcription factor activity and sequence-specific DNA binding (GO:0003700). Further evaluation of GO molecular functions revealed nucleotide binding (20.185 %), nucleic acid binding (21.226 %), DNA binding (12.549 %), and transcription factor activity (sequence-specific DNA binding) (4.217 %).

### Genes with WRKY13 Binding Sites in their Promoter

The descriptions of the genes whose promoters had 4 or more WRKY13 bm was carried out using RAP-DB. The distribution of W-box cis-elements within the promoter region of rice disease resistance-related genes is represented by Figure 5i-5iii. The list of genes having WRKY13 bm in the promoter sequence was narrowed down to nine genes that have a known role in disease resistance (Table 3). These genes were manually searched to find their probable role in defense response against pathogens. The genes *ankyrin repeat*, *aminotransferase*, *PR*2, *WRKY*12, *PR*5, β-*Glycosidase* 14, germin 8-7 (*ger*6), *PR*1b, and shikimate biosynthesis protein (*aroDE*) enumerated in the Table 3 have existing reports of role in disease resistance. The nine genes resulting from the promoter analysis are found to be located in different chromosomes (4, 8, 1, 11, 7, 6, and 9). The *β-Glycosidase*14 gene located in chromosome 4 contained TGACT and TTGACT bm, whereas, Germin 8-7 (*ger*6) located in chromosome 6 contained TGACT, TTGAC, and TTGACT bm. TTGAC bm was found in the genes *WRKY*12, *Ankyrin*, *PR*1b, *PR*5, *aminotransferase* and shikimate dehydrogenase protein (*aroDE*) located in chromosomes 1, 11, 7, 6, 9 and 1 respectively. *PR*2 located in chromosome 1 contained TGACT bm.

**Figure 5.**
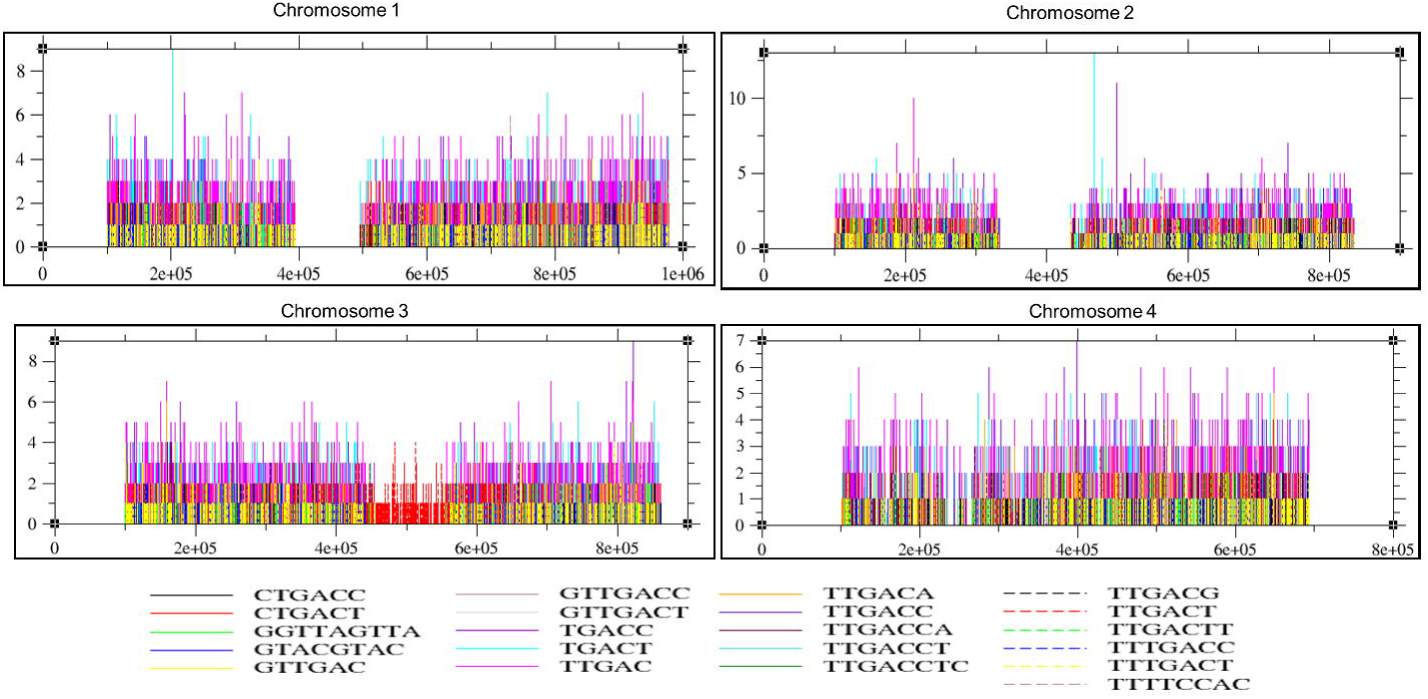
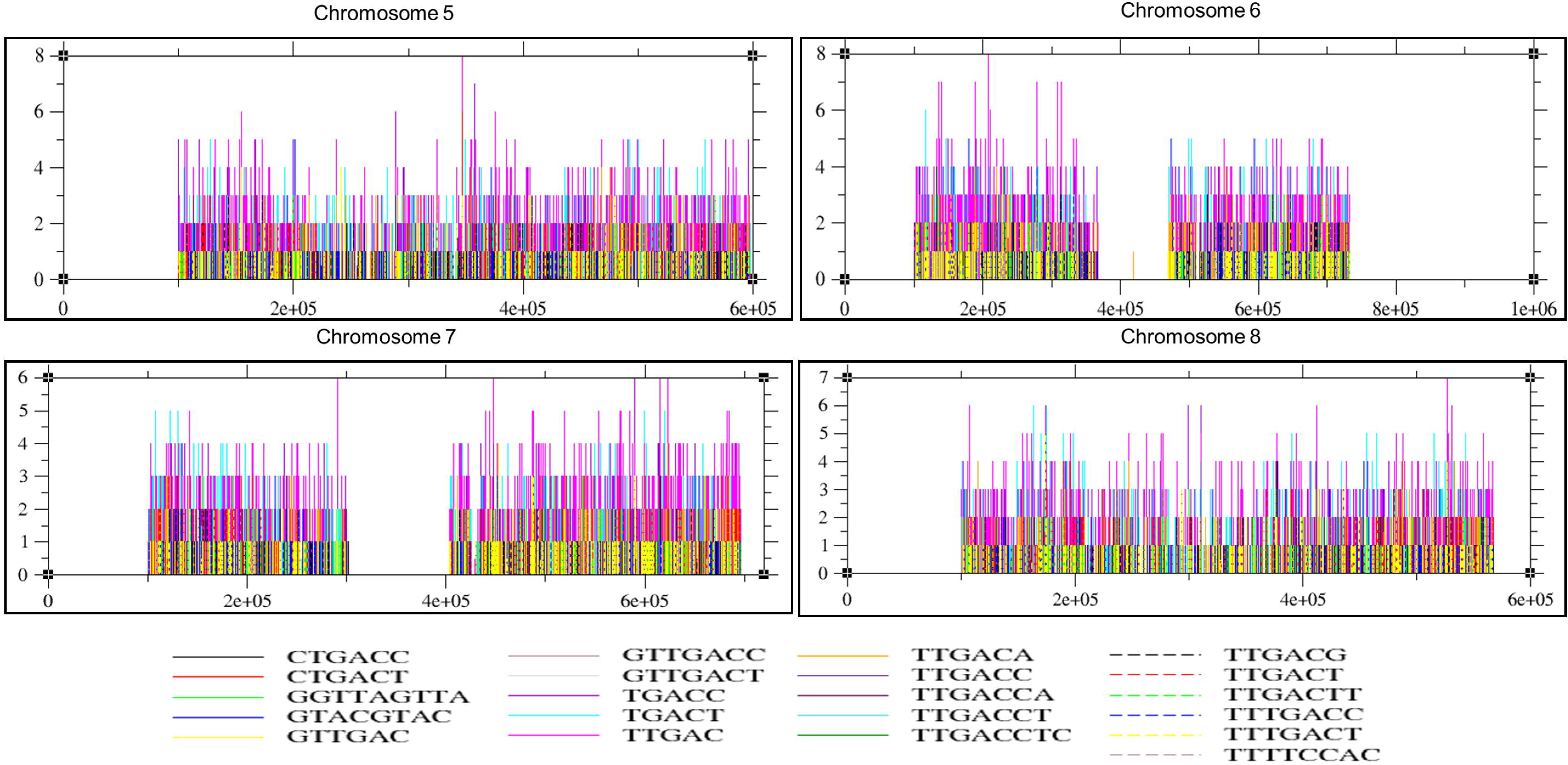
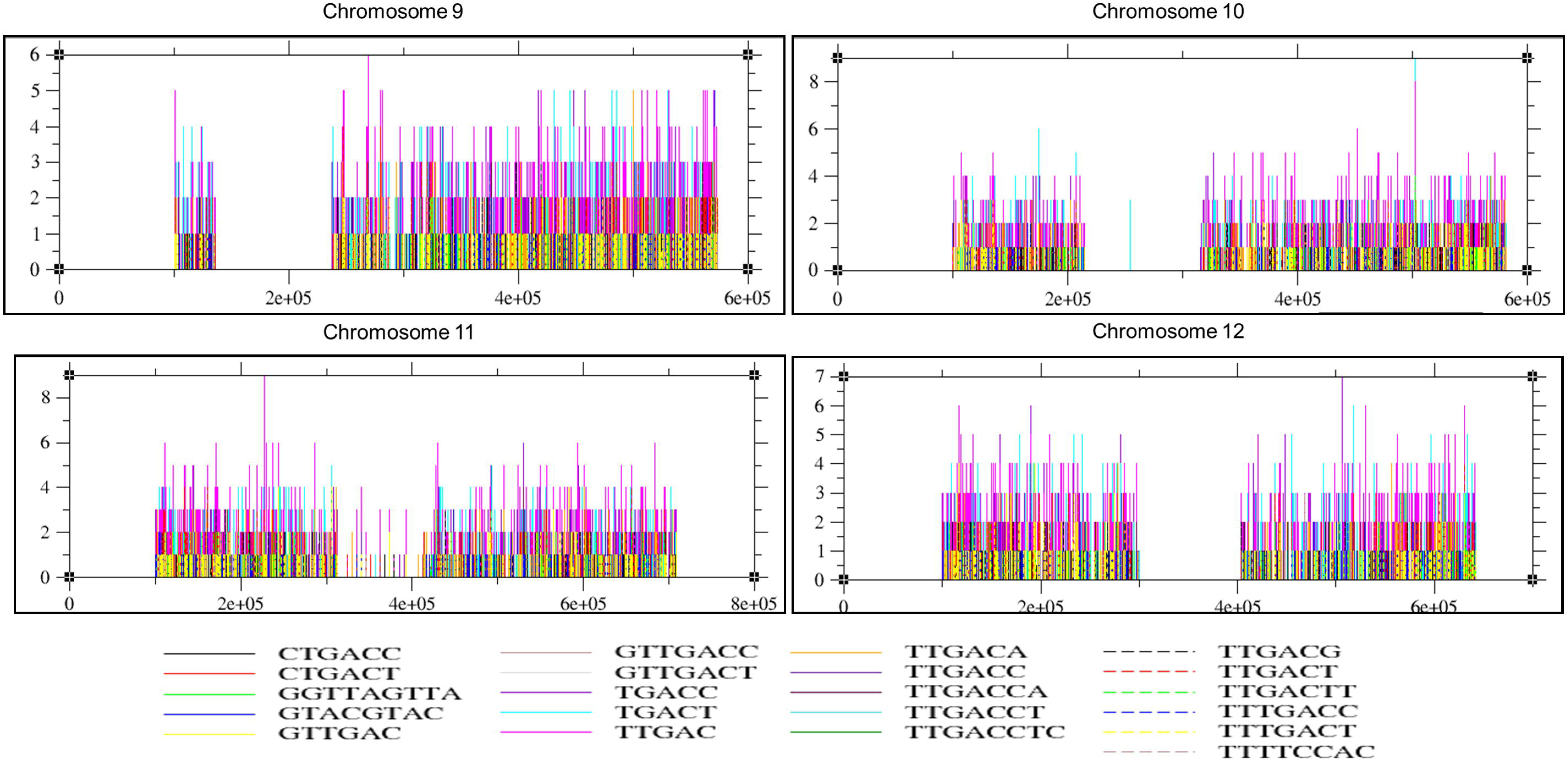
Graphical Representation of Occurrence of WRKY13 TF bm in Rice Chromosomes X-axis—combined 1 kb promoter sequence of all the genes in a given chromosome. Y axis— number of occurrences of motifs.

**Table 3.** Disease Resistance-Related Genes found using Genome-Wide Computational Analysis

### W-Box *Cis*-Acting Regulatory Elements in Promoter Sequences

The *cis*-acting regulatory elements that may regulate disease resistance-related genes were obtained from the PLACE database using the FASTA format of the promoter sequences of all nine sequences as an input. The analysis provides the name of each factor or site names, signal sequence (motif), and the location of the motif if in sense or antisense strand and was tabulated using Table 4. The promoter sequences were used as input to obtain the visualization of the position of the cis-elements in the promoter sequences from plantPAN. The organism ‘*Oryza sativa*’ was selected and the motifs were entered. The distribution of W-box cis-elements within the promoter region of rice disease-related genes is represented using Figure 6. *WRKY*13 was included due to reports of auto-regulation, while TIFY9 was previously reported by the group to interact closely with WRKY13 to mediate an interaction with the WRKY core network that interacts with MAPK cascade genes. A*nkyrin repeat* revealed 11 signal sequences, *aminotransferase* revealed 8 signal sequences, *PR*2 revealed 20 signal sequences, *WRKY*12 revealed 23 signal sequences, *PR*5 revealed 9 signal sequences, β-*Glycosidase* 14 revealed 12 signal sequences, germin 8-7 (*ger*6) revealed 33 signal sequences, *PR*1b revealed 21 signal sequences, and shikimate biosynthesis protein (*aroDE*) revealed eight signal sequences. *WRKY*13 revealed three signal sequences, while *TIFY*9 revealed one signal sequence. The W-box factors site corresponding to their signal sequences were WBBOXPCWRKY1 - TTTGACY, WBOXATNPR1 - TTGAC, WBOXHVISO1 - TGACT, WBOXNTERF3 - TGACY and WBOXNTCHN48 - CTGACY. The site factor WBBOXPCWRKY1 was not present in the promoter regions of *Ankyrin* and *WRKY*13, whereas WBOXNTCHN48 was only found in the promoter region of *PR*1b and *PR*2. All five W-box factor sites were found in the promoter regions of *PR*1b and *PR*2.

**Figure 6.**
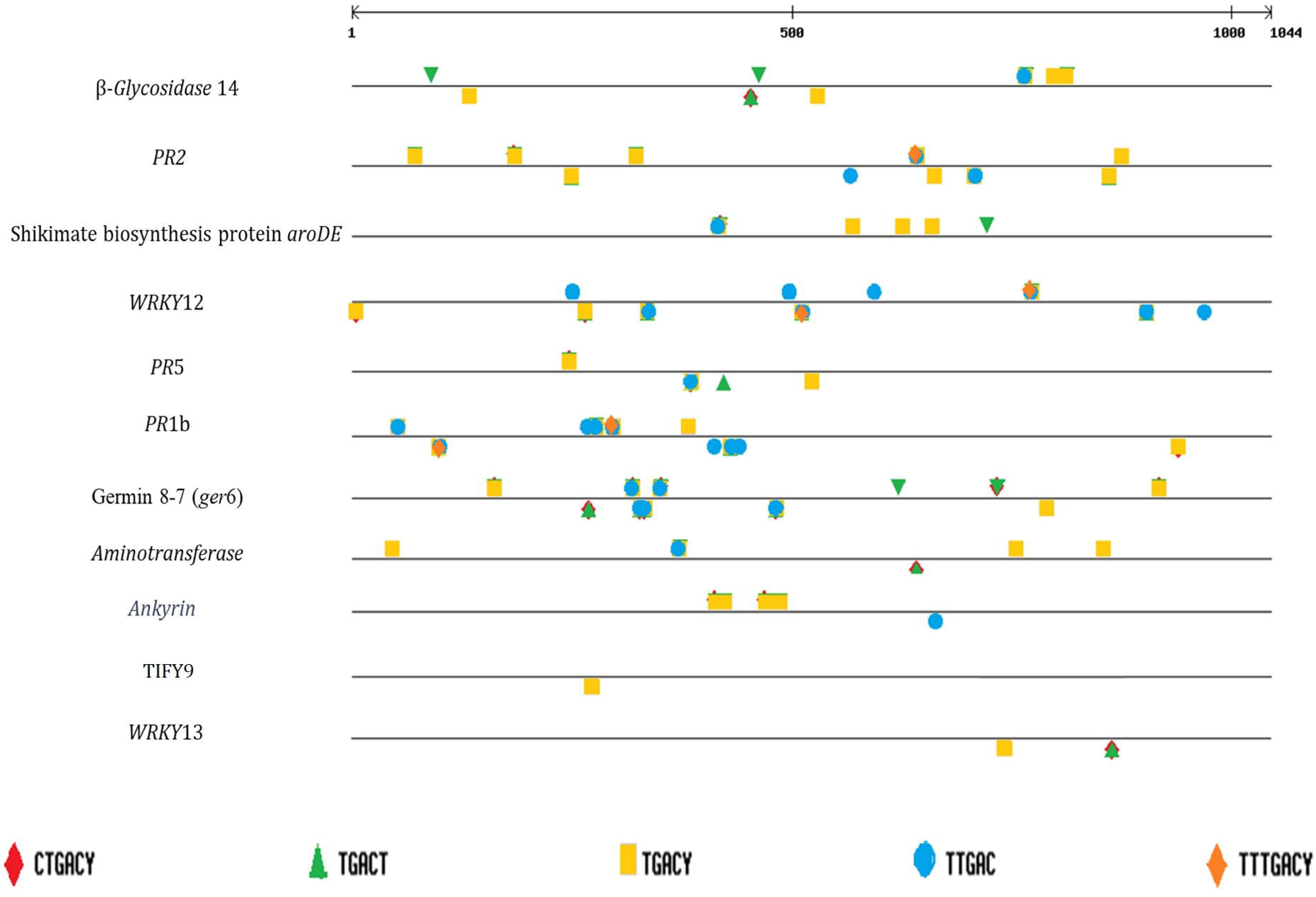
Distribution of W-box Cis-Elements within Promoter Region of Rice Disease Resistance-Related Genes

**Table 4.** Cis-Elements of selected Disease Resistance Related Genes

### WRKY13 Binds to the Promoters of the Selected Genes

The protein structure was predicted using Swiss-model which built it using the target-template alignment using inbuilt ProMod3 (Version 1.0.2.). The tool identified a closely related template named 1wj2.1.A and the template alignment reported a sequence similarity of 0.50 and an identity of 55.17. The predicted structure of WRKY13 binding with DNA sequences is represented in Figure 7i-7xi. The promoter binding site pattern of WRKY13 was according to the respective motifs found in the promoter sequences of these genes since no common motif was found. Promoter-WRKY13 docking was carried out using Autodock and the docked complex with the best score was selected for further analysis. The interaction energies (binding energy, Van der Waals energy, and electrostatic energy) were estimated. The individual energy components of the best docking mode for each promoter structure were tabulated in Table 5. As provided by the AutoDock tool, among all these molecules, WRKY13 showed affinity to these protein molecules in the following order: *Aminotransferase*, shikimate biosynthesis protein (*aroDE*), *ankyrin, PR*2, *WRKY13, WRKY12, TIFY*9, *β-Glycosidase*14, Germin 8-7 (*ger*6), *PR*1b and *PR*5. The interacting residues predicted by NUCPLOT for all the 11 docked molecules were predicted. The predictions were documented for all the complexes (b of Figure 7(i)-7(xi). Transcription factor AA interacts with the target DNA bases (5’, 3’ and one/two in the between) and with ribose and phosphate groups. These interactions were predicted and each provided unit was recorded.

**Figure 7.**
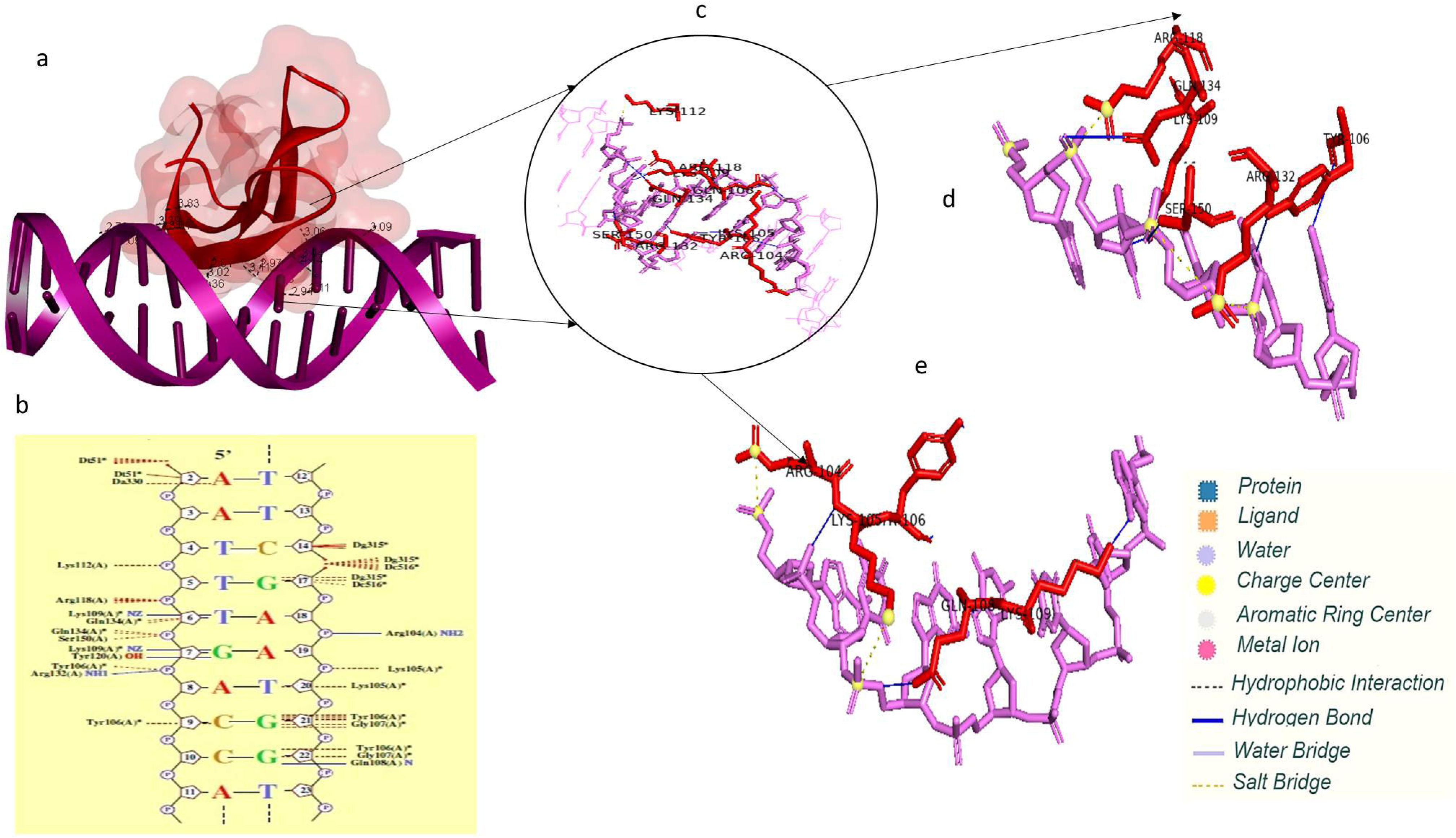
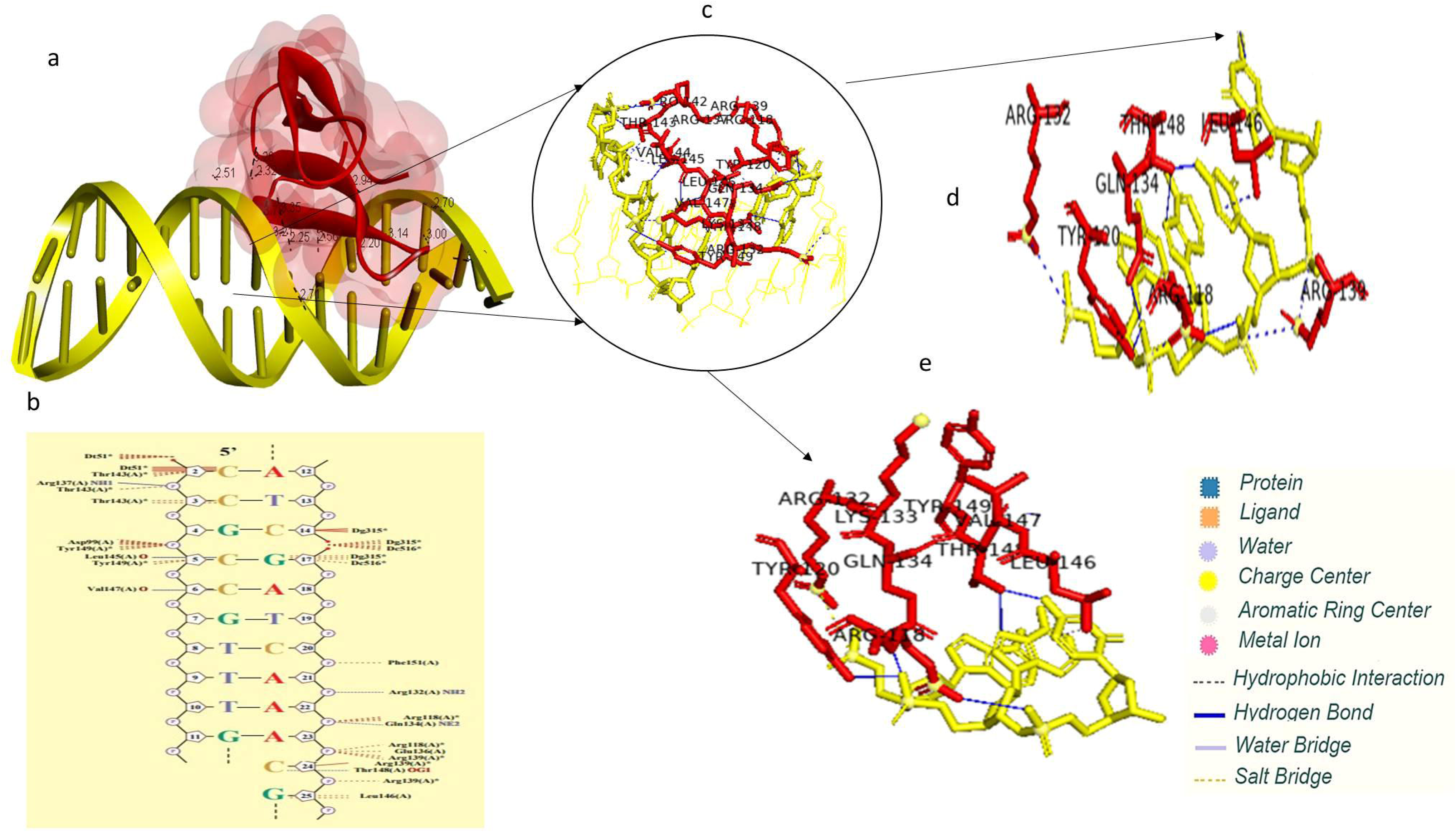
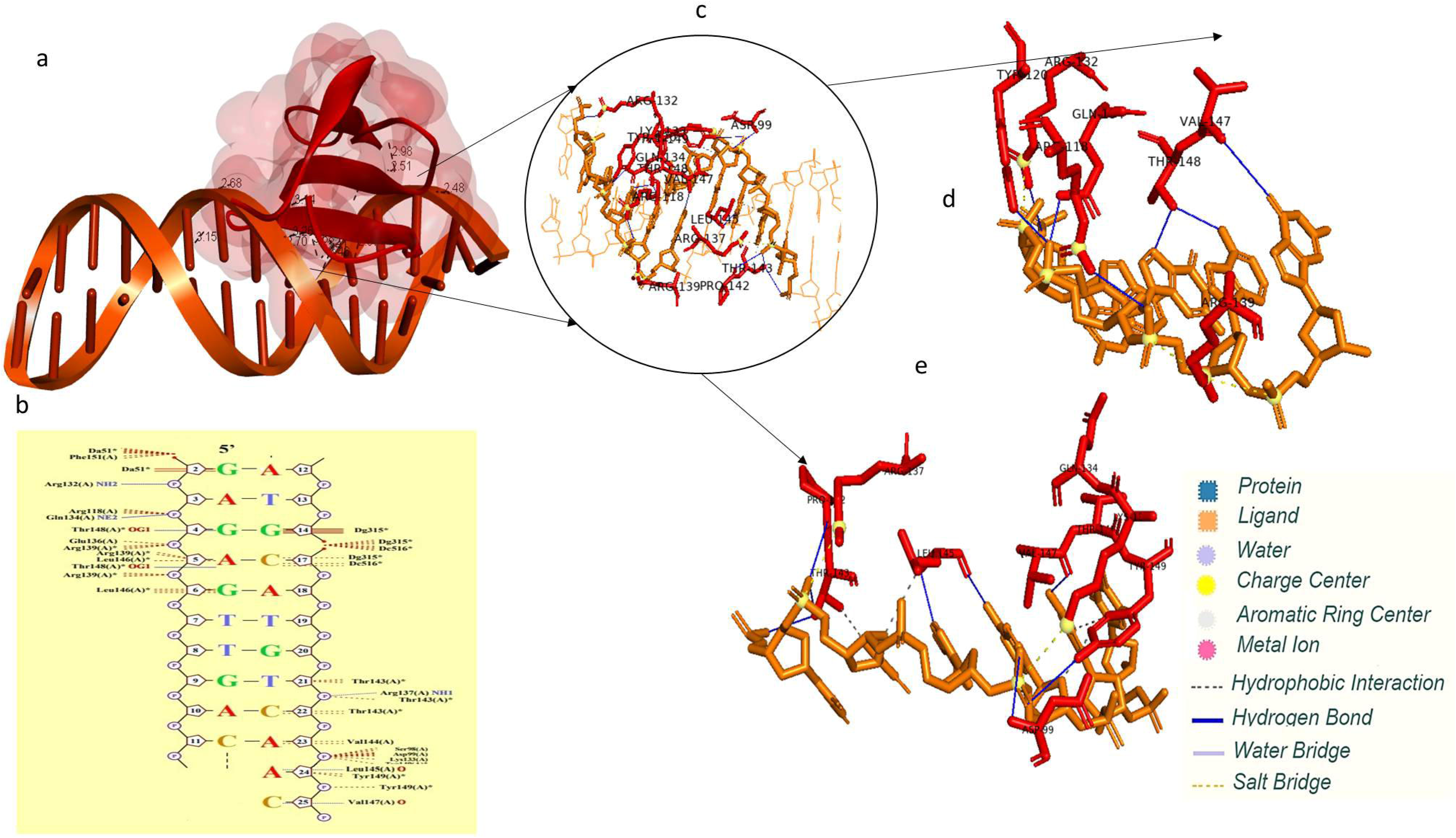
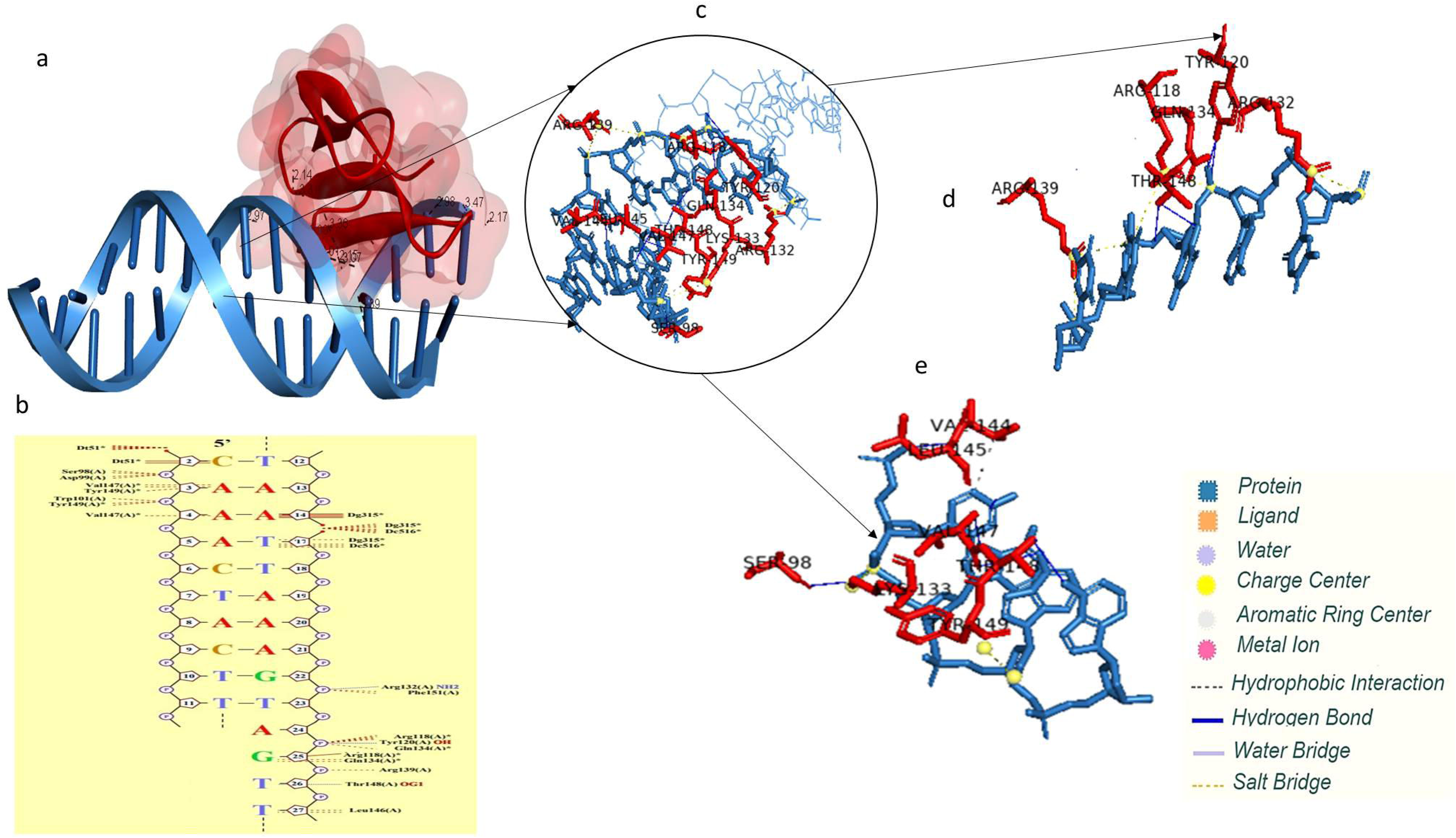
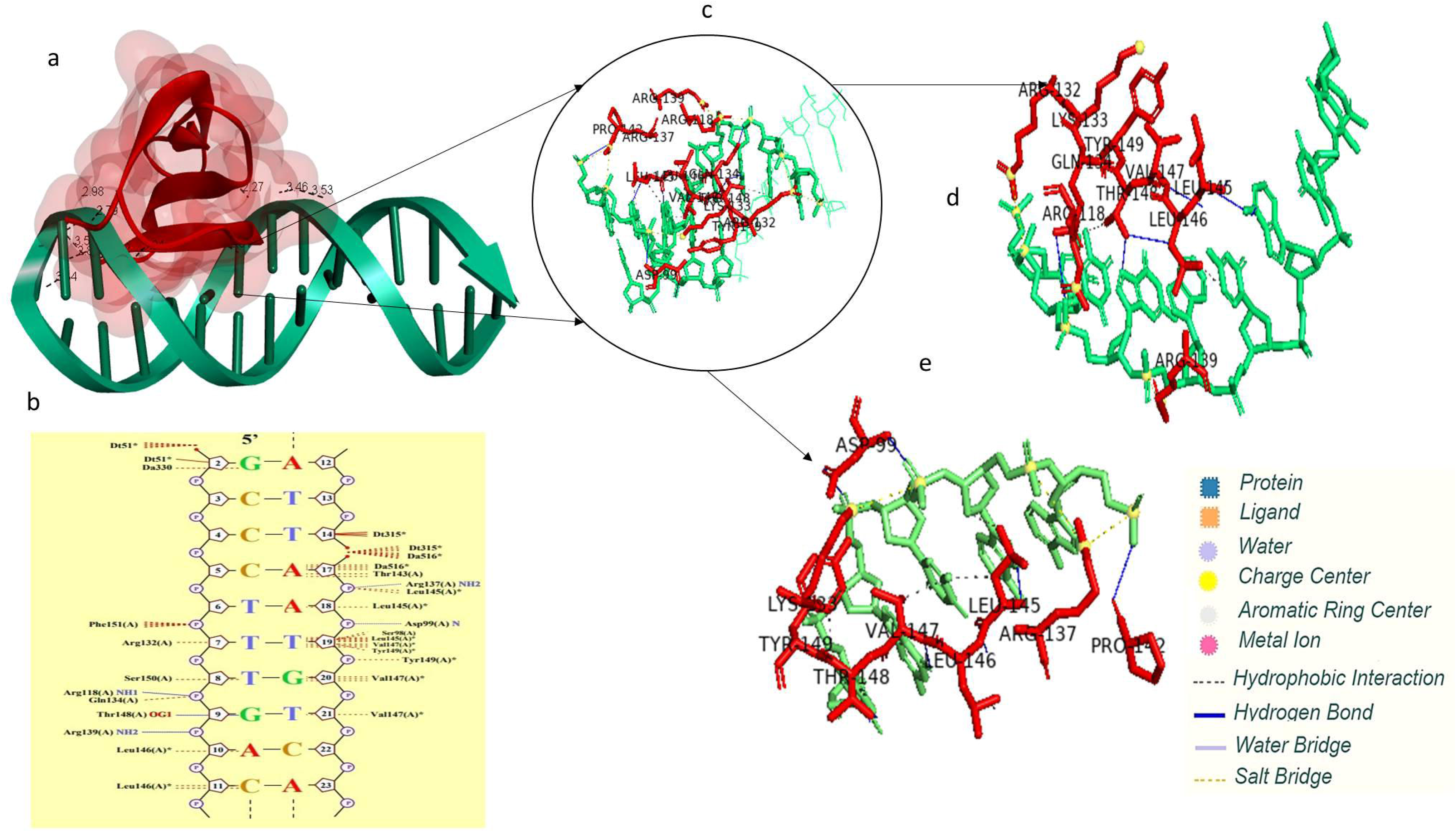
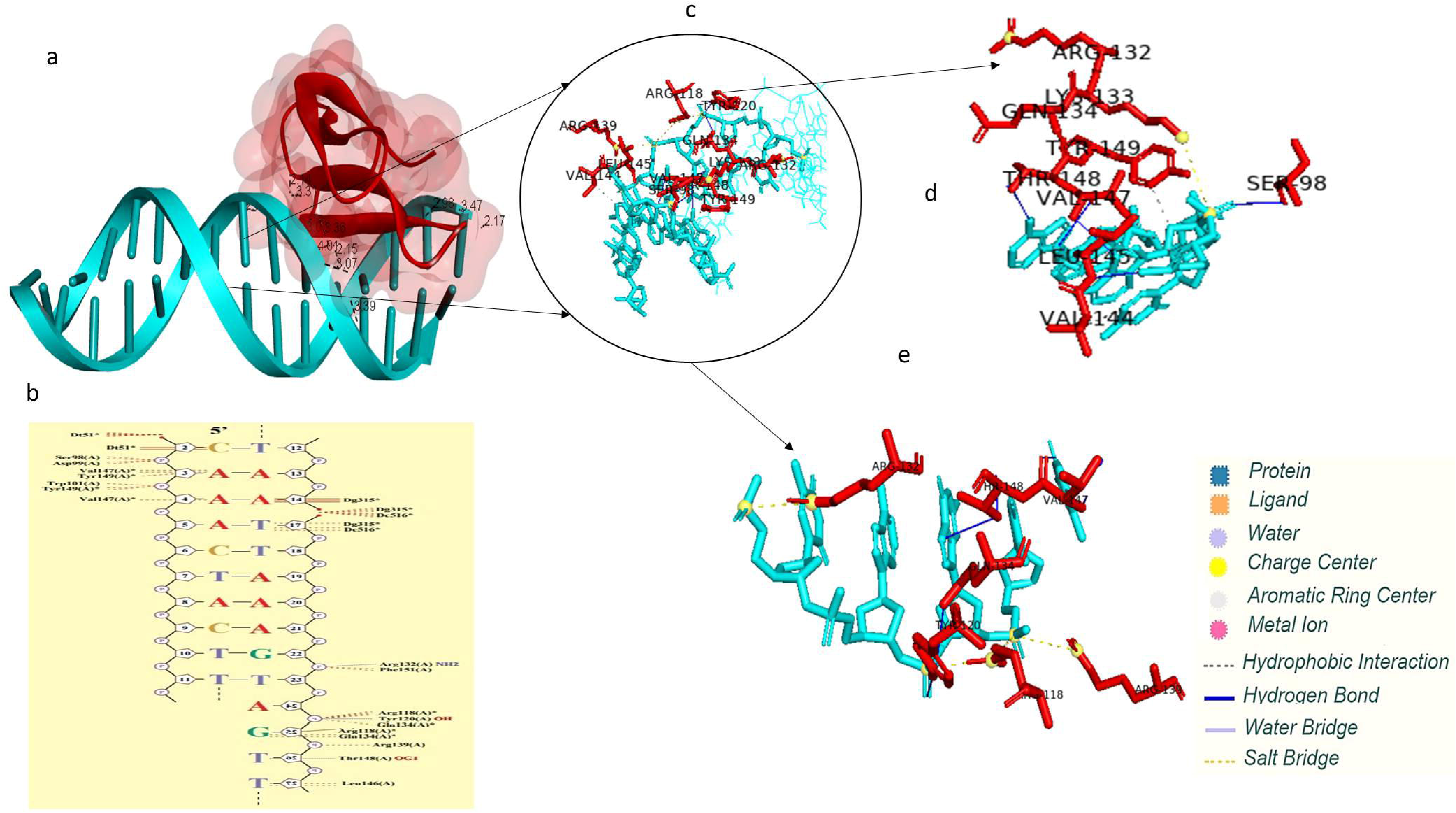
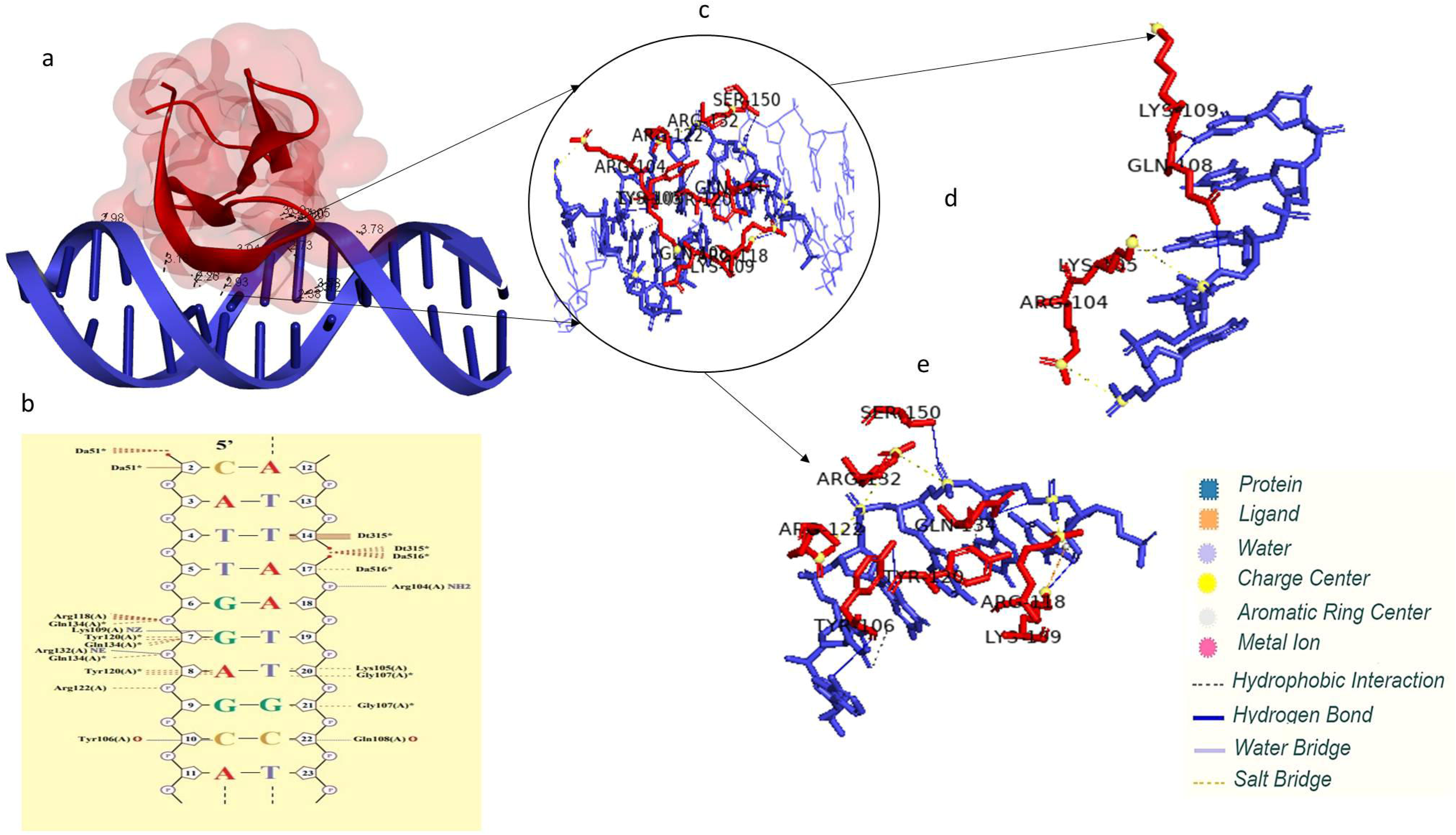
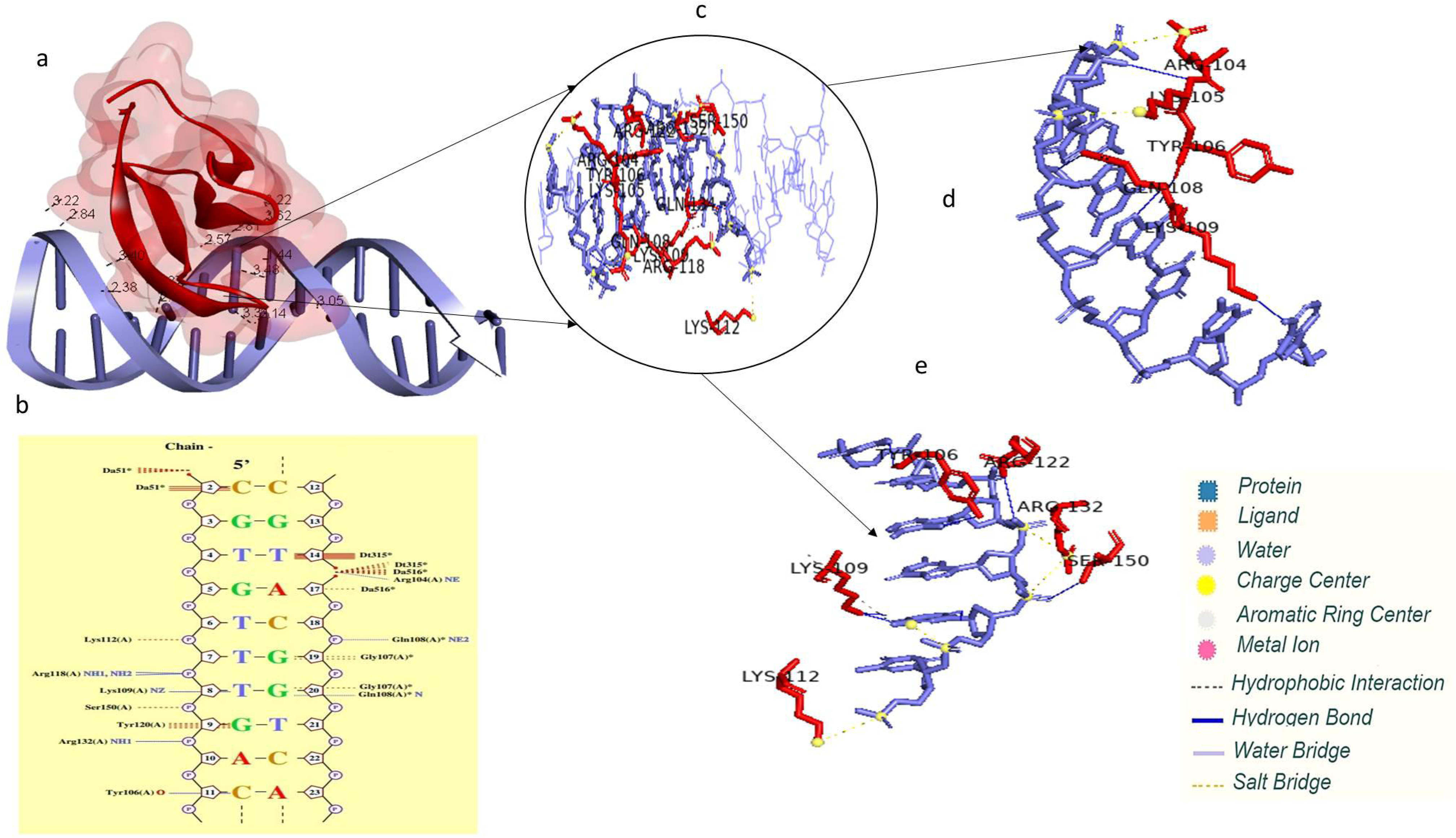
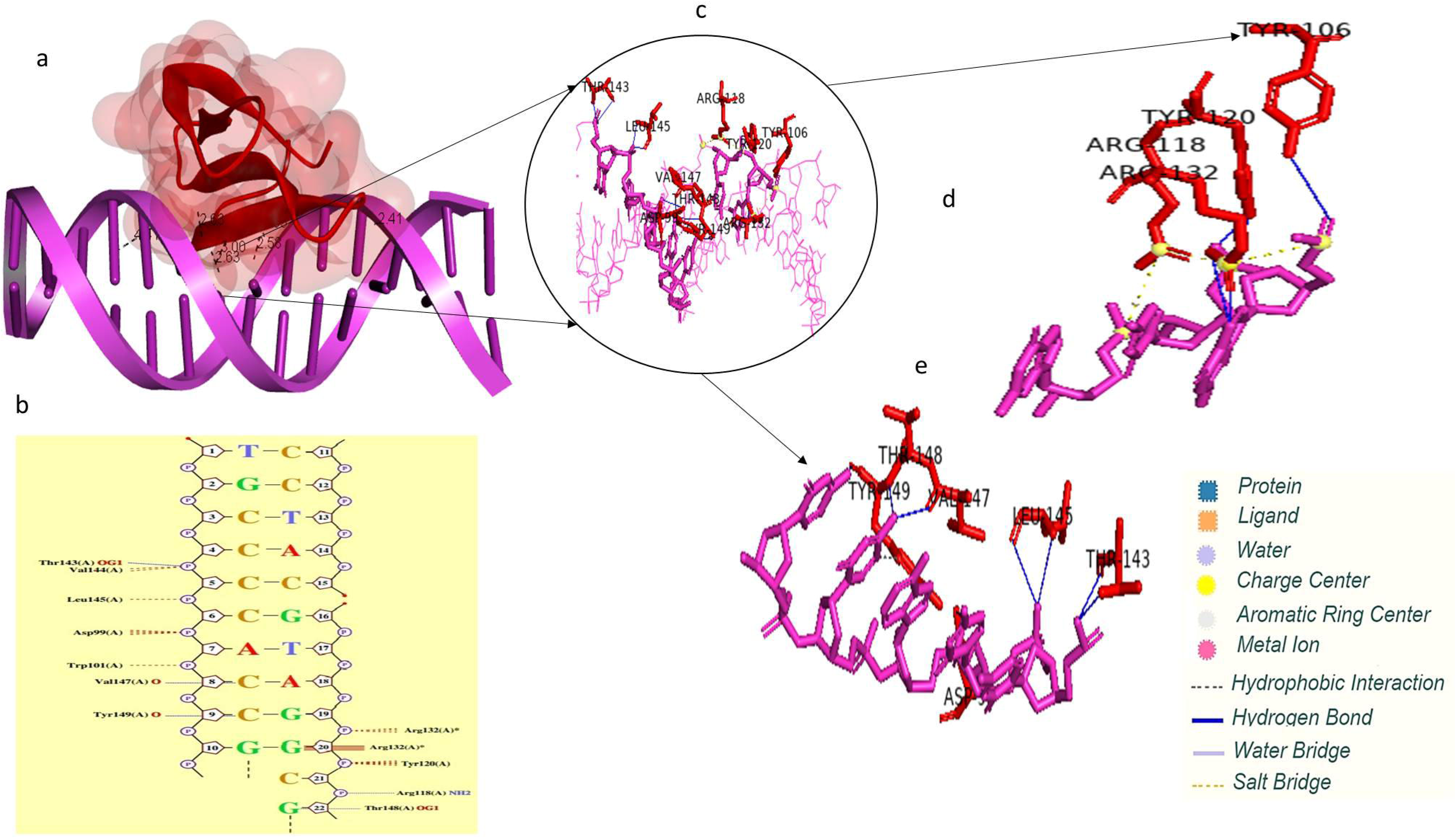
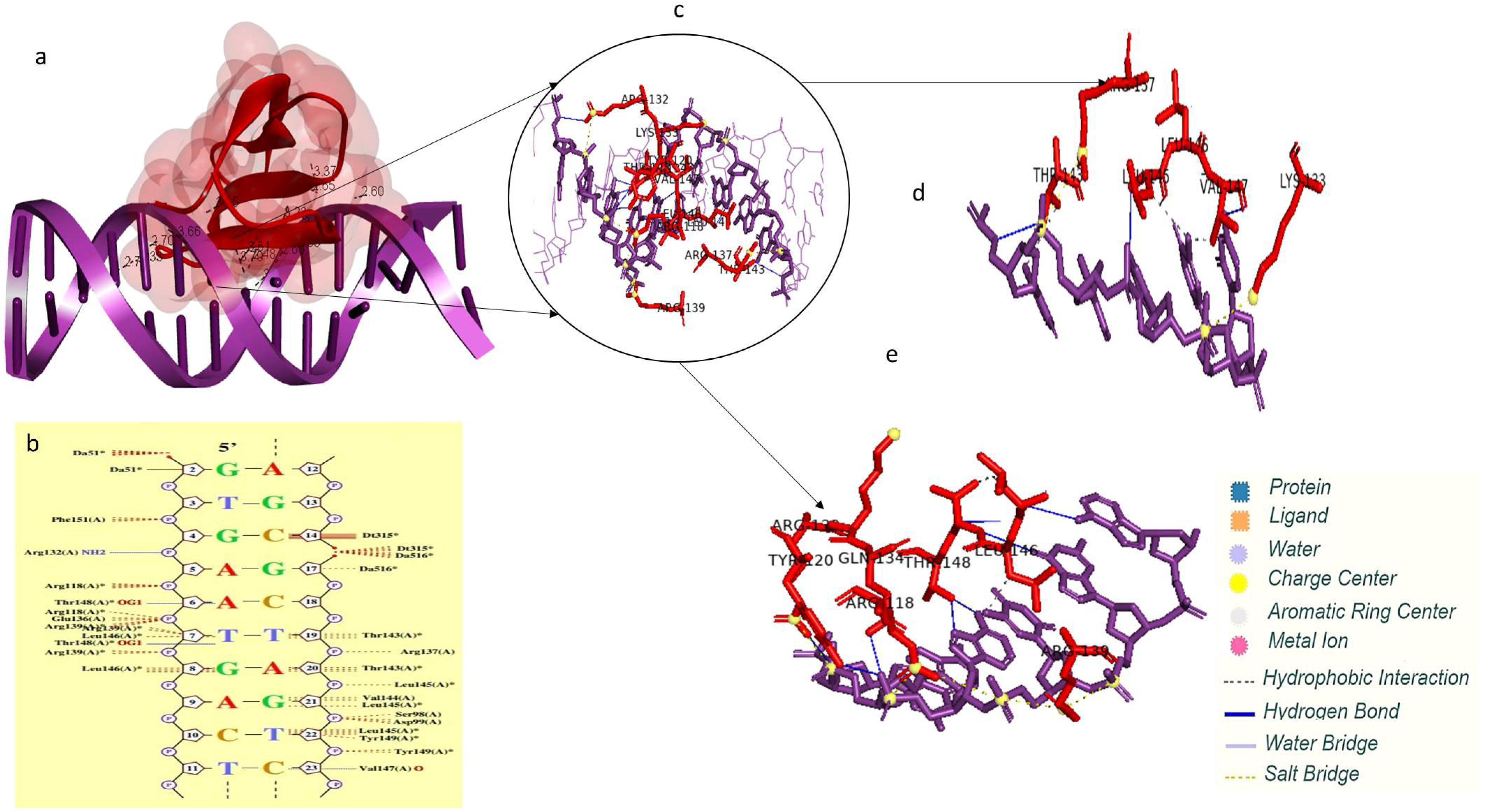
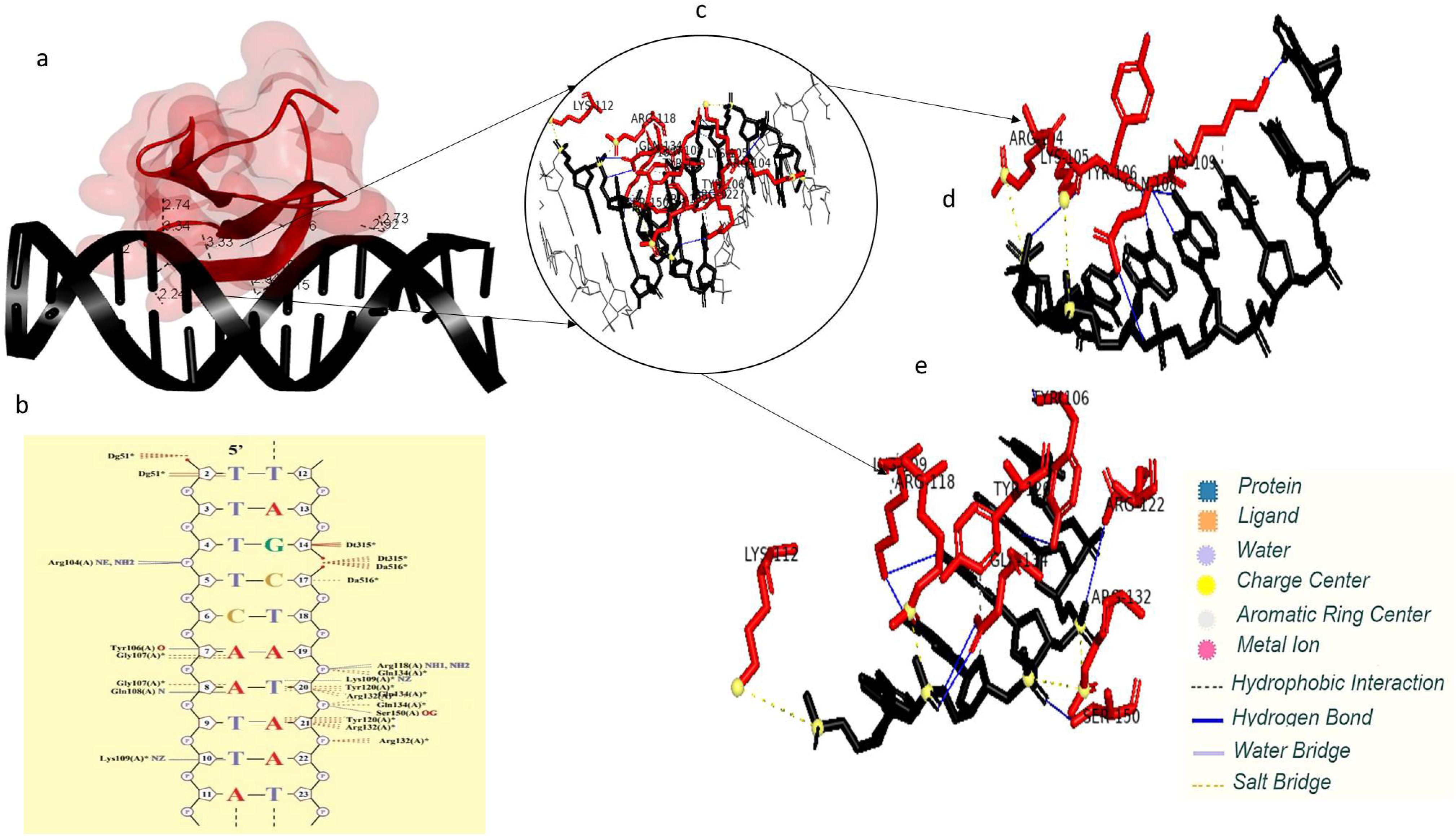
Predicted Structures of Binding of WRKY13 with Promoter Sequences of Selected Rice Disease Resistance-Related Genes and interacting residues a. WRKY13-DNA complex; b. predicted interacting residues from Nucplot; c-e. interacting residues i- *β-Glycosidase*14-WRKY13; ii- *ger*6-WRKY13; iii- *WRKY*12-WRKY13; iv- *Ankyrin repeat*- WRKY13; v- *PR*1b-WRKY13; vi- *PR*5-WRKY13; vii- *PR*2-WRKY13; viii- *Aminotransferase*- WRKY13; ix- *aro*DE-WRKY13; x- *TIFY*9-WRKY13; xi- *WRKY*13-WRKY13

**Table 5.** Interaction energies of Protein–DNA Complex of WRKY13 TF with Promoters of Selected Rice Genes

The interactions in DNA-WRKY13 complexes were also analyzed using PLIP server that filters the interacting residues. The interaction of WRKY13 TF protein binding with the promoter sequences of selected disease resistance genes was studied using the PLIP server (Figure 7(i)-7(xi). It provides other details such as interacting residues and distance between the hydrogen and acceptor, type of bonds formed, donor and acceptor, the distance between interaction in carbon atoms in case of hydrophobic bonds and distance between centres of charge in case of salt bridges. It also provides the IDs of donor and acceptor atoms. The PLIP server provides the details of hydrogen bonds and hydrophobic bonds. It also gives details of the salt bridge formed between the residues. The interacting residues of each predicted structure of binding of WRKY13 with the promoter sequences of selected disease resistance-related genes are tabulated using Supplementary Tables 2-12. In the case of WRKY13-*aminotransferase* interaction, the interacting residues were glycine (GLN108A) and Lysine (LYS109A). Whereas the residues forming salt bridges with the phosphate groups were lysine (LYS105A and LYS109A), Tyrosine (TYR106A), serine (SER150A) and glutamine (GLN108A, GLN134A). The residues arginine (ARG104A, ARG118A, ARG132A) and lysine (LYS105A, LYS112A) were involved in hydrophobic interactions. Whereas, WRKY13-*ankyrin* interaction revealed threonine (THR143A), valine (VAL144A), leucine (LEU145A, LEU146A) and Tyrosine (TYR149A) to be the interacting residues. Arginine (ARG118A), tyrosine (TYR120A, TYR149A), glutamine (GLN134A), proline (PRO142A), threonine (THR143A, THR148A), leucine (LEU145A), and Valine (VAL147A) formed a salt bridge with the phosphate group. The residues arginine (ARG118A, ARG132A, ARG137A, ARG139A) and lysine (LYS133A) were involved in hydrophobic interactions.

The interacting residues in the case of WRKY13-Shikimate biosynthesis protein (*aroDE*) interaction were tyrosine (TYR106A), glutamine (GLN108A) and Lysine (LYS109A). Residues interacting in hydrophobic interactions were Lysine (LYS105A, LYS109A), tyrosine (TYR106A, TYR108A), glutamine (GLN108A, GLN134A), Arginine (ARG122A), and (SER150A). Arginine (ARG104A, ARG118A, ARG132A), and Lysine (LYS105A, LYS112A) formed a salt bridge with the phosphate groups. Whereas, in the case of WRKY13-*WRKY*13 interaction, the interacting residues were glutamine (GLN108A), tyrosine (TYR106A), lysine (LYS109A), tyrosine (TYR120A). Arginine (ARG104A, ARG118A, ARG132A) and Lysine (LYS105A, LYS112A) formed a salt bridge with the phosphate groups. The residues glutamine (GLN108A, GLN134A), Lysine (LYS105A, LYS109A), tyrosine (TYR106A), Arginine (ARG122A) and serine (SER150A) were involved in hydrophobic interactions. For WRKY13-*WRKY*12 interaction revealed the interacting residues threonine (THR148A) and tyrosine (TYR149A). Hydrophobic interaction was seen between aspartic acid (ASP99A), tyrosine (TYR106A, TYR120A), arginine (ARG132A), threonine (THR143A), leucine (LEU145A), valine (VAL147A) and tyrosine (TYR149A). The resides Arginine (ARG118A, ARG132A) to be forming salt bridges with the phosphate group. In the case of WRKY13-*PR*2, the interacting residues of the protein were valine (VAL144A, VAL147A) and tyrosine (TYR149A). The residues serine (SER98A), tyrosine (TYR120A), glutamine (GLN134A), leucine (LEU145A), Valine (VAL147A), and threonine (THR148A) were involved in hydrophobic interaction. Arginine (ARG118A, ARG132A, ARG139A) and Lysine (LYS133A) formed a salt bridge with the phosphate group. In the case of WRKY13-*β-Glycosidase* 14, the interacting residues were Valine (VAL144A) and tyrosine (TYR149A). Residues involved in the hydrophobic interaction were serine (SER98A), tyrosine (TYR120A), glutamine (GLN134A), leucine (LEU145A), Valine (VAL147A) and threonine (THR148A). Arginine (ARG118A, ARG132A, ARG139A) and lysine (LYS133A) formed a salt bridge with the phosphate groups.

In the case of WRKY13-Germin 8-7 (*ger*6), the interacting residues of the protein were leucine (LEU145A), and tyrosine (TYR143A, TYR149A). The residues aspartic acid (ASP99A), proline (PRO6A, PRO142A), arginine (ARG118A, ARG132A), tyrosine (TYR120A, TYR149A), glutamine (GLN134A) leucine (LEU54A, LEU 55A, LEU145A), threonine (THR143A, THR148A), valine (VAL147A) was involved in hydrophobic interaction. Lysine (LYS133A) and arginine (ARG118A, ARG132A, ARG137A, ARG139A) formed a salt bridge with the phosphate group. For the WRK13-*PR*1b interaction, the interacting residues were leucine (LEU145A, LEU146A), valine (VAL147A), threonine (THR148A) and tyrosine (TYR149A). Aspartic acid (ASP99A), glutamine (GLN134A), proline (PRO142A), leucine (LEU145A), (VAL147A), and threonine (THR148A) were the residues that formed a salt bridge with phosphate groups. Arginine (ARG118A, ARG132A, ARG137A, ARG139A), and lysine LYS133A were involved in hydrophobic interactions. In the case of WRKY13-*TIFY*9 interaction, the interacting residues were threonine (THR143A), Leucine (LEU145A), and Valine (VAL147A). Arginine (ARG118A, ARG132A, ARG137A, ARG139A) and Lysine (LYS133A) formed a salt bridge with the phosphate groups. The residues glutamine (GLN134A), tyrosine (TYR120A), Arginine (ARG132A), threonine (THR143A, THR148A), Leucine (LEU145A), and Valine (VAL147a) were involved in hydrophobic interactions. For the WRKY13-*PR*5 interaction, the interacting residues were lysine (LYS105A) and tyrosine (TYR120A). The residues tyrosine (TYR106A), glycine (GLN108A, GLN134A), Lysine (LYS109A) and serein (SER150A) were involved in hydrophobic interactions. The residues lysine (LYS105A) and arginine (ARG104A, ARG118A, ARG122A, ARG132A) formed salt bridges with phosphate groups. Lysine (LYS109A) formed π-cation interactions with ligand atoms. It is noted that the salt bridges in all the interactions (usually involving arginine) were formed due to the electrostatic energy caused due to the interaction of the amino acid with the phosphate groups.

### Root-Mean-Square Deviation of Atomic Positions (RMSD)

RMSD values were estimated to understand the effect of the interaction of gene-WRKY13 complexes. RMSD is the scalar distance measurement between the atoms from their initial conformation to the final conformation, providing the estimation of the stability of the system. Here, RMSD was plotted for all the studied genes with the WRKY13 protein complex. It can be observed that the stable trajectories were attained quickly from the beginning of the simulations for all the complexes and were found to be fluctuating between 0.1nm to 0.35nm except for aroDE. Figures for RMSD of each of the interactions are attached in supplementary figures 1.1-1.11. The RMSD of all the interactions is provided in Figure 8. The RMSD values of all interactions are provided in Table 6 that revealed *aroDE*-WRKY13 (0.358) has the higher average RMSD, swiftly followed by *TIFY9*-WRKY13 (0.329), *PR*1b-WRKY13 (0.287), *WRKY*12-WRKY13 (0.273), *PR*5-WRKY13 (0.251), *ger*6-WRKY13 (0.238), *PR*2-WRKY13 (0.236), *WRKY*13-WRKY13 (0.245), *ankyrin*-WRKY13 (0.221) and β-*glucosidase*-WRKY13 (0.200) and *Aminotransferase*-WRKY13 (0.198). these observations suggest that out of compared complexes, *Aminotransferase*-WRKY13 forms the most stable confirmation, with the highest structural stability, while the *aroDE*-WRKY13 complex has the least structural stability.

**Figure 8.**
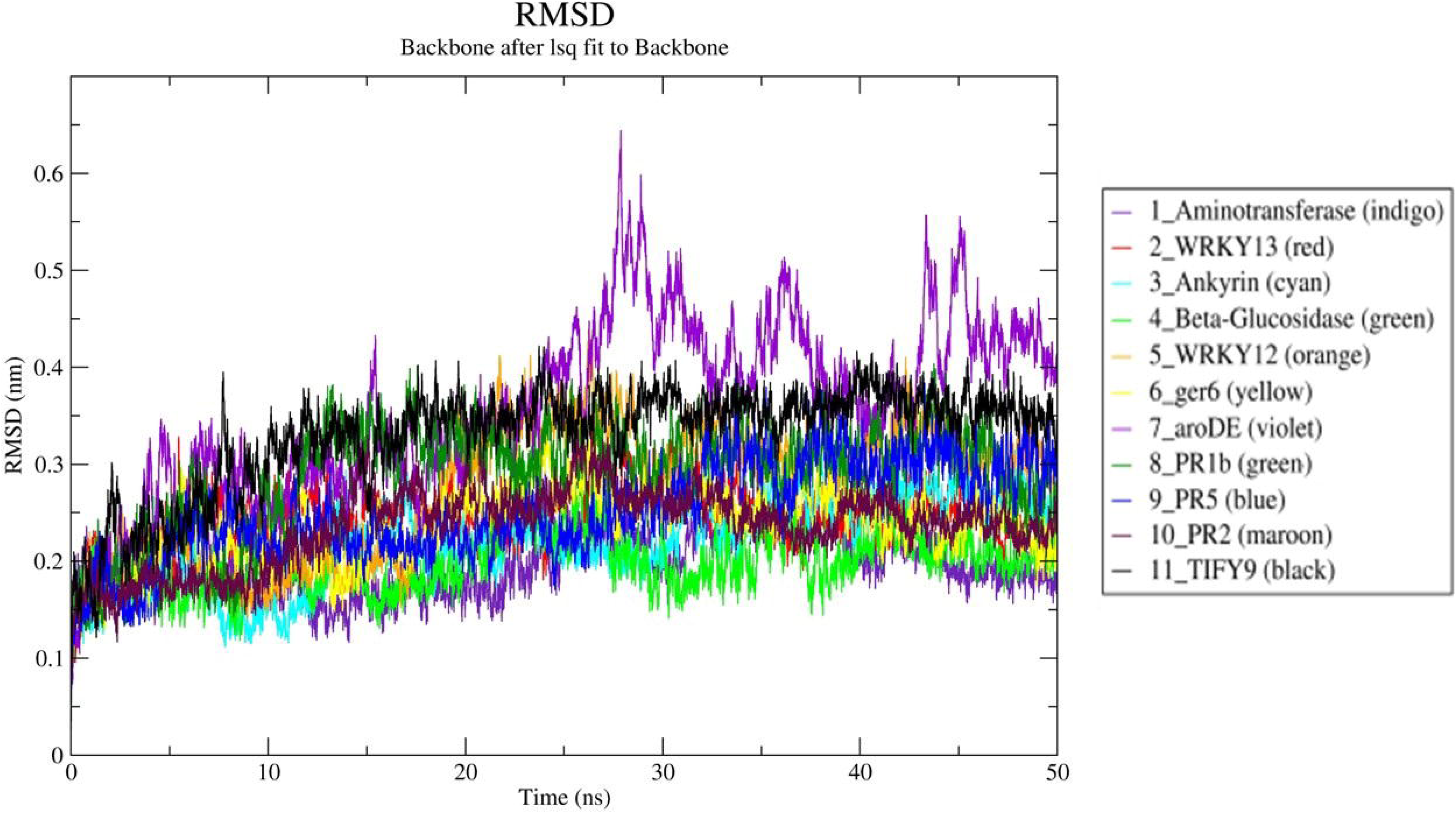
Plot of the Root Mean Square Deviation (RMSD) of WRKY13 Protein with Selected Disease Resistance Related Genes.

**Table 6.** Statistical analysis of MD simulation for Protein-DNA complexes.

### Root Mean Square Fluctuation (RMSF)

RMSF values were estimated to understand the dynamic behaviour of each protein residue during the interaction of gene-WRKY13 complexes. Figures for RMSF of each of the interactions are attached in supplementary figures 2.1-2.11. The RMSF of all the interactions is provided in Figure 9. The RMSF values of all interactions are provided in Table 6 that revealed *aroDE*-WRKY13 (0.167) has the highest mean RMSF value, swiftly followed by *WRKY*12-WRKY13 (0.153), *TIFY*9-WRKY13 (0.130), *ankyrin*-WRKY13 (0.125), *WRKY*13-WRKY13 (0.121), *ger*6-WRKY13 (0.117), *PR*1b-WRKY13 (0.114), *PR*5-WRKY13 (0.113), *PR*2-WRKY13 (0.108), β-*glucosidase*-WRKY13 (0.0995) and *Aminotransferase*-WRKY13 (0.094). In Figure 9, it can be observed that apart from the high fluctuating terminal regions of the protein, significantly fluctuating residues are present in the central regions which are involved in the interaction of DNA. Common fluctuating residues were SER-98, ALA-140, ASN-154, GLU-152, GLY-107, GLY-113, TRP-103, GLY-128, LYS-127, PHE-151, PRO-115, PRO-142, SER-126, THR-143, VAL-147 and PRO158. The residues TRP-103 and GLY107 (Figure 10) from the protein domain WRKYGQK (beta-strand) which has a role in DNA binding, while the nearby fluctuating residues were SER-98, GLY-113 and PRO-115. The other residue present in the beta strands was VAL-147. The other residues are present in the loops and turns of the protein structure.

**Figure 9.**
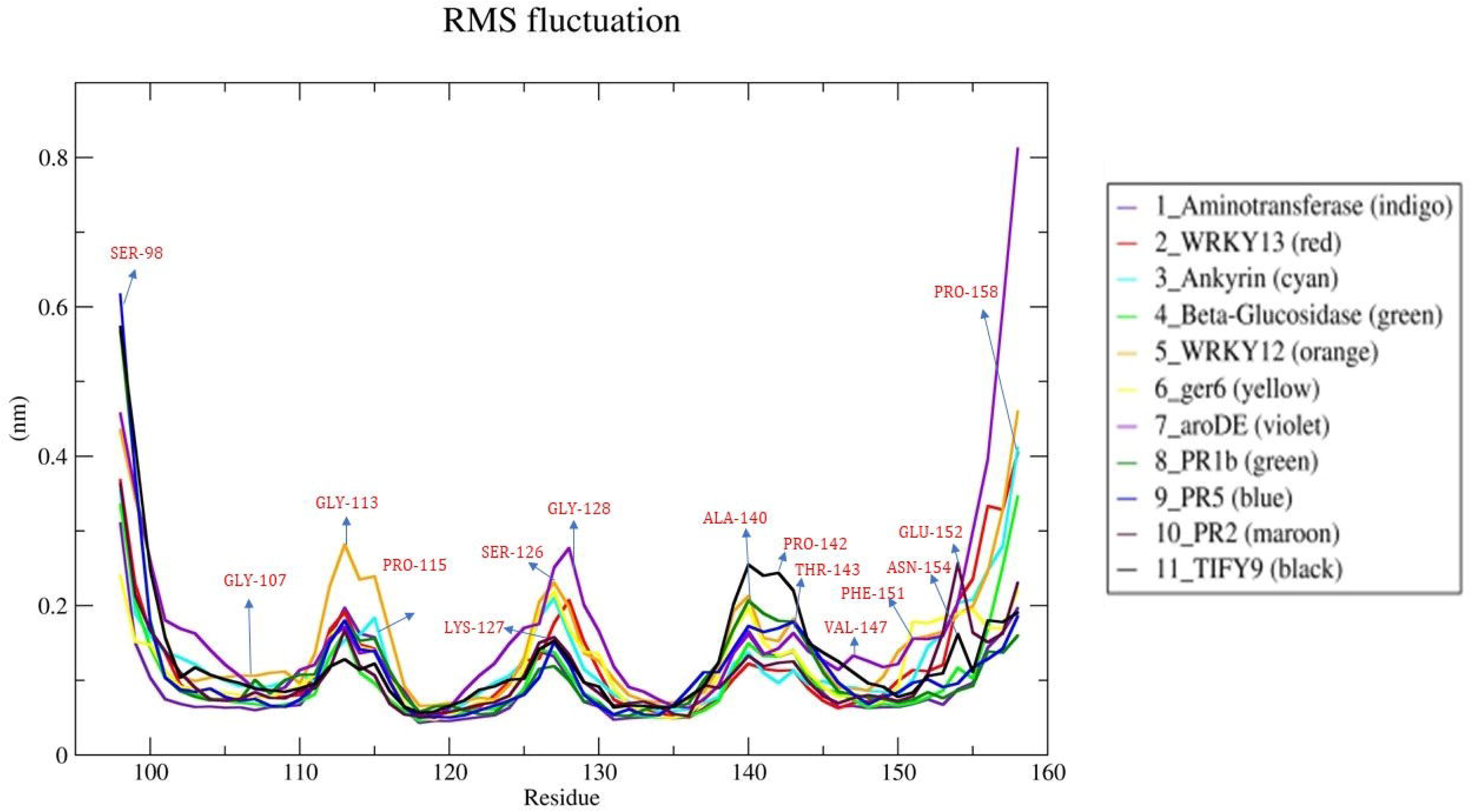
Root Mean Square Fluctuations (RMSF) of WRKY13 Protein with Selected Disease Resistance Related Genes. The fluctuating residues as per the WRKY13 TF protein are marked

**Figure 10.**
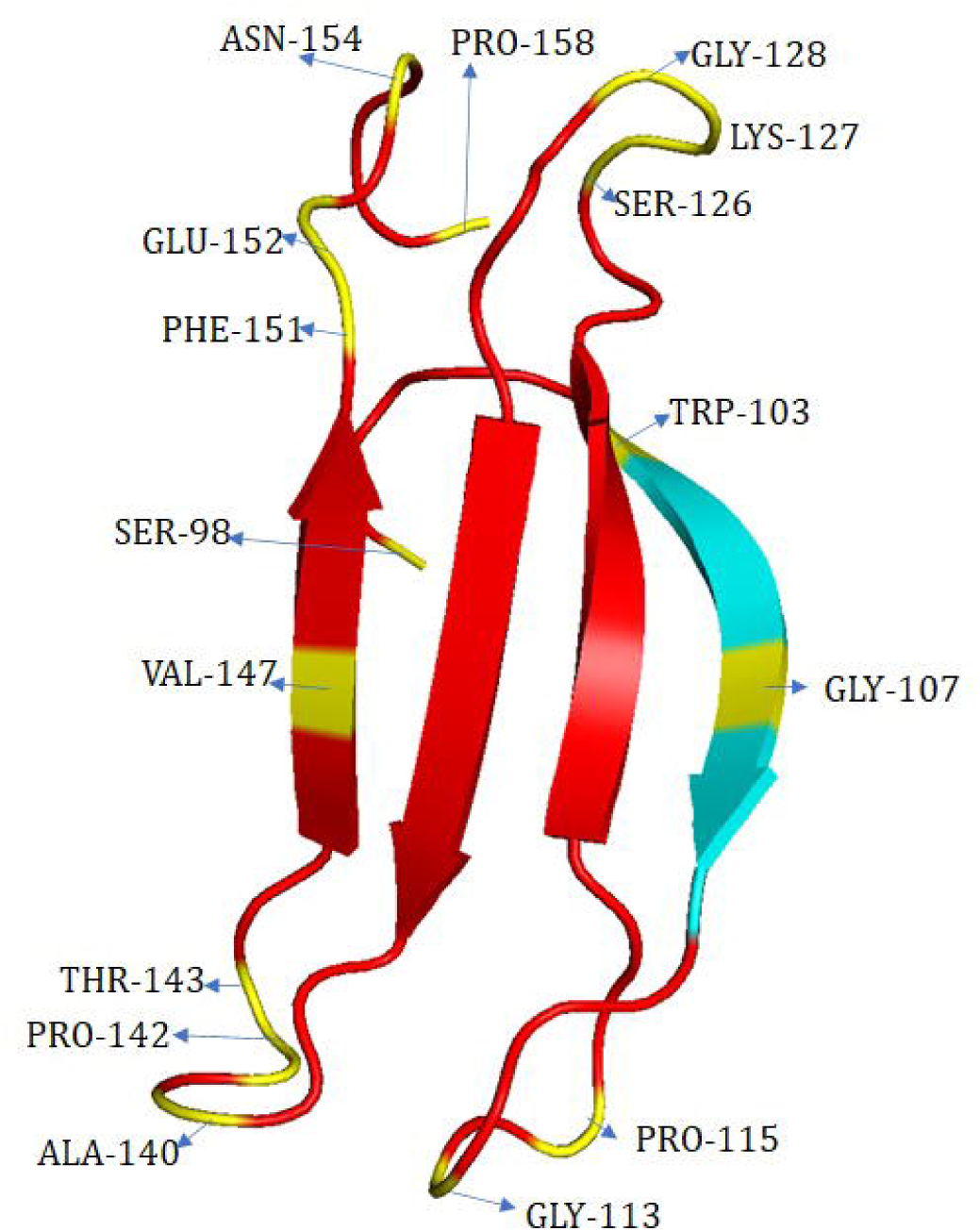
Fluctuating peak residues and their positions in WRKY13 protein. WRKY13 protein in red, fluctuating residues marked in yellow, WRKYGQK conserved domain marked in blue

### The radius of Gyration (ROG)

The compactness if the structure or the complex is given by the radius of gyration (ROG). ROG is the mass-weight root mean square distance from the common centre of mass. ROG was calculated to know the stability of the formed complexes. The larger the distance of the molecule present at the extremities from the centre of the mass implies a higher value of ROG. This reflects that the structure is loosely packed or folded. Therefore, the relation directly corresponds to higher ROG values, the lesser the compactness and vice-versa. Figures for ROG of each of the interactions are attached in supplementary figures 3.1-3.11. The ROG of all the interactions is provided in Figure 11. The ROG values are given in Table 6 that revealed *WRKY*12-WRKY13 (1.389Å) has the highest ROG, followed by *PR*1b-WRKY13 (1.328Å), *WRKY*13-WRKY13 (1.298Å), *PR*5-WRKY13(1.294Å), *aroDE*-WRKY13 (1.315Å), β-*glucosidase-*WRKY13 (1.314Å), *Ankyrin*-WRKY13 (1.307Å), *TIFY*9-WRKY13 (1.310Å), *Aminotransferase*-WRKY13 (1.276Å), *PR*2-WRKY13 (1.276Å) and *ger*6-WRKY13 (1.275Å).

**Figure 11.**
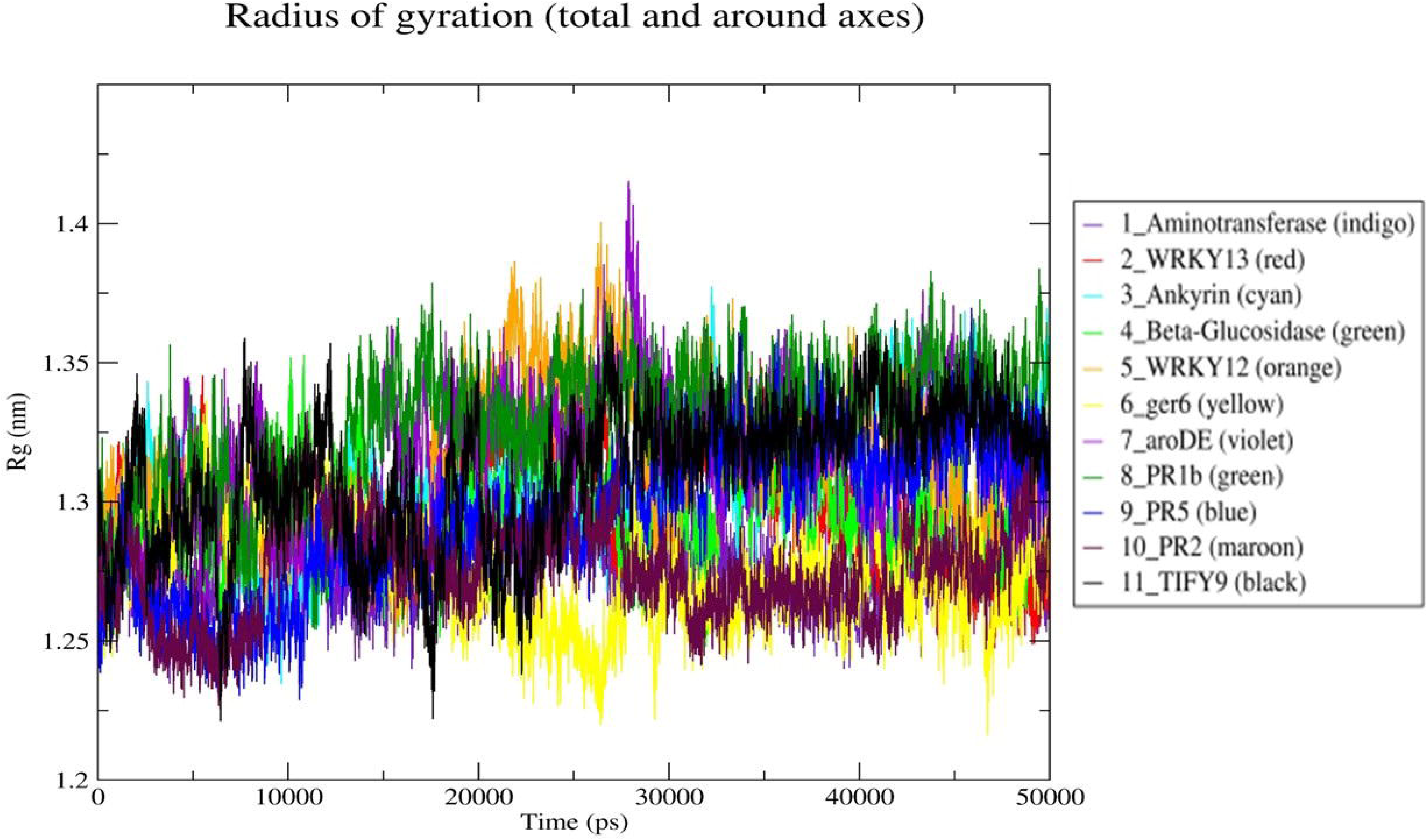
Radius of Gyration (ROG) of WRKY13 Protein with Selected Disease Resistance Related Genes.

### Solvent Accessible Surface Area (SASA)

The surface area that is accessible to the solvent is measured by estimating SASA values. The amino acids buried inside protein molecules are mostly hydrophobic and have a lower surface area in comparison to that of the exposed amino acids to the solvent which are hydrophilic. SASA of the complex is estimated to gain information on the surface area. SASA for all the interactions is provided in Figure 12. Figures for SASA of each of the interactions are attached in supplementary figures 4.1-4.11. The SASA values given in Table 6 revealed *WRKY*13-WRKY13 (51.252) had the highest SASA, followed by *aroDE*-WRKY13 (51.076), *WRKY*12-WRKY13 (50.561), *PR*1*b*-WRKY13 (50.344), *Ankyrin*-WRKY13 (50.018), *β-glucosidase*-WRKY13 (49.925), *ger*6-WRKY13 (49.946), *PR*5-WRKY13 (49.792), *TIFY*9-WRKY13 (49.097), *Aminotransferase*-WRKY13 (48.522), and *PR*2-WRKY13 (47.834). It was observed that after 2 ns, 5ns, 10ns and 10.5ns there was a sharp increase in the SASA plot for all the complexes that is possibly due to folding at the WRKY motif. This may be the reason for the further reduction in the exposed area as evident from each screenshot of the simulation.

**Figure 12.**
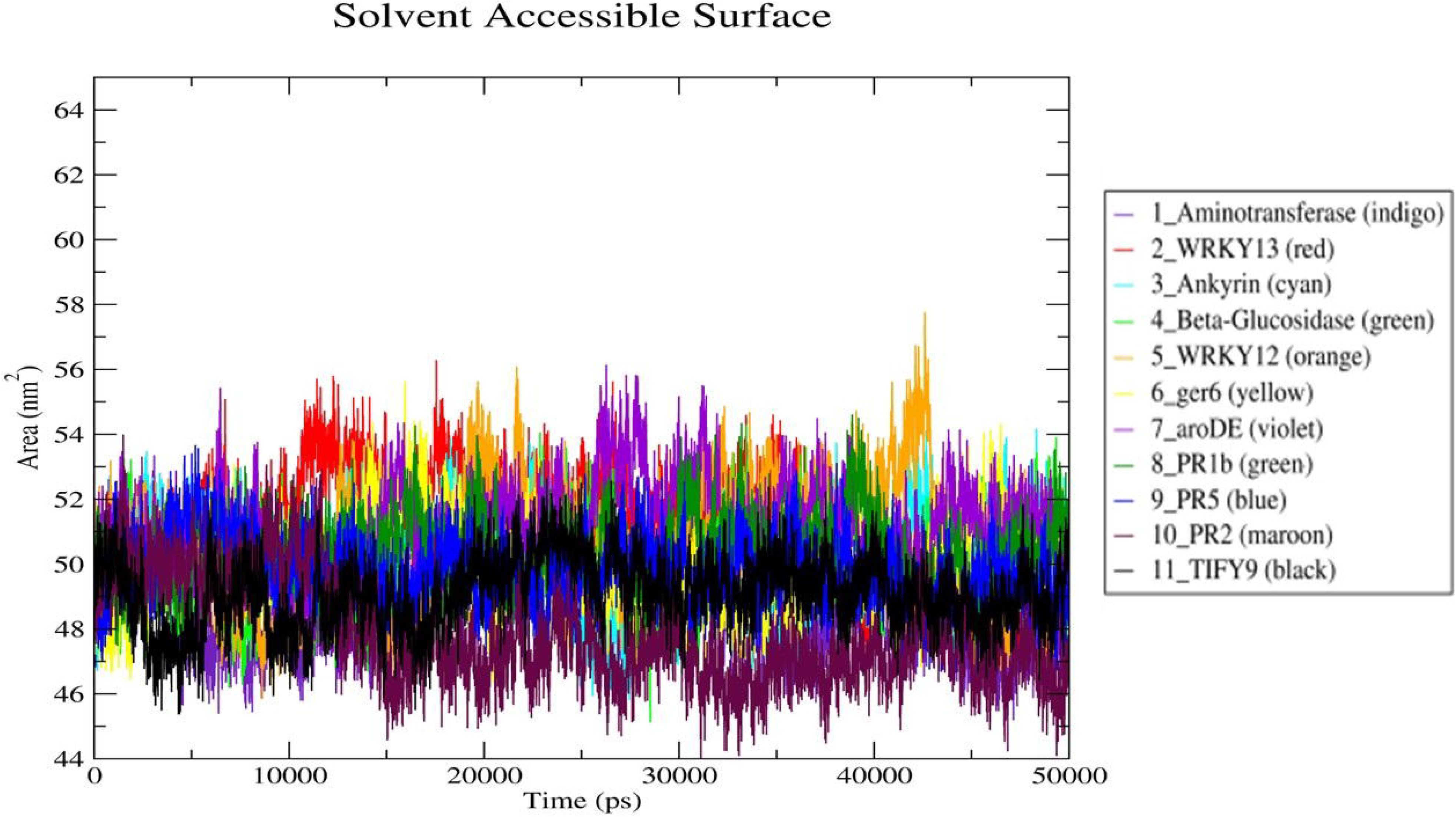
Solvent Accessible Surface Area (SASA) of WRKY13 Protein with Selected Disease Resistance Related Genes.

### Hydrogen Bonds

The stability of the complex was analyzed by calculating the total number of intramolecular hydrogen bonds. A higher number of hydrogen bonds is equivalent to higher stability of the binding complex. Figures for Hydrogen bonds of each of the interactions are attached in supplementary figures number 5.1-5.11. The hydrogen bond for all the interactions is provided in Figure 13. The hydrogen bond values are given in Table 6 that revealed *PR*2-WRKY13 (4.527) has a higher number of hydrogen bonds, followed by *WRKY*13-WRKY13 (2.469), *β-glucosidase*-WRKY13 (0.877), *aroDE*-WRKY13 (1.515), *Ankyrin*-WRKY13 (1.456), *TIFY*9-WRKY13 (1.449), *PR*5-WRKY13 (1.155), *WRKY*12-WRKY13 (0.656), *ger*6-WRKY13 (0.385), *aminotransferase*-WRKY13 (0.012) and *PR*1b-WRKY13 (0.003). Both a*minotransferase*-WRKY13 and *PR*1b-WRKY13 interactions had a lower number of hydrogen bonds. An increased number of hydrogen bonds implies a more stable DNA-protein complex. The trajectories of MD simulations were analyzed using VMD and at every 10ns, the frames were captured to indicate the structural changes across the simulations (Figure 14).

**Figure 13.**
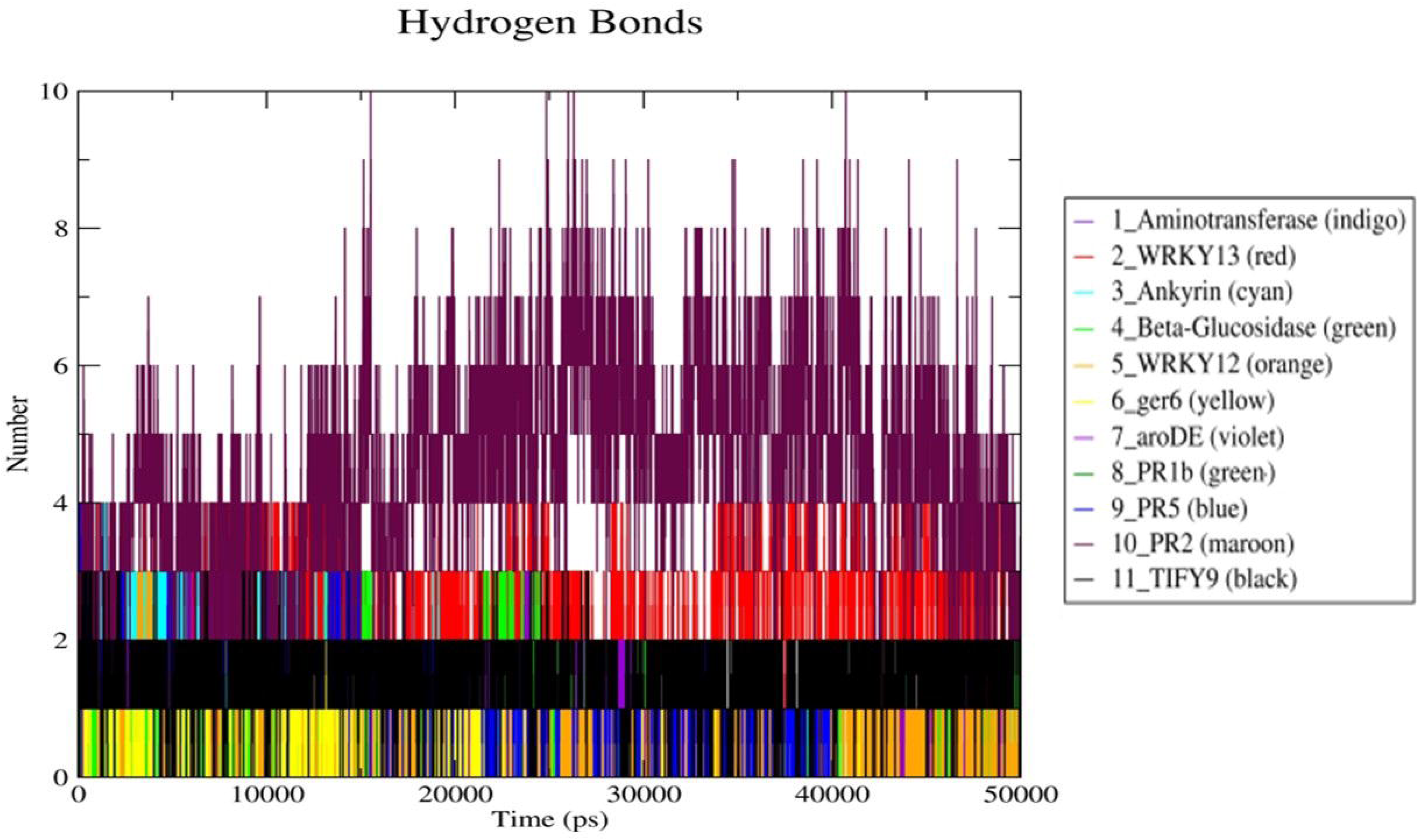
Hydrogen Bonds of WRKY13 Protein with Selected Disease Resistance Related Genes.

**Figure 14.**
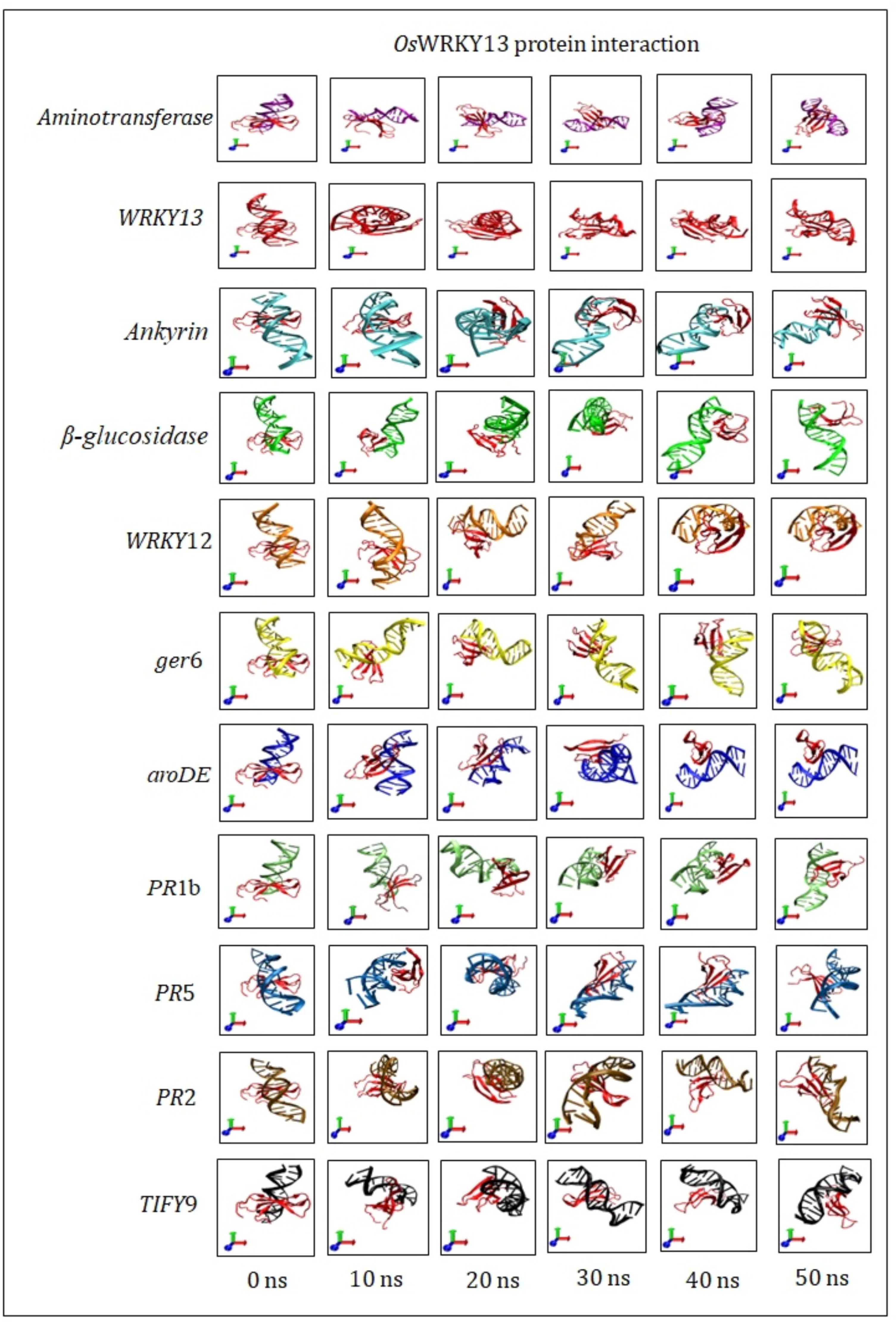
Snapshots of Simulations at 0-10 ns Time Interval for WRKY13 and Interacting Rice Disease Resistance Related Genes.

### Co-expression network of selected disease resistance-related genes

Microarray data of all 11 genes for pathogen infection (*M. grisea* and *Xoo*) was collected from the ROAD database (Cao *et al*.,2012). The heatmap of the expression of all 11 genes was obtained from the ROAD database indicating that the genes showed significant change (fold change ≥ 2 or fold change ≤ 0.5) in response to both pathogens. The heatmap of the selected platforms is represented in Figure 15. The expression values were further evaluated using a cystoscope to evaluate their interaction with each other. The network (Figure 16) revealed that *Aminotransferase* and *PR*1b may not be easy targets of WRKY13 TF but an extension of the network. Whereas *PR*5, *WRKY*12, *ger*6, *PR*2, *Ankyrin*, *glucosidase*14, W*RKY*13, *aroDE* and *TIFY*9 interact directly with each other. The gene network provides a probable mechanism through which the WRKY13 TF protein interacts with other genes.

**Figure 15.**
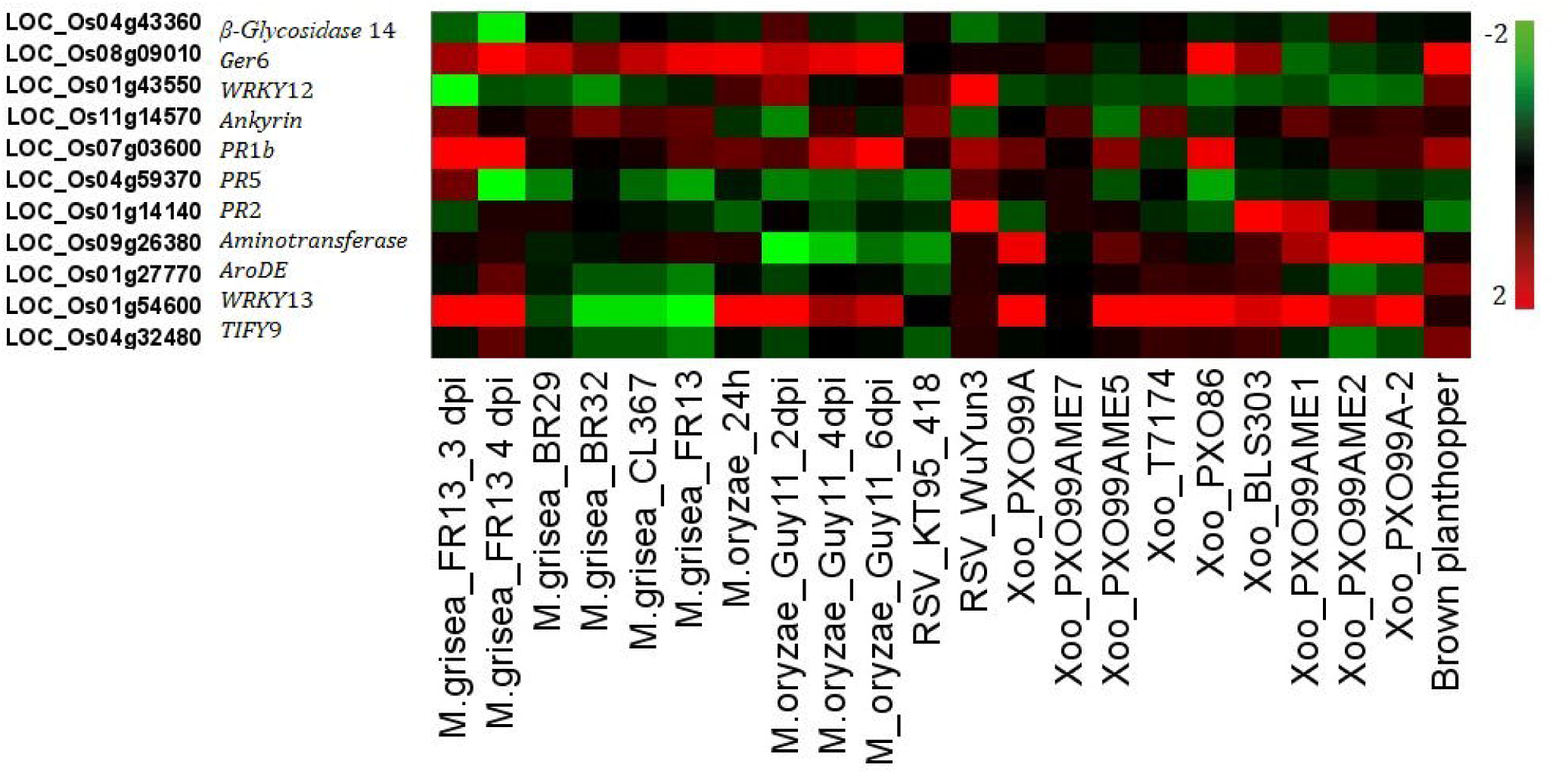
Expression patterns of Selected Disease Resistance Related Genes identified for rice infected with *Magnoparthe grisea* and *Xanthomonas oryzae* pv. *oryzae*.

**Figure 16.**
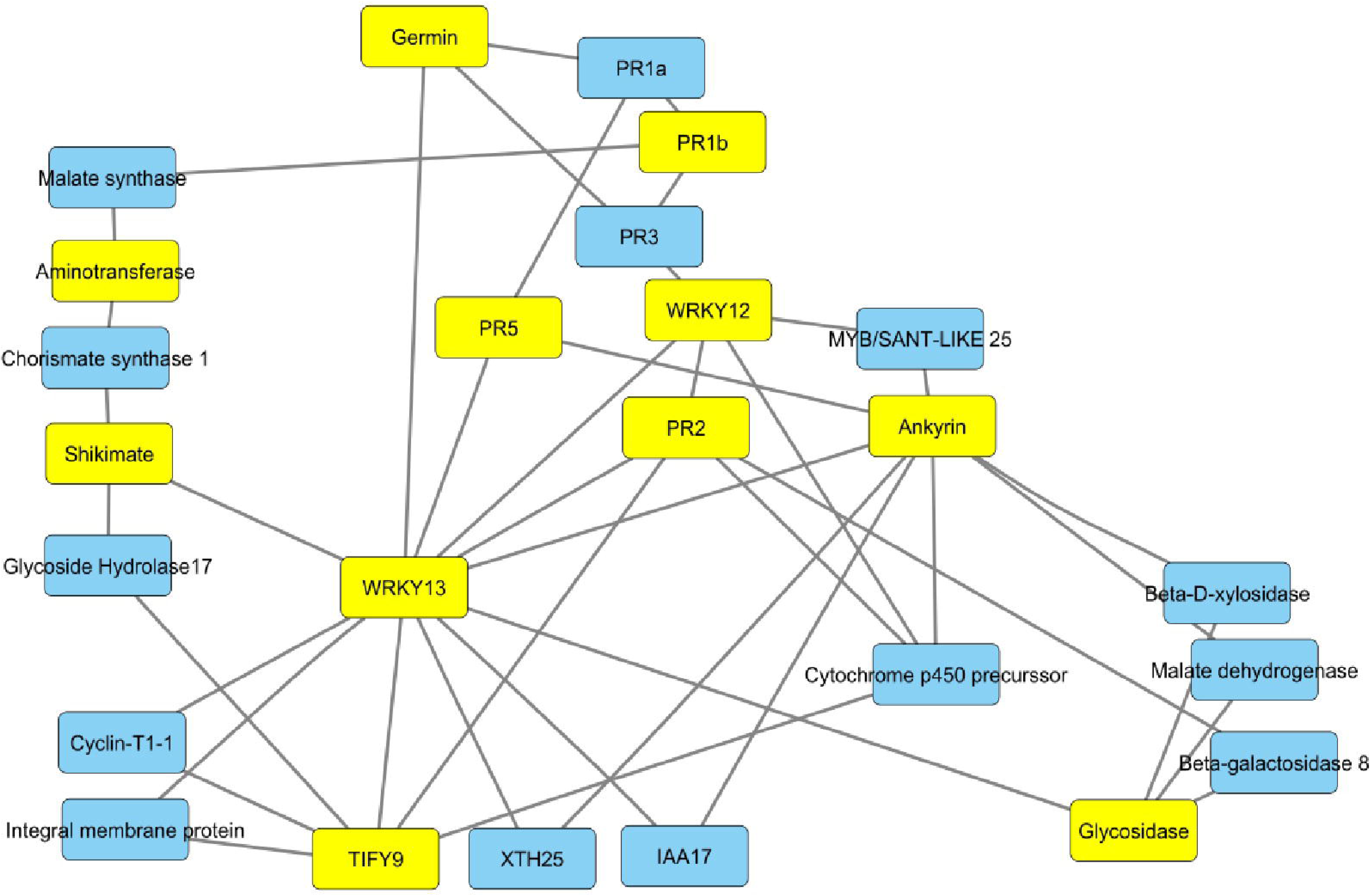
Gene network revealing the interaction of the Rice Disease Resistance-Related Genes. Yellow rectangles represent selected disease resistance genes, blue rectangles represent interacting genes

## Discussion

The overexpression of WRKY13 in rice resulted in the expression and suppression of 236 and 273 genes respectively (Qiu *et al*., 2008). The promoter analysis of these target genes helped to understand that the up-regulated genes were controlled by other WRKY proteins and the down-regulated genes were regulated by MYB and AP2/EREBP proteins (Qiu *et al*., 2009). WRKY TF family are important regulators in defence response pathways against pathogen infection (Yang *et al*., 2022). However, the promoter analysis in the entire genome will help to identify the possible target genes in the entire rice genome controlled by WRKY13 TF. Promoter analysis of many pathogen-inducible genes has helped to find possible gene targets in the past. Promoter analysis in rice helped to identify pathogen-inducible cis-regulatory elements (PICE) such as GCC-box, G-box, AS-1 and H-box in the genes that were regulated during a pathogen attack (Kong *et al*., 2018). Promoter analysis revealed six different binding sites of transcription factors in the promoters of GLP (Germin-like proteins) in rice (Das *et al*., 2019). The mechanism of *myb*4 against sheath blight in rice was explored using genome-wide promoter analysis in rice (Pooja *et al*., 2015). In the present study, genome-wide promoter analysis was carried out to find the 1,012 target genes having 4 or more WRKY13 TF binding sites in their promoter regions. All these genes were subjected to gene ontology and enrichment analysis to classify them. Disease resistance-related genes were selected and the presence of a W-box *cis*-acting regulatory element was estimated. These genes were also subjected to docking studies to find the interacting residues.

The WRKY binding site is characterized by the presence of a W-box, a hexamer of TTGAC(C/T). Variants of the W-box and W-box-like cis-element are reported such as TTTGACA, TTTGAC(C/T), TTGACTT, TTGAC(A/C), TTGAC(A/C)A, and TTGAC(A/C)(C/G/T), TTGACG and a W-box like cis-element: TGAC(C/T) (Maleck *et al*., 2000; Eulgem *et al*., 2000; Kankainen *et al*., 2004; Qiu *et al*., 2007). *In vitro* studies carried out using rice WRKY13 revealed that rice WRKY13 TF protein bound to PRE4 a cis-element present in its promoter and thus auto-regulates. PRE4 is characterized by TACTGCGCTTAGT sequence (Cai *et al*., 2008). Tobacco *Nt*WRKY12 bound to the WK-box (TTTTCAC) in the *PR*1a promoter, a variant of W-box (Van Verk *et al*., 2008). Rice WRKY13-downregulated genes had promoters enriched with the GGTTAGTTA element that harboured the Myb1 element (Chakravarthy *et al*., 2003). The GTACGTAC motif containing the ACGTATERD1 and ACGTABOX elements required for the binding of AP2/EREBP or bZIP proteins was found to be enriched in WRKY13-downregulated genes. The promoters of the WRKY13-upregulated genes were enriched with GTTGAC(C/T) variant of TTGACCTC that harboured the W-box (Qiu *et al*., 2009). The GTTGACC motif was found to be more enriched than the GTTGACT motif in both strands of the promoters. The GTTGAC motif that harboured the W-box (TTGAC) was enriched in both strands of the upregulated gene promoters (Qiu *et al*., 2009). This explains the important role played by WRKY transcription factors in the regulation of various differentially expressed genes.

Genes related to disease resistance were selected from the 1,012 genes obtained after fixing a threshold of four or more motifs in the promoter regions of the genes. Even though, the present study is motivated to explore the target genes responsible for disease resistance in rice, the classification of the genes using gene ontology and enrichment analysis draws light toward the genes regulated by the WRKY13 TF protein. These genes have various biological, cellular and molecular functions. Focusing on the genes responsive to biotic stress with existing reports of their role in rice disease resistance were considered for further analysis. Hence the genes *PR*2, *WRKY*12, *PR*5, *β-Glycosidase*14, *aminotransferase*, germin 8-7 (*ger*6), *PR*1b, *ankyrin*, shikimate biosynthesis protein (*aroDE*) were further subjected to analysis. The algorithm used in the present study also revealed that the promoter region of *WRKY*13 had 3 motifs (TGACT, TGACY, CTGACY) in its promoter sequence. *WRKY*13 had existing reports of autoregulating its promoter regions in rice (Xiao et al., 2013; Kong *et al*., 2018). Therefore, *WRKY*13 was also considered for further analysis. Previously, we proved that the *WRKY*13 gene-mediated network was proven to mediate disease resistance through the *TIFY*9 gene against both sheath blight and sheath rot diseases (Jimmy and Babu, 2019). *TIFY*9 was further analyzed to study the regulation by the *WRKY*13 TF gene. C*is*-acting regulatory elements analysis of the selected genes helped to find the type of cis-elements and to validate the presence of the motifs. These elements may be present in the direct or the reverse strand of the promoter (Kong *et al*., 2018). It was reported through various experiments that promoters of a particular *cis*-acting regulatory element respond to a specific trigger. The factors found here (W-box) were reported to be responsible for the binding of WRKY TFs (Kong *et al*., 2018) and also direct pathogen-inducible expression.

The WB box (WBBOXPCWRKY1) was reported to allow specific binding of WRKY protein to the DNA sequence motif W-box (T)(T)TGAC(C/T). It was found in the amylase gene in sweet potato, *αAmy*2 genes in wheat, barley, and wild oat, the *PR*1 gene in parsley, and a transcription factor gene in *Arabidopsis*. The motif was initially registered in PLACE database as TTGACT and was corrected to TTGACY (Y=C/T) (Ishiguro and Nakamura, 1994; Rushton *et al*., 1995, Rushton *et al*., 1996; De Pater *et al*., 1996; Eulgem *et al*., 2000). Whereas TTTGACY (WBOXATNPR1) was the W-box found in the promoter of the *Arabidopsis NPR*1 gene and was recognized specifically by salicylic acid (SA)-induced WRKY DNA binding proteins. It acts as the negative regulatory element for the inducible expression of *AtWRKY*18 (Chena and Chen, 2002, Yu *et al*., 2001; Chen *et al.,* 2002; Eulgem *et al.,* 2000; Maleck *et al.,* 2000; Xu *et al*., 2004). In cotton *delta-cadinene synthase*-A gene (*CAD*1A) promoter region was reported to have the motif TTGAC/ TTGACC (WBOXGACAD1A), a W-box that allows the binding of *Ga*WRKY1. *Ga*WRKY1 regulates sesquiterpene biosynthesis in cotton. (Xu *et al*., 2004). In barley, SUSIBA2 was reported to bind to the W-box element (AGTCAAAATTGACC) found in the promoter region of the *isoamylase*1 (*iso*1) promoter (Sun *et al*., 2003). Whereas, in tobacco, the W box in the promoter region of the class I basic chitinase gene was reported to have the WBOXNTCHN48 harboured in the TGACT motif. These sites were reported to act as binding sites of WRKY1, WRKY2 and WRKY4. These WRKYs led to elicitor-responsive transcription of defense genes in tobacco (Yamamoto *et al.,* 2004). Also, in tobacco, the CTGACY harbouring the cis-element WBOXNTERF3 found in the promoter region of a transcriptional repressor *ERF*3 gene in tobacco leads to activation of the ERF3 gene due to wounding (Nishiuchi *et al*., 2004).

*WRKY*13 gene was reported to play a mediatory role in signalling in plant physiological pathways related to either development or disease resistance. *WRKY*13 gene is a potential regulator of multiple physiological processes since it influences more than 500 genes that are involved in multiple signalling pathways belonging to disease resistance, redox homeostatic, abiotic stress responses and plant development. *WRKY*13 gene also regulates antagonistic cross-talk between abiotic and biotic stress. Qiu *et al*. (2008) determined the active role of *WRKY*13 in rice resistance against both bacterial and fungal pathogens and its function as a mediator of crosstalk between the genes of pathogen-induced salicylic-dependent defense pathway and other five physiological pathways. In a parallel study, chitinase and germin-like protein were reported as over-lapping proteins during both biotic and abiotic stress responses in rice (Sruthilaxmi and Babu, 2020). This indicated a common intricate network in play during cross-talk that regulated the functions of both these genes. Similarly, the present study of genome-wide promoter analysis revealed the bm of WRKY13 protein in the promoters of various target genes that were involved in a wide array of physiological processes and disease resistance mechanisms. This suggested the functioning of specific gene networks. The gene network mediated by *WRKY*13 against sheath infecting fungi sheath blight and sheath rot revealed such a network in action with other transcription factors (*TIFY*9) playing a mediatory role in activating the core genes involved in MAPK cascade. The activation of the MAPK pathway then amplifies the signal by phosphorylating other MAPK genes sequentially (Shi *et al*., 2020). It was revealed that the expression of *WRKY*13 induced other disease resistance-related genes such as *WRKY*12 and *PR*2 (Jimmy and Babu, 2019). Qiu *et al*., (2007) reported that *WRKY*13-overexpressing rice has increased resistance against bacterial leaf blight (*Xanthomonas oryzae* pv *oryzae*) and rice blast disease (*M. oryzae*). Also, *WRKY*13-overexpressing rice reported increased resistance against both the sheath-infecting fungal pathogens: *R. solani* and *S. oryzae* (Jimmy and Babu, 2019). Therefore, if a single TF can be regulated it could help modulate the expression of several other genes in the genome. The genome-wide promoter analysis to search rice WRKY13 TF bm indicates the mechanism involved in the functioning of the network which is a result of the binding of WRKY13 TF proteins to the promoter regions of these disease resistance-related genes. The presence of cis-acting regulatory elements in the promoter regions of these nine selected genes indicates their probable role in rice disease resistance.

*WRKY*12, *aminotransferase*, and *ankyrin* are either directly involved in disease resistance or are activated by other genes/TFs. Pooja *et al*., (2015) stated during *R. solani* infection, *WRKY*12, *aminotransferase*, and *ankyrin* are expressed. WRKY12 TF was demonstrated to play several roles in plant systems including plant development and immunity. WRKY12 TF triggered the expression of *NPR*1, *PR*1b, phenylalanine ammonia-lyase (*ZB*8) and peroxidase (*POX*22.3). WRKY12 was also described to be a transcriptional regulator in SA-or JA-dependent defense signalling cascades (Liu *et al*., 2005). Expression of *WRKY*12 resulted in the upregulation of PR genes (Jimmy and Babu, 2015), and was described to co-express along with *WRKY*13 (Jimmy and Babu, 2019) and *myb*4 (Pooja and Babu, 2015). Aminotransferase catalyzes the conversion of amino acids and oxoacids by the transfer of amino groups. Infected plant cells show deposition of callose, thereby, getting thickened and encasing fungal mycelia within them. Therefore, massive lignification of the plant cells prevents the progression of mycelia (Cohen *et al*., 1989). Ankyrins, on the other hand, are intracellular adaptor proteins, that target diverse proteins in both the endoplasmic reticulum and the plasma membrane. *M. oryzae* infection in rice induced the expression of Ankyrin repeat-containing protein (*PIANK*1) (Zang *et al*., 2010). Ankyrin proteins are also proved to target the function of other genes downstream of the receptor proteins to regulate plant immunity (Yang *et al*., 2012). Over-expression of ankyrin repeat-containing protein (*PIANK*1) in rice was shown to enhance the expression of *WRKY13, PR1b and PR5* against *M. oryzae infection* (Mou *et al*., 2013).

*PR*5, *PR*1 and *PR*2 expression were enhanced in *Xa*21-mediated resistance response after inoculation with *Xoo* in the resistance response, whereas, *PR*5 showed changes in susceptible cultivars. (Hou *et al*., 2012). During *M. oryzae* infection, *PR*5 expressed in *WRKY*53 over-expressing rice (Chujo *et al*., 2007) while *PR*2 revealed enhanced expression during gene pyramiding (Fukuoka *et al*., 2015). Exogenous application of JA and ET significantly increased the expression level of *PR*1b in the seedlings. Whereas, the control sample was treated with a pathogen (*Xoo*) and water treatment (Luan and Zhou, 2015). *M. oryzae* induced the expression of PR genes such as *PR*1b, *PR*2 and *PR*5 (Schweizer *et al*., 1997). Individually, *PR*1b was described as an important link of a defense signalling molecule-mediated network that results in the regulation of genes involved in disease resistance. Xie *et al*., (2011) demonstrated that *PR*1b has basal transcription levels in older rice leaves infected with *M. oryzae. PR*2 coexpressed along with *NAC*111, a blast-responsive gene, in response to *M. oryzae* (Yokotani *et al*., 2014).

The Shikimate dehydrogenase (*aroDE*) plays a role in the biosynthesis of the chorismate which leads to the biosynthesis of aromatic amino acids. It catalyzes the reversible NADPH-linked reduction of 3-dehydroshikimate (DHSA) to yield shikimate (SA) (UniProt Consortium, 2015). Suppressed aroDE expression in tobacco reduced the content of aromatic amino acids and downstream products (Hittinger and Carroll, 2007). Biosynthetic pathways including the shikimate, phenylpropanoid, and arylmonoamine pathways are reported to coordinately activate phenolic phytoalexin synthesis, and related genes induced by biotic or abiotic stresses in rice (Cho and Lee, 2015). *Ger*6 is expressed during *M. oryzae* infection (Manosalva *et al*., 2009). Germin 8-7 (g*er*6) was described to be induced in broad-spectrum disease resistance (UniProt Consortium, 2015; Sakai *et al*., 2013). β-Glycosidase14 (*Bglu*14) along with β-Glycosidase16 and β-Glycosidase18 hydrolyze the monolignol glucosides coniferin and syringin (Baiya *et al*., 2018). All these genes have WRKY13 TF binding sites harbouring W-box cis-elements in their promoter region. These genes were further analyzed using docking studies.

DNA-protein docking involves comparative modelling of transcription factors and has proven to be an efficient method to provide detailed information regarding their interactions (Kolinski and Skolnick, 1994; Sánchez and Šali, 1997). Proteins of WRKY family members are reported to specifically bind to varying DNA motifs involving a common binding consensus core (Ciolkowski *et al*., 2008; Yamasaki *et al*., 2012; Brand *et al*., 2013). In the WRKYGQK conserved domain, the highly conserved lysine (lys) favours the negatively charged DNA phosphate backbone (Yamasaki *et al*., 2012). The WRKY conserved residues are the major interacting residues in WRKY13 protein-gene interaction. These conserved residues have evolutionary significance as their interaction with specific ligand molecules triggers a response to various stimuli that play a vital role in disease resistance (Karkute *et al*., 2015). Hydrogen bonds and hydrophobic interactions play an essential role in stabilizing the protein-DNA complex (Pandey *et al*., 2018). The interaction of TF protein and DNA is necessary for the regulation of transcription. Therefore, the characterization of TF binding sites throughout the rice genome is essential to find the genes regulated by the corresponding TF, thus determining the regulatory network (Hannenhalli, 2008).

A molecular dynamics simulation for 11 rice genes with WRKY13 transcription factor complexes was carried out. Molecular docking was first carried out for all 11 DNA models and the WRKY13 TF protein. The interactions predicted using nucplot matched with the interactions as predicated by the RMSF plot. Lower stability was observed in the *Aminotransferase*-WRKY13 complex, as the mean value of scalar distance (RMSD), and residue-level fluctuation (RMSF), were least from other docked complexes. Whereas ROG was lower for *Ge*r6, SASA was lower for *PR*2-WRKY13 and H-bond was low for both *PR*1b-WRKY13 and *Aminotransferase*-WRKY13 complex. The lowered Hydrogen bond value was next to negligible which indicates that even though the motifs are present in the promoter regions of these genes, these might be resent further away to be targets of WRKY13 TF protein. The interactions for *PR*2-WRKY13, *WRKY*12-WRKY13, *WRKY*13-WRKY13 and *aroDE-WRKY13* were reported to have the highest simulation values. These genes might be the easy targets of WRKY13 TF proteins. A similar MD simulation study on WRKY DNA binding domain was performed on barley to understand the effect of mutation on the DNA recognition of the WRKY family. This study considered only interactions among the WRKY family, but one study provides insights into different gene interactions WRKY13 transcription factor (Pandey *et al*.,2018). Another study has shown the effect of structural diversity on the DNA binding of WRKY protein, where they have illustrated the significant role of H-bonding in stabilizing the protein-DNA complex. The results from this study reveal the correlation of the role of H-bonding in protein-DNA complex during recognition (Pooja *et al*., 2015). While Dai *et al*. (2021) revealed that a 1 kb stepping motion was seen where the run for 10ms long time intervals. In our study, no such stepping was observed and protein was seen to be attached to the same grove till the completion of the run. The use of high-performance computation can be applied to simulating the system in a long run for more insights into the binding mechanism of WRKY protein.

The interaction of all the 11 genes revealed that *Aminotransferase* and *PR*1b support that these genes are not easy targets of WRKY13 but an extension of the network. Whereas *other genes* interact directly with WRKY13. The gene network provides the probable mechanism through which the WRKY13 TF protein interacts with other genes. Based on these findings, the complex network of WRKY-mediated signalling pathways towards biotic stress in rice was elaborated. WRKY13 interaction with other genes is reported here. The network mechanism of WRKY13 TF protein was derived from rice biotic stress microarray data and confirms the previous WRKY13 network against biotic stress. This study provides WRKY13 TF-regulated gene targets involved in disease resistance mechanism and thus traces the molecular mechanism of WRKY13 against rice pathogens.

## Supporting information

Manuscript_Tables

## SUPPLEMENTARY FIGURES

**Supplementary Figure 1.1 – 1.11.** Root Mean Square Fluctuations (RMSF) of WRKY13 Protein with Selected Disease Resistance Related Genes.

*1.Aminotransferase*-WRKY13; 2. *WRKY*13-WRKY13**; 3.** *Ankyrin repeat*-WRKY13; 4. *β-Glycosidase*14-WRKY13; 5. *WRKY*12-WRKY13; 6. *ger*6-WRKY13; 7. *aro*DE-WRKY13; 8. *PR*1b-WRKY13; 9. *PR*5-WRKY13; 10. *PR*2-WRKY13; 11. *TIFY*9-WRKY13

**Supplementary Figure 2.1 – 2.11.** Root Mean Square Fluctuations (RMSF) of WRKY13 Protein with Selected Disease Resistance Related Genes.

*1.Aminotransferase*-WRKY13; 2. *WRKY*13-WRKY13**; 3.** *Ankyrin repeat*-WRKY13; 4. *β-Glycosidase*14-WRKY13; 5. *WRKY*12-WRKY13; 6. *ger*6-WRKY13; 7. *aro*DE-WRKY13; 8. *PR*1b-WRKY13; 9. *PR*5-WRKY13; 10. *PR*2-WRKY13; 11. *TIFY*9-WRKY13

**Supplementary Figure 3.1 – 3.11.** Solvent Accessible Surface Area (SASA) of WRKY13 Protein with Selected Disease Resistance Related Genes.

*1.Aminotransferase*-WRKY13; 2. *WRKY*13-WRKY13**; 3.** *Ankyrin repeat*-WRKY13; 4. *β-Glycosidase*14-WRKY13; 5. *WRKY*12-WRKY13; 6. *ger*6-WRKY13; 7. *aro*DE-WRKY13; 8. *PR*1b-WRKY13; 9. *PR*5-WRKY13; 10. *PR*2-WRKY13; 11. *TIFY*9-WRKY13

**Supplementary Figure 4.1 – 4.11.** Radius of Gyration (ROG) of WRKY13 Protein with Selected Disease Resistance Related Genes.

*1.Aminotransferase*-WRKY13; 2. *WRKY*13-WRKY13**; 3.** *Ankyrin repeat*-WRKY13; 4. *β-Glycosidase*14-WRKY13; 5. *WRKY*12-WRKY13; 6. *ger*6-WRKY13; 7. *aro*DE-WRKY13; 8. *PR*1b-WRKY13; 9. *PR*5-WRKY13; 10. *PR*2-WRKY13; 11. *TIFY*9-WRKY13

**Supplementary Figure 5.1 – 5.11.** Hydrogen Bonds of WRKY13 Protein with Selected Disease Resistance Related Genes.

*1.Aminotransferase*-WRKY13; 2. *WRKY*13-WRKY13**; 3.** *Ankyrin repeat*-WRKY13; 4. *β-Glycosidase*14-WRKY13; 5. *WRKY*12-WRKY13; 6. *ger*6-WRKY13; 7. *aro*DE-WRKY13; 8. *PR*1b-WRKY13; 9. *PR*5-WRKY13; 10. *PR*2-WRKY13; 11. *TIFY*9-WRKY13

## SUPPLEMENTARY TABLES

**Supplementary Table 1.** List of rice genes having WRKY13 bm in their promoters

**Supplementary Table 2-12.** Interacting residues of WRKY13- *aminotransferase*, WRKY13- *ankyrin,* WRKY13- *ger6,* WRKY13- *βGlycosidase*14. WRKY13- *PR1b,* WRKY13- *PR2,* WRKY13- *PR5,* WRKY13- *aroDE,* WRKY13- *WRKY12*, WRKY13- *TIFY9,* and WRKY13-*WRKY*13 complexes

