## Supplementary material for "Rice WRKY13 TF protein binds to motifs in the promoter region to regulate downstream disease resistance-related genes": Manuscript_Tables: Manuscript_tables_Jimmy et al. 2023.pdf

**Table 1.** Binding Motifs of WRKY13 TF

| S. no | Motif |
|---|---|
| 1 | TTGAC |
| 2 | TTGACC |
| 3 | TTGACT |
| 4 | TTTGACC |
| 5 | TTTGACT |
| 6 | TTGACA |
| 7 | GTTGACC |
| 8 | GTTGACT |
| 9 | GTTGAC |
| 10 | TTGACCTC |
| 11 | CTGACC |
| 12 | CTGACT |
| 13 | TTGACCA |
| 14 | TTGACCT |
| 15 | TTTTCCAC |
| 16 | GGTTAGTTA |
| 17 | GTACGTAC |
| 18 | TTGACTT |
| 19 | TTGACG |
| 20 | TGACC |
| 21 | TGACT |

**Table 2.** Occurrences and Chromosome-Wise Distribution of WRKY13 Bm in Rice Genome

| Motif | CHR 1 | CHR 2 | CHR 3 | CHR 4 | CHR 5 | CHR 6 | CHR 7 | CHR 8 | CHR 9 | CHR 10 | CHR 11 | CHR 12 |
|---|---|---|---|---|---|---|---|---|---|---|---|---|
| TTGAC | 23647 | 19172 | 20462 | 15051 | 13418 | 14587 | 12559 | 11253 | 9272 | 9128 | 10619 | 9769 |
| TGACC | 16265 | 12984 | 14056 | 10177 | 8740 | 9433 | 8425 | 7483 | 6157 | 6004 | 7206 | 6289 |
| TGACT | 18720 | 15443 | 16419 | 12557 | 10791 | 11432 | 10313 | 9521 | 7788 | 7578 | 8775 | 7844 |
| TTGACC | 5500 | 4490 | 4528 | 3188 | 3030 | 3338 | 2842 | 2559 | 2182 | 2052 | 2475 | 2279 |
| TTGACT | 6664 | 5722 | 5820 | 4456 | 3873 | 4207 | 3677 | 3520 | 2775 | 2736 | 3231 | 2950 |
| TTGACA | 7991 | 6136 | 6938 | 5084 | 4456 | 4899 | 4201 | 3670 | 3012 | 2988 | 3462 | 3159 |
| GTTGAC | 4608 | 3575 | 3920 | 2874 | 2579 | 2745 | 2245 | 2213 | 1786 | 1716 | 2088 | 1858 |
| CTGACC | 3594 | 1765 | 3336 | 2377 | 1984 | 1880 | 1824 | 1467 | 1227 | 1192 | 1555 | 1315 |
| CTGACT | 3520 | 2771 | 3164 | 2527 | 1979 | 2079 | 1857 | 1739 | 1328 | 1358 | 1564 | 1444 |
| TTGACG | 3472 | 2796 | 3164 | 2311 | 2051 | 2139 | 1835 | 1500 | 1299 | 1352 | 1443 | 1369 |
| TTTGACC | 2592 | 1922 | 2036 | 1324 | 1460 | 1588 | 1407 | 1234 | 1022 | 992 | 1091 | 1035 |
| TTTGACT | 2896 | 2427 | 2620 | 1866 | 1635 | 1767 | 1787 | 1518 | 1201 | 1244 | 1335 | 1215 |
| GTTGACC | 972 | 884 | 840 | 552 | 466 | 512 | 380 | 439 | 292 | 292 | 448 | 368 |
| GTTGACT | 1304 | 1063 | 1068 | 962 | 862 | 844 | 632 | 759 | 568 | 548 | 728 | 595 |
| TTGACCA | 2068 | 1715 | 1628 | 1192 | 1188 | 1156 | 1155 | 971 | 843 | 808 | 1039 | 971 |
| TTGACCT | 1320 | 1004 | 992 | 816 | 700 | 815 | 588 | 619 | 512 | 484 | 604 | 488 |
| TTGACTT | 2496 | 2407 | 2252 | 1754 | 1455 | 1679 | 1442 | 1507 | 1065 | 1012 | 1321 | 1194 |
| TTGACCTC | 300 | 248 | 232 | 172 | 172 | 224 | 132 | 184 | 116 | 148 | 156 | 124 |
| TTTTCCAC | 752 | 499 | 588 | 420 | 428 | 400 | 376 | 412 | 223 | 288 | 348 | 336 |
| GTACGTAC | 784 | 540 | 640 | 476 | 326 | 340 | 280 | 364 | 188 | 232 | 296 | 271 |
| GGTTAGTTA | 80 | 40 | 44 | 60 | 36 | 20 | 24 | 48 | 40 | 36 | 24 | 28 |

**Table 3.** Disease Resistance-Related Genes found using Genome Wide Computational Analysis

| Motif | Locus ID | Chromosome no. | Gene description |
|---|---|---|---|
| TGACT, TTGACT | LOC_Os04g43360 | 4 | *β-Glycosidase* 14 |
| TGACT, TTGAC, TTGACT | LOC_Os08g09010 | 8 | Germin 8-7 (*ger*6) |
| TTGAC | LOC_Os01g43550 | 1 | *WRKY*12 |
| TTGAC | LOC_Os11g14570 | 11 | *Ankyrin* |
| TTGAC | LOC_Os07g03600 | 7 | *PR*1b |
| TTGAC | LOC_Os04g59370 | 6 | *PR*5 |
| TGACT | LOC_Os01g14140 | 1 | *PR*2 |
| TTGAC | LOC_Os09g26380 | 9 | *Aminotransferase* |
| TTGAC | LOC_Os01g27770 | 1 | Shikimate biosynthesis protein *aroDE* |
| TTGAC | LOC_Os04g32480 | 4 | *TIFY*9 |
| TTGAC | LOC_Os01g54600 | 1 | *WRKY*13 |

**Table 4.** Cis-Elements of selected Disease Resistance Related Genes

| Disease resistance related genes | Factor or Site Name | Loc. (Str.) Sense – (+) Anti-sense - (-) | Signal Sequence |
|---|---|---|---|
| **Germin 8-7 (*ger*6) (LOC_Os08g09010)** | WBOXATNPR1 | 161 (+) | TTGAC |
| | WBOXHVISO1 | 162 (+) | TGACT |
| | WBOXNTERF3 | 162 (+) | TGACY |
| | WBOXHVISO1 | 269 (-) | TGACT |
| | WBOXHVISO1 | 269 (-) | TGACT |
| | WBOXNTERF3 | 269 (-) | TGACY |
| | WBBOXPCWRKY1 | 318 (+) | TTTGACY |
| | WBOXATNPR1 | 319 (+) | TTGAC |
| | WBOXHVISO1 | 320 (+) | TGACT |
| | WBOXNTERF3 | 320 (+) | TGACY |
| | WBBOXPCWRKY1 | 327 (-) | TTTGACY |
| | WBOXHVISO1 | 327 (-) | TGACT |
| | WBOXNTERF3 | 327 (-) | TGACY |
| | WBOXATNPR1 | 328 (-) | TTGAC |
| | WBOXHVISO1 | 333 (-) | TGACT |
| | WBOXNTERF3 | 333 (-) | TGACY |
| | WBBOXPCWRKY1 | 333 (-) | TTTGACY |
| | WBOXATNPR1 | 334 (-) | TTGAC |
| | WBBOXPCWRKY1 | 350 (+) | TTTGACY |
| | WBOXATNPR1 | 351 (+) | TTGAC |
| | WBOXHVISO1 | 352 (+) | TGACT |
| | WBOXNTERF3 | 352 (+) | TGACY |
| | WBOXHVISO1 | 483 (-) | TGACT |
| | WBOXNTERF3 | 483 (-) | TGACY |

| | | | |
|---|---|---|---|
| | WBBOXPCWRKY1 | 483(-) | TTTGACY |
| | WBOXATNPR1 | 484 (-) | TTGAC |
| | WBOXNTERF3 | 622 (+) | TGACY |
| | WBOXHVISO1 | 735 (+) | TGACT |
| | WBOXNTERF3 | 735 (+) | TGACY |
| | WBOXATNPR1 | 792 (-) | TTGAC |
| | WBOXATNPR1 | 920 (+) | TTGAC |
| | WBOXHVISO1 | 921 (+) | TGACT |
| | WBOXNTERF3 | 921 (+) | TGACY |
| **β-*glycosidase*14 (LOC_Os04g43360)** | WBOXNTERF3 | 89 (+) | TGACY |
| | WBOXATNPR1 | 133 (-) | TTGAC |
| | WBOXHVISO1 | 452 (-) | TGACT |
| | WBOXNTERF3 | 452 (-) | TGACY |
| | WBOXNTERF3 | 461 (+) | TGACY |
| | WBOXATNPR1 | 528 (-) | TTGAC |
| | WBBOXPCWRKY1 | 764 (+) | TTTGACY |
| | WBOXATNPR1 | 765 (+) | TTGAC |
| | WBOXNTERF3 | 766 (+) | TGACY |
| | WBOXATNPR1 | 797 (+) | TTGAC |
| | WBOXATNPR1 | 811 (+) | TTGAC |
| | WBOXNTERF3+ | 812 (+) | TGACY |
| | WBOXATNPR1 | 45 (+) | TTGAC |
| | WBOXATNPR1 | 373 (+) | TTGAC |
| ***Aminotransferase* (LOC_Os09g26380)** | WBBOXPCWRKY1 | 372 (+) | TTTGACY |
| | WBOXNTERF3 | 374 (+) | TGACY |
| | WBOXHVISO1 | 644 (-) | TGACT |
| | WBOXNTERF3 | 644 (-) | TGACY |
| | WBOXATNPR1 | 758 (+) | TTGAC |

| | | | |
|---|---|---|---|
| | WBOXATNPR1 | 858 (+) | TTGAC |
| ***WRKY*12** **(LOC_Os01g43550)** | WBOXNTERF3 | 3 (-) | TGACY |
| | WBOXNTCHN48 | 3(-) | CTGACY |
| | WBOXATNPR1 | 250 (+) | TTGAC |
| | WBOXHVISO1 | 265 (-) | TGACT |
| | WBOXNTERF3 | 265 (-) | TGACY |
| | WBOXNTCHN48 | 265 (-) | CTGACY |
| | WBOXHVISO1 | 336 (-) | TGACT |
| | WBOXNTERF3 | 336 (-) | TGACY |
| | WBOXATNPR1 | 337 (-) | TTGAC |
| | WBOXATNPR1 | 497 (+) | TTGAC |
| | WBBOXPCWRKY1 | 511 (-) | TTTGACY |
| | WBOXHVISO1 | 511 (-) | TGACT |
| | WBOXNTERF3 | 511 (-) | TGACY |
| | WBOXATNPR1 | 512 (-) | TTGAC |
| | WBOXATNPR1 | 594 (+) | TTGAC |
| | WBBOXPCWRKY1 | 771 (+) | TTTGACY |
| | WBOXATNPR1 | 772 (+) | TTGAC |
| | WBOXHVISO1 | 773 (+) | TGACT |
| | WBOXNTERF3 | 773 (+) | TGACY |
| | WBOXHVISO1 | 903 (-) | TGACT |
| | WBOXNTERF3 | 903 (-) | TGACY |
| | WBOXATNPR1 | 904 (-) | TTGAC |
| | WBOXATNPR1 | 969 (-) | TTGAC |
| ***WRKY*13** **(LOC_Os01g54600)** | WBOXNTERF3 | 841 (-) | TGACY |
| | WBOXNTCHN48 | 841 (-) | CTGACY |
| | WBOXATNPR1 | 722 (-) | TTGAC |
| ***TIFY9*** **(LOC_Os04g32480)** | TGACY | 361(+) | TGACC |

| | | | |
|---|---|---|---|
| ***Ankyrin* (LOC_Os11g14570)** | WBOXNTCHN48 | 414 (+) | CTGACY |
| | WBOXHVISO1 | 415 (+) | TGACT |
| | WBOXNTERF3 | 415 (+) | TGACY |
| | WBOXHVISO1 | 425 (+) | TGAC |
| | WBOXNTERF3 | 425 (+) | TGACY |
| | WBOXNTCHN48 | 471 (+) | CTGACY |
| | WBOXHVISO1 | 472 (+) | TGACT |
| | WBOXNTERF3 | 472 (+) | TGACY |
| | WBOXHVISO1 | 489 (+) | TGACT |
| | WBOXNTERF3 | 489 (+) | TGACY |
| | WBOXATNPR1 | 666 (-) | TTGAC |
| **Shikimate biosynthesis protein (LOC_Os01g27770)** | WBOXATNPR1 | 416 (+) | TTGAC |
| | WBOXHVISO1 | 417 (+) | TGACT |
| | WBBOXPCWRKY1 | 415 (+) | TTTGACY |
| | WBOXNTERF3 | 417 (+) | TGACY |
| | WBOXATNPR1 | 569 (+) | TTGAC |
| | WBOXATNPR1 | 625 (+) | TTGAC |
| | WBOXATNPR1 | 659 (+) | TTGAC |
| | WBOXNTERF3 | 721 (+) | TGACY |
| ***PR*1b (LOC_Os07g03600)** | WBOXATNPR1 | 51 (+) | TTGAC |
| | WBOXNTERF3 | 52 (+) | TGACY |
| | WBOXHVISO1 | 98 (-) | TGACT |
| | WBOXNTERF3 | 98 (-) | TGACY |
| | WBBOXPCWRKY1 | 98 (-) | TTTGACY |
| | WBOXATNPR1 | 99 (-) | TTGAC |
| | WBOXATNPR1 | 268 (+) | TTGAC |
| | WBOXATNPR1 | 277 (+) | TTGAC |
| | WBOXHVISO1 | 278 (+) | TGACT |

| | | | |
|---|---|---|---|
| | WBOXNTERF3 | 278 (+) | TGACY |
| | WBBOXPCWRKY1 | 295 (+) | TTTGACY |
| | WBOXATNPR1 | 296 (+) | TTGAC |
| | WBOXNTERF3 | 297 (+) | TGACY |
| | WBOXNTERF3 | 383 (+) | TGACY |
| | WBOXATNPR1 | 413 (-) | TTGAC |
| | WBOXHVISO1 | 431 (-) | TGACT |
| | WBOXNTERF3 | 431 (-) | TGACY |
| | WBOXATNPR1 | 432 (-) | TTGAC |
| | WBOXATNPR1 | 441 (-) | TTGAC |
| | WBOXNTERF3 | 942 (-) | TGACY |
| | WBOXNTCHN48 | 942 (-) | CTGACY |
| **PR5 (LOC_Os04g59370)** | WBOXATNPR1 | 246 (+) | TTGAC |
| | WBOXHVISO1 | 247 (+) | TGACT |
| | WBOXNTERF3 | 247 (+) | TGACY |
| | WBBOXPCWRKY1 | 385 (-) | TTTGACY |
| | WBOXHVISO1 | 385 (-) | TGACT |
| | WBOXNTERF3 | 385 (-) | TGACY |
| | WBOXATNPR1 | 386 (-) | TTGAC |
| | WBOXNTERF3 | 422 (-) | TGACY |
| | WBOXATNPR1 | 523 (-) | TTGAC |
| | WBOXHVISO1 | 71 (+) | TGACT |
| | WBOXNTERF3 | 71 (+) | TGACY |
| | WBOXNTCHN48 | 183 (+) | CTGACY |
| | WBOXHVISO1 | 184 (+) | TGACT |
| | WBOXNTERF3 | 184 (+) | TGACY |
| | WBOXHVISO1 | 248 (-) | TGACT |
| | WBOXNTERF3 | 248 (-) | TGACY |

| | | | |
|---|---|---|---|
| ***PR*2 (LOC_Os01g14140)** | WBOXHVISO1 | 322 (+) | TGACT |
| | WBOXNTERF3 | 322 (+) | TGACY |
| | WBOXATNPR1 | 566 (-) | TTGAC |
| | WBBOXPCWRKY1 | 640 (+) | TTTGACY |
| | WBOXATNPR1 | 641 (+) | TTGAC |
| | WBOXHVISO1 | 642 (+) | TGACT |
| | WBOXNTERF3 | 642 (+) | TGACY |
| | WBOXNTERF3 | 662 (-) | TGACY |
| | WBOXNTERF3 | 707 (-) | TGACY |
| | WBOXATNPR1 | 708 (-) | TTGAC |
| | WBOXHVISO1 | 860 (-) | TGACT |
| | WBOXNTERF3 | 860 (-) | TGACY |
| | WBOXNTERF3 | 874 (+) | TGACY |

**Table 5.** Interaction energies of Protein–DNA Complex of WRKY13 TF with Promoters of Selected Rice Genes.

| Molecule | Interaction energy | Vander waals energy | Electrostatic energy | Interacting residues with the binding motif |
|---|---|---|---|---|
| ***β-Glycosidase*14** | -163.41 | -93.58 | -6.24 | GLN67, PRO6, TYR25 |
| **Germin 8-7 (*ger*6)** | -226.87 | -158.85 | -8.04 | THR52 |
| ***WRKY*12** | -170.44 | -100.77 | -6.39 | TYR25 |
| ***Ankyrin*** | -82.12 | -10.87 | -4.82 | PR06, SER7 |
| ***PR*1b** | -240.73 | -170.5 | -5.91 | ARG46 |
| ***PR*5** | -304.0 | -233.53 | -5.59 | SER7 |
| ***PR*2** | -94.81 | -25.19 | -6.45 | GLN67, TYR, THR52 |
| ***Aminotransferase*** | -62.37 | -7.71 | -5.99 | ARG41, ARG31, ARG41, GLN17 |
| **Shikimate biosynthesis protein (*aro*DE)** | -82.98 | -13.83 | -6.62 | TYR25, SER7 |
| ***TIFY9*** | -135.2 | -45.7 | -6.41 | LEU149, LYS133 |
| ***WRKY*13** | -101.6 | -30.67 | -5.14 | THR52, ARG46 |

**Table 6.** Statistical analysis of MD simulation for Protein-DNA complexes.

| MOL | RMSD | RMSF | ROG | SASA | H-bond |
|---|---|---|---|---|---|
| ***Aminotransferase*** | 0.0004979 to 0.2977097 **Avg** 0.198 | 0.0432 to 0.3102 **Avg** 0.093 | 1 to 1.34398 **Avg** 1.276 | 45.006 to 53.337 **Avg**48.522 | 0 to 1 **Avg** 0.012 |
| ***Ankyrin*** | 0.0004859 to 0.3290237 **Avg** 0.221 | 0.0551 to 0.4116 **Avg** 0.125 | 1.23477 to 1.37713 **Avg** 1.307 | 45.827 to 54.147 **Avg** 50.018 | 0 to 4 **Avg** 1.456 |
| ***aroDE*** | 0.0005053 to 0.643639 **Avg** 0.358 | 0.0498 to 0.812 **Avg** 0.167 | 1.24185 to 1.41514 **Avg** 1.315 | 46.465 to 56.118 **Avg** 51.076 | 0 to 3 **Avg** 1.5149 |
| **β-*glucosidase*** | 0.0005143 to 0.3160566 **Avg** 0.200 | 0.0458 to 0.3461 **Avg** 0.099 | 1.24155 to 1.40038 **Avg** 1.313 | 45.145 to 54.274 **Avg** 49.925 | 0 to 3 **Avg** 0.877 |
| ***ger*6** | 0.0005012 to 0.3396654 **Avg** 0.238 | 0.0475 to 0.2406 **Avg** 0.117 | 1.21619 to 1.33635 **Avg** 1.275 | 46.38 to 55.606 **Avg** 49.946 | 0 to 3 **Avg** 0.385 |
| ***PR*1b** | 0.0005075 to 0.4030854 **Avg** 0.287 | 0.0524 to 0.5678 **Avg** 0.114 | 1.25069 to 1.38367 **Avg** 1.328 | 46.733 to 54.591 **Avg** 50.344 | 0 to 1 **Avg** 0.003 |
| ***PR*5** | 0.0004924 to 0.3880028 **Avg** 0.251 | 0.0501 to 0.6167 **Avg** 0.113 | 1.22892 to 1.36953 **Avg** 1.294 | 46.433 to 53.646 **Avg** 49.792 | 0 to 4 **Avg** 1.1548 |
| ***PR*2** | 0.0004991 to 0.3453989 **Avg** 0.236 | 0.0515 to 0.3633 **Avg** 0.108 | 1.2269 to 1.33269 **Avg** 1.276 | 44.03 to 53.974 **Avg** 47.834 | 0 to 10 **Avg** 4.528 |
| ***TIFY*9** | 0.0004938 to 0.4217459 **Avg** 0.329 | 0.0565 to 0.5734 **Avg** 0.131 | 1.22131 to 1.36528 **Avg** 1.310 | 45.387 to 52.638 **Avg** 49.097 | 0 to 3 **Avg** 1.449 |
| ***WRKY*12** | 0.0005043 to 0.4386873 **Avg** 0.273 | 0.0656 to 0.46 **Avg** 0.153 | 0.644672 to 1.40038 **Avg** 1.389 | 45.708 to 57.744 **Avg** 50.561 | 0 to 3 **Avg** 0.656 |
| ***WRKY*13** | 0.005183 to 0.357719 **Avg** 0.245 | 0.0469 0.4052 **Avg** 0.121 | 1.24637 to 1.3544 **Avg** 1.298 | 46.131 to 56.269 **Avg** 51.252 | 0 to 4 **Avg** 2.469 |
