## Supplementary figures and images for "Rice WRKY13 TF protein binds to motifs in the promoter region to regulate downstream disease resistance-related genes"

### Supplementary figure 2.1.jpg

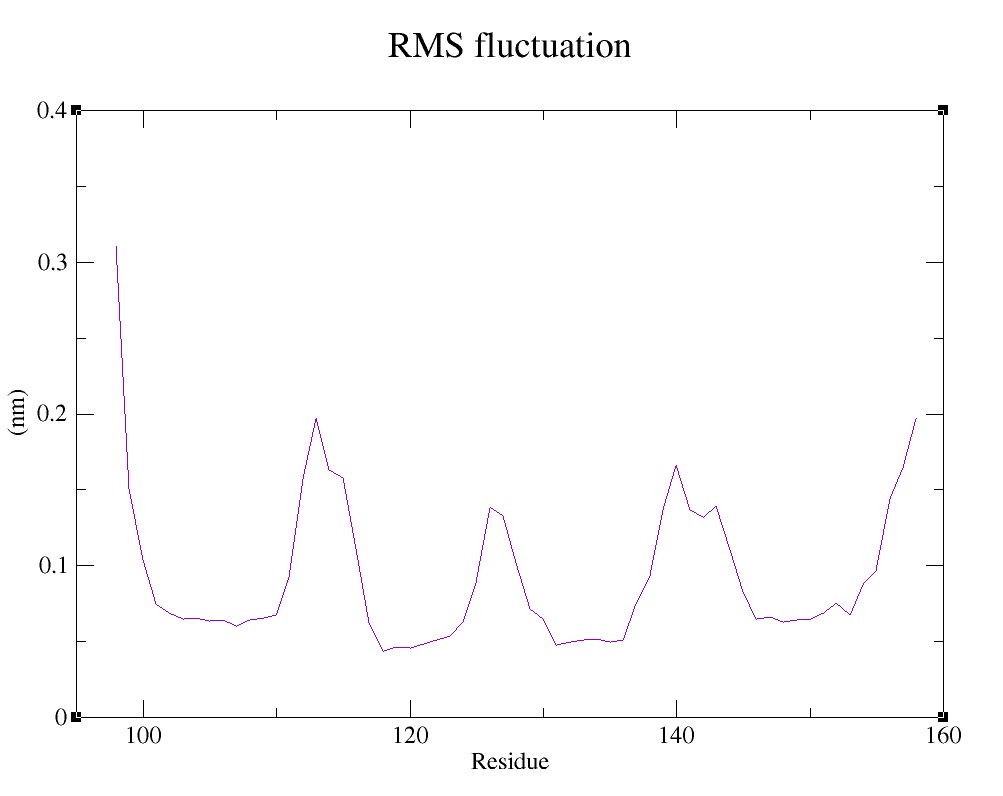

### Supplementary figure 2.2.jpg

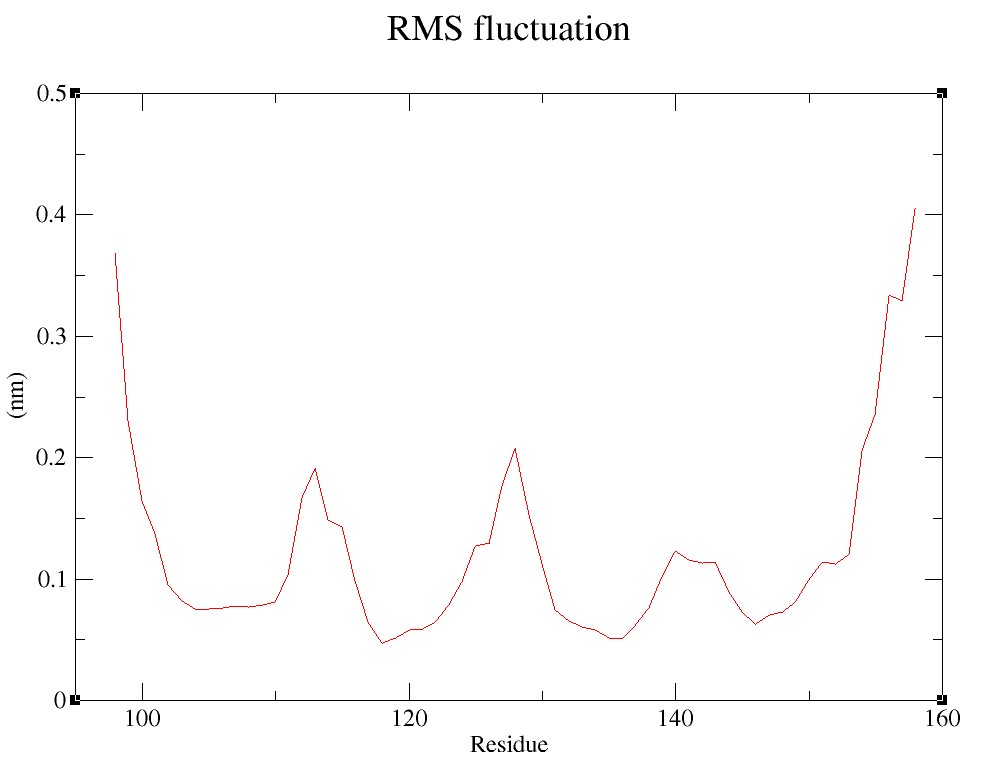

### Supplementary figure 2.3.jpg

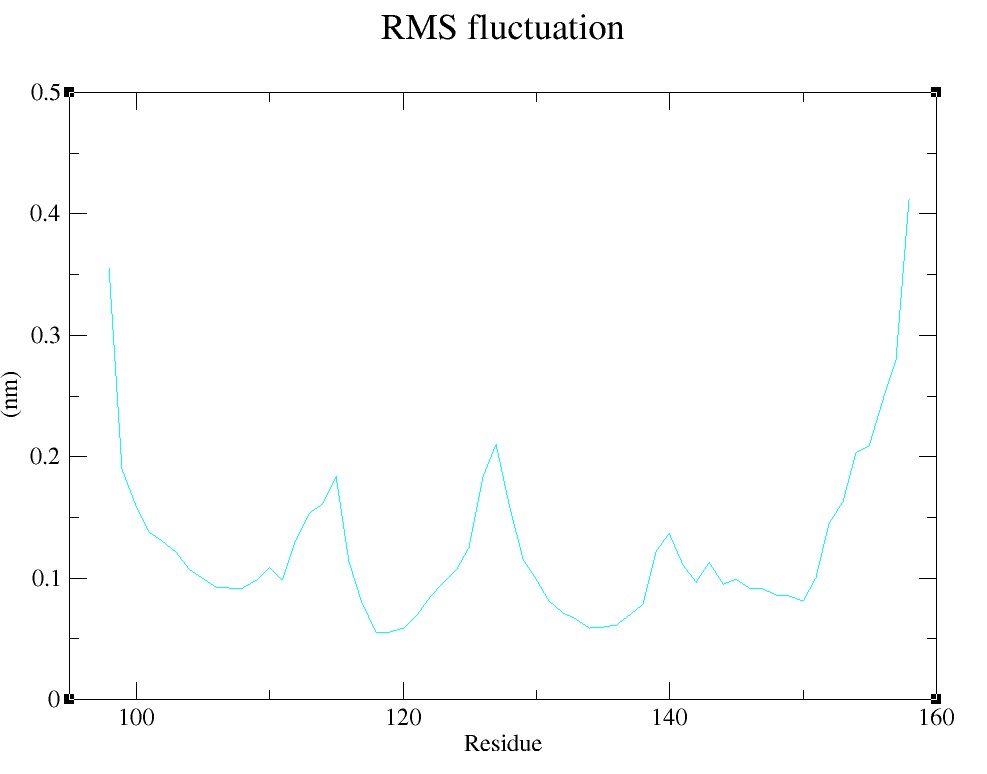

### Supplementary figure 2.4.jpg

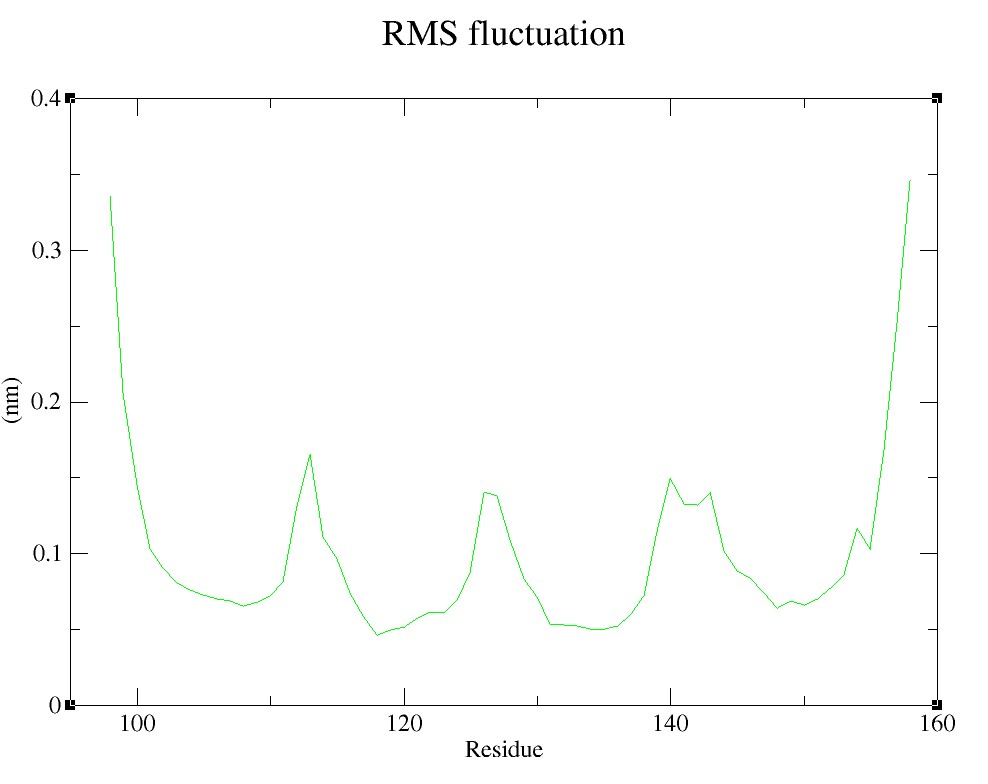

### Supplementary figure 2.5.jpg

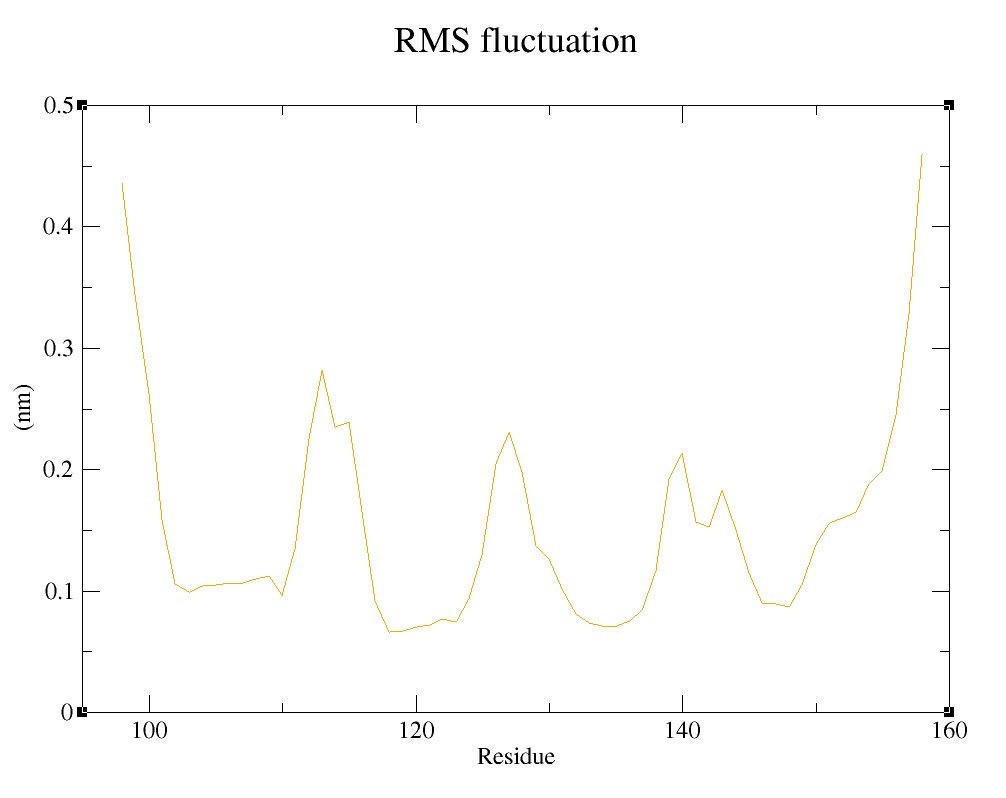

### Supplementary figure 2.6.jpg

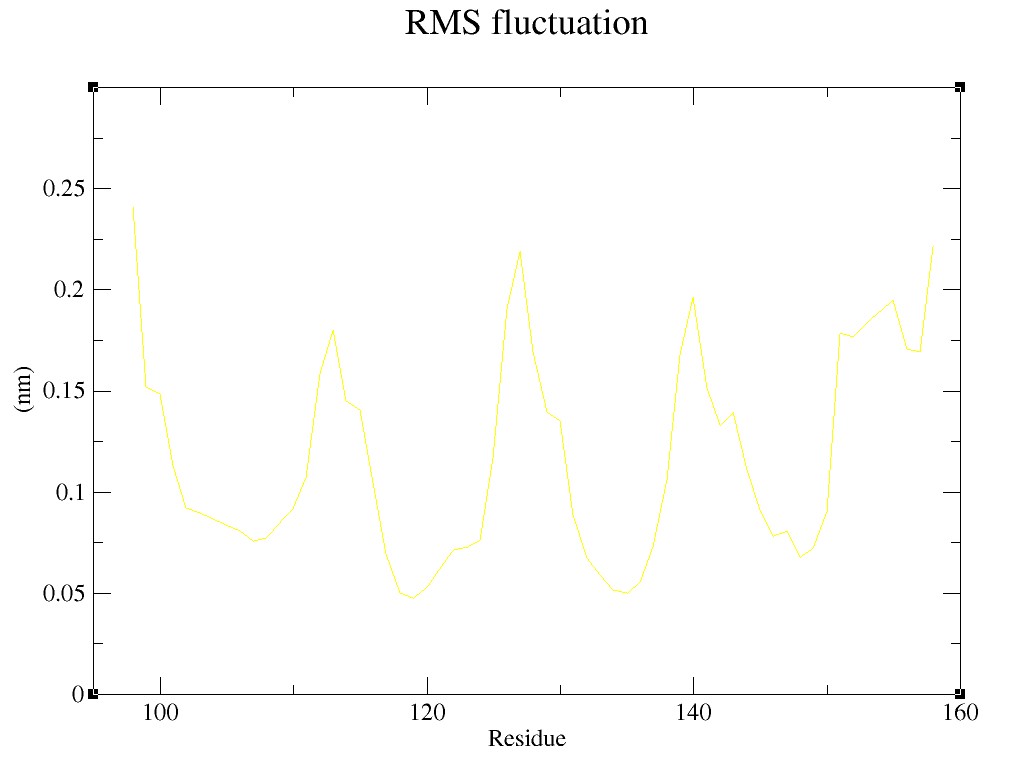

### Supplementary figure 2.7.jpg

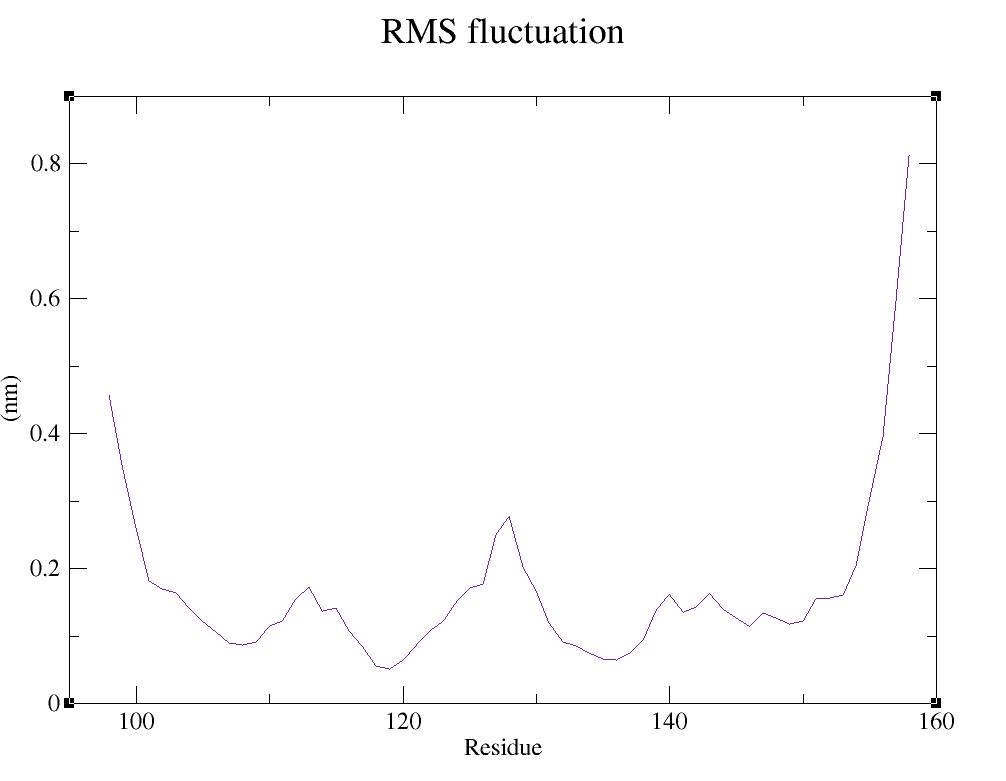

### Supplementary figure 2.8.jpg

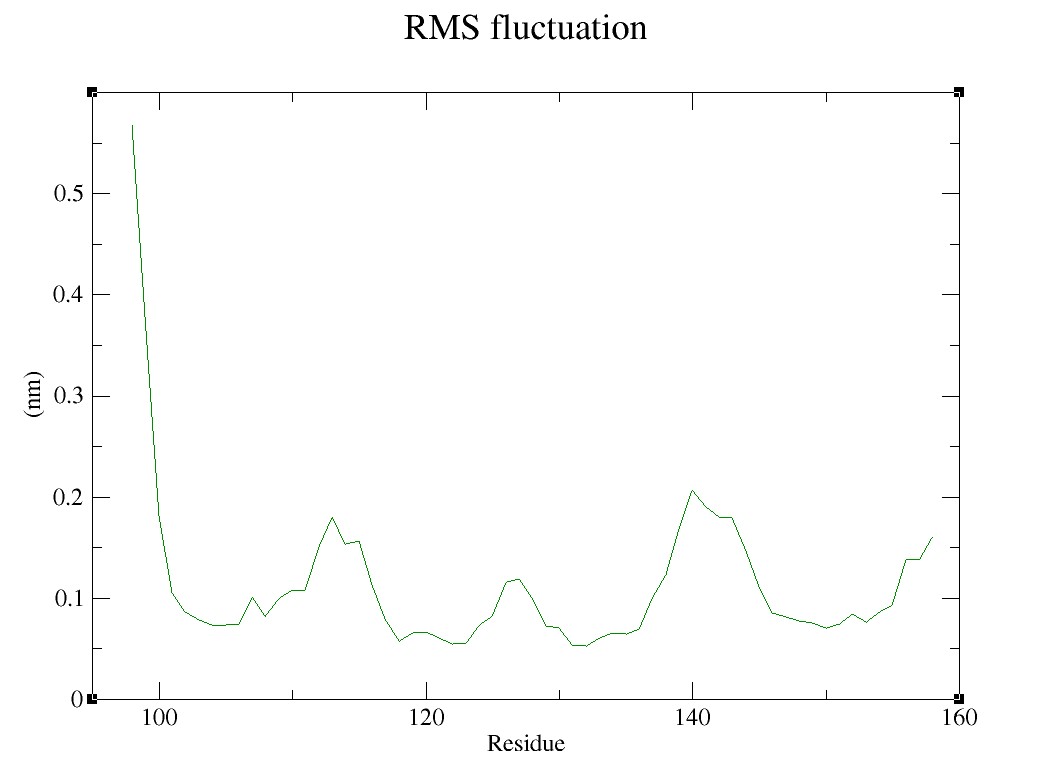

### Supplementary figure 2.9.jpg

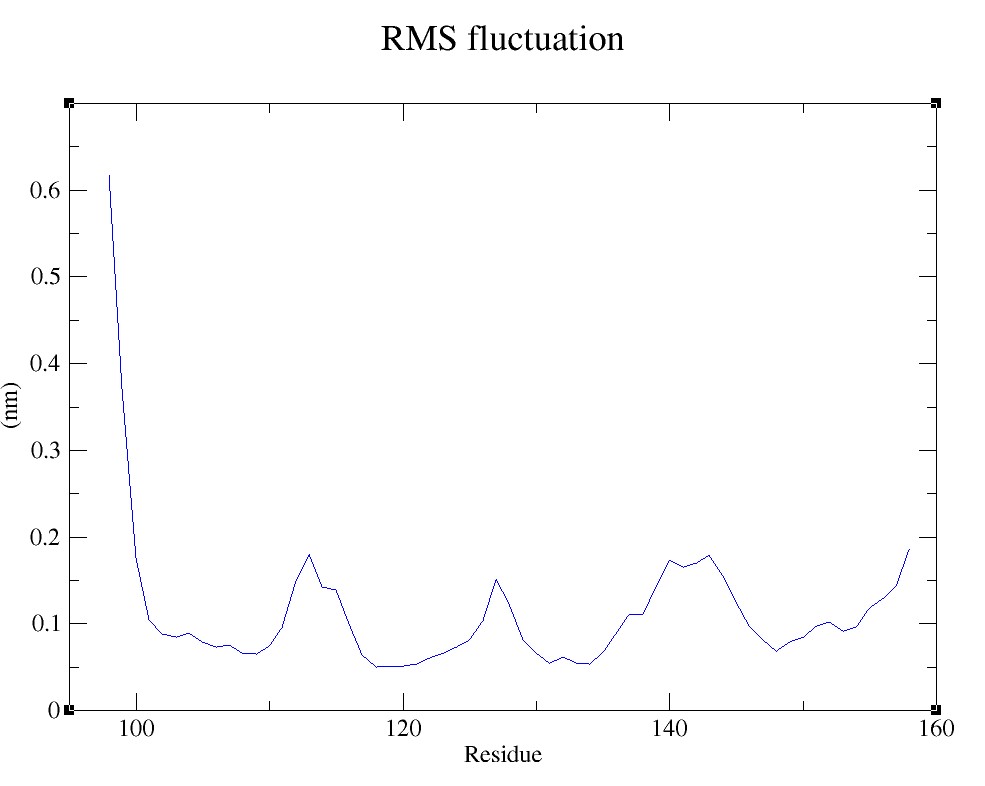

### Supplementary figure 2.10.jpg

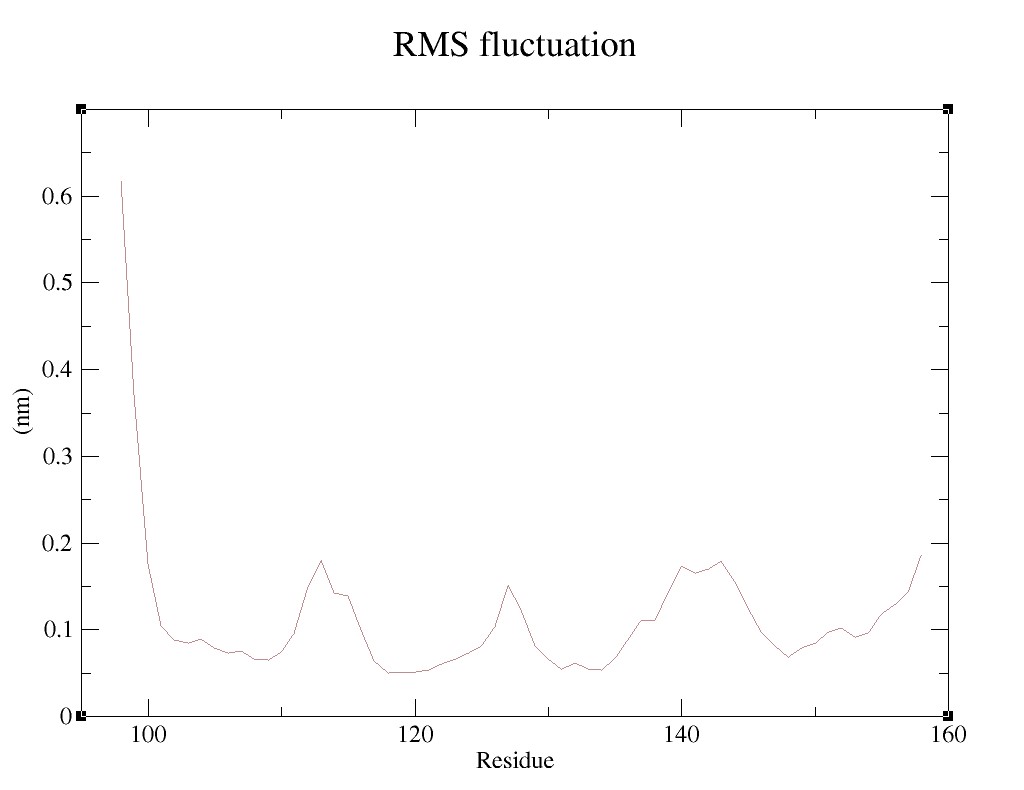

### Supplementary figure 2.11.jpg

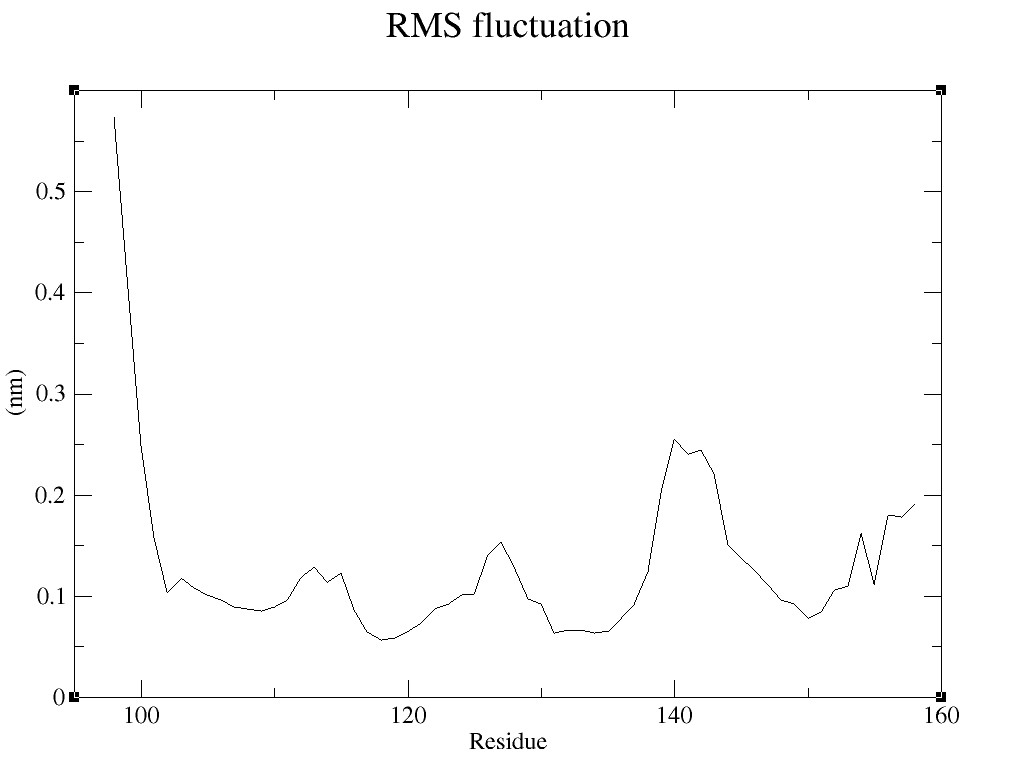

### Supplementary figure 3.1.jpg

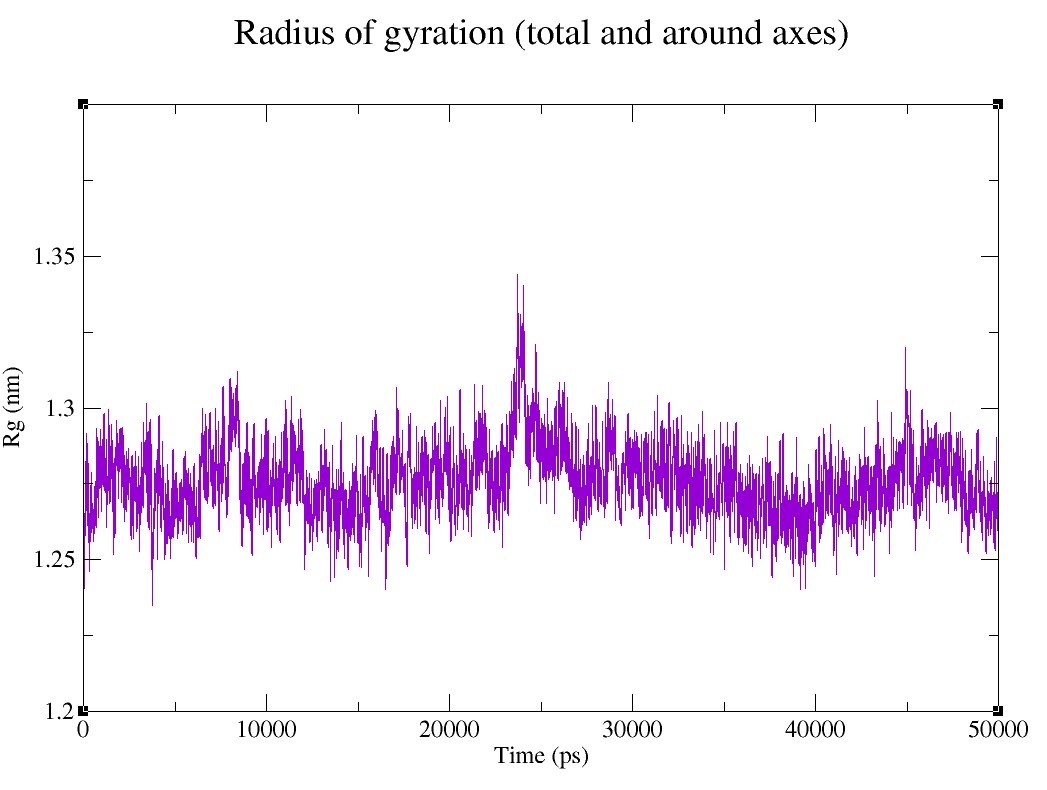

### Supplementary figure 3.2.jpg

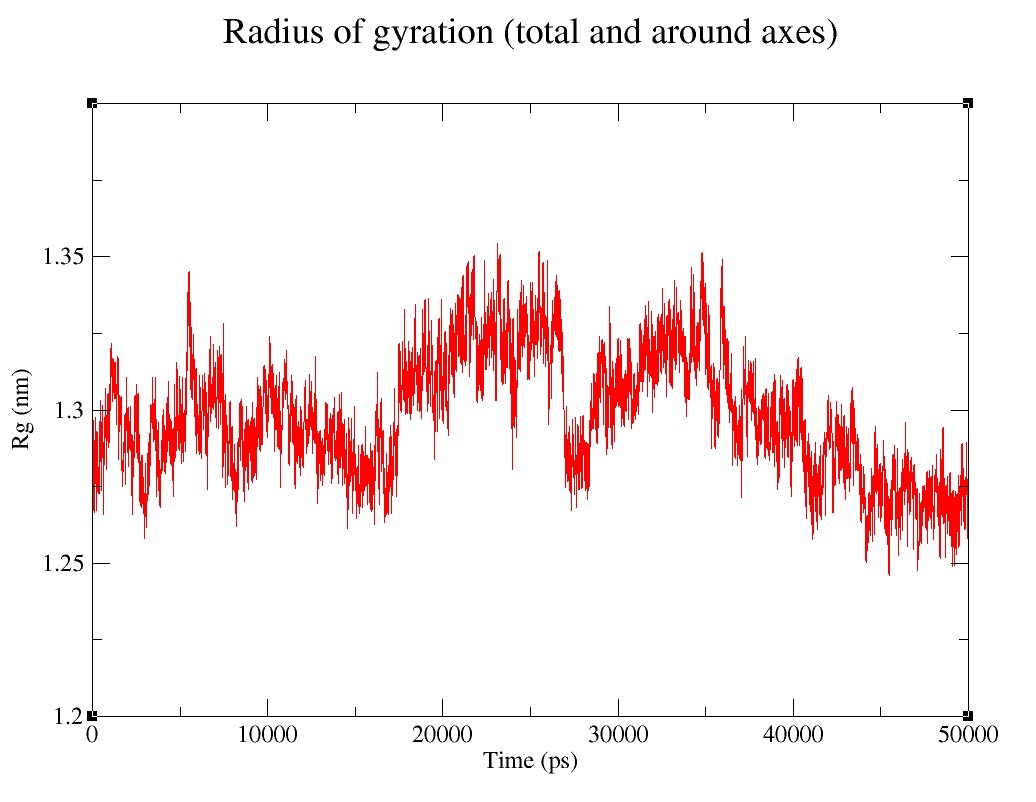

### Supplementary figure 3.3.jpg

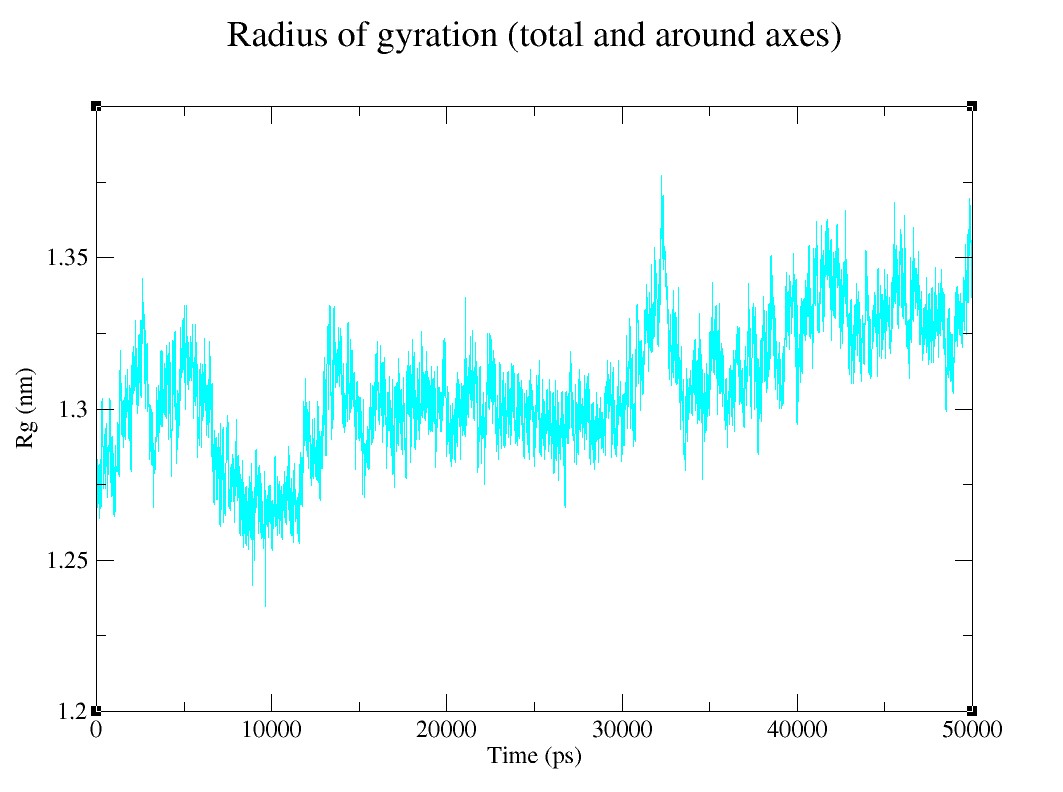

### Supplementary figure 3.4.jpg

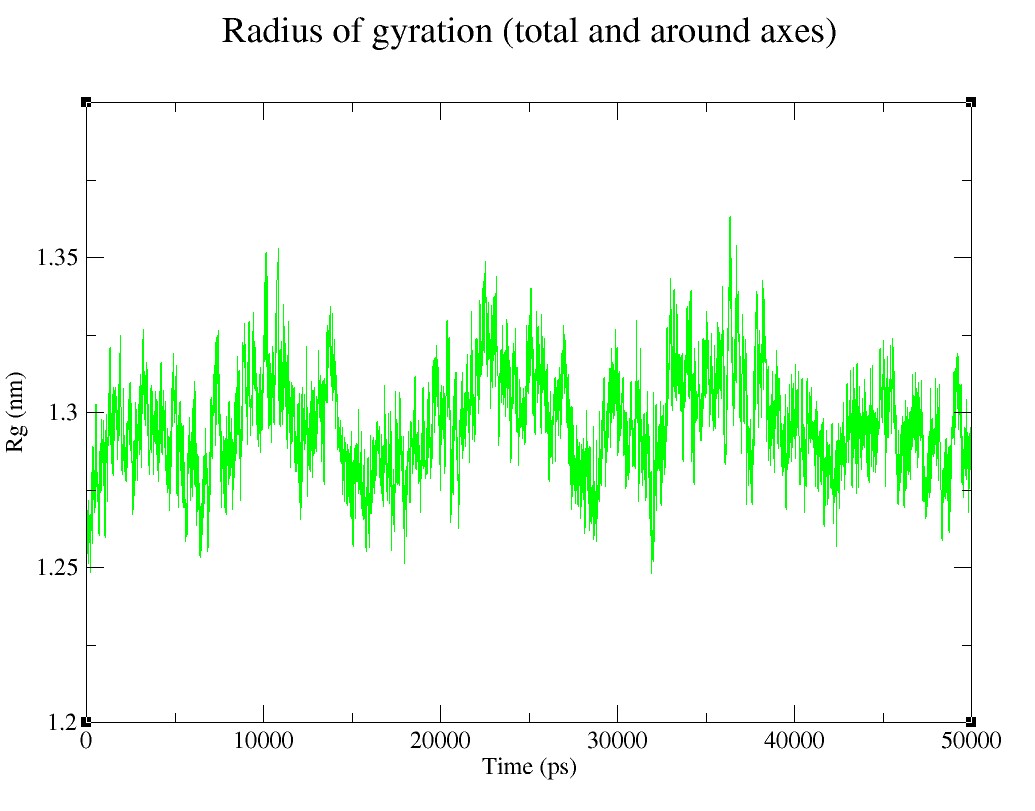

### Supplementary figure 3.5.jpg

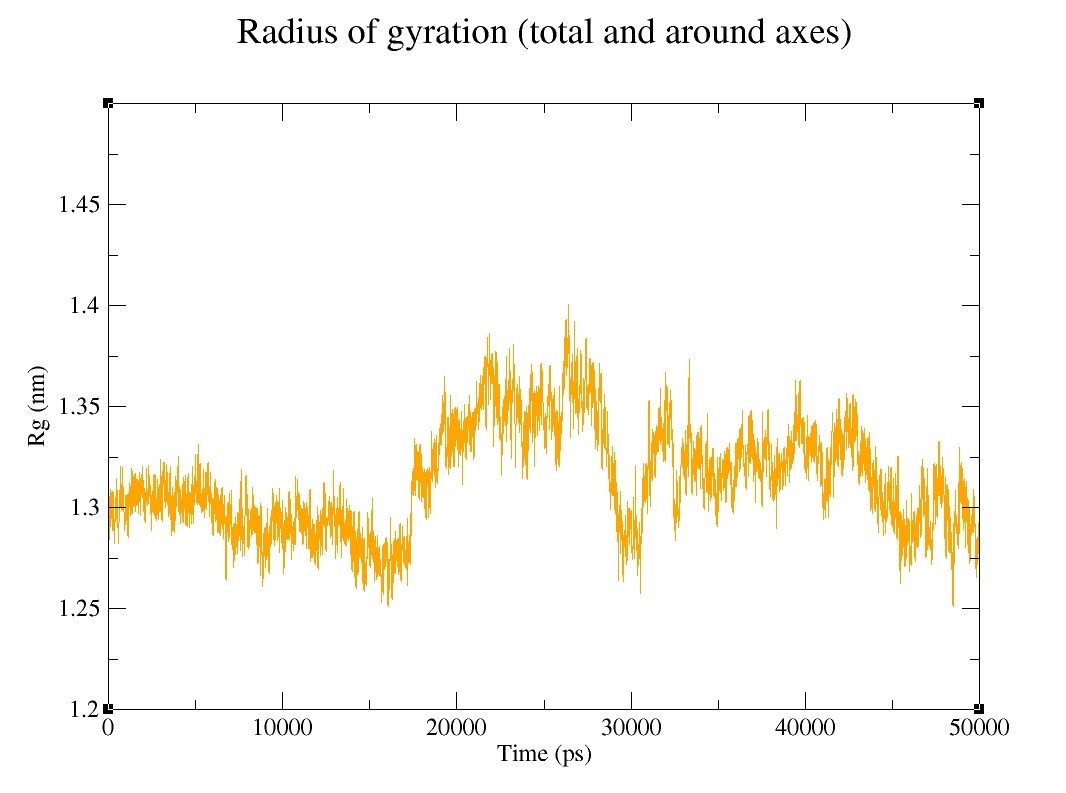

### Supplementary figure 3.6.jpg

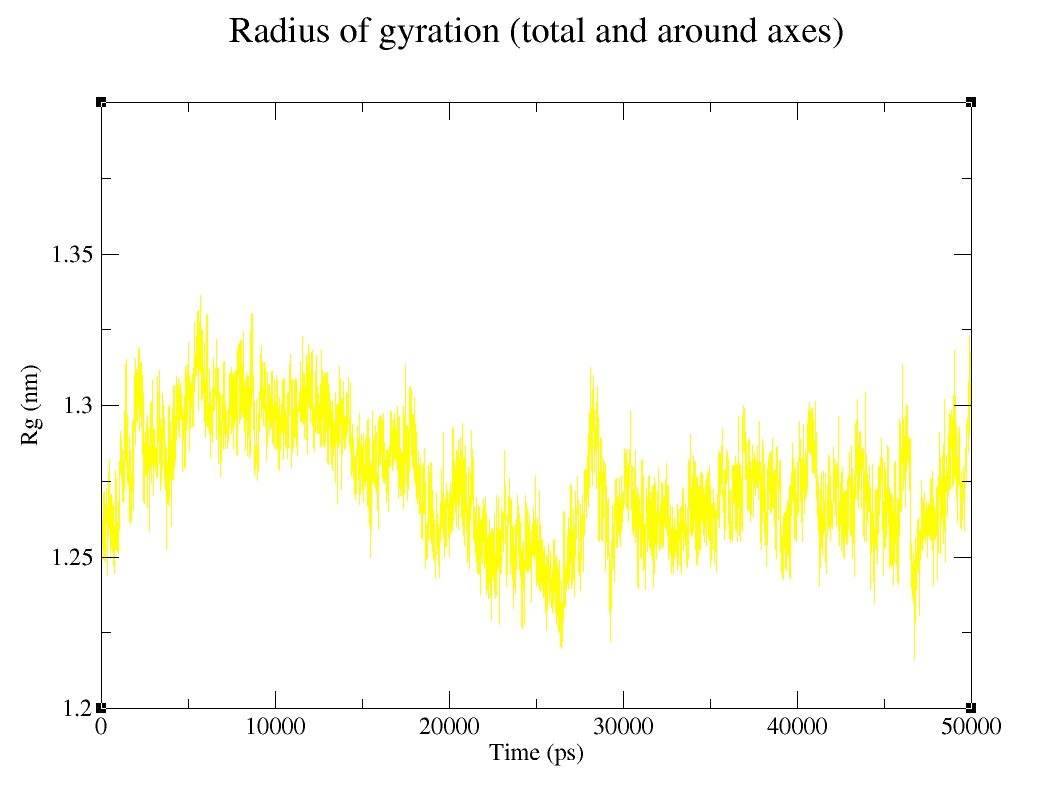

### Supplementary figure 3.7.jpg

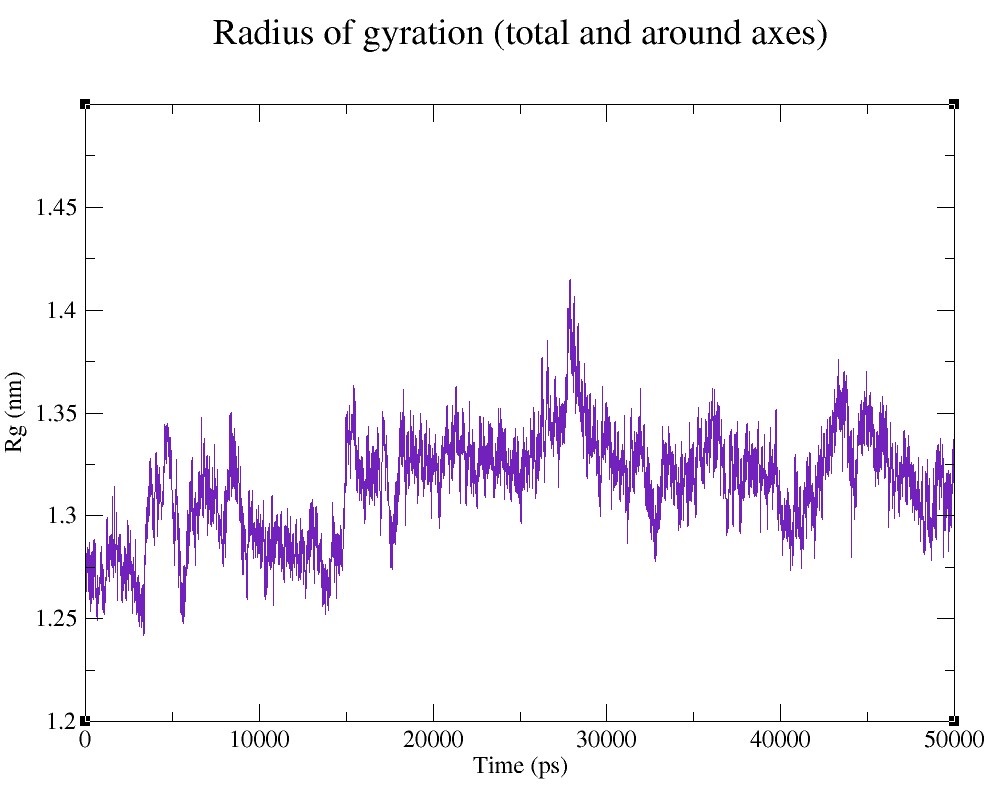

### Supplementary figure 3.8.jpg

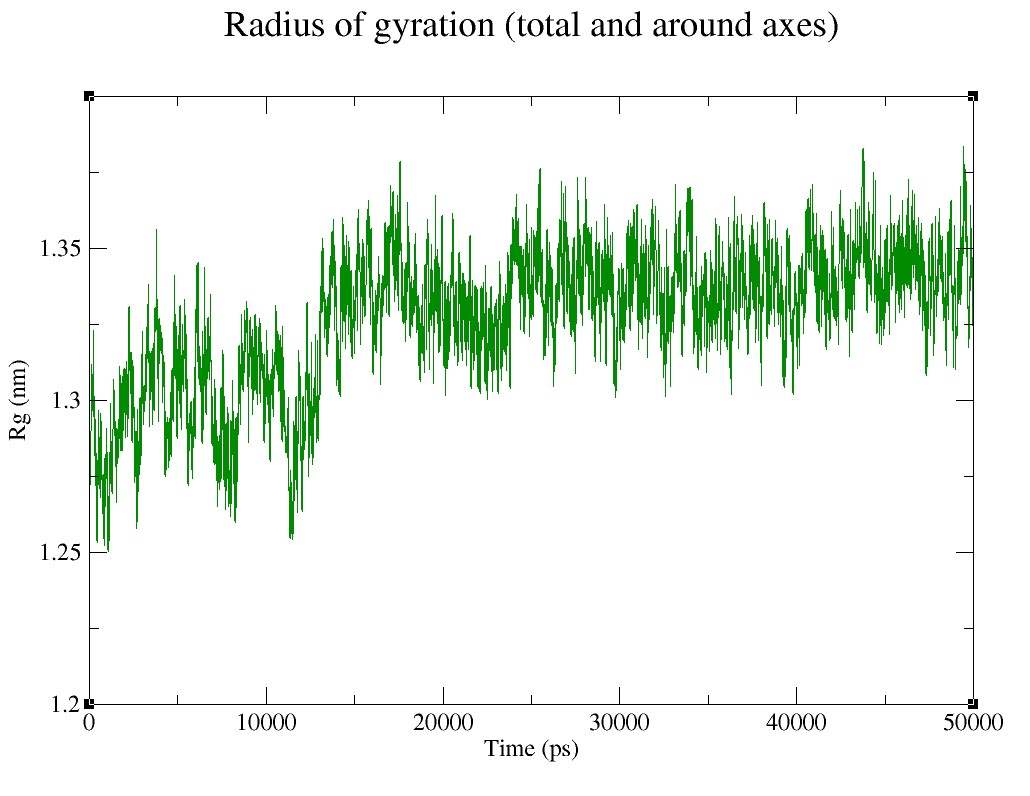

### Supplementary figure 3.9.jpg

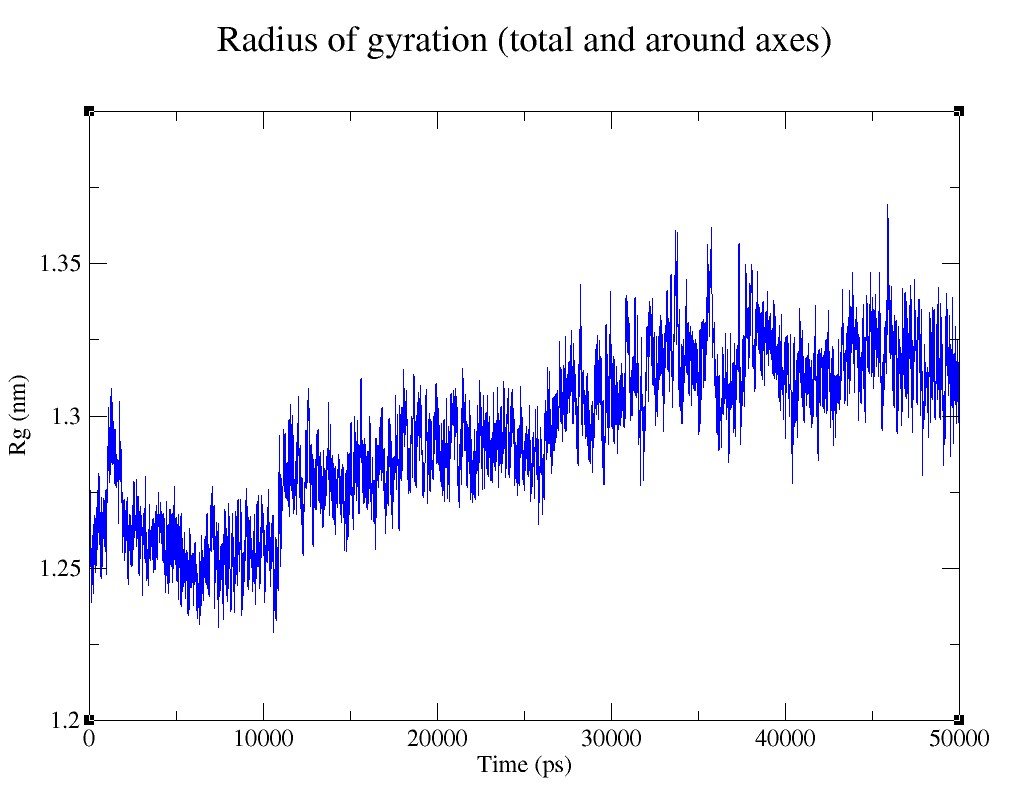

### Supplementary figure 3.10.jpg

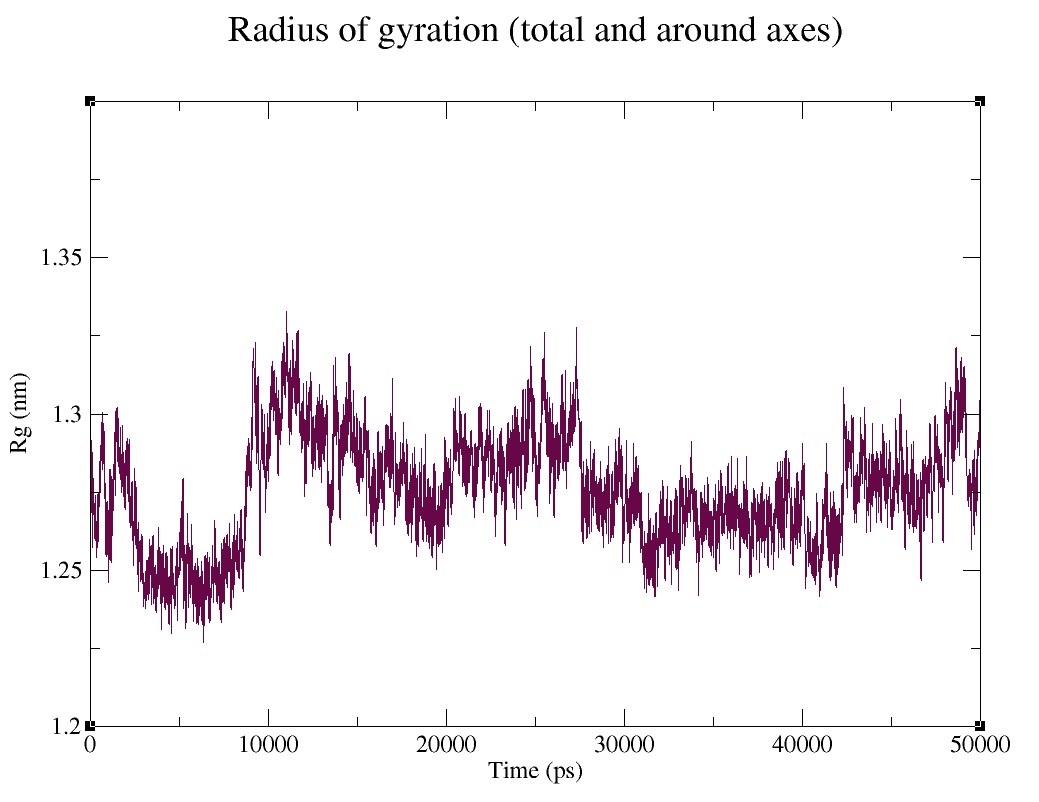

### Supplementary figure 3.11.jpg

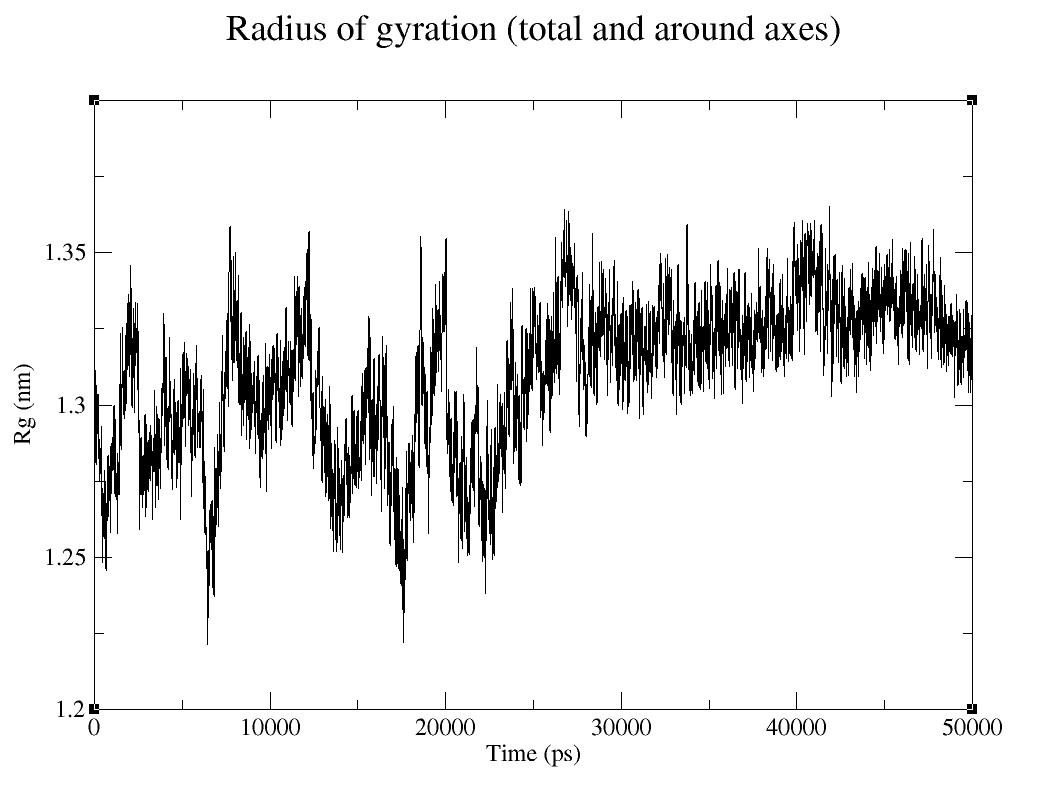

### Supplementary figure 4.1.jpg

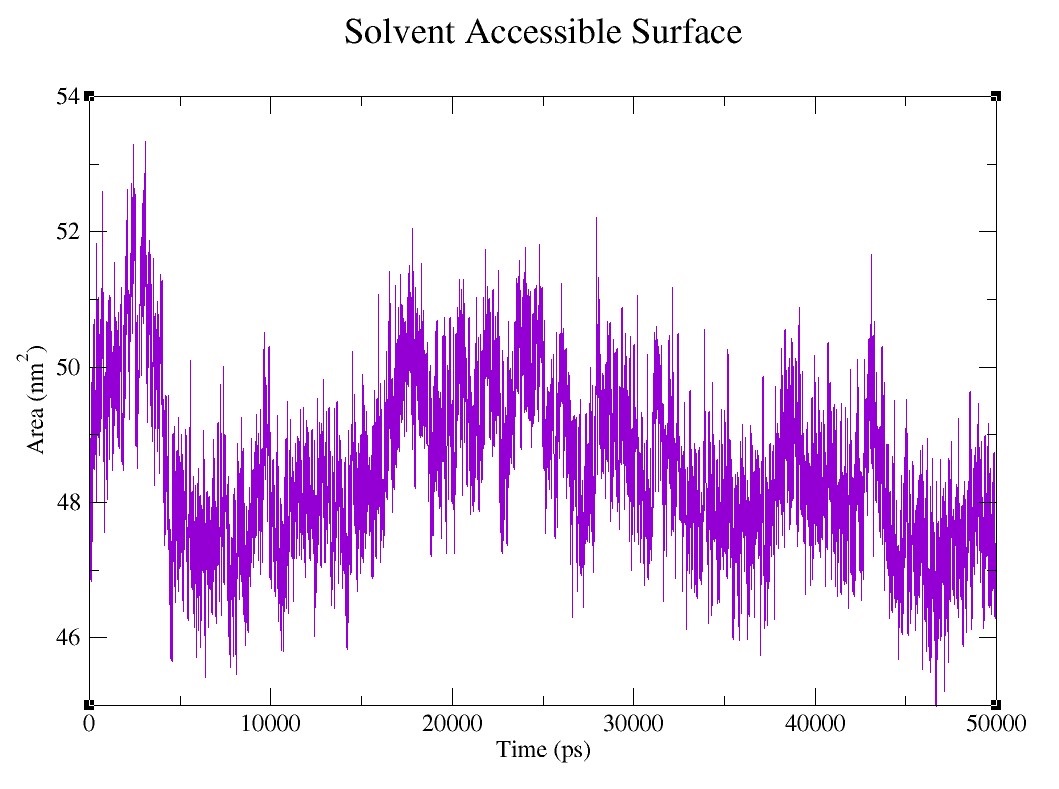

### Supplementary figure 4.2.jpg

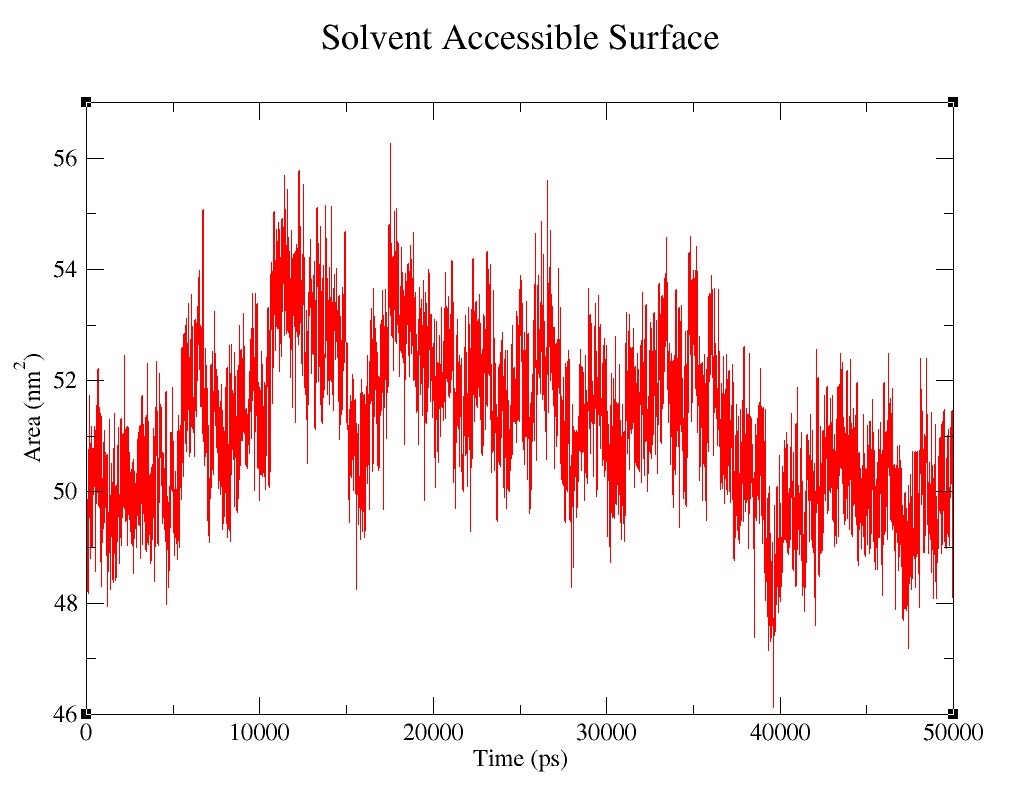

### Supplementary figure 4.3.jpg

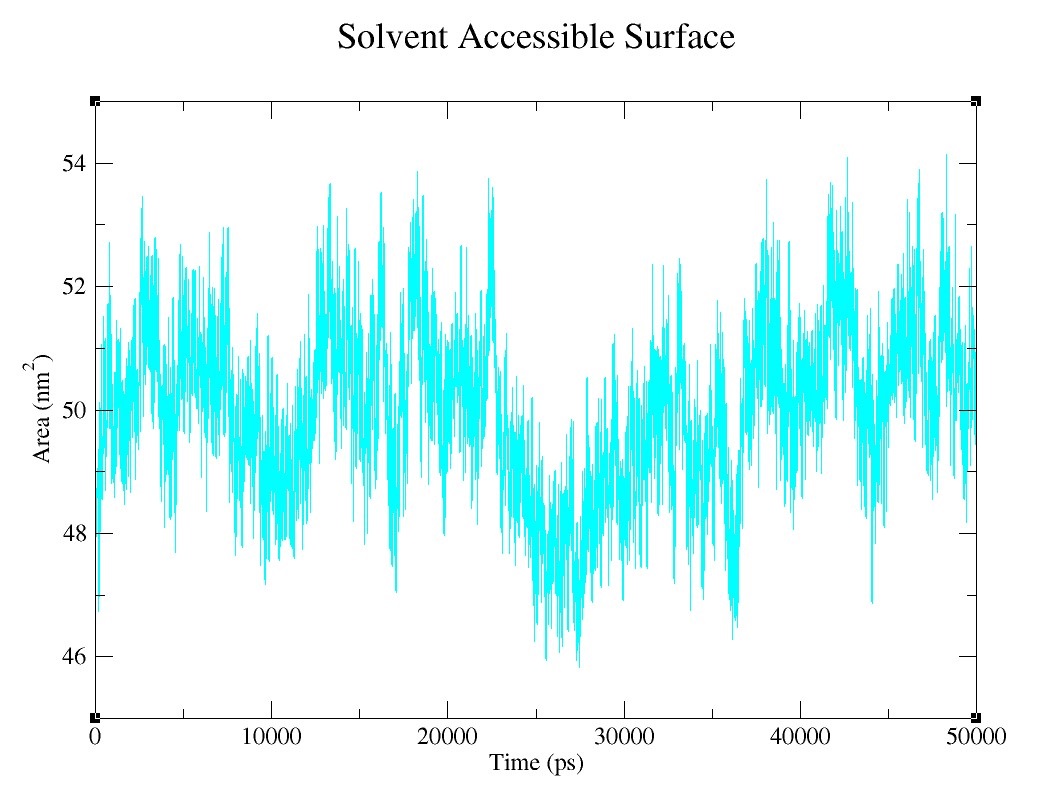

### Supplementary figure 4.4.jpg

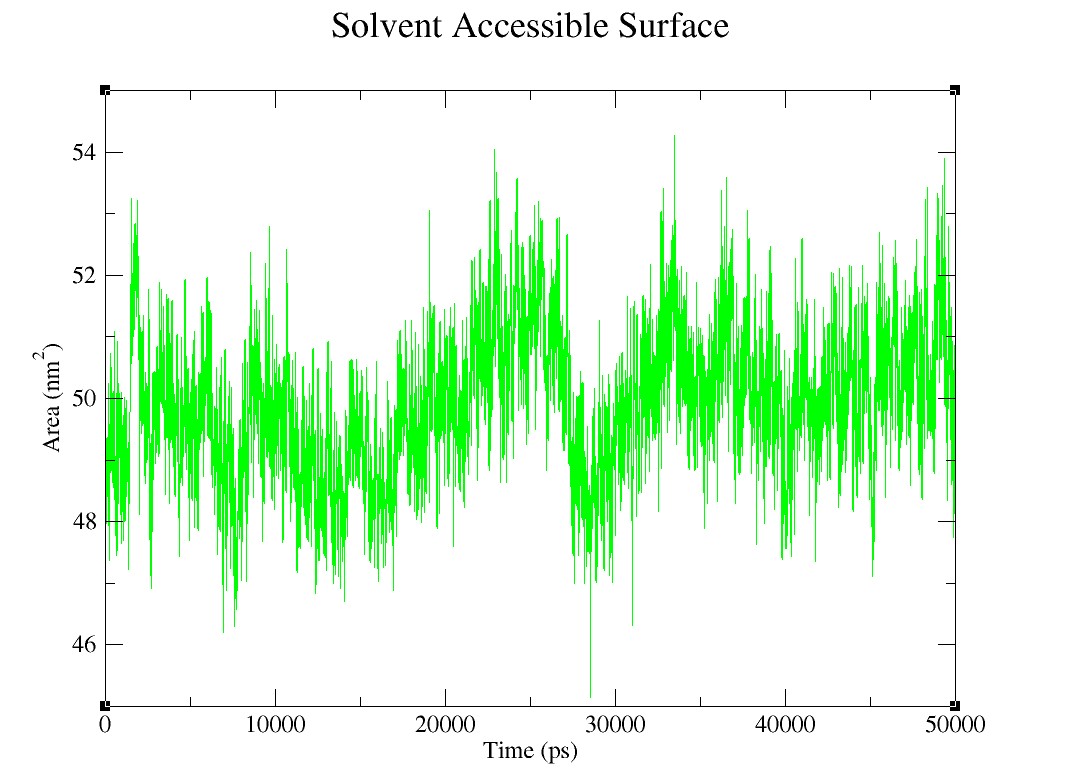

### Supplementary figure 4.5.jpg

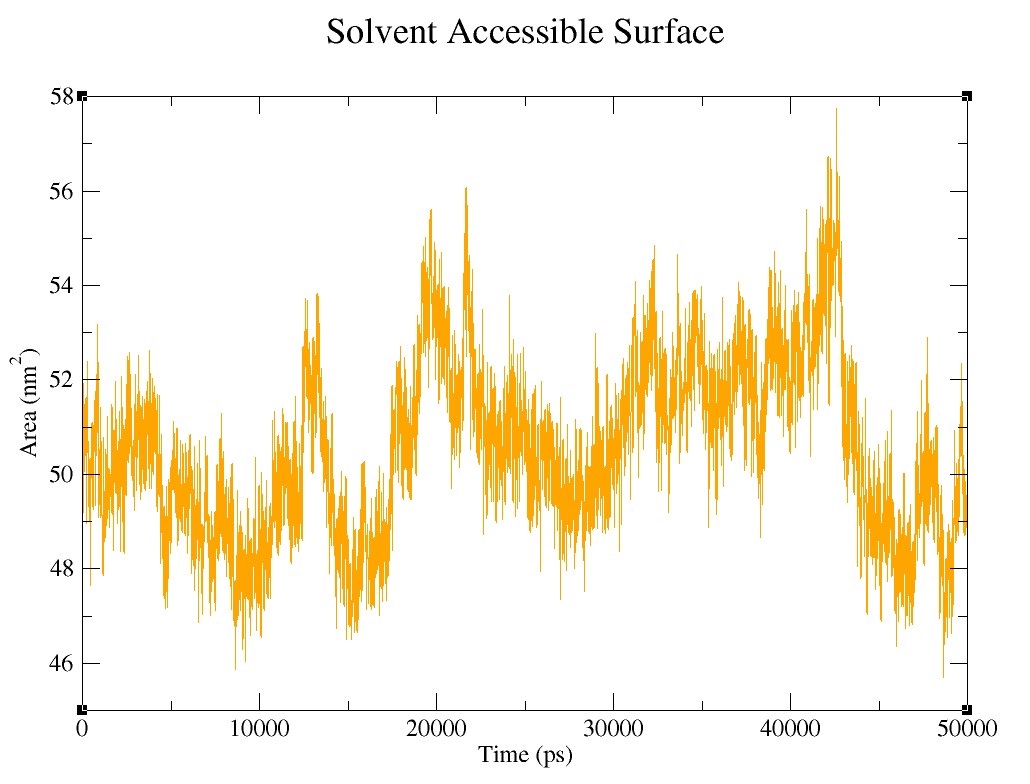

### Supplementary figure 4.6.jpg

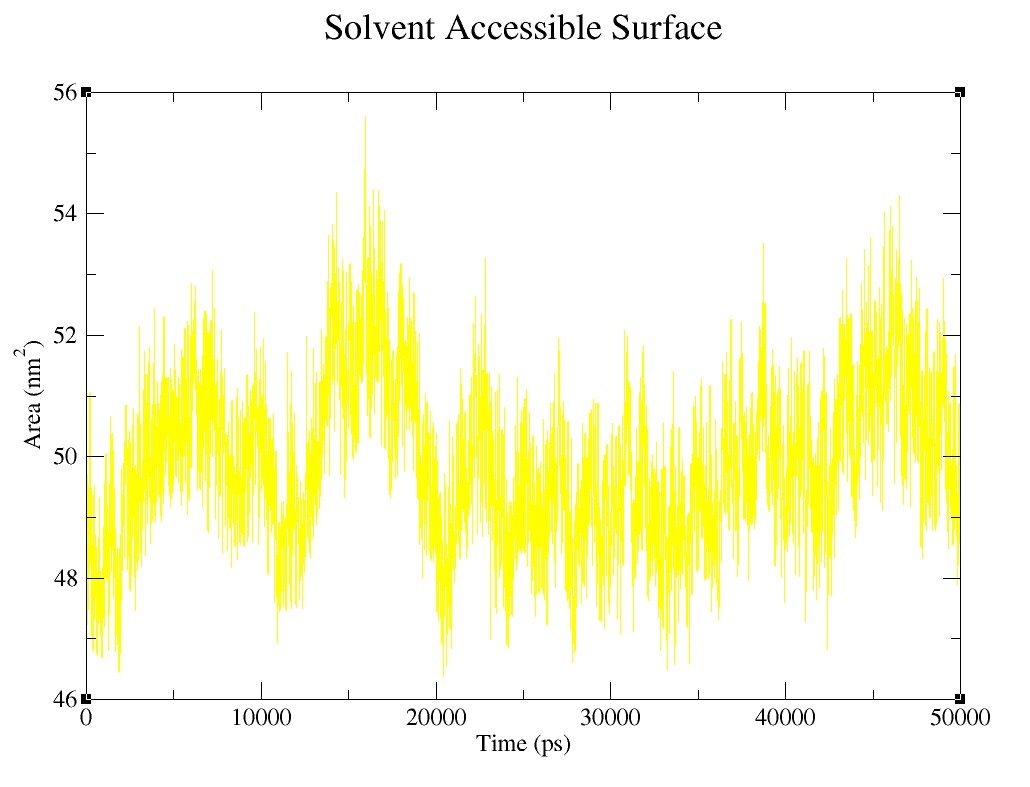

### Supplementary figure 4.7.jpg

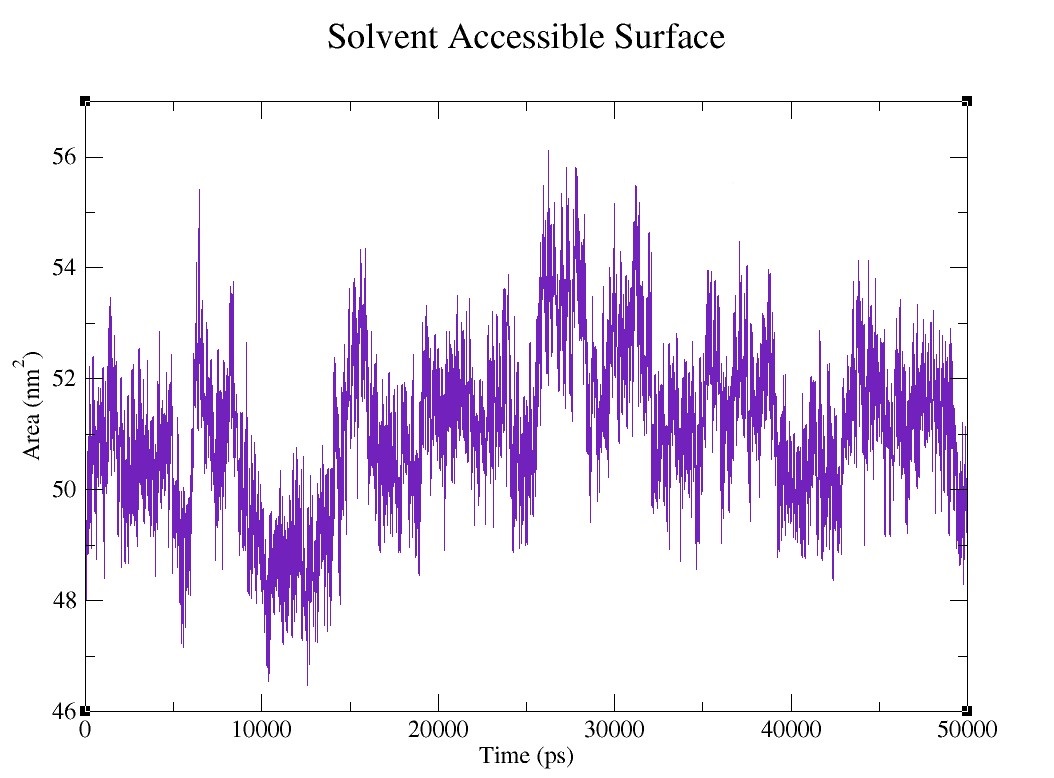

### Supplementary figure 4.8.jpg

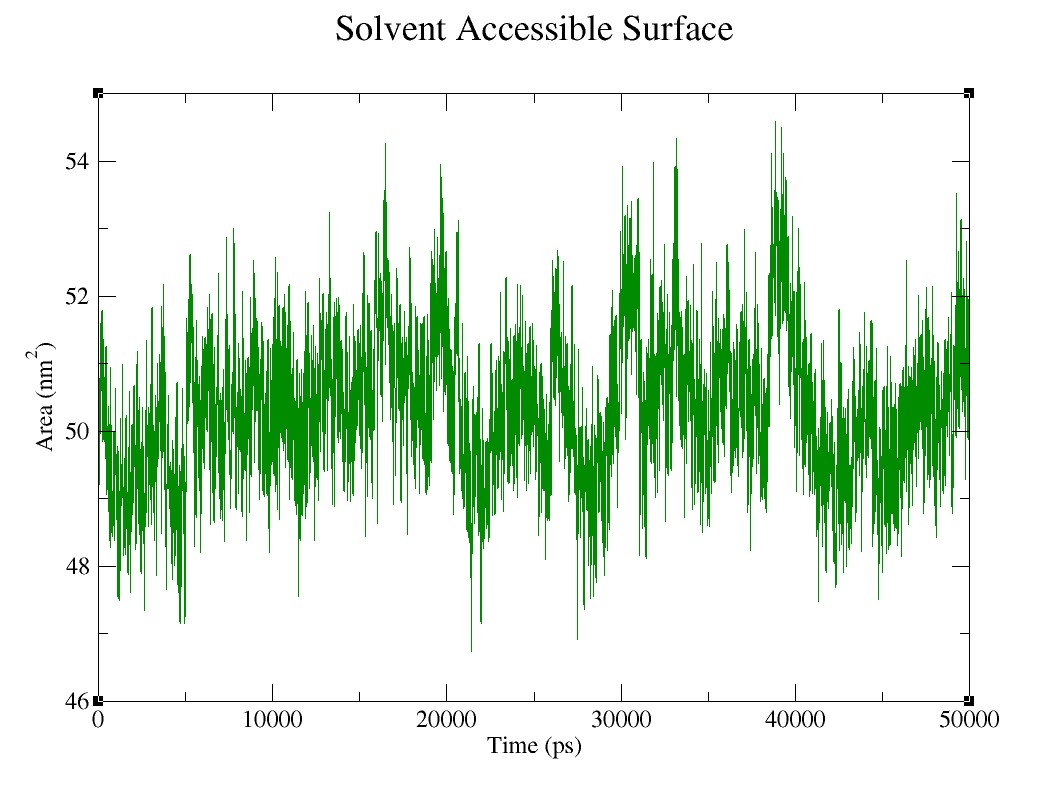

### Supplementary figure 4.9.jpg

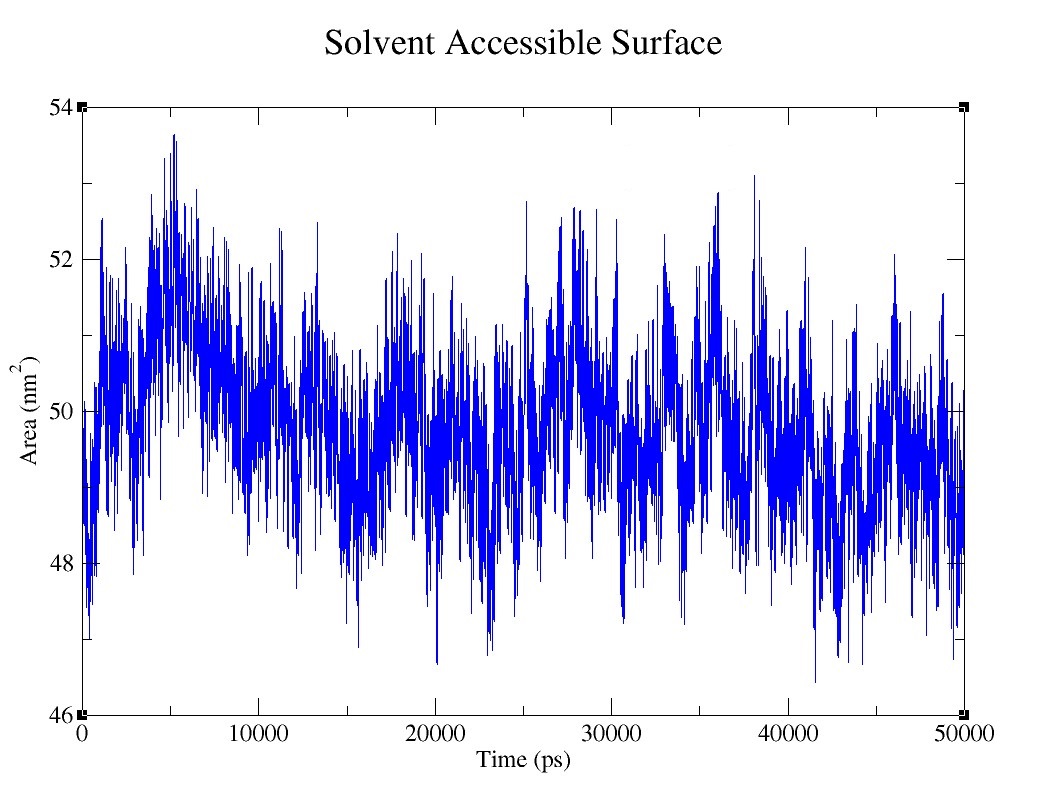

### Supplementary figure 4.10.jpg

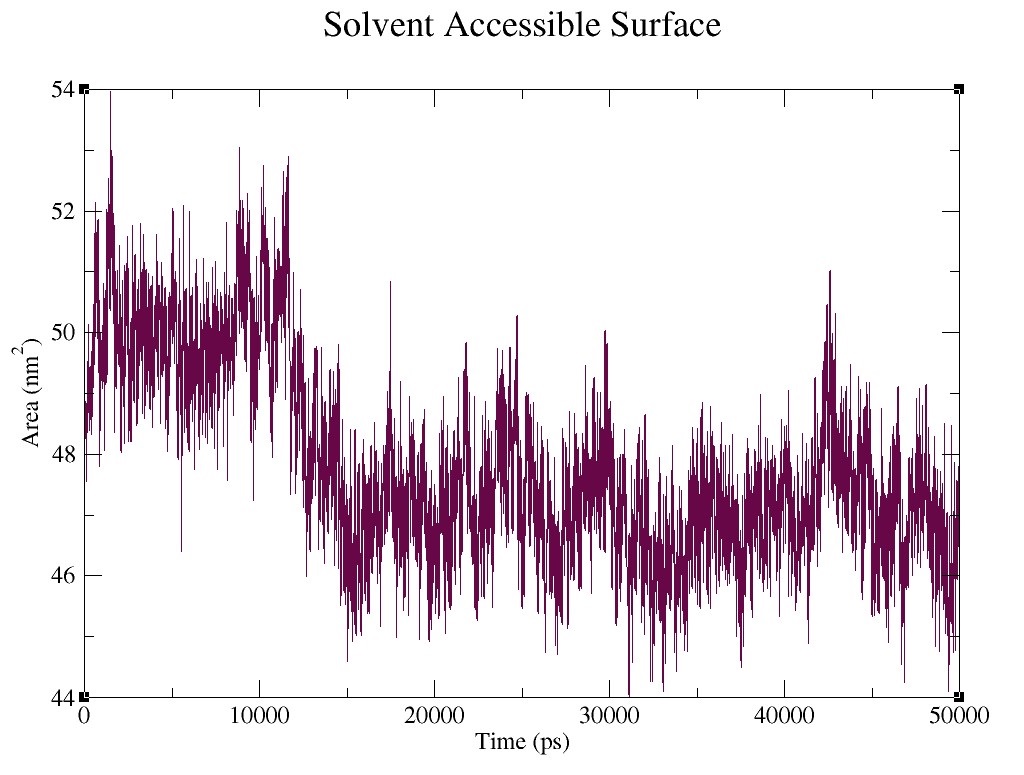

### Supplementary figure 4.11.jpg

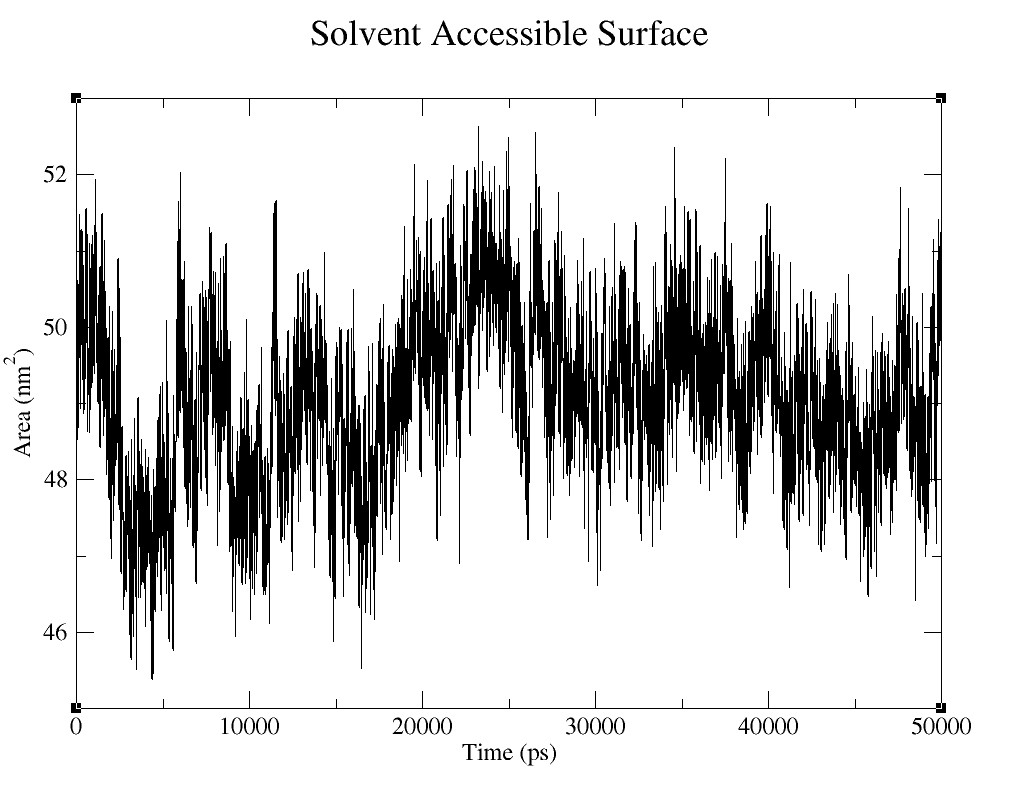

### Supplementary figure 5.1.jpg

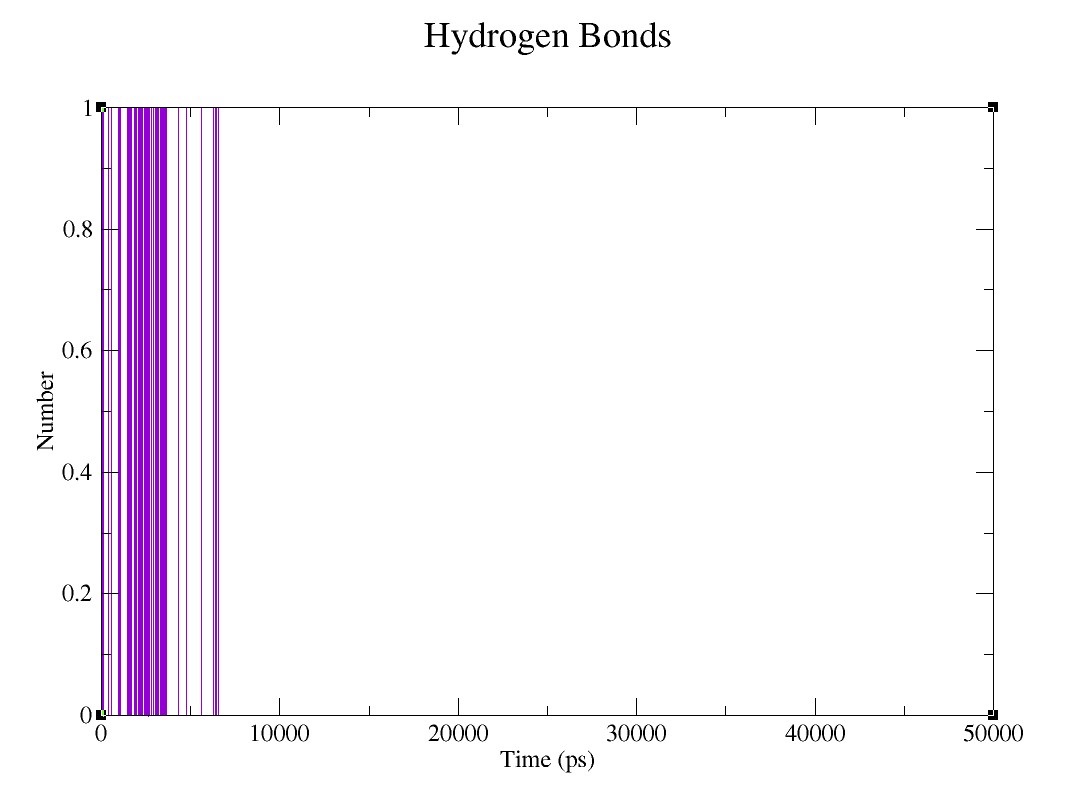

### Supplementary figure 5.2.jpg

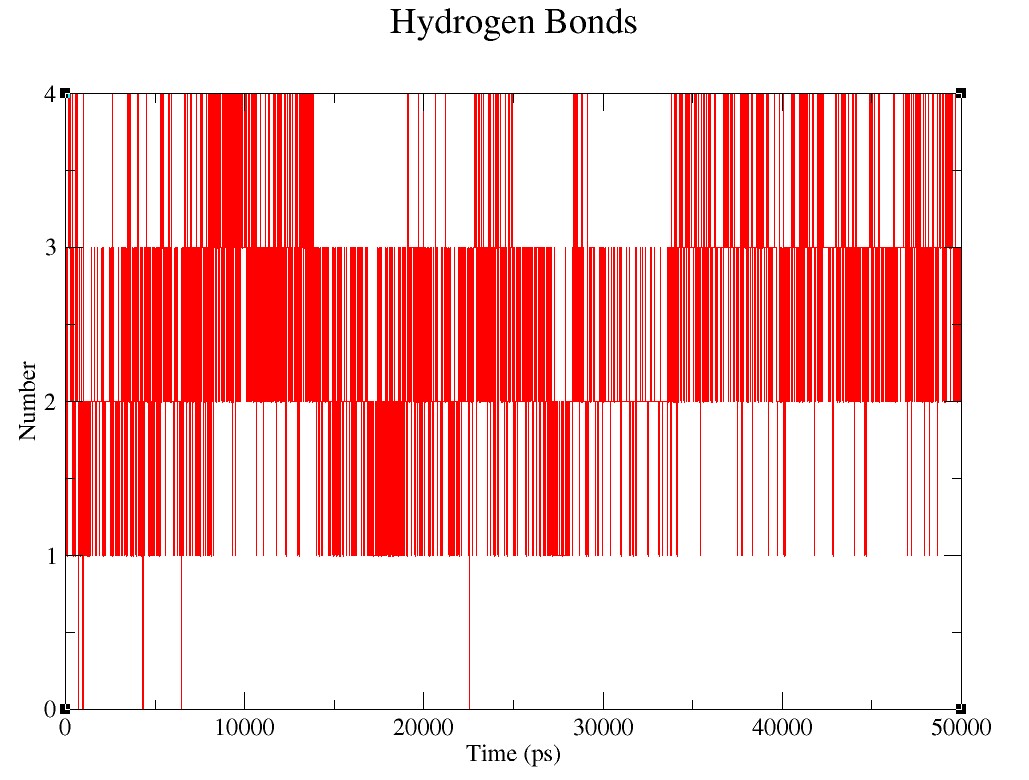

### Supplementary figure 5.3.jpg

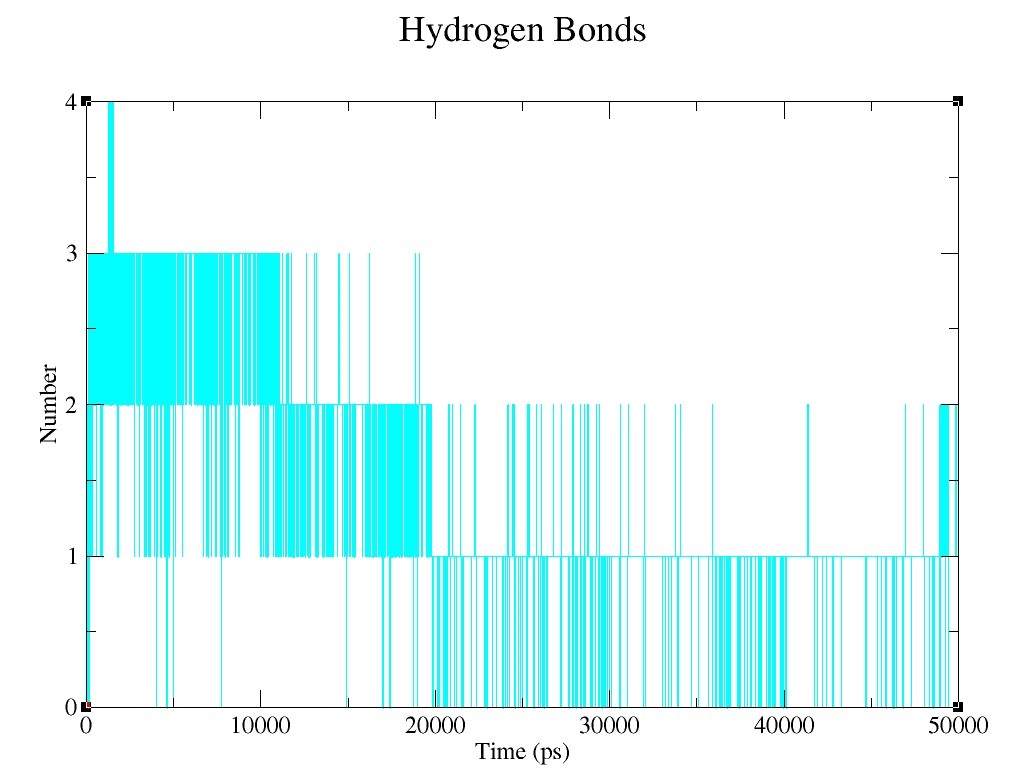

### Supplementary figure 5.4.jpg

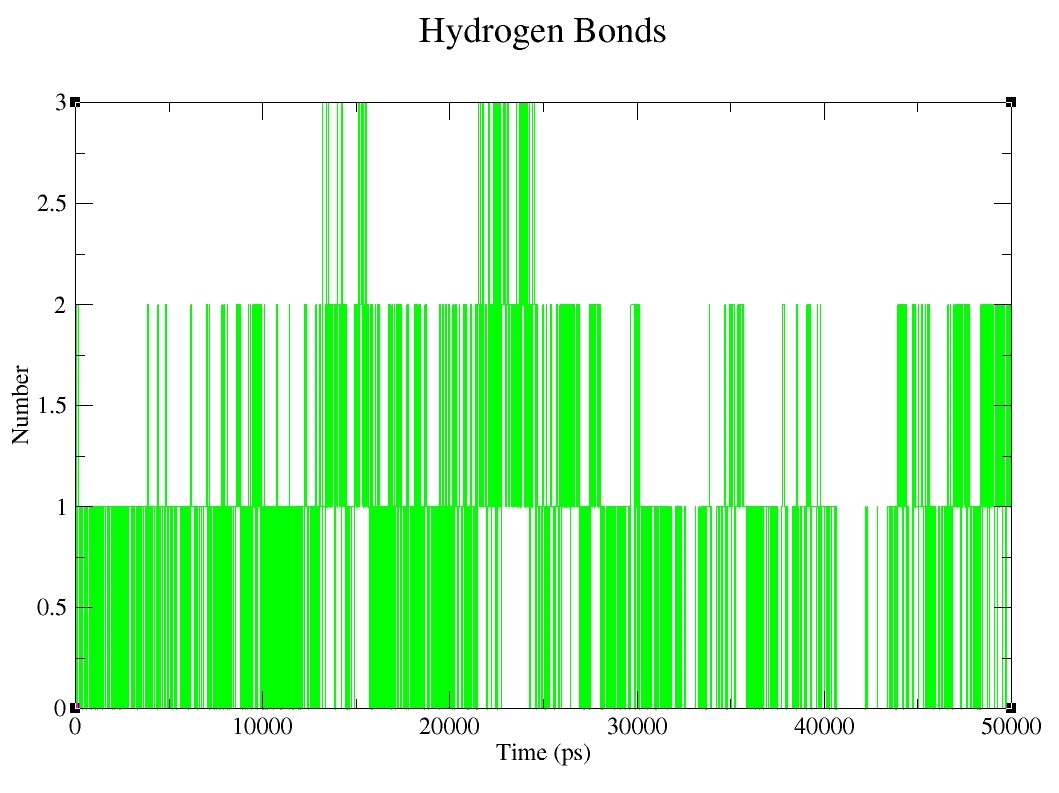

### Supplementary figure 5.5.jpg

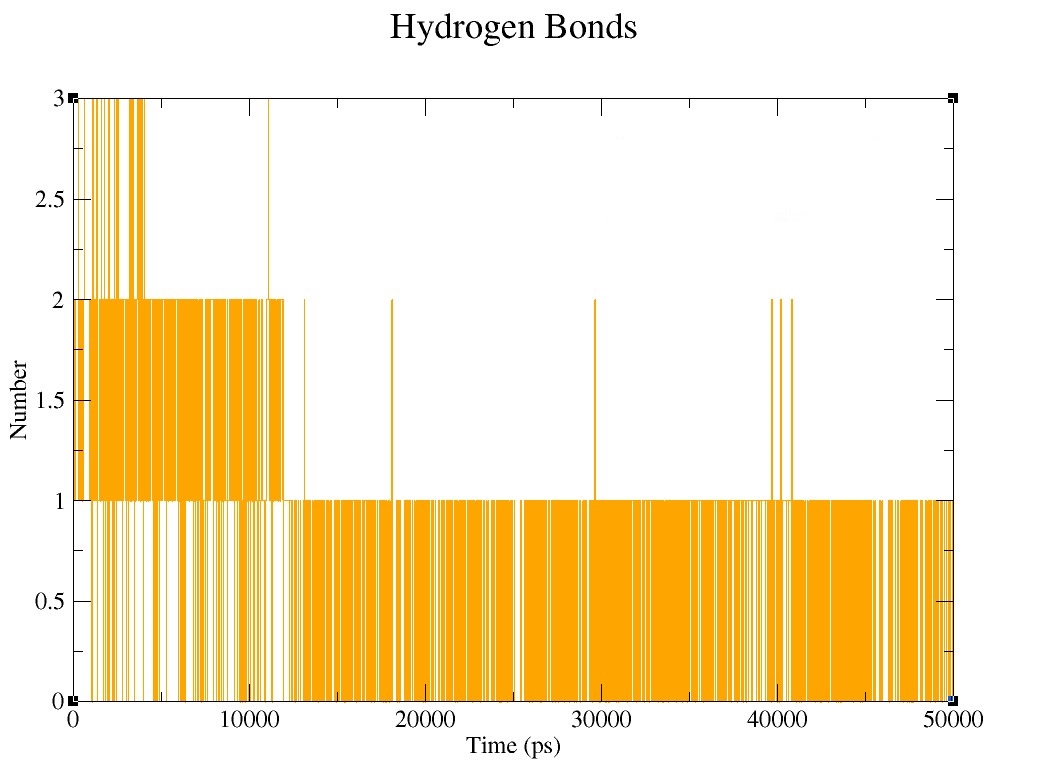

### Supplementary figure 5.6.jpg

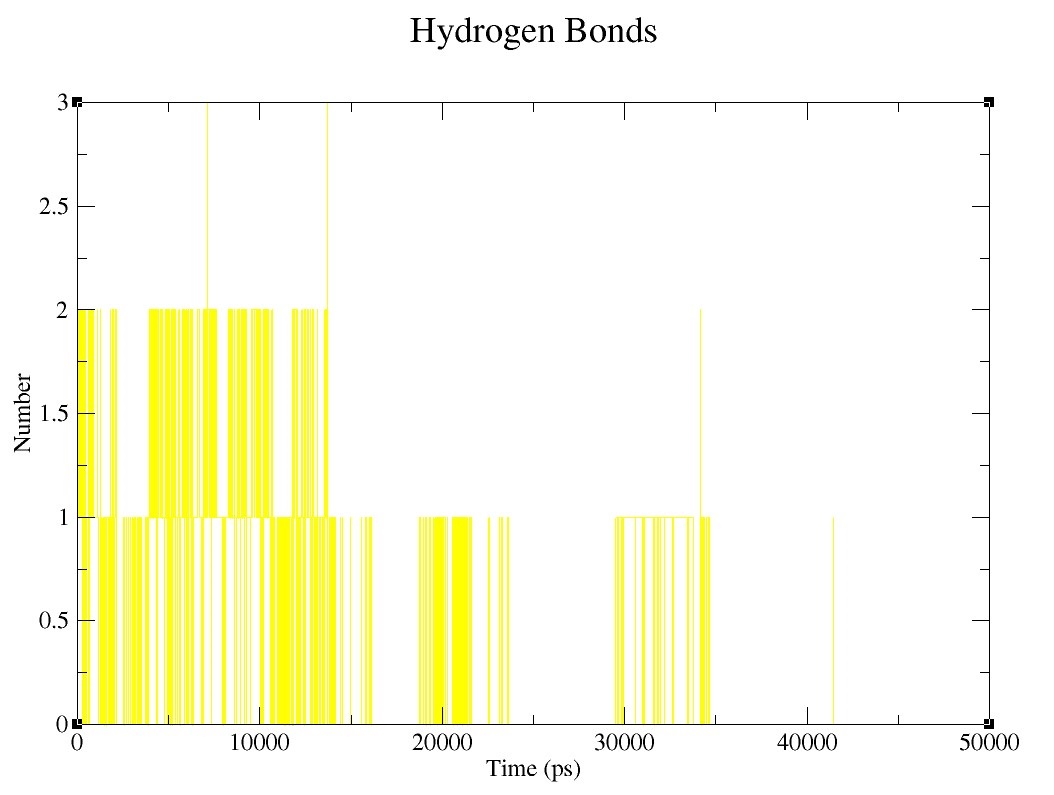

### Supplementary figure 5.7.jpg

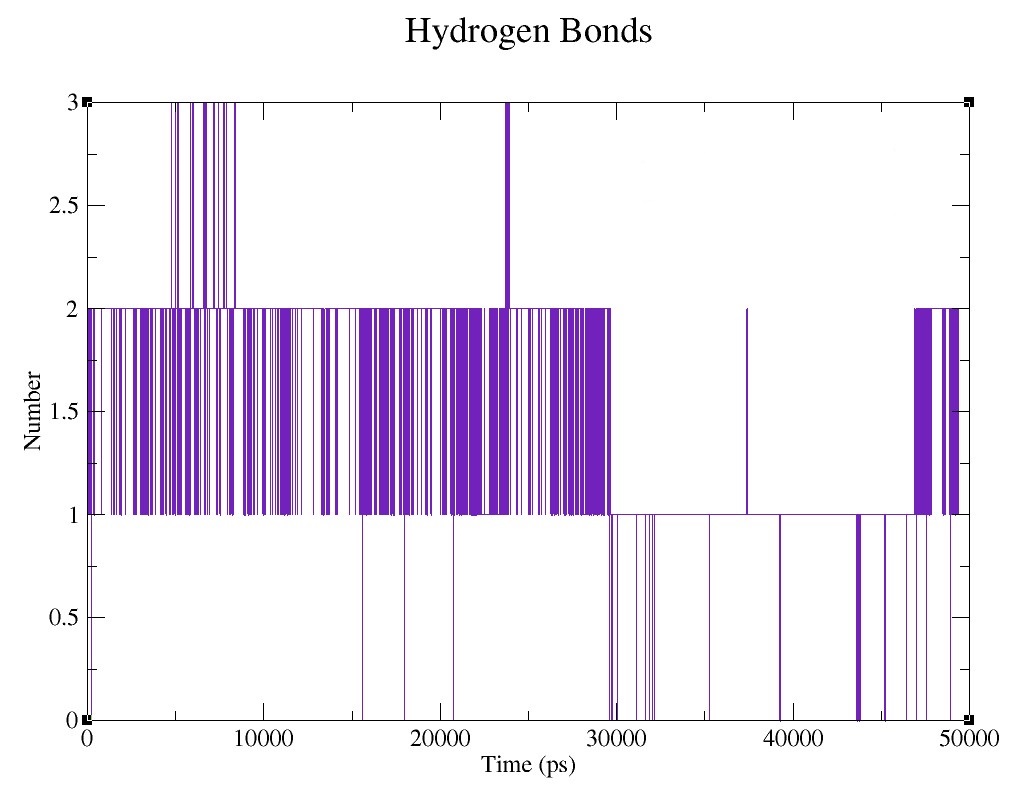

### Supplementary figure 5.8.jpg

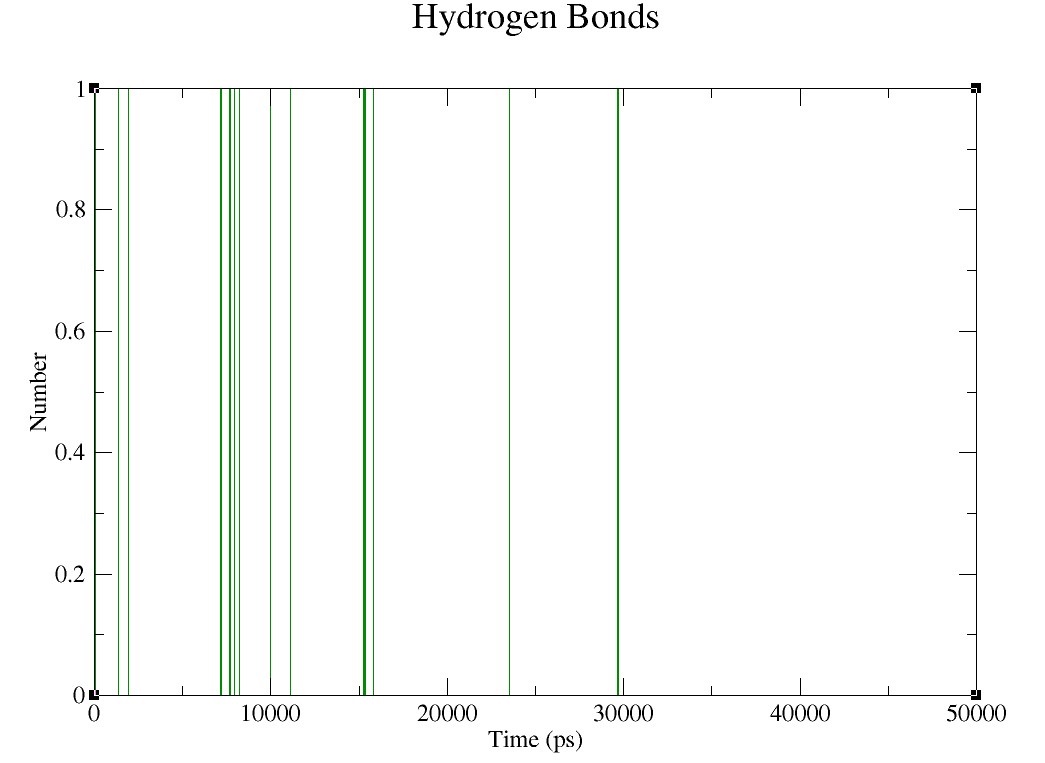

### Supplementary figure 5.9.jpg

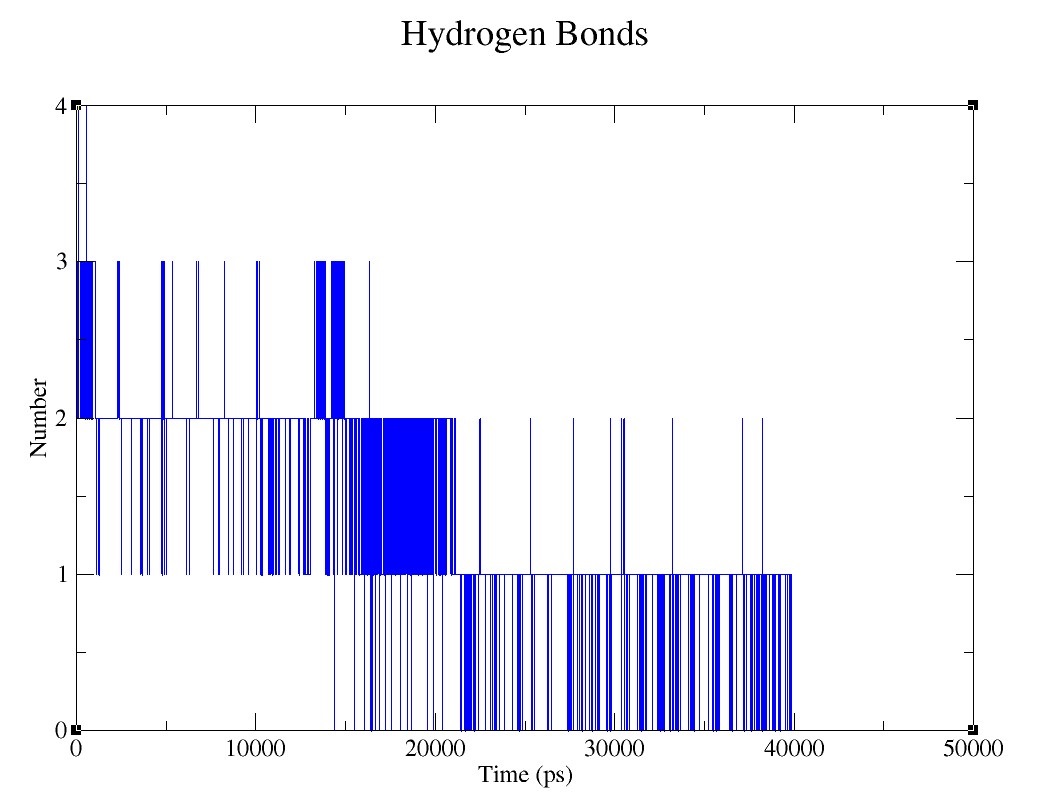

### Supplementary figure 5.10.jpg

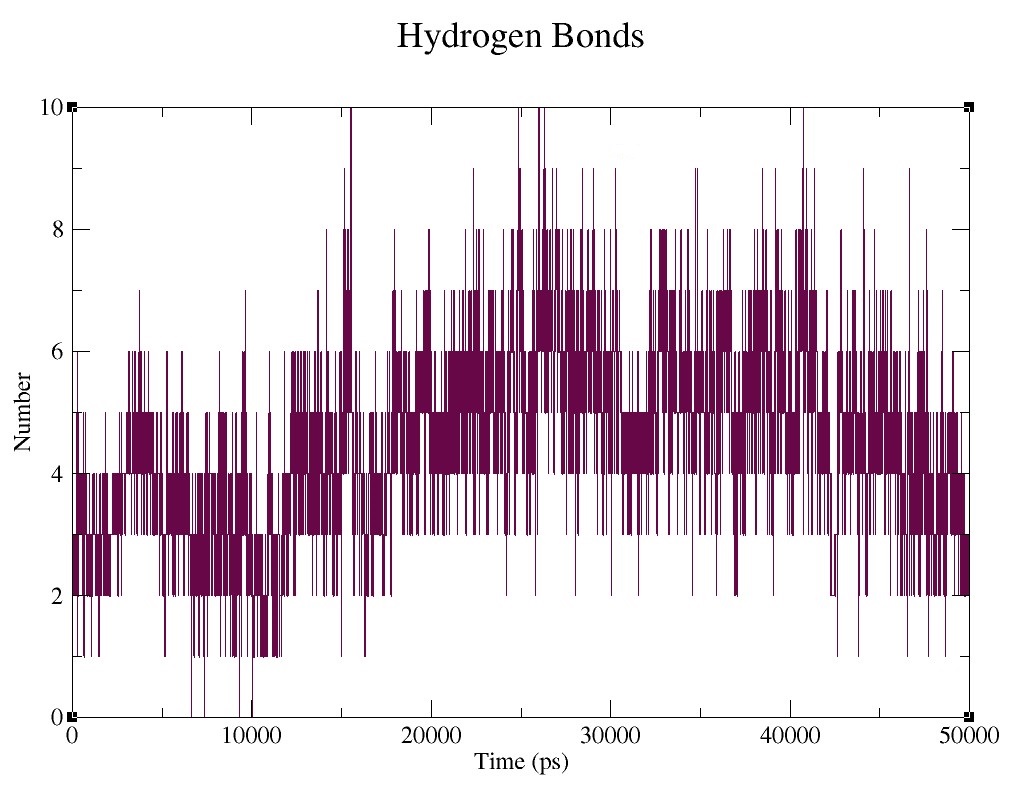

### Supplementary figure 5.11.jpg

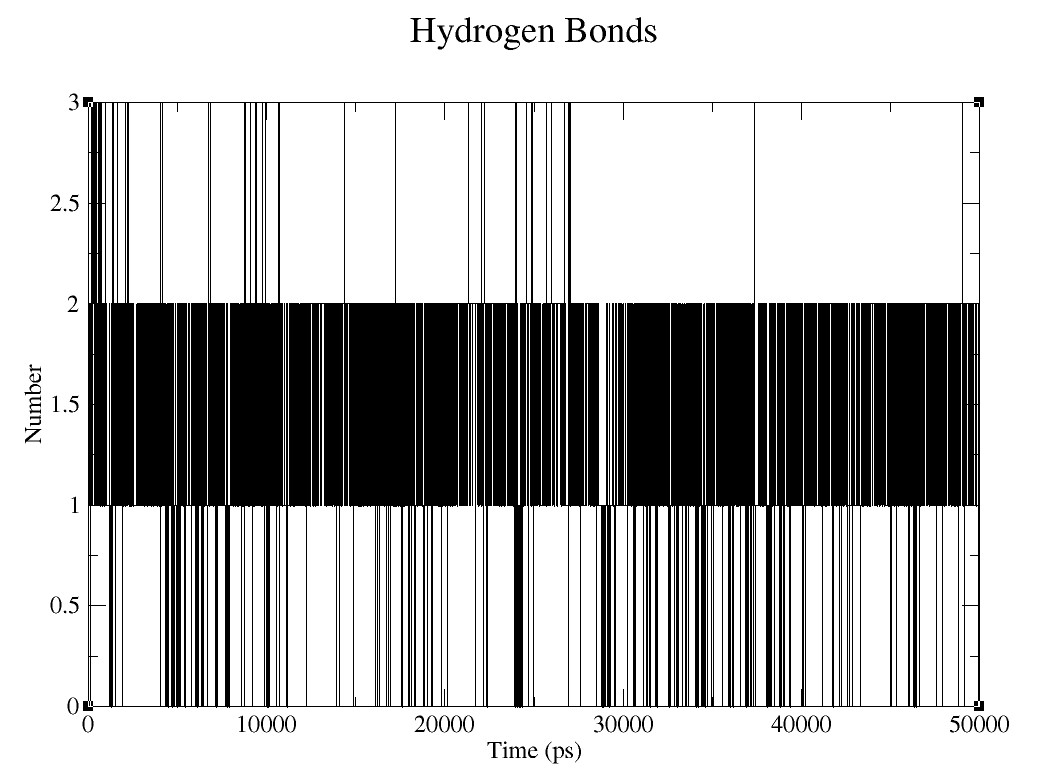

### Supplementary figures 1.1.jpg

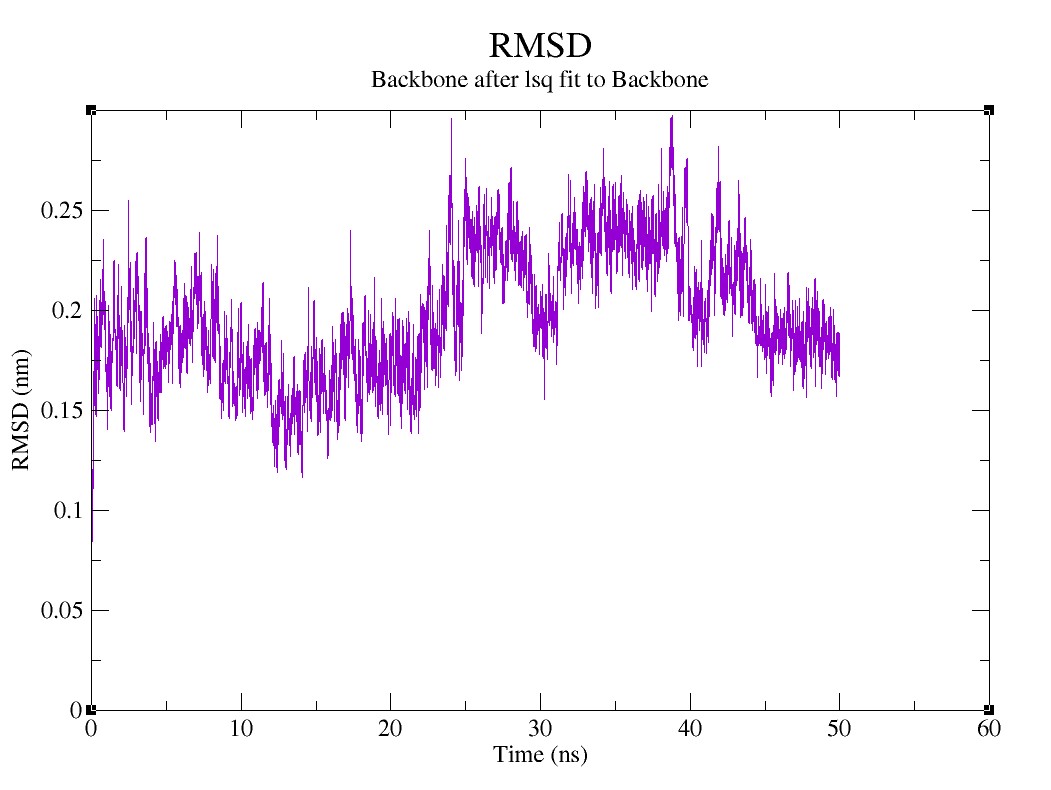

### Supplementary figures 1.2.jpg

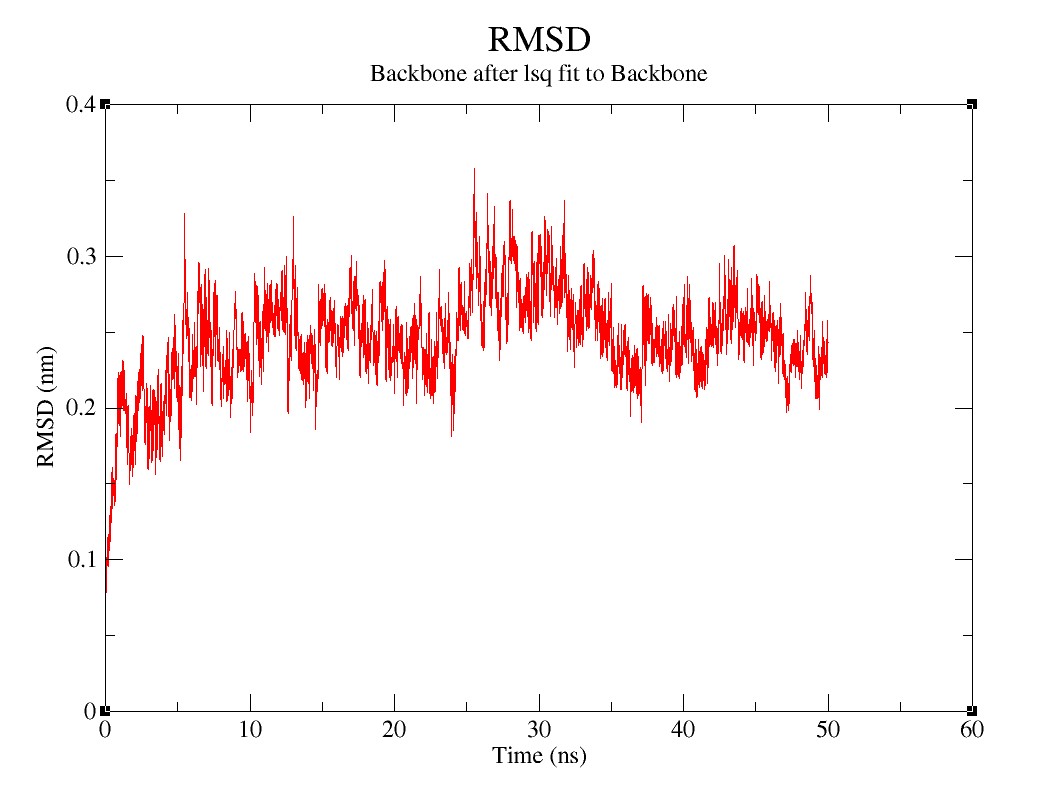

### Supplementary figures 1.3.jpg

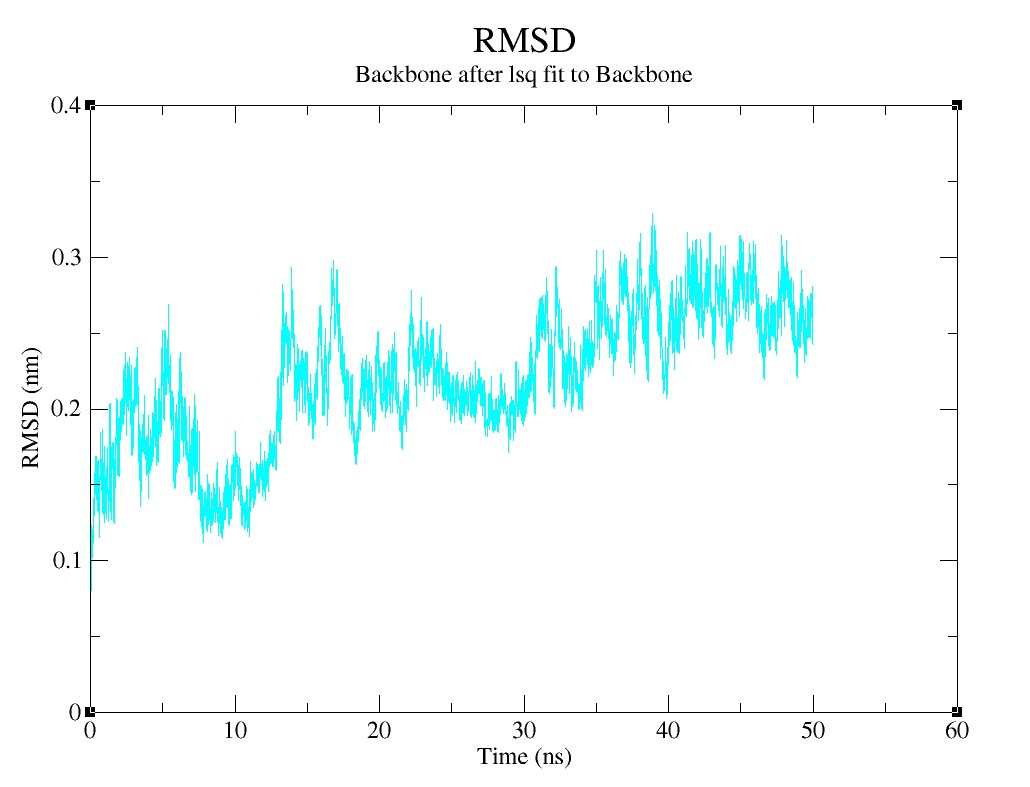

### Supplementary figures 1.4.jpg

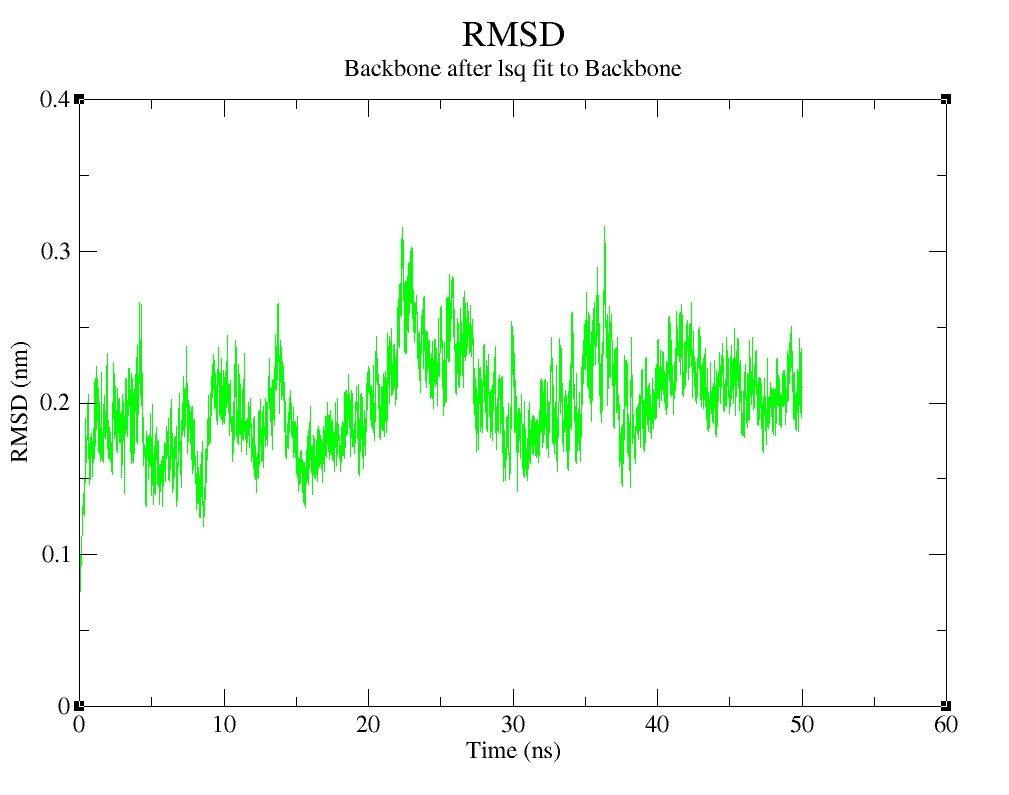

### Supplementary figures 1.5.jpg

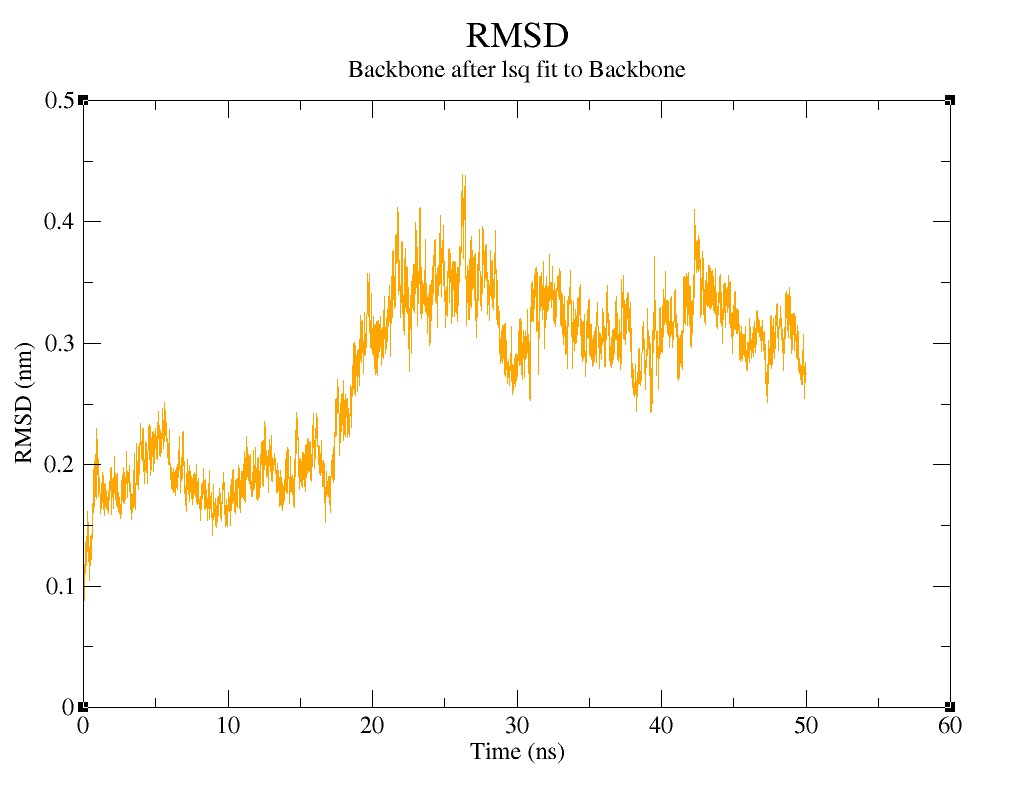

### Supplementary figures 1.6.jpg

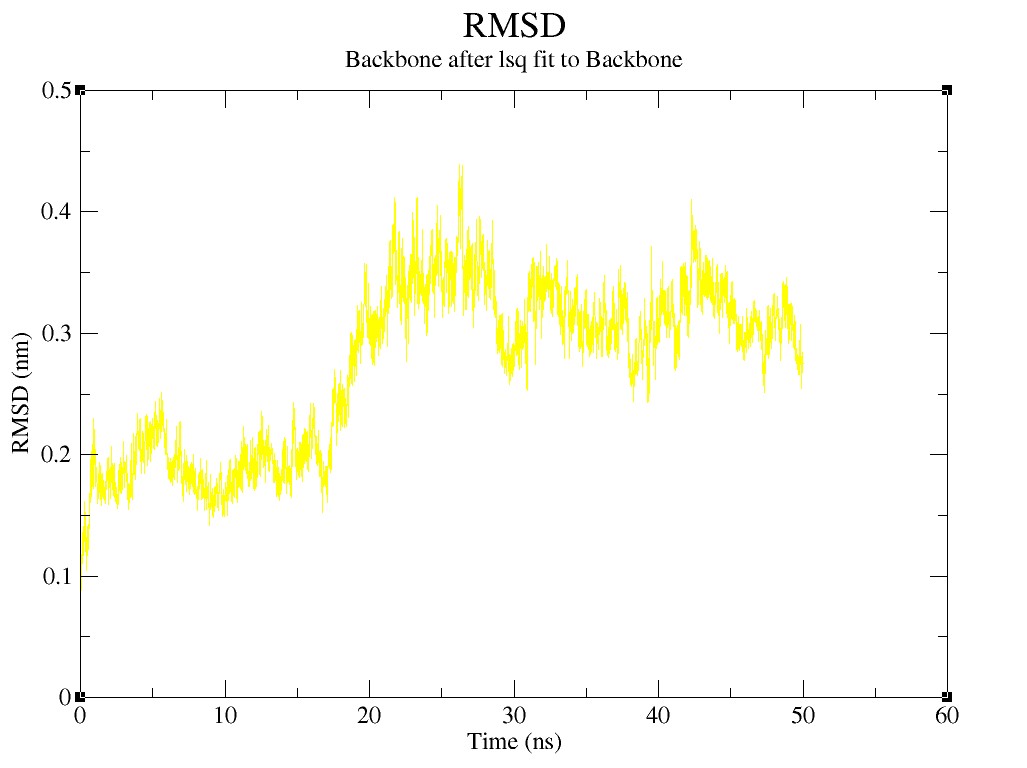

### Supplementary figures 1.7.jpg

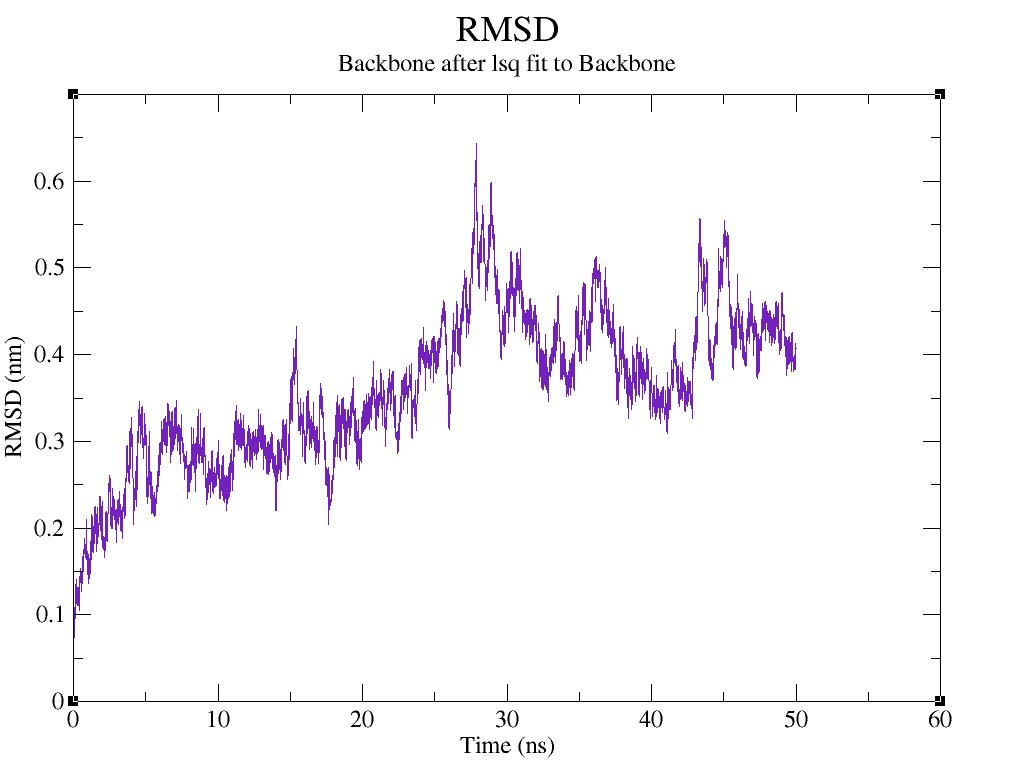

### Supplementary figures 1.8.jpg

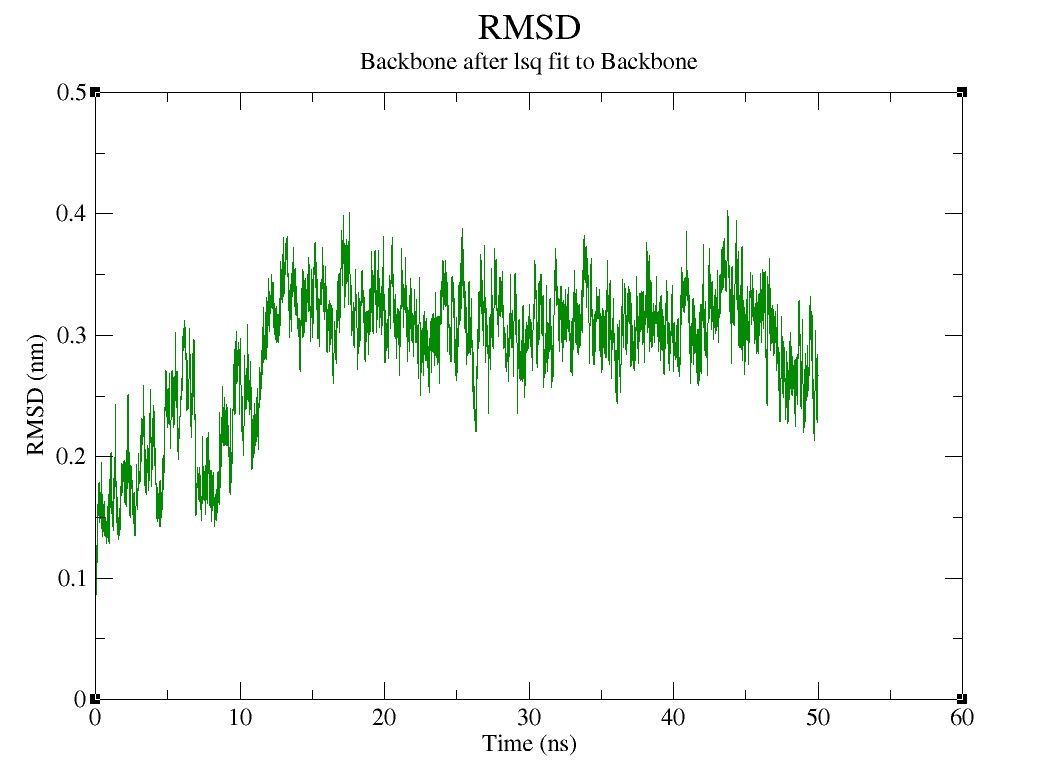

### Supplementary figures 1.9.jpg

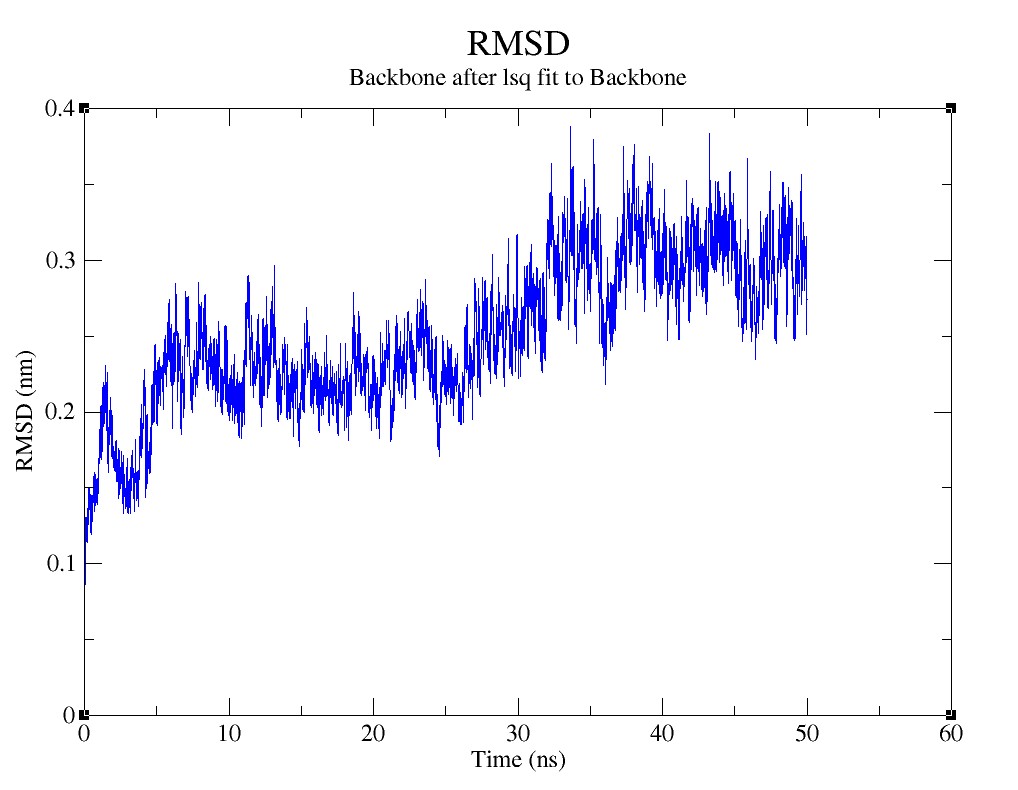

### Supplementary figures 1.10.jpg

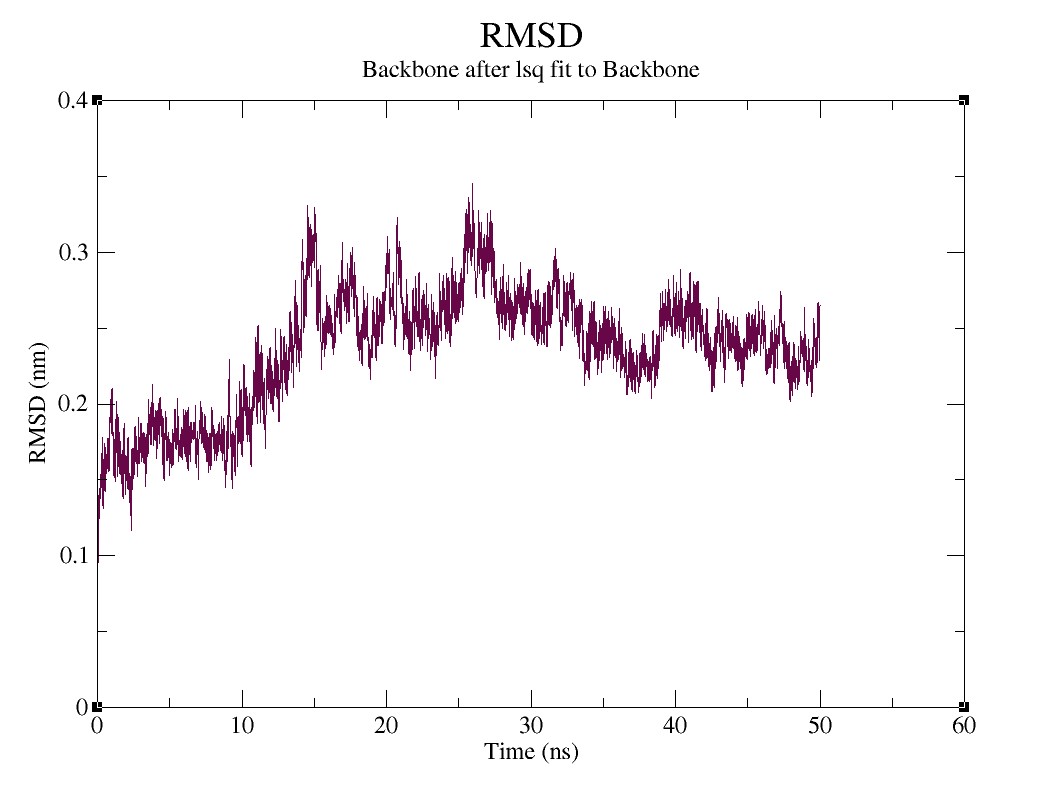

### Supplementary figures 1.11.jpg

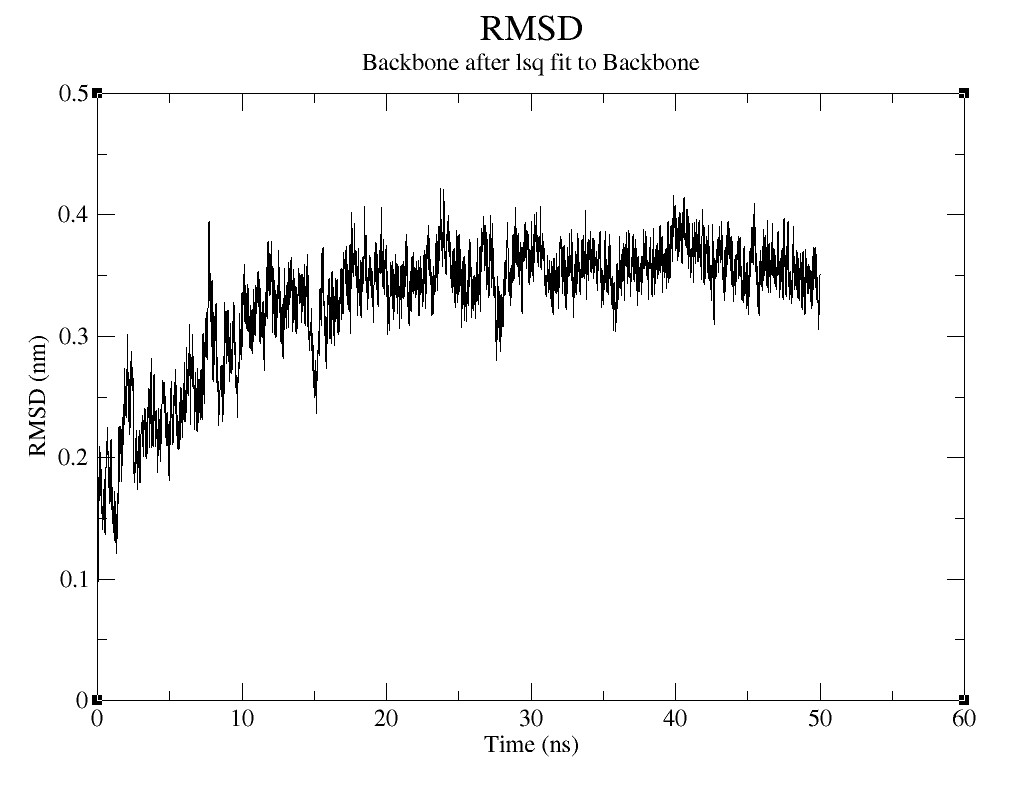
