## SupplementaryTables for "Rice WRKY13 TF protein binds to motifs in the promoter region to regulate downstream disease resistance-related genes": Manuscript_Supplementarytables_Jimmy et al. 2023.pdf

**Supplementary Table 1.** List of rice genes having WRKY13 binding motifs in their promoters

| Chromosome 1 | | | | | |
|---|---|---|---|---|---|
| **Motif** | **ID** | **Frequency** | **Locus ID** | **Protein name** | **Positions in 1k promoter region** |
| **TTTGACT** | >Os01t0633000-01 | 4 | LOC_Os01g44210 | 50S ribosomal protein L31 | 408 to 414, 429 to 435, 520 to 526, 587 to 593 |
| **TTGACTT** | >Os01t0633000-01 | 4 | LOC_Os01g44210 | 50S ribosomal protein L31 | 409 to 415, 430 to 436, 521 to 527, 588 to 594 |
| | >Os01t0633000-01 | 4 | LOC_Os01g44210 | 50S ribosomal protein L31 | 409 to 415, 430 to 436, 521 to 527, 588 to 594 |
| **CTGACT** | >Os01t0202800-00 | 5 | LOC_Os01g10600 | NOD26 like intrinsic protein 1;2 | 242 to 247, 388 to 393, 392 to 397, 396 to 401, 400 to 405 |
| **TGACC** | >Os01t0104900-01 | 6 | LOC_Os01g01520 | Transferase family protein | 93 to 97, 353 to 357, 367 to 371, 415 to 419, 421 to 425, 575 to 579 |
| | >Os01t0136400-01 | 4 | LOC_Os01g04409 | Wall-Associated kinase 1 | 143 to 147, 401 to 405, 768 to 772, 862 to 866 |
| | >Os01t0153000-00 | 4 | LOC_Os01g05980 | Protein kinase, catalytic domain | 30 to 34, 330 to 334, 678 to 682, 740 to 744 |
| | >Os01t0174500-01 | 4 | LOC_Os01g07920 | Metridin-like ShK toxin domain | 168 to 172, 270 to 274, 450 to 454, 483 to 487 |
| | >Os01t0243000-01 | 5 | LOC_Os01g14080 | Triacylglycerol lipase | 268 to 272, 782 to 786, 846 to 850, 885 to 889, 893 to 897 |
| | >Os01t0246400-01 | 4 | LOC_Os01g14410 | EARLY LIGHT INDUCIBLE PROTEIN | 581 to 585, 792 to 796, 858 to 862, 947 to 951 |
| | >Os01t0249700-00 | 4 | LOC_Os01g14700 | Heavy metal-associated domain | 37 to 41, 374 to 378, 399 to 403, 615 to 619 |
| | >Os01t0252100-01 | 4 | LOC_Os01g14860 | Shaggy-like kinase gamma | 387 to 391, 482 to 486, 666 to 670, 779 to 783 |
| | >Os01t0252100-02 | 4 | LOC_Os01g14860 | Glycogen synthase kinase-3 homolog MsK-3 | 221 to 225, 316 to 320, 500 to 504, 613 to 617 |
| | >Os01t0252200-01 | 4 | LOC_Os01g14870 | Zinc finger CCCH domain 3 | 147 to 151, 159 to 163, 530 to 534, 535 to 539 |
| | >Os01t0384450-01 | 5 | LOC_Os01g28744 | H0124E07.4 protein | 209 to 213, 397 to 401, 620 to 624, 879 to 883, 910 to 914 |
| | >Os01t0386700-01 | 4 | LOC_Os01g28989 | BSK1 (BR-SIGNALING KINASE 1); ATP binding / binding / kinase/ protein kinase/ protein tyrosine kinase | 417 to 421, 572 to 576, 729 to 733, 953 to 957 |
| | >Os01t0536950-00 | 4 | LOC_Os01g35300 | Rust resistance protein rp3-1 | 367 to 371, 400 to 404, 424 to 428, 533 |

| | | | | | |
|---|---|---|---|---|---|
| | | | | | to 537 |
| | >Os01t0558600-01 | 4 | LOC_Os01g37800 | Small GTP-binding protein OsRab1B2 | 223 to 227, 229 to 233, 249 to 253, 617 to 621 |
| | >Os01t0625900-01 | 5 | LOC_Os01g43610 | Ovate family protein 3 | 372 to 376, 404 to 408, 416 to 420, 648 to 652, 656 to 660 |
| | >Os01t0665750-01 | 4 | LOC_Os01g47560 | WRKY transcription factor 16 | 167 to 171, 478 to 482, 487 to 491, 530 to 534 |
| | >Os01t0718300-03 | 5 | LOC_Os01g52050 | BRASSINOSTEROID INSENSITIVE 1 | 155 to 159, 341 to 345, 368 to 372, 785 to 789, 932 to 936 |
| | >Os01t0718900-01 | 4 | LOC_Os01g52090 | MADF domain | 345 to 349, 934 to 938, 942 to 946, 966 to 970 |
| | >Os01t0746400-00 | 4 | LOC_Os01g54270 | Carotenoid cleavage dioxygenase 8 | 348 to 352, 424 to 428, 799 to 803, 853 to 857 |
| | >Os01t0757900-01 | 4 | LOC_Os01g55310 | Haloacid dehalogenase/epoxide hydrolase domain | 289 to 293, 553 to 557, 692 to 696, 882 to 886 |
| | >Os01t0758900-01 | 4 | LOC_Os01g55410 | Protein of unknown function DUF688 family protein | 5 to 9, 460 to 464, 516 to 520, 545 to 549 |
| | >Os01t0764950-01 | 4 | LOC_Os01g55960 | Pentatricopeptide repeat domain | 75 to 79, 383 to 387, 617 to 621, 742 to 746 |
| | >Os01t0772200-01 | 5 | LOC_Os01g56550 | OsTFIIF2-2 of RNA polymerase II | 66 to 70, 239 to 243, 417 to 421, 428 to 432, 709 to 713 |
| | >Os01t0772200-02 | 5 | LOC_Os01g56550 | Transcription initiation factor IIF, beta subunit family protein | 66 to 70, 239 to 243, 417 to 421, 428 to 432, 709 to 713 |
| | >Os01t0772700-00 | 4 | LOC_Os01g56590 | Armadillo-type fold domain | 495 to 499, 502 to 506, 652 to 656, 657 to 661 |
| | >Os01t0823700-01 | 4 | LOC_Os01g60850 | Protein of unknown function DUF641 | 171 to 175, 183 to 187, 242 to 246, 567 to 571 |
| | >Os01t0893700-00 | 4 | LOC_Os01g66900 | DOMON domain domain | 675 to 679, 698 to 702, 721 to 725, 744 to 748 |
| | >Os01t0927000-01 | 4 | LOC_Os01g70220 | SET domain SET118. histone-lysine N-methyltransferase | 9 to 13, 257 to 261, 493 to 497, 721 to 725 |
| | >Os01t0936900-00 | 5 | LOC_Os01g71060 | Peptidase A1 domain. xylanase inhibitor | 88 to 92, 99 to 103, 184 to 188, 260 to 264, 907 to 911 |
| **TGACT** | >Os01t0100900-01 | 4 | LOC_Os01g01080 | Pyyridoxal phosphate-dependent decarboxylase domain decarboxylase | 353 to 357, 365 to 369, 678 to 682, 728 to 732 |

| | | | | | |
|---|---|---|---|---|---|
| | >Os01t0102600-02 | 4 | LOC_Os01g01302 | Shikimate kinase domain | 878 to 882, 902 to 906, 924 to 928, 956 to 960 |
| | >Os01t0114700-00 | 5 | LOC_Os01g02440 | Ser/Thr receptor-like kinase | 43 to 47, 506 to 510, 609 to 613, 630 to 634, 790 to 794 |
| | >Os01t0115800-01 | 4 | LOC_Os01g02590 | Receptor-like kinase | 580 to 584, 659 to 663, 871 to 875, 875 to 879 |
| | >Os01t0128400-02 | 4 | LOC_Os01g03760 | Protein of unknown function DUF2359, TMEM214 domain | 592 to 596, 683 to 687, 727 to 731, 973 to 977 |
| | >Os01t0140500-01 | 4 | LOC_Os01g04730 | 60S ribosomal protein L26B | 69 to 73, 76 to 80, 441 to 445, 460 to 464 |
| | >Os01t0160700-00 | 5 | LOC_Os01g06730 | Leucine-rich repeat, N-terminal domain verticillium wilt disease resistance protein | 172 to 176, 282 to 286, 543 to 547, 604 to 608, 825 to 829 |
| | >Os01t0160800-01 | 4 | LOC_Os01g06740 | Protein synthesis inhibitor II. ribosome inactivating protein | 10 to 14, 697 to 701, 810 to 814, 934 to 938 |
| | >Os01t0167500-01 | 4 | LOC_Os01g07310 | Protein of unknown function DUF250 domain transporter-related | 240 to 244, 416 to 420, 424 to 428, 457 to 461 |
| | >Os01t0182300-01 | 4 | LOC_Os01g08670 | Lipocalin domain | 152 to 156, 412 to 416, 670 to 674, 806 to 810 |
| | >Os01t0198702-01 | 4 | LOC_Os01g10600 | NOD26-like membrane integral protein ZmNIP1-1. aquaporin protein | 106 to 110, 701 to 705, 764 to 768, 795 to 799 |
| | >Os01t0202800-00 | 9 | LOC_Os01g12800 | Mpv17/PMP22 family protein. peroxisomal membrane protein | 243 to 247, 317 to 321, 325 to 329, 385 to 389, 389 to 393, 393 to 397, 397 to 401, 401 to 405, 669 to 673 |
| | >Os01t0228300-01 | 4 | LOC_Os01g12860 | Transcription factor myb. MYB family transcription factor | 553 to 557, 579 to 583, 853 to 857, 857 to 861 |
| | >Os01t0229000-00 | 4 | LOC_Os01g12880 | VHS domain VHS and GAT domain | 125 to 129, 297 to 301, 563 to 567, 812 to 816 |
| | >Os01t0229200-03 | 4 | LOC_Os01g14140 | Beta-1,3-glucanase | 403 to 407, 564 to 568, 568 to 572, 914 to 918 |
| | >Os01t0243700-01 | 4 | LOC_Os01g14140 | Beta-1,3-glucanase | 71 to 75, 184 to 188, 322 to 326, 642 to 646 |
| | >Os01t0245532-00 | 4 | LOC_Os01g14630 | Terpenoid synthase domain. polyprenyl synthetase | 228 to 232, 423 to 427, 519 to 523, 851 to 855 |
| | >Os01t0248701-00 | 4 | LOC_Os01g14840 | Zinc finger, C2H2-like domain. ZOS1-06 - C2H2 zinc finger protein | 149 to 153, 833 to 837, 957 to 961, 982 to 986 |

| | | | | | |
|---|---|---|---|---|---|
| | >Os01t0251200-02 | 5 | LOC_Os01g14932 | Avr9/Cf-9 induced kinase 1. NAK-like ser/thr protein kinase | 356 to 360, 643 to 647, 682 to 686, 703 to 707, 867 to 871 |
| | >Os01t0253100-02 | 4 | LOC_Os01g15010 | NTGB2 | 563 to 567, 572 to 576, 762 to 766, 860 to 864 |
| | >Os01t0272800-01 | 4 | LOC_Os01g18640 | Pyridoxal phosphate-dependent transferase, major region, subdomain 1 domain. aminotransferase | 209 to 213, 624 to 628, 734 to 738, 849 to 853 |
| | >Os01t0290100-01 | 4 | | Sorbitol transporter | 22 to 26, 86 to 90, 211 to 215, 757 to 761 |
| | >Os01t0337180-01 | 5 | LOC_Os01g25760 | NB-ARC domain. powdery mildew resistance protein PM3F | 297 to 301, 414 to 418, 630 to 634, 777 to 781, 870 to 874 |
| | >Os01t0359800-00 | 4 | LOC_Os01g28600 | exo70 exocyst complex subunit family protein | 303 to 307, 466 to 470, 590 to 594, 825 to 829 |
| | >Os01t0383100-01 | 4 | LOC_Os01g32350 | NClpP3 (ATP-dependent Clp protease proteolytic subunit ClpP3). OsClp2 - Putative Clp protease homologue | 345 to 349, 428 to 432, 734 to 738, 874 to 878 |
| | >Os01t0511700-01 | 4 | LOC_Os01g33784 | Lipase family protein | 510 to 514, 784 to 788, 814 to 818, 843 to 847 |
| | >Os01t0521400-01 | 4 | LOC_Os01g33784 | Lipase family protein | 45 to 49, 141 to 145, 213 to 217, 307 to 311 |
| | >Os01t0532200-01 | 5 | LOC_Os01g34780 | FF domain domain | 204 to 208, 251 to 255, 733 to 737, 742 to 746, 819 to 823 |
| | >Os01t0532200-02 | 5 | LOC_Os01g34780 | FF domain domain | 283 to 287, 292 to 296, 369 to 373, 780 to 784, 785 to 789 |
| | >Os01t0561600-01 | 4 | LOC_Os01g38110 | Cytochrome P450 family protein | 236 to 240, 319 to 323, 773 to 777, 864 to 868 |
| | >Os01t0629400-01 | 4 | LOC_Os01g43870 | NLI interacting factor domain | 91 to 95, 154 to 158, 523 to 527, 937 to 941 |
| | >Os01t0633000-01 | 4 | LOC_Os01g44210 | 50S ribosomal protein L31 | 410 to 414, 431 to 435, 522 to 526, 589 to 593 |
| | >Os01t0665200-03 | 4 | LOC_Os01g47530 | Blast and wounding induced mitogen-activated protein kinase | 185 to 189, 243 to 247, 872 to 876, 937 to 941 |
| | >Os01t0673900-01 | 4 | LOC_Os01g48310 | Zinc finger, RING-type domain | 83 to 87, 232 to 236, 956 to 960, 989 to 993 |
| | >Os01t0730700-01 | 4 | LOC_Os01g53040 | WRKY transcription factor 14 | 270 to 274, 532 to 536, 539 to 543, 586 to 590 |

| | | | | | |
|---|---|---|---|---|---|
| | >Os01t0762900-01 | 4 | LOC_Os01g55720 | UPA24 | 80 to 84, 170 to174, 229 to 233, 867 to 871 |
| | >Os01t0788900-01 | 7 | LOC_Os01g57900 | Pentatricopeptide repeat domain. PPR repeat domain | 257 to 261, 288 to 292, 324 to 328, 451 to 455, 535 to 539, 567 to 571, 716 to 720 |
| | >Os01t0788950-01 | 4 | LOC_Os01g57910 | Partner of Nob1. KH domain | 400 to 404, 487 to 491, 550 to 554, 801 to 805 |
| | >Os01t0814700-01 | 4 | LOC_Os01g59910 | Cyclin-like F-box domain OsFBX29 - F-box domain | 140 to 144, 240 to 244, 324 to 328, 454 to 458 |
| | >Os01t0816100-01 | 4 | LOC_Os01g60020 | NAC4 | 500 to 504, 504 to 508, 508 to 512, 802 to 806 |
| | >Os01t0835800-01 | 4 | LOC_Os01g61910 | KIP1 | 4 to 8, 168 to 172, 361 to 365, 954 to 958 |
| | >Os01t0852100-01 | 4 | LOC_Os01g63280 | Tyrosine protein kinase domain | 4 to 8, 196 to 200, 371 to 375, 688 to 692 |
| | >Os01t0871100-01 | 4 | LOC_Os01g65070 | Alpha/beta hydrolase fold-1 domain | 331 to 335, 504 to 508, 610 to 614, 679 to 683 |
| | >Os01t0894300-01 | 4 | LOC_Os01g66940 | Fructokinase 1. kinase, pfkB family | 340 to 344, 624 to 628, 731 to 735, 835 to 839 |
| | >Os01t0894500-01 | 4 | LOC_Os01g66960 | Sep15/SelM redox domain | 161 to 165, 377 to 381, 474 to 478, 843 to 847 |
| | >Os01t0914400-01 | 4 | LOC_Os01g68598 | EPIDERMAL PATTERNING FACTOR-like protein 9 | 65 to 69, 99 to 103, 532 to 536, 660 to 664 |
| | >Os01t0925100-01 | 4 | LOC_Os01g70080 | NB-ARC domain | 76 to 80, 174 to 178, 919 to 923, 931 to 935 |
| | >Os01t0928800-01 | 6 | LOC_Os01g70380 | Serine palmitoyltransferase | 88 to 92, 232 to 236, 316 to 320, 438 to 442, 521 to 525, 746 to 750 |
| | >Os01t0930400-00 | 4 | LOC_Os01g70490 | Potassium transporter 5 | 11 to 15, 49 to 53, 202 to 206, 765 to 769 |
| | >Os01t0940800-01 | 4 | LOC_Os01g71350 | Beta-1, 3-glucanase precursor. glycosyl hydrolases family 17 | 470 to 474, 642 to 646, 731 to 735, 990 to 994 |
| | >Os01t0949800-01 | 4 | LOC_Os01g72140 | Glutathione S-transferase GST 28 | 61 to 65, 196 to 200, 751 to 755, 854 to 858 |
| | >Os01t0951000-02 | 4 | LOC_Os01g72220 | WD domain, G-beta repeat domain | 26 to 30, 172 to 176, 601 to 605, 611 to 615 |

| | | | | | |
|---|---|---|---|---|---|
| | >Os01t0958100-01 | 4 | LOC_Os01g72800 | chloroplast SRP receptor cpFtsY precursor | 222to226250to254281to285541to545 |
| | >Os01t0958100-02 | 4 | LOC_Os01g72800 | chloroplast SRP receptor cpFtsY precursor | 159to163187to191218to222478to482 |
| | >Os01t0962700-01 | 4 | LOC_Os01g73170 | Peroxidase 12 precursor | 113 to 117, 162 to 166, 212 to 216, 814 to 818 |
| | >Os01t0973300-01 | 4 | LOC_Os01g74180 | Armadillo-like helical domain. Adaptin | 505 to 509, 658 to 662, 717 to 721, 733 to 737 |
| | >Os01t0975000-01 | 4 | LOC_Os01g74370 | Protein of unknown function DUF966 family protein | 720 to 724, 733 to 737, 871 to 875, 948 to 952 |
| | >Os01t0975000-02 | 4 | LOC_Os01g74370 | Protein of unknown function DUF966 family protein | 665 to 669, 678 to 682, 816 to 820, 893 to 897 |
| **TTGAC** | >Os01t0104200-00 | 4 | LOC_Os01g01430 | NAC16 | 71 to 75, 575 to 579, 713 to 717, 736 to 740 |
| | >Os01t0116600-01 | 4 | LOC_Os01g02720 | Protein synthesis factor, GTP-binding domain. elongation factor Tu | 81 to 85, 184 to 188, 277 to 281, 322 to 326 |
| | >Os01t0117600-01 | 4 | LOC_Os01g02830 | Protein kinase, catalytic domain domain. receptor-like kinase ARK1AS | 589 to 593, 785 to 789, 821 to 825, 858 to 862 |
| | >Os01t0130000-01 | 4 | LOC_Os01g03914 | Cation efflux protein family protein. metal tolerance protein C3 | 8 to 12, 17 to 21, 36 to 40, 419 to 423 |
| | >Os01t0136400-01 | 4 | LOC_Os01g04409 | Protein kinase, core domain | 142 to 146, 186 to 190, 400 to 404, 861 to 865 |
| | >Os01t0142800-01 | 4 | LOC_Os01g04950 | Peptide transporter, peptide transporter PTR2 | 137 to 141, 251 to 255, 313 to 317, 868 to 872 |
| | >Os01t0142800-02 | 5 | LOC_Os01g04950 | peptide transporter PTR2 | 34 to 38, 148 to 152, 210 to 214, 765 to 769, 903 to 907 |
| | >Os01t0158400-01 | 4 | LOC_Os01g06500 | Galactose-binding like domain. PHLOEM 2-LIKE A5 | 8 to 12, 176 to 180, 291 to 295, 480 to 484 |
| | >Os01t0158500-01 | 5 | LOC_Os01g06510 | arginyl-tRNA synthetase | 124 to 128, 184 to 188, 532 to 536, 647 to 651, 912 to 916 |
| | >Os01t0160800-01 | 5 | | Protein synthesis inhibitor II , ribosome inactivating protein | 474 to 478, 796 to 800, 809 to 813, 905 to 909, 933 to 937 |
| | >Os01t0162101-00 | 4 | | MATE | 48 to 52, 230 to 234, 239 to 243, 854 to 858 |

| | | | | | |
|---|---|---|---|---|---|
| | >Os01t0162500-01 | 4 | LOC_Os01g06890 | Leucine-rich repeat-containing N-terminal, type 2 domain | 113 to 117, 318 to 322, 646 to 650, 712 to 716 |
| | >Os01t0180800-01 | 4 | LOC_Os01g08560 | Heat shock protein Hsp70 family protein, DnaK family protein | 288 to 292, 319 to 323, 630 to 634, 677 to 681 |
| | >Os01t0180800-02 | 4 | LOC_Os01g08560 | Heat shock protein Hsp70 family protein, DnaK family | 233 to 237, 264 to 268, 575 to 579, 622 to 626 |
| | >Os01t0200200-00 | 4 | LOC_Os01g10310; | Protein of unknown function DUF1677, plant domain | 409 to 413, 562 to 566, 659 to 663, 845 to 849 |
| | >Os01t0222900-01 | 6 | | Curculin-like (mannose-binding) lectin domain | 269 to 273, 768 to 772, 800 to 804, 878 to 882, 898 to 902, 930 to 934 |
| | >Os01t0225500-00 | 4 | LOC_Os01g12570 | 3-methyl-2-oxobutanoate hydroxymethyltransferase | 581 to 585, 761 to 765, 878 to 882, 906 to 910 |
| | >Os01t0234700-01 | 4 | LOC_Os01g13340 | harpin-induced protein 1 domain | 367 to 371, 521 to 525, 530 to 534, 549 to 553 |
| | >Os01t0235200-01 | 4 | LOC_Os01g13404 | ATP binding protein | 684 to 688, 716 to 720, 858 to 862, 864 to 868 |
| | >Os01t0235200-02 | 4 | LOC_Os01g13404 | ATP binding protein | 650 to 654, 682 to 686, 824 to 828, 830 to 834 |
| | >Os01t0242200-01 | 5 | LOC_Os01g14010 | indeterminate domain 8 | 109 to 113, 377 to 381, 449 to 453, 458 to 462, 477 to 481 |
| | >Os01t0243000-01 | 4 | LOC_Os01g14080 | Triacylglycerol lipase. lipase class 3 family protein | 267 to 271, 522 to 526, 884 to 888, 892 to 896 |
| | >Os01t0254000-01 | 4 | LOC_Os01g15010 | NTGB2 , miro | 237 to 241, 474 to 478, 493 to 497, 725 to 729 |
| | >Os01t0269800-00 | 4 | LOC_Os01g16400 | NB-ARC domain | 148 to 152, 167 to 171, 696 to 700, 953 to 957 |
| | >Os01t0304200-00 | 5 | | Transcription factor PCF7 | 43 to 47, 115 to 119, 229 to 233, 308 to 312, 632 to 636 |
| | >Os01t0311300-01 | 7 | | Sorbitol transporter | 43 to 47, 115 to 119, 229 to 233, 308 to 312, 632 to 636 |
| | >Os01t0311500-01 | 4 | LOC_Os01g20940 | PHS1 (PROPYZAMIDE-HYPERSENSITIVE 1); phosphoprotein phosphatase/ protein tyrosine/serine/threonine phosphatase. PHS1 | 489 to 493, 498 to 502, 517 to 521, 945 to 949 |

| | | | | | |
|---|---|---|---|---|---|
| | >Os01t0329400-01 | 4 | LOC_Os01g22600 | Mitochondrial substrate carrier family protein | 90 to 94, 248 to 252, 294 to 298, 679 to 683 |
| | >Os01t0329400-02 | 4 | LOC_Os01g22600 | Mitochondrial substrate carrier family protein | 202 to 206, 411 to 415, 429 to 433, 599 to 603 |
| | >Os01t0351100-00 | 4 | LOC_Os01g24920; | Poly. poly synthetase 2-B | 135 to 139, 224 to 228, 538 to 542, 557 to 561 |
| | >Os01t0375500-00 | 4 | LOC_Os01g27770 | Shikimate biosynthesis protein aroDE | 416 to 420, 569 to 573, 625 to 629, 659 to 663 |
| | >Os01t0530366-01 | 4 | LOC_Os01g34614; | RNA polymerase Rbp10 domain. DNA directed RNA polymerase, 7 kDa subunit domain | 204 to 208, 807 to 811, 842 to 846, 928 to 932 |
| | >Os01t0532200-01 | 4 | LOC_Os01g34780; | FF domain domain. | 203 to 207, 250 to 254, 371 to 375, 732 to 736 |
| | >Os01t0536000-01 | 4 | LOC_Os01g35184 | CBL (CALCINEURIN B-LIKE )-interacting serine/threonine-protein kinase 24 PROTEIN-INTERACTING PROTEIN KINASE 8;CGSNL. 10 - CAMK includes calcium/calmodulin depedent | 313 to 317, 372 to 376, 713 to 717, 781 to 785 |
| | >Os01t0549400-04 | 5 | LOC_Os01g36860 | RNA helicase-like protein DB10. DEAD-box ATP-dependent RNA helicase 40 | 249 to 253, 704 to 708, 730 to 734, 926 to 930, 949 to 953 |
| | >Os01t0558600-01 | 4 | LOC_Os01g37800; | Ras-related protein RIC | 222 to 226, 228 to 232, 248 to 252, 517 to 521 |
| | >Os01t0563000-05 | 5 | LOC_Os01g38229 | Peptidyl-prolyl cis-trans isomerase, FKBP-type domain | 459 to 463, 477 to 481, 683 to 687, 891 to 895, 965 to 969 |
| | >Os01t0584032-00 | 4 | | Skin secretory protein xP2 | 68 to 72, 77 to 81, 96 to 100, 345 to 349 |
| | >Os01t0587900-00 | 5 | LOC_Os01g40540 | OSIGBa0125M19.6 protein. lectin receptor-type protein kinase | 80 to 84, 112 to 116, 264 to 268, 369 to 373, 535 to 539 |
| | >Os01t0588400-01 | 4 | LOC_Os01g40580 | Band 7 protein family protein. hypersensitive-induced response protein | 315 to 319, 364 to 368, 629 to 633, 724 to 728 |
| | >Os01t0598200-01 | 4 | LOC_Os01g41510 | Calcineurin B-like protein 8. calcineurin B | 59 to 63, 76 to 80, 301 to 305, 762 to 766 |

| | | | | | |
|---|---|---|---|---|---|
| | >Os01t0603500-00 | 4 | LOC_Os01g41910 | Cf2/Cf5-like disease resistance protein receptor-like protein kinase 5 precursor | 655 to 659, 730 to 734, 761 to 765, 978 to 982 |
| | >Os01t0624700-01 | 4 | LOC_Os01g43550 | WRKY 12 | 250 to 254, 497 to 501, 594 to 598, 772 to 776 |
| | >Os01t0633000-01 | 5 | LOC_Os01g44210 | 50S ribosomal protein L31 | 65 to 69, 409 to 413, 430 to 434, 521 to 525, 588 to 592 |
| | >Os01t0633500-00 | 4 | LOC_Os01g44260; | Dihydroflavonol-4-reductase | 61 to 65, 125 to 129, 378 to 382, 718 to 722 |
| | >Os01t0635200-01 | 4 | LOC_Os01g44390 | Homeodomain-like | 112 to 116, 129 to 133, 553 to 557, 895 to 899 |
| | >Os01t0646700-03 | 4 | LOC_Os01g45880 | OSIGBa0105P02.4 protein. retrotransposon protein | 465 to 469, 594 to 598, 667 to 671, 804 to 808 |
| | >Os01t0653100-00 | 5 | LOC_Os01g46410 | Leaf senescence related protein | 76 to 80, 236 to 240, 443 to 447, 502 to 506, 529 to 533 |
| | >Os01t0678400-01 | 4 | LOC_Os01g48660 | Protein of unknown function DUF2921 domain | 153 to 157, 382 to 386, 522 to 526, 850 to 854 |
| | >Os01t0694300-01 | 4 | LOC_Os01g49940 | T17H3.9. ESP4 | 420 to 424, 528 to 532, 789 to 793, 926 to 930 |
| | >Os01t0712600-00 | 4 | LOC_Os01g51530 | Methyltransferase | 211 to 215, 360 to 364, 392 to 396, 538 to 542 |
| | >Os01t0712800-00 | 5 | LOC_Os01g51550 | peroxidase family protein | 46 to 50, 415 to 419, 722 to 726, 736 to 740, 921 to 925 |
| | >Os01t0721800-01 | 4 | LOC_Os01g52380 | Protein kinase-like domain | 380 to 384, 519 to 523, 729 to 733, 810 to 814 |
| | >Os01t0721800-01 | 4 | LOC_Os01g52380 | Protein kinase-like domain | 380 to 384, 519 to 523, 729 to 733, 810 to 814 |
| | >Os01t0737600-00 | 4 | LOC_Os01g53580 | 3-5 exonuclease domain | 263 to 267, 392 to 396, 413 to 417, 419 to 423 |
| | >Os01t0742400-01 | 4 | LOC_Os01g53920 | Protein kinase, core domain. receptor-like protein kinase 5 precursor | 265 to 269, 272 to 276, 342 to 346, 782 to 786 |
| | >Os01t0742400-02 | 4 | LOC_Os01g53920 | Protein kinase, core domain. receptor-like protein kinase 5 precursor | 262 to 266, 269 to 273, 339 to 343, 779 to 783 |

| | | | | | |
|---|---|---|---|---|---|
| | >Os01t0748950-01 | 4 | LOC_Os01g54515 | Peptide transporter PTR2 | 406 to 410, 613 to 617, 937 to 941, 968 to 972 |
| | >Os01t0760300-01 | 4 | LOC_Os01g55520 | Pollen-specific kinase partner protein. ATROPGEF7/ROPGEF7 | 391 to 395, 619 to 623, 735 to 739, 830 to 834 |
| | >Os01t0763750-02 | 4 | LOC_Os01g55799 | Exo70 exocyst complex subunit domain | 54 to 58, 842 to 846, 923 to 927, 928 to 932 |
| | >Os01t0764950-01 | 5 | LOC_Os01g55960 | Pentatricopeptide repeat domain | 2 to 6, 74 to 78, 382 to 386, 616 to 620, 741 to 745 |
| | >Os01t0772700-00 | 4 | LOC_Os01g56590 | Armadillo-type fold domain. guanine nucleotide exchange family protein | 205 to 209, 501 to 505, 651 to 655, 770 to 774 |
| | >Os01t0784700-00 | 4 | LOC_Os01g57560 | Serine/threonine protein kinase-related domain | 130 to 134, 266 to 270, 309 to 313, 751 to 755 |
| | >Os01t0788900-01 | 5 | LOC_Os01g57900 | Pentatricopeptide repeat domain | 287 to 291, 534 to 538, 566 to 570, 715 to 719, 721 to 725 |
| | >Os01t0788900-02 | 4 | LOC_Os01g57900 | Pentatricopeptide repeat domain | 115 to 119, 147 to 151, 296 to 300, 302 to 306 |
| | >Os01t0793100-00 | 4 | LOC_Os01g58070 | Polyphenol oxidase | 345 to 349, 503 to 507, 621 to 625, 995 to 999 |
| | >Os01t0802000-01 | 4 | LOC_Os01g58780 | Zinc finger, RING/FYVE/PHD-type domain. zinc finger, C3HC4 type | 60 to 64, 167 to 171, 237 to 241, 962 to 966 |
| | >Os01t0810600-00 | 4 | LOC_Os01g59560 | Protein kinase domain | 483 to 487, 502 to 506, 514 to 518, 533 to 537 |
| | >Os01t0813500-01 | 4 | LOC_Os01g59560; | cDNA clone: J100063K14 | 189 to 193, 725 to 729, 749 to 753, 785 to 789 |
| | >Os01t0817000-01 | 6 | LOC_Os01g60110 | Protein of unknown function DUF607 family protein | 11 to 15, 46 to 50, 159 to 163, 401 to 405, 650 to 654, 948 to 952 |
| | >Os01t0827300-01 | 4 | | Laccase precursor | 45 to 49, 256 to 260, 532 to 536, 668 to 672 |
| | >Os01t0842500-01 | 4 | LOC_Os01g61160; | Laccase precursor protein | 41 to 45, 69 to 73, 366 to 370, 717 to 721 |
| | >Os01t0852100-01 | 4 | LOC_Os01g62490; | Laccase precursor protein | 80 to 84, 370 to 374, 687 to 691, 848 to 852 |
| | >Os01t0853700-01 | 5 | LOC_Os01g63280 | Tyrosine protein kinase domain | 86 to 90, 332 to 336, 599 to 603, 643 to 647, 726 to 730 |
| | >Os01t0855900-01 | 4 | LOC_Os01g63460 | MCB1 protein. MYB family | 251 to 255, 387 to 391, 394 to 398, 500 |

| | | | | | |
|---|---|---|---|---|---|
| | | | | transcription factor | to 504 |
| | >Os01t0856850-00 | 4 | LOC_Os01g63710 | CDC6 - Putative DNA replication initiation protein | 100 to 104, 109 to 113, 683 to 687, 833 to 837 |
| | >Os01t0857000-01 | 5 | LOC_Os01g63820 | Double-stranded RNA-binding-like domain | 134 to 138, 389 to 393, 419 to 423, 449 to 453, 807 to 811 |
| | >Os01t0857000-02 | 4 | LOC_Os01g63820 | Double-stranded RNA-binding-like domain | 126 to 130, 156 to 160, 186 to 190, 544 to 548 |
| | >Os01t0872300-00 | 4 | LOC_Os01g65190; | POT domain containing peptide transporter | 281 to 285, 430 to 434, 635 to 639, 676 to 680 |
| | >Os01t0873200-01 | 4 | | Amidophosphoribosyltransferase, chloroplast precursor (Glutamine phosphoribosylpyrophosphate amidotransferase) | 100 to 104, 162 to 166, 260 to 264, 486 to 490 |
| | >Os01t0878300-02 | 4 | LOC_Os01g65650 | Protein kinase, core domain. receptor-like protein kinase HAIKU2 precursor | 365 to 369, 414 to 418, 758 to 762, 993 to 997 |
| | >Os01t0878400-01 | 4 | LOC_Os01g65660; | Amino acid transporter, transmembrane domain. amino acid transporter | 139 to 143, 201 to 205, 241 to 245, 386 to 390 |
| | >Os01t0883000-00 | 5 | LOC_Os01g66020 | protein kinase family protein | 149 to 153, 192 to 196, 473 to 477, 519 to 523, 871 to 875 |
| | >Os01t0899425-00 | 4 | | Ribulose 1,5-bisphosphate carboxylase/oxygenase large subunit | 87 to 91, 259 to 263, 272 to 276, 346 to 350 |
| | >Os01t0904400-01 | 6 | LOC_Os01g67740 | Chromosome assembly protein homolog - segregation protein | 141 to 145, 302 to 306, 450 to 454, 482 to 486, 631 to 635, 637 to 64 |
| | >Os01t0911000-01 | 4 | LOC_Os01g68310 | sas10/Utp3 family protein | 154 to 158, 345 to 349, 507 to 511, 564 to 568 |
| | >Os01t0916000-01 | 4 | LOC_Os01g68730 | RNA-binding protein FUS | 21 to 25, 35 to 39, 244 to 248, 931 to 935 |
| | >Os01t0917500-01 | 6 | LOC_Os01g68870 | leucine-rich repeat receptor protein kinase EXS precursor | 55 to 59, 318 to 322, 330 to 334, 543 to 547, 553 to 557, 574 to 578 |
| | >Os01t0921000-01 | 5 | LOC_Os01g69190; | Nucleotide-diphospho-sugar transferase domain. regulatory protein | 430 to 434, 437 to 441, 583 to 587, 670 to 674, 843 to 847 |

| | | | | | |
|---|---|---|---|---|---|
| | >Os01t0921400-01 | 4 | LOC_Os01g69230 | exo70 exocyst complex subunit domain | 132 to 136, 281 to 285, 383 to 387, 404 to 408 |
| | >Os01t0925100-01 | 4 | LOC_Os01g70080 | NB-ARC domain | 75 to 79, 321 to 325, 918 to 922 , 930 to 934 |
| | >Os01t0928700-00 | 4 | LOC_Os01g70370 | Serine palmitoy ltransferase 2 | 304 to 308, 351 to 355, 360 to 364, 378 to 382 |
| | >Os01t0928800-01 | 4 | LOC_Os01g70380 | Serine palmitoyl transferase 2 | 231 to 235, 437 to 441, 520 to 524, 745 to 749 |
| | >Os01t0931100-01 | 4 | LOC_Os01g70550 | Heparan-alpha-glucosaminide N-acetyltransferase | 51 to 55, 409 to 413, 604 to 608, 691 to 695 |
| | >Os01t0937300-01 | 7 | LOC_Os01g71100 | NB-ARC domain | 165 to 169, 175 to 179, 277 to 281, 538 to 542, 566 to 570, 744 to 748, 786 to 790 |
| | >Os01t0937300-02 | 7 | LOC_Os01g71100 | NB-ARC domain | 159 to 163, 169 to 173, 271 to 275, 532 to 536 560 to 564, 738 to 742, 780 to 784 |
| | >Os01t0940800-01 | 4 | LOC_Os01g71350 | Beta-1,3-glucanase precursor. glycosyl hydrolases family 17 | 220 to 224, 469 to 473, 641 to 645, 730 to 734 |
| | >Os01t0946700-01 | 4 | LOC_Os01g71830 | endo-1, 3-beta-glucanase 1 | 176 to 180, 221 to 225, 548 to 552, 620 to 624 |
| | >Os01t0952800-02 | 4 | LOC_Os01g72370 | Achaete-scute transcription factor related domain. helix-loop-helix DNA-binding domain | 169 to 173, 459 to 463, 634 to 638, 817 to 821 |
| | >Os01t0953801-00 | 4 | LOC_Os01g72450; | Domain of unknown function DUF296 domain. | 201 to 205, 602 to 606, 712 to 716, 743 to 747 |
| | >Os01t0955400-00 | 5 | LOC_Os01g72540; | OsCML23 - Calmodulin-related calcium sensor protein | 393 to 397, 429 to 433, 641 to 645, 766 to 770, 825 to 829 |
| | >Os01t0955500-00 | 4 | LOC_Os01g72550 | OsCML19 - Calmodulin-related calcium sensor protein | 78 to 82, 277 to 281, 789 to 793, 802 to 806 |
| | >Os01t0966000-01 | 5 | LOC_Os01g73514 | Plasma membrane H – ATPase. 5-oxoprolinase | 236 to 240, 442 to 446, 615 to 619, 624 to 628, 643 to 647 |
| | >Os01t0966000-02 | 5 | LOC_Os01g73514 | Plasma membrane H – ATPase. 5-oxoprolinase | 204 to 208, 410 to 414, 583 to 587, 592 to 596, 611 to 615 |

| | | | | | |
|---|---|---|---|---|---|
| | >Os01t0966000-03 | 4 | LOC_Os01g73514 | Plasma membrane H – ATPase. 5-oxoprolinase | 517 to 521, 642 to 646, 675 to 679, 916 to 920 |
| | >Os01t0966100-01 | 4 | LOC_Os01g73530 | Peroxisomal ABC transporter | 239 to 243, 288 to 292, 568 to 572, 598 to 602 |
| | >Os01t0969200-01 | 4 | LOC_Os01g73970 | Protein of unknown function DUF707 family protein. lysine ketoglutarate reductase trans-splicing related 1 | 83 to 87, 167 to 171, 326 to 330, 637 to 641 |
| | >Os01t0975000-02 | 5 | LOC_Os01g74370 | Domain of unknown function DUF966 domain | 375 to 379, 643 to 647, 664 to 668, 755 to 759, 815 to 819 |
| **TTGACA** | >Os01t0853700-01 | 4 | LOC_Os01g63460 | MCB1 protein | 86 to 91, 332 to 337, 643 to 648, 726 to 731 |
| | >Os01t0857000-01 | 4 | LOC_Os01g63820 | C-Terminal Domain Phosphatase-Like 2 | 134 to 139, 389 to 394, 419 to 424, 449 to 454 |
| | >Os01t0917500-01 | 4 | LOC_Os01g68870 | multiple sporogeneous cells-1 | 318 to 323, 330 to 335, 543 to 548, 553 to 558 |
| | >Os01t0921400-01 | 4 | LOC_Os01g69230 | exo70 exocyst complex subunit | 132 to 137, 281 to 286, 383 to 388, 404 to 409 |
| **TTGACC** | >Os01t0764950-01 | 4 | LOC_Os01g55960 | Pentatricopeptide repeat domain | 74 to 79, 382 to 387, 616 to 621, 741 to 746 |
| **TTGACT** | >Os01t0633000-01 | 4 | LOC_Os01g44210 | 50S ribosomal protein L31 | 409 to 414, 430 to 435, 521 to 526, 588 to 593 |
| | >Os01t0788900-01 | 4 | LOC_Os01g57900 | Pentatricopeptide repeat domain | 287 to 292, 534 to 539, 566 to 571, 715 to 720 |
| | >Os01t0928800-01 | 4 | LOC_Os01g70380 | serine palmitoyltransferase 2 | 231 to 236, 437 to 442, 520 to 525, 745 to 750 |
| **Chromosome 2** | | | | | |
| **Motif** | **ID** | **Frequency** | **Locus ID** | **Protein name** | **Positions in 1k promoter region** |
| **CTGACC** | >Os02t0741900-01 | 7 | LOC_Os02g50830 | T23E23.20. | 111 to 116, 134 to 139, 156 to 161, 179 to 184, 274 to 279, 294 to 299, 393 to 398 |
| | >Os02t0674800-01 | 4 | LOC_Os02g45250 | Rice outmost cell-specific gene5 (ROC5) | 423 to 428, 592 to 597, 780 to 785, 796 to 801 |
| **TTTGACT** | >Os02t0561900-00 | 4 | LOC_Os02g35440 | E3 ubiquitin-protein ligase EL5 | 407 to 413, 439 to 445, 581 to 587, 587 to 593 |

| | | | | | |
|---|---|---|---|---|---|
| | >Os02t0158100-01 | 5 | LOC_Os02g06340 | EF hand domain | 241 to 246, 245 to 250, 249 to 254, 253 to 258, 337 to 342 |
| | >Os02t0272800-01 | 4 | LOC_Os02g17292 | Retrotransposon protein | 176 to 181, 216 to 221, 775 to 780, 923 to 928 |
| **GTACGTAC** | >Os02t0791400-01 | 4 | LOC_Os02g54880 | Cytochrome c oxidase | 449 to 456, 453 to 460, 457 to 464, 600 to 607 |
| **TGACC** | >Os02t0152800-01 | 5 | LOC_Os02g05880 | RNA polymerase Rpb1, domain 1 | 820 to 824, 830 to 834, 932 to 936, 942 to 946, 972 to 976 |
| | >Os02t0193000-01 | 5 | LOC_Os02g09960 | Protein kinase-like domain. LYK8 | 16 to 20, 249 to 253, 290 to 294, 425 to 429, 947 to 951 |
| | >Os02t0224800-01 | 4 | LOC_Os02g13150 | Ndr family protein. pollen-specific protein SF21 | 169 to 173, 177 to 181, 200 to 204, 678 to 682 |
| | >Os02t0227700-01 | 4 | LOC_Os02g13430 | Serine/threonine protein kinase-related domain. receptor-like protein kinase 5 precursor | 577 to 581, 608 to 612, 885 to 889, 891 to 895 |
| | >Os02t0251900-01 | 4 | LOC_Os02g15290 | Tobacco rattle virus-induced protein variant 2. VQ domain | 764 to 768, 781 to 785, 805 to 809, 945 to 949 |
| | >Os02t0277600-01 | 4 | LOC_Os02g17700 | Iron-sulfur cluster protein 21 | 66 to 70, 243 to 247, 308 to 312, 686 to 690 |
| | >Os02t0281000-02 | 5 | LOC_Os02g17970 | Predicted protein | 154 to 158, 298 to 302, 406 to 410, 502 to 506, 709 to 713 |
| | >Os02t0311150-00 | 5 | LOC_Os02g20720 | BTB/POZ-like domain | 95 to 99, 384 to 388, 446 to 450, 623 to 627, 662 to 666 |
| | >Os02t0510100-01 | 4 | LOC_Os02g30624 | Cyclophilin 7 | 97 to 101, 334 to 338, 511 to 515, 535 to 539 |
| | >Os02t0510100-02 | 4 | LOC_Os02g30624 | U6 snRNA-associated Sm-like protein LSm6 | 38 to 42, 275 to 279, 452 to 456, 476 to 480 |
| | >Os02t0511900-00 | 4 | LOC_Os02g30800 | DNA polymerase | 457 to 461, 495 to 499, 645 to 649, 669 to 673 |
| | >Os02t0626400-03 | 4 | LOC_Os02g41650 | Phenylalanine ammonia-lyase | 252 to 256, 431 to 435, 807 to 811, 993 to 997 |
| | >Os02t0627100-01 | 5 | LOC_Os02g41680 | Phenylalanine ammonia-lyase | 104 to 108, 333 to 337, 362 to 366, 466 to 470, 784 to 788 |
| | >Os02t0628200-01 | 5 | LOC_Os02g41780 | DUF250 domain | 53 to 57, 220 to 224, 280 to 284, 457 to 461, 881 to 885 |

| | | | | | |
|---|---|---|---|---|---|
| | >Os02t0628800-01 | 4 | LOC_Os02g41820 | Ubiquitin-like protein 5 | 113 to 117, 293 to 297, 330 to 334, 595 to 599 |
| | >Os02t0658900-01 | 4 | LOC_Os02g44111 | sec20 domain | 85 to 89, 545 to 549, 674 to 678, 795 to 799 |
| | >Os02t0661400-00 | 4 | LOC_Os02g44260 | Protein of unknown function DUF597 domain | 253 to 257, 379 to 383, 940 to 944, 992 to 996 |
| | >Os02t0674800-01 | 5 | LOC_Os02g45250 | OCL1 homeobox protein. homeobox and START domains | 424 to 428, 593 to 597, 781 to 785, 797 to 801, 911 to 915 |
| | >Os02t0675700-01 | 4 | LOC_Os02g45310 | DUF248, methyltransferase putative family protein. dehydration response related protein | 201 to 205, 329 to 333, 885 to 889, 956 to 960 |
| | >Os02t0684000-00 | 4 | LOC_Os02g45880 | Tetratricopeptide-like helical domain | 79 to 83, 94 to 98, 470 to 474, 917 to 921 |
| | >Os02t0741900-01 | 7 | LOC_Os02g50830 | T23E23.20 | 112 to 116, 135 to 139, 157 to 161, 180 to 184, 275 to 279, 295 to 299, 394 to 398 |
| | >Os02t0744000-01 | 4 | LOC_Os02g51020 | NDF6 (NDH DEPENDENT FLOW 6) | 80 to 84, 835 to 839, 841 to 845, 946 to 950 |
| | >Os02t0748300-02 | 4 | LOC_Os02g51350 | VMP3 protein. OsFBK10 - F-box domain and kelch repeat | 274 to 278, 539 to 543, 629 to 633, 921 to 925 |
| | >Os02t0771400-00 | 5 | LOC_Os02g53160 | Protein-tyrosine phosphatase, SIW14-like domain. tyrosine phosphatase family protein | 48 to 52, 53 to 57, 74to78, 576 to 580, 665 to 669 |
| | >Os02t0822300-00 | 4 | LOC_Os02g57640 | RNA-binding protein Nova-1.KH domain | 123 to 127, 480 to 484, 498 to 502, 855 to 859 |
| | >Os02t0823800-02 | 4 | LOC_Os02g57780 | Smg8/Smg9 domain | 12 to 16, 41 to 45, 356 to 360, 983 to 987 |
| **TGACT** | >Os02+B34:F63t0106 600-01 | 5 | LOC_Os02g01700 | Zinc finger, RING/FYVE/PHD-type domain. RNA recognition motif | 8 to 12, 92 to 96, 603 to 607 634 to 638, 991 to 995 |
| | >Os02t0115600-02 | 4 | LOC_Os02g02390 | S1, RNA binding domain | 117 to 121, 527 to 531, 578 to 582, 897 to 901 |
| | >Os02t0153200-01 | 5 | LOC_Os02g05920 | Protein kinase, core domain. phytosulfokine receptor precursor | 262 to 266, 529 to 533, 925 to 929, 941 to 945, 947 to 951 |

| | | | | | |
|---|---|---|---|---|---|
| | >Os02t0158100-01 | 6 | LOC_Os02g06340 | EF hand domain. EH domain 1 | 238 to 242, 242 to 246, 246 to 250, 250 to 254, 254 to 258, 338 to 342 |
| | >Os02t0186800-00 | 4 | LOC_Os02g09390 | Cytochrome P450 family protein | 231 to 235, 412 to 416, 418 to 422, 979 to 983 |
| | >Os02t0204000-01 | 4 | LOC_Os02g10940 | Tetratricopeptide-like helical domain | 46 to 50, 192 to 196, 198 to 202, 713 to 717 |
| | >Os02t0259900-01 | 5 | LOC_Os02g14840 | Inwardly rectifying potassium channel. potassium channel KAT1 | 2 to 6, 34 to 38, 183 to 187, 761 to 765, 982 to 986 |
| | >Os02t0281200-02 | 4 | LOC_Os02g17292 | Retrotransposon protein | 22 to 26, 80 to 84, 569 to 573, 957 to 961 |
| | >Os02t0293050-00 | 4 | LOC_Os02g18000 | NBS-LRR protein disease resistance protein RGA2 | 129 to 133, 599 to 603, 779 to 783, 786 to 790 |
| | >Os02t0311150-00 | 4 | LOC_Os02g19470 | Protein of unknown function DUF594 family protein | 268 to 272, 296 to 300, 310 to 314, 923 to 927 |
| | >Os02t0326500-01 | 4 | LOC_Os02g20720 | BTB/POZ-like domain. MBTB2 - Bric-a-Brac, Tramtrack, Broad Complex BTB domain with Meprin and TRAF Homology MATH domain | 340 to 344, 361 to 365, 379 to 383, 879 to 883 |
| | >Os02t0466600-01 | 4 | LOC_Os02g22084 | Targeting protein-related | 248 to 252, 277 to 281, 291 to 295, 719 to 723 |
| | >Os02t0477100-01 | 4 | LOC_Os02g26800 | Aminotransferase-like, plant mobile domain | 263 to 267, 393 to 397, 523 to 527, 640 to 644 |
| | >Os02t0508500-01 | 4 | LOC_Os02g28580 | WRC domain | 263 to 267, 393 to 397, 523 to 527, 640 to 644 |
| | >Os02t0537775-00 | 4 | LOC_Os02g30530 | Transposon protein | 14 to 18, 188 to 192, 199 to 203, 520 to 524 |
| | >Os02t0549850-00 | 5 | | Myosin | 434 to 438, 451 to 455, 496 to 500, 527 to 531, 544 to 548 |
| | >Os02t0554200-00 | 4 | LOC_Os02g34830 | Tetratricopeptide-like helical | 42 to 46, 222 to 226, 428 to 432, 820 to 824 |
| | >Os02t0556900-01 | 4 | LOC_Os02g35100 | H0525C06.6 protein. DDT domain | 302 to 306, 447 to 451, 451 to 455, 455 to 459 |

| | | | | | |
|---|---|---|---|---|---|
| | >Os02t0561900-00 | 5 | LOC_Os02g35100 | H0525C06.6 protein. DDT domain | 128 to 132, 409 to 413, 441 to 445, 583 to 587, 589 to 593 |
| | >Os02t0573400-01 | 4 | LOC_Os02g35440 | E3 ubiquitin-protein ligase EL5. RING-H2 finger protein ATL4O precursor | 65 to 69, 95 to 99, 175 to 179, 751 to 755 |
| | >Os02t0593400-01 | 4 | LOC_Os02g36400 | Peptidase C19, ubiquitin carboxyl-terminal hydrolase 2 family protein. ubiquitin carboxyl-terminal hydrolase domain | 23 to 27, 185 to 189, 515 to 519, 684 to 688 |
| | >Os02t0672200-02 | 4 | LOC_Os02g44108 | Beta-expansin | 292 to 296, 296 to 300, 807 to 811, 892 to 896 |
| | >Os02t0690550-00 | 4 | LOC_Os02g45070 | AGO1 homologous protein. PINHEAD | 5 to 9, 238 to 242, 273 to 277, 597 to 601 |
| | >Os02t0694400-01 | 5 | LOC_Os02g46500 | Zinc finger, RING/FYVE/PHD-type domain | 210 to 214, 472 to 476, 516 to 520, 982 to 986, 986 to 990 |
| | >Os02t0694400-02 | 5 | LOC_Os02g46720 | ara54-like RING finger protein | 210 to 214, 472 to 476, 516 to 520, 982 to 986, 986 to 990 |
| | >Os02t0700600-03 | 4 | LOC_Os02g47220 | GAMYB-binding protein. CPuORF20 - conserved peptide uORF | 4 to 8, 715 to 719, 790 to 794, 866 to 870 |
| | >Os02t0704800-01 | 6 | LOC_Os02g47590 | Ornithine carbamoyltransferase | 157 to 161, 161 to 165, 208 to 212, 215 to 219, 629 to 633, 901 to 905 |
| | >Os02t0708300-01 | 4 | LOC_Os02g47870 | Zinc finger, RING/FYVE/PHD-type domain. anaphase-promoting complex subunit 11 | 364 to 368, 498 to 502, 529 to 533, 683 to 687 |
| | >Os02t0714500-01 | 4 | LOC_Os02g48380 | Tetratricopeptide-like helical domain | 623 to 627, 774 to 778, 865 to 869, 924 to 928 |
| | >Os02t0722500-01 | 4 | LOC_Os02g49070 | Arf GTPase activating protein family protein. ZAC | 212 to 216, 231 to 235, 395 to 399, 504 to 508 |
| | >Os02t0730800-01 | 4 | LOC_Os02g49820 | Nucleotide-binding, alpha-beta plait domain. RNA recognition motif family protein | 201 to 205, 206 to 210, 246 to 250, 660 to 664 |
| | >Os02t0756600-01 | 4 | LOC_Os02g52000 | Phi-1 protein. phosphate-induced protein 1 conserved region domain | 126 to 130, 361 to 365, 856 to 860, 985 to 989 |

| | | | | | |
|---|---|---|---|---|---|
| | >Os02t0767500-01 | 4 | LOC_Os02g52860 | Mitochondrial phosphate transporter. phosphate carrier protein, mitochondrial precursor | 180 to 184, 397 to 401, 593 to 597, 759 to 763 |
| | >Os02t0769200-01 | 4 | LOC_Os02g53000 | Erwinia induced protein 1. lysM domain-containing GPI-anchored protein precursor | 34 to 38, 329 to 333, 620 to 624, 888 to 892 |
| | >Os02t0786400-00 | 4 | LOC_Os02g54520 | DNA-binding transcription factor blind 1. MYB family transcription factor | 342 to 346, 538 to 542, 757 to 761, 762 to 766 |
| | >Os02t0802200-01 | 5 | LOC_Os02g55870 | Glycoside hydrolase family 79, N-terminal protein. heparanase-like protein precursor | 76 to 80, 582 to 586, 762 to 766, 768 to 772, 889 to 893 |
| | >Os02t0809700-01 | 4 | | Pi homeostasis | 172 to 176, 179 to 183, 691 to 695, 765 to 769 |
| | >Os02t0813166-00 | 4 | LOC_Os02g56820 | Cyclin-like F-box domain OsFBX69 - F-box domain | 144 to 148, 174 to 178, 486 to 490, 816 to 820 |
| **TTGAC** | >Os02t0110000-01 | 4 | LOC_Os02g01980 | Lipase, GDSL domain. GDSL-like lipase/acylhydrolase | 187 to 191, 311 to 315, 324 to 328, 929 to 933 |
| | >Os02t0111600-01 | 4 | LOC_Os02g02120 | Serine/threonine protein kinase-related domain. OsWAK11 - OsWAK receptor-like protein kinase | 27 to 31, 98 to 102, 159 to 163, 248 to 252 |
| | >Os02t0113200-00 | 5 | LOC_Os02g02230 | Cytochrome P450-like protein | 160 to 164, 614 to 618, 638 to 642, 957 to 961, 979 to 983 |
| | >Os02t0114000-01 | 4 | LOC_Os02g02290 | Bromodomain domain SNF2 family N-terminal domain | 114 to 118, 207 to 211, 552 to 556, 894 to 898 |
| | >Os02t0115600-02 | 4 | LOC_Os02g02390 | S1, RNA binding domain | 116 to 120, 526 to 530, 577 to 581, 896 to 900 |
| | >Os02t0126400-01 | 4 | LOC_Os02g03410 | Protein kinase CPK1. CAMK_CAMK_like.12 - CAMK includes calcium/calmodulin depedent protein kinases | 451 to 455, 911 to 915, 927 to 931, 945 to 949 |
| | >Os02t0126400-02 | 4 | LOC_Os02g03410 | Protein kinase CPK1. (Os02t0126400-01); Calcium-dependent protein kinase. CAMK_CAMK_like.12 - CAMK includes calcium/calmodulin depedent | 439 to 443, 899 to 903, 915 to 919, 933 to 937 |

| | | | | | |
|---|---|---|---|---|---|
| | | | | protein kinases | |
| | >Os02t0133100-01 | 5 | LOC_Os02g04020 | Phosphatidylinositol transfer-like protein III | 13 to 17, 627 to 631, 668 to 672, 891 to 895, 936 to 940 |
| | >Os02t0136000-00 | 4 | LOC_Os02g04340 | Phox/Bem1p domain. NIN-like protein 2 | 106 to 110, 221 to 225, 242 to 246, 920 to 924 |
| | >Os02t0149900-00 | 4 | LOC_Os02g05640 | Putative homeobox-leucine zipper protein HOX26 | 30 to 34, 220 to 224, 343 to 347, 707 to 711 |
| | >Os02t0149900-00 | 4 | LOC_Os02g05640 | Putative homeobox-leucine zipper protein HOX26 | 30 to 34, 220 to 224, 343 to 347, 707 to 711 |
| | >Os02t0158400-01 | 4 | LOC_Os02g06370 | ssDNA-binding transcriptional regulator family protein. whirly transcription factor domain | 418 to 422, 521 to 525, 530 to 534, 549 to 553 |
| | >Os02t0159700-03 | 4 | LOC_Os02g06480 | Electron transport SCO1/SenC family protein. SCO1 protein homolog, mitochondrial precursor | 417 to 421, 530 to 534, 539 to 543, 558 to 562 |
| | >Os02t0162733-00 | 4 | LOC_Os02g06720 | WD-40 repeat family protein / beige-related. WD domain | 194 to 198, 309 to 313,454 to 458, 857 to 861 |
| | >Os02t0164800-01 | 4 | LOC_Os02g06890 | Zinc finger, C2H2-type domain. OTU-like cysteine protease family protein | 89 to 93, 324 to 328, 641 to 645, 935 to 939 |
| | >Os02t0173500-01 | 4 | LOC_Os02g07720 | Cholinephosphate cytidylyltransferase | 390 to 394, 588 to 592, 694 to 698, 763 to 767 |
| | >Os02t0174100-02 | 4 | LOC_Os02g07780 | Soform 2 of Squamosa promoter-binding-like protein 4. OsSPL4 - SBP-box gene family member | 116 to 120, 125 to 129, 144 to 148, 565 to 569 |
| | >Os02t0174100-03 | 4 | LOC_Os02g07780 | Isoform 2 of Squamosa promoter-binding-like protein 4. OsSPL4 - SBP-box gene family member | 80 to 84, 89 to 93, 108 to 112, 529 to 533 |
| | >Os02t0174100-01 | 4 | LOC_Os02g07780 | Isoform 2 of Squamosa promoter-binding-like protein 4. OsSPL4 - SBP-box gene family member | 123 to 127, 132 to 136, 151 to 155, 572 to 576 |
| | >Os02t0184400-01 | 4 | LOC_Os02g09170 | sucrose phosphate synthase 2 | 62 to 66, 98 to 102, 103 to 107, 899 to 903 |
| | >Os02t0184400-02 | 4 | LOC_Os02g09170 | sucrose phosphate synthase 2 | 27 to 31, 63 to 67, 68 to 72, 864 to 868 |

| | | | | | |
|---|---|---|---|---|---|
| | >Os02t0186800-00 | 4 | LOC_Os02g09390 | Cytochrome P450 family protein | 230 to 234, 411 to 415, 417 to 421, 978 to 982 |
| | >Os02t0191700-00 | 5 | LOC_Os02g09840 | EF-HAND 2 domain. serine/threonine-protein phosphatase 2A regulatory subunit B subunitbeta | 258 to 262, 365 to 369, 703 to 707, 832 to 836, 884 to 888 |
| | >Os02t0193000-01 | 4 | LOC_Os02g09960 | Protein kinase-like domain LYK8 | 248 to 252, 289 to 293, 932 to 936, 946 to 950 |
| | >Os02t0204000-01 | 5 | LOC_Os02g10940 | Tetratricopeptide-like helical domain | 45 to 49, 191 to 195, 197 to 201, 642 to 646, 661 to 665 |
| | >Os02t0216000-00 | 5 | LOC_Os02g12450 | Receptor-like protein kinase 2 precursor | 393 to 397, 654 to 658, 703 to 707, 762 to 766, 829 to 833 |
| | >Os02t0216300-01 | 4 | LOC_Os02g12480 | cDNA clone:J023088C01, full insert sequence | 112 to 116, 444 to 448, 717 to 721, 916 to 920 |
| | >Os02t0221900-01 | 4 | LOC_Os02g12890 | Cytochrome P450 family protein | 59 to 63, 71 to 75, 128 to 132, 432 to 436 |
| | >Os02t0231700-00 | 4 | LOC_Os02g13780 | Protein kinase, catalytic domain domain. receptor-like protein kinase 2 precursor | 165 to 169, 288 to 292, 824 to 828, 905 to 909 |
| | >Os02t0233333-00 | 4 | LOC_Os02g13910 | Predicted protein. retrotransposon protein, Ty1-copia subclass | 188 to 192, 206 to 210, 782 to 786, 969 to 973 |
| | >Os02t0244700-04 | 4 | LOC_Os02g14770 | Phosphoenolpyruvate carboxylase 1 | 4 to 8, 165 to 169, 337 to 341, 601 to 605 |
| | >Os02t0245800-01 | 4 | LOC_Os02g14840 | Inwardly rectifying potassium channel. potassium channel KAT1 | 376 to 380, 431 to 435, 739 to 743, 923 to 927 |
| | >Os02t0254700-01 | 5 | LOC_Os02g15550 | Spermidine synthase 3 | 213 to 217, 352 to 356, 369 to 373, 519 to 523, 747 to 751 |
| | >Os02t0258900-02 | 4 | LOC_Os02g15870 | Molybdopterin biosynthesis CNX2 protein (Molybdenum cofactor biosynthesis enzyme CNX2) | 73 to 77, 279 to 283, 434 to 438, 973 to 977 |
| | >Os02t0297200-01 | 5 | LOC_Os02g19470 | Protein of unknown function DUF594 family protein | 706 to 710, 742 to 746, 774 to 778, 843 to 847, 848 to 852 |
| | >Os02t0309500-00 | 4 | LOC_Os02g20620 | BTB/POZ-like domain. speckle-type POZ protein | 445 to 449, 524 to 528, 533 to 537, 553 to 557 |
| | >Os02t0326500-01 | 4 | LOC_Os02g22084 | Targeting protein-related | 339 to 343, 360 to 364, 505 to 509, 535 to 539 |
| | >Os02t0329933-00 | 4 | | UDP-glycosyltransferase UGT99C4 | 320 to 324, 369 to 373, 401 to 405, 549 to 553 |

| | | | | | |
|---|---|---|---|---|---|
| | >Os02t0461600-03 | 4 | LOC_Os02g26349 | Protein of unknown function DUF1777 domain. transposon protein, putative, CACTA, En/Spm sub-class | 409 to 413, 610 to 614, 730 to 734, 736 to 740 |
| | >Os02t0491600-00 | 4 | LOC_Os02g29000 | Oxalate oxidase-like protein or germin-like protein (Germin-like 8) (Germin-like 12). Cupin domain | 124 to 128, 303 to 307, 563 to 567, 959 to 963 |
| | >Os02t0512400-01 | 4 | LOC_Os02g30850 | Glutaredoxin. OsGrx_C8 - glutaredoxin subgroup III | 90 to 94, 291 to 295, 523 to 527, 779 to 783 |
| | >Os02t0528900-01 | 4 | LOC_Os02g32690 | PDR-like ABC transporter. pleiotropic drug resistance protein 15 | 301 to 305, 591 to 595, 600 to 604, 619 to 623 |
| | >Os02t0530100-01 | 4 | LOC_Os02g32814 | C4-dicarboxylate transporter/malic acid transport protein. (Os02t0530100-01); Copper chaperone homolog CCH. heavy metal-associated domain | 66 to 70, 326 to 330, 654 to 658, 746 to 750 |
| | >Os02t0538000-01 | 6 | LOC_Os02g33500 | Threonyl-tRNA synthetase. | 17 to 21, 365 to 369, 374 to 378, 394 to 398, 743 to 747,<br>752 to 756 |
| | >Os02t0561900-00 | 5 | LOC_Os02g35440 | E3 ubiquitin-protein ligase EL5. RING-H2 finger protein ATL4O precursor | 276 to 280, 408 to 412, 440 to 444, 582 to 586, 588 to 592 |
| | >Os02t0575700-01 | 4 | LOC_Os02g36590 | OSIGBa0097P08.1 protein. CPuORF19 - conserved peptide uORF | 170 to 174, 382 to 386, 732 to 736, 794 to 798 |
| | >Os02t0577300-00 | 4 | LOC_Os02g36770 | Galactosyltransferase family protein | 478 to 482, 617 to 621, 826 to 830, 929 to 933 |
| | >Os02t0581900-01 | 4 | LOC_Os02g37109 | Pentatricopeptide repeat protein PPR986-12. vegetative cell wall protein gp1 precursor | 321 to 325, 411 to 415, 545 to 549, 823 to 827 |
| | >Os02t0615800-01 | 4 | LOC_Os02g40240 | Protein kinase, core domain | 137 to 141, 189 to 193, 278 to 282, 536 to 540 |
| | >Os02t0627100-01 | 5 | LOC_Os02g41680 | Phenylalanine ammonia-lyase | 3 to 7, 146 to 150, 332 to 336, 361 to 365, 465 to 469 |
| | >Os02t0631500-00 | 4 | LOC_Os02g42040 | Initiation factor eIF-4 gamma, middle; Up-frameshift suppressor 2. MIF4G domain | 109 to 113, 166 to 170, 178 to 182, 778 to 782 |

| | | | | | |
|---|---|---|---|---|---|
| | >Os02t0640500-01 | 4 | LOC_Os02g42780 | Concanavalin A-like lectin/glucanase, subgroup domain. lectin receptor-type protein kinase | 239 to 243, 360 to 364, 679 to 683, 922 to 926 |
| | >Os02t0654000-01 | 4 | LOC_Os02g43710 | Methylglutaconyl-CoA hydratase. enoyl-CoA hydratase/isomerase family protein | 473 to 477, 583 to 587, 690 to 694, 756 to 760 |
| | >Os02t0654300-01 | 4 | LOC_Os02g43740 | Protein kinase KIPK. AGC_PVPK_like_kin82y.6 - ACG kinases include homologs to PKA, PKG and PKC | 75 to 79, 94 to 98, 409 to 413, 984 to 988 |
| | >Os02t0662200-01 | 4 | LOC_Os02g44330 | YbaK/aminoacyl-tRNA synthetase associated region domain. rho guanine nucleotide exchange factor | 33 to 37, 592 to 596, 657 to 661, 891 to 895 |
| | >Os02t0668400-00 | 4 | LOC_Os02g44810 | tRNA pseudouridine synthase family protein | 25 to 29, 63 to 67, 216 to 220, 587 to 591 |
| | >Os02t0673600-01 | 4 | LOC_Os02g45180 | ORMDL family protein. ORM1 | 275 to 279, 564 to 568, 572 to 576, 903 to 907 |
| | >Os02t0684000-00 | 4 | LOC_Os02g45880 | Tetratricopeptide-like helical domain | 24 to 28, 66 to 70, 93 to 97, 916 to 920 |
| | >Os02t0690600-01 | 4 | LOC_Os02g46500 | Zinc finger, RING/FYVE/PHD-type domain | 283 to 287, 314 to 318, 925 to 929, 932 to 936 |
| | >Os02t0693400-01 | 5 | LOC_Os02g46650 | Peptidase C19, ubiquitin carboxyl-terminal hydrolase 2 family protein. ubiquitin carboxyl-terminal hydrolase domain | 196 to 200, 646 to 650, 724 to 728, 770 to 774 |
| | >Os02t0704800-01 | 6 | LOC_Os02g47590 | Ornithine carbamoyl transferase | 207 to 211, 214 to 218, 314 to 318, 628 to 632, 816 to 820, 900 to 904 |
| | >Os02t0708300-01 | 5 | LOC_Os02g47870 | Zinc finger, RING/FYVE/PHD-type domain. anaphase-promoting complex subunit 11 | 209 to 213, 363 to 367, 497 to 501, 528 to 532, 682 to 686 |
| | >Os02t0712700-01 | 4 | LOC_Os02g48210 | Concanavalin A-like lectin/glucanase domain. lectin-like protein kinase | 133 to 137, 158 to 162, 667 to 671, 692 to 696 |
| | >Os02t0714500-01 | 4 | LOC_Os02g48380 | Tetratricopeptide-like helical domain | 329 to 333, 353 to 357, 622 to 626, 773 to 777 |

| | | | | | |
|---|---|---|---|---|---|
| | >Os02t0722500-01 | 4 | LOC_Os02g49070 | Arf GTPase activating protein family protein. ZAC | 211 to 215, 494 to 498, 503 to 507, 522 to 526 |
| | >Os02t0725100-01 | 4 | LOC_Os02g49320 | Thioredoxin fold domain. sucrase-related | 404 to 408, 463 to 467, 501 to 505, 834 to 838 |
| | >Os02t0727300-01 | 4 | LOC_Os02g49520 | Beta-catenin repeat family protein. armadillo/beta-catenin repeat family protein | 298 to 302, 355 to 359, 754 to 758, 760 to 764 |
| | >Os02t0728100-01 | 4 | LOC_Os02g49570 | SPPA; serine-type endopeptidase. OsProtIV1 - Putative Protease IV homologue; Domain 2 is a SPH | 22 to 26, 310 to 314, 786 to 790, 966 to 970 |
| | >Os02t0728100-02 | 4 | LOC_Os02g49570 | SPPA; serine-type endopeptidase. OsProtIV1 - Putative Protease IV homologue; Domain 2 is a SPH | 5 to 9, 293 to 297, 769 to 773, 949 to 953 |
| | >Os02t0730400-02 | 4 | LOC_Os02g49770 | Six-bladed beta-propeller, TolB-like domain. NHL repeat | 542 to 546, 695 to 699, 828 to 832, 902 to 906 |
| | >Os02t0751100-02 | 4 | LOC_Os02g51540 | Peptidase A1 domain. eukaryotic aspartyl protease domain | 49 to 53, 326 to 330, 661 to 665, 794 to 798 |
| | >Os02t0767400-01 | 4 | LOC_Os02g52850 | Serine/threonine protein kinase-related domain. receptor-like protein kinase like protein | 487 to 491, 600 to 604, 606 to 610, 806 to 810 |
| | >Os02t0771100-01 | 4 | LOC_Os02g53140 | COP1 | 63 to 67, 114 to 118, 505 to 509, 525 to 529 |
| | >Os02t0776900-01 | 5 | LOC_Os02g53690 | Transcription activator, Gibberellin (GA) - induced stem elongation growth regulating factor protein | 119 to 123, 432 to 436, 441 to 445, 460 to 464, 735 to 739 |
| | >Os02t0776900-02 | 5 | LOC_Os02g53690 | Transcription activator, Gibberellin (GA) - induced stem elongation growth regulating factor protein | 86 to 90, 399 to 403, 408 to 412, 427 to 431, 702 to 706 |
| | >Os02t0778500-01 | 4 | LOC_Os02g53810 | Chalcone isomerase domain | 294 to 298, 378 to 382, 693 to 697, 822 to 826 |
| | >Os02t0796700-01 | 4 | LOC_Os02g55340 | Protein kinase family protein / WD-40 repeat family protein. WD domain and HEAT domain | 75 to 79, 532 to 536, 540 to 544, 879 to 883 |
| | >Os02t0807100-02 | 5 | LOC_Os02g56320 | Starch synthase V. | 187 to 191, 238 to 242, 250 to 254, 391 |

| | | | | | |
|---|---|---|---|---|---|
| | | | | OsWAK18b | to 395, 471 to 475 |
| | >Os02t0815200-01 | 4 | LOC_Os02g57010 | 29 kDa ribonucleoprotein, chloroplast precursor (RNA-binding protein cp29). RNA recognition motif | 470 to 474, 478 to 482, 489 to 493, 557 to 561 |
| | >Os02t0819400-01 | 4 | LOC_Os02g57400 | Mutant low phytic acid protein 1. 2-phosphoglycerate kinase-related | 251 to 255, 444 to 448, 517 to 521, 582 to 586 |
| | >Os02t0821400-01 | 5 | LOC_Os02g57560 | Protein kinase, core domain. tyrosine protein kinase domain | 191 to 195, 200 to 204, 219 to 223, 508 to 512, 972 to 976 |
| | >Os02t0821400-02 | 5 | LOC_Os02g57560 | Protein kinase, core domain. tyrosine protein kinase domain | 168 to 172, 177 to 181, 196 to 200, 485 to 489, 949 to 953 |
| **TTGACA** | >Os02t0712700-01 | 4 | LOC_Os02g48210 | Concanavalin A-like lectin/ glucanase domain | 133 to 138, 158 to 163, 667 to 672, 692 to 697 |
| | >Os02t0767400-01 | 4 | LOC_Os02g52850 | Serine/threonine protein kinase-related domain. receptor-like protein kinase like protein | 487 to 492, 600 to 605, 606 to 611, 806 to 811 |
| **TTGACT** | >Os02t0115600-02 | 4 | LOC_Os02g02390 | S1, RNA binding domain | 116 to 121, 526 to 531, 577 to 582, 896 to 901 |
| | >Os02t0186800-00 | 4 | LOC_Os02g09390 | Cytochrome P450 family protein | 230 to 235, 411 to 416, 417 to 422, 978 to 983 |
| | >Os02t0297200-01 | 4 | LOC_Os02g19470 | Protein of unknown function DUF594 family protein | 742 to 747, 774 to 779, 843 to 848, 848 to 853 |
| | >Os02t0561900-00 | 4 | LOC_Os02g35440 | E3 ubiquitin-protein ligase EL5. RING-H2 finger protein ATL4O precursor | 408 to 413, 440 to 445, 582 to 587, 588 to 593 |
| | >Os02t0690600-01 | 4 | LOC_Os02g46500 | plant U-box protein 36 | 283 to 288, 314 to 319, 925 to 930, 932 to 937 |
| | >Os02t0704800-01 | 4 | LOC_Os02g47590 | Ornithine carbamoyltransferase | 207 to 212, 214 to 219, 628 to 633, 900 to 905 |
| | >Os02t0708300-01 | 4 | LOC_Os02g47870 | Zinc finger, RING/FYVE/PHD-type domain. anaphase-promoting complex subunit 11 | 363 to 368, 497 to 502, 528 to 533, 682 to 687 |
| **Chromosome 3** | | | | | |
| **Motif** | **ID** | **Frequency** | **Locus ID** | **Protein name** | **Positions in 1k promoter region** |

| | | | | | |
|---|---|---|---|---|---|
| **CTGACC** | >Os03t0812800-00 | 7 | LOC_Os03g59790 | Calcium-binding allergen Ole e 8 | 389 to 394, 412 to 417, 435 to 440, 458 to 463, 500 to 505, 523 to 528, 546 to 551 |
| **TTTGACT** | >Os03t0821100-01 | 5 | LOC_Os03g60620 | Non-cell-autonomous heat shock cognate protein 70. DnaK family protein | 130 to 366, 2 to 68, 211 to 217, 217 to 223, 365 to 371 |
| **TTGACG** | >Os03t0164700-01 | 4 | LOC_Os03g06880 | Carbohydrate/purine kinase domain. kinase, pfkB family | 103 to 108, 119 to 124, 438 to 443, 823 to 828 |
| | >Os03t0365000-00 | 4 | LOC_Os03g24960 | GDSL esterase/lipase protein 47 | 702 to 707, 778 to 783, 806 to 811, 868 to 873 |
| **GTTGACC** | >Os03t0258500-00 | 4 | LOC_Os03g15290 | Autophagy Associated Gene 6C | 438 to 444, 501 to 507, 570 to 576, 791 to 797 |
| **GTTGAC** | >Os03t0258500-00 | 4 | LOC_Os03g15290 | Autophagy Associated gene 6C | 438 to 443, 501 to 506, 570 to 575, 791 to 796 |
| **CTGACT** | >Os03t0428800-01 | 4 | LOC_Os03g31490 | Tetratricopeptide-like helical domain | 116 to 121, 161 to 166, 371 to 376, 488 to 493 |
| | >Os03t0851900-02 | 4 | LOC_Os03g63500 | ATPase, AFG1-like domain | 617 to 622, 624 to 629, 703 to 708, 710 to 715 |
| **GTACGTAC** | >Os03t0416200-01 | 4 | LOC_Os03g30250 | Brittle Culm 1 | 87 to 94, 109 to 116, 113 to 120, 117 to 124 |
| | >Os03t0416200-02 | 4 | LOC_Os03g30250 | Brittle Culm 1 | 57 to 647, 9 to 86, 83 to 90, 87 to 94 |
| **TGACC** | >Os03t0113100-01 | 5 | LOC_Os03g02200 | Thymidine kinase | 161 to 165, 332 to 336, 511 to 515, 699 to 703, 883 to 887 |
| | >Os03t0113100-02 | 5 | LOC_Os03g02200 | Thymidine kinase | 148 to 152, 319 to 323, 498 to 502, 686 to 690, 870 to 874 |
| | >Os03t0130500-02 | 4 | LOC_Os03g03830 | EF hand domain | 515 to 519, 629 to 633, 695 to 699, 828 to 832 |
| | >Os03t0131000-01 | 4 | LOC_Os03g03890 | H0409D10.5 protein | 241 to 245, 269 to 273, 667 to 671, 777 to 781 |
| | >Os03t0146100-05 | 4 | LOC_Os03g05290 | Tonoplast intrinsic protein (Tonoplast water channel) aquaporin protein | 329 to 333, 635 to 639, 753 to 757, 876 to 880 |
| | >Os03t0177900-03 | 6 | LOC_Os03g08050 | EF-1 alpha. elongation factor Tu, | 181 to 185, 318 to 322, 339 to 343, 478 to 482, 612 to 616, 837 to 841 |
| | >Os03t0244700-01 | 4 | LOC_Os03g14090 | Armadillo-like helical domain. armadillo/beta-catenin repeat family protein | 78 to 82, 228 to 232, 286 to 290, 782 to 786 |
| | >Os03t0258500-00 | 6 | LOC_Os03g15290 | Autopagy | 33 to 37, 275 to 279, 440 to 444, 503 to 507, 572 to 576, 793 to 797 |

| | | | | | |
|---|---|---|---|---|---|
| | >Os03t0258900-01 | 4 | LOC_Os03g15320 | Galactose oxidase, beta-propeller domain. glyoxal oxidase-related | 666 to 670, 780 to 784, 792 to 796, 841 to 845 |
| | >Os03t0296800-01 | 4 | LOC_Os03g18550 | Mitochondrial carrier protein domain | 386 to 390, 716 to 720, 724 to 728, 966 to 970 |
| | >Os03t0296800-02 | 4 | LOC_Os03g18550 | Mitochondrial carrier protein domain | 382 to 386, 712 to 716, 720 to 724, 962 to 966 |
| | >Os03t0319100-01 | 4 | LOC_Os03g20340 | SNF1-related protein kinase regulatory subunit beta-1 | 526 to 530, 664 to 668, 741 to 745, 874 to 878 |
| | >Os03t0350100-01 | 4 | LOC_Os03g22730 | SAR DNA-binding protein-2 | 84 to 88, 221 to 225, 602 to 606, 693 to 697 |
| | >Os03t0350100-02 | 4 | LOC_Os03g22730 | SAR DNA-binding protein-2, nucleolar protein NOP5-1 | 81 to 85, 218 to 222, 599 to 603, 690 to 694 |
| | >Os03t0371300-00 | 5 | LOC_Os03g25500 | Cytochrome P450 CYP709E4. cytochrome P450 72A1 | 235 to 239, 262 to 266, 336 to 340, 365 to 369, 916 to 920 |
| | >Os03t0397700-02 | 4 | LOC_Os03g27990 | Protein kinase, catalytic domain domain. STRUBBELIG-RECEPTOR FAMILY 7 precursor | 150 to 154, 467 to 471, 722 to 726, 888 to 892 |
| | >Os03t0401300-03 | 5 | LOC_Os03g28330 | Sucrose synthase 2 sucrose synthase | 14 to 18, 325 to 329, 603 to 607, 693 to 697, 750 to 754 |
| | >Os03t0577200-01 | 5 | LOC_Os03g38020 | Mps one binder kinase activator-like 1A mps one binder kinase activator-like 1A | 5 to 9, 217 to 221, 820 to 824, 850 to 854, 855 to 859 |
| | >Os03t0596400-01 | 5 | LOC_Os03g39910 | XH domain | 263 to 267, 424 to 428, 480 to 484, 889 to 893, 923 to 927 |
| | >Os03t0647400-01 | 4 | LOC_Os03g44520 | GCK domain | 60 to 64, 89 to 93, 188 to 192, 844 to 848 |
| | >Os03t0694500-01 | 4 | LOC_Os03g48810 | Permease 1. nucleobase-ascorbate transporter | 233 to 237, 338 to 342, 627 to 631, 830 to 834 |
| | >Os03t0712400-02 | 4 | LOC_Os03g50450 | A typical receptor-like kinase MARK. inactive receptor kinase At2g26730 precursor | 517 to 521, 757 to 761, 763 to 767, 782 to 786 |
| | >Os03t0734300-01 | 4 | LOC_Os03g52390 | Proteinase inhibitor type II CEVI57 precursor. PIII1 - Proteinase inhibitor II family protein | 550 to 554, 788 to 792, 794 to 798, 957 to 961 |
| | >Os03t0744600-01 | 4 | LOC_Os03g53270 | Ripening-associated protein stem-specific protein TSJT1 | 253 to 257, 399 to 403, 868 to 872, 904 to 908 |

| | | | | | |
|---|---|---|---|---|---|
| | >Os03t0745600-01 | 4 | LOC_Os03g53400 | UPF0005 domain. transmembrane BAX inhibitor motif | 202 to 206, 237 to 241, 295 to 299, 563 to 567 |
| | >Os03t0812800-00 | 7 | LOC_Os03g59790 | Calcium-binding allergen Ole e 8. EF hand family protein | 390 to 394, 413 to 417, 436 to 440, 459 to 463, 501 to 505, 524 to 528, 547 to 551 |
| | >Os03t0822100-01 | 9 | LOC_Os03g60730 | Transposase | 195 to 199, 248 to 252, 272 to 276, 462 to 466, 524 to 528, 599 to 603, 700 to 704, 859 to 863, 992 to 996 |
| | >Os03t0825800-01 | 4 | LOC_Os03g61060 | Protein kinase, core domain | 19 to 23, 349 to 353, 552 to 556, 897 to 901 |
| **TGACT** | >Os03t0103000-01 | 4 | LOC_Os03g01290 | Nitrate transporter. peptide transporter PTR2 | 323 to 327, 769 to 773, 777 to 781, 781 to 785 |
| | >Os03t0124300-01 | 4 | LOC_Os03g03290 | ATP binding protein. receptor-like protein kinase | 472 to 476, 610 to 614, 647 to 651, 661 to 665 |
| | >Os03t0124300-02 | 4 | LOC_Os03g03290 | ATP binding protein. receptor-like protein kinase | 429 to 433, 567 to 571, 604 to 608, 618 to 622 |
| | >Os03t0133400-01 | 4 | LOC_Os03g04110 | Peptidoglycan-binding Lysin subgroup domain | 154 to 158, 703 to 707, 747 to 751, 879 to 883 |
| | >Os03t0150600-01 | 4 | LOC_Os03g05620 | Pi transporter, Pi homeostasis inorganic phosphate transporter | 310 to 314, 459 to 463, 544 to 548, 922 to 926 |
| | >Os03t0150600-02 | 4 | LOC_Os03g05620 | Pi transporter, Pi homeostasis inorganic phosphate transporter | 311 to 315, 460 to 464, 545 to 549, 923 to 927 |
| | >Os03t0162650-01 | 4 | LOC_Os03g06700 | fatty acid elongase. 3-ketoacyl-CoA synthase | 220 to 224, 234 to 238, 546 to 550, 781 to 785 |
| | >Os03t0173500-01 | 4 | LOC_Os03g07739 | Kinase associated protein phosphatase | 348 to 352, 355 to 359, 403 to 407, 410 to 414 |
| | >Os03t0173500-02 | 4 | LOC_Os03g07739 | Kinase associated protein phosphatase | 344 to 348, 351 to 355, 399 to 403, 406 to 410 |
| | >Os03t0184000-00 | 4 | LOC_Os03g08570 | Phytoene desaturase amine oxidase, flavin | 57 to 61, 271 to 275, 333 to 337, 879 to 883 |
| | >Os03t0215000-03 | 4 | LOC_Os03g11590 | integral membrane family protein. | 33 to 37, 110 to 114, 845 to 849, 955 to 959 |
| | >Os03t0223900-01 | 4 | LOC_Os03g12320 | UDP-3-O-acyl N-acetylglucosamine deacetylase domain | 314 to 318, 695 to 699, 740 to 744, 859 to 863 |
| | >Os03t0279816-00 | 4 | LOC_Os03g17164 | Kinesin-related protein | 141 to 145, 448 to 452, 477 to 481, 895 to 899 |

| | | | | | |
|---|---|---|---|---|---|
| | >Os03t0329500-01 | 4 | LOC_Os03g21210 | Endo-1,4-beta-glucanase endoglucanase | 45 to 49, 496 to 500, 724 to 728, 728 to 732 |
| | >Os03t0352500-01 | 4 | LOC_Os03g22900 | AGR379Wp. | 117 to 121, 241 to 245, 511 to 515, 941 to 945 |
| | >Os03t0353900-01 | 5 | LOC_Os03g23050 | Argonaute and Dicer protein, PAZ domain | 93 to 97, 216 to 220, 228 to 232, 509 to 513, 759 to 763 |
| | >Os03t0368300-01 | 4 | LOC_Os03g25300 | Peroxidase 1. peroxidase precursor | 460 to 464, 505 to 509, 570 to 574, 579 to 583 |
| | >Os03t0375717-00 | 5 | LOC_Os03g25890 | Retrotransposon protein | 219 to 223, 277 to 281, 291 to 295, 699 to 703, 981 to 985 |
| | >Os03t0375966-00 | 4 | LOC_Os03g25920 | Amino acid permease family protein. | 178 to 182, 524 to 528, 847 to 851, 855 to 859 |
| | >Os03t0389100-01 | 4 | LOC_Os03g27190 | Peptidase C14, caspase catalytic domain. ICE-like protease p20 domain | 578 to 582, 597 to 601, 648 to 652, 979 to 983 |
| | >Os03t0401300-02 | 5 | LOC_Os03g28330 | Sucrose synthase 2 sucrose synthase | 481 to 485, 545 to 549, 631 to 635, 805 to 809, 829 to 833 |
| | >Os03t0401300-03 | 4 | LOC_Os03g28330 | Sucrose synthase 2 sucrose synthase | 71 to 75, 95 to 99, 528 to 532, 620 to 624 |
| | >Os03t0425200-01 | 4 | LOC_Os03g31170 | Nosine/uridine-preferring nucleoside hydrolase domain. inosine-uridine preferring nucleoside hydrolase family protein | 455 to 459, 653 to 657, 737 to 741, 937 to 941 |
| | >Os03t0425200-02 | 4 | LOC_Os03g31170 | Inosine/uridine-preferring nucleoside hydrolase domain. inosine-uridine preferring nucleoside hydrolase family protein | 419 to 423, 617 to 621, 701 to 705, 901 to 905 |
| | >Os03t0426300-00 | 4 | LOC_Os03g31260 | Cysteine-rich receptor-like protein kinase 28 precursor | 413 to 417, 458 to 462, 872 to 876, 893 to 897 |
| | >Os03t0428800-01 | 4 | LOC_Os03g31490 | Tetratricopeptide-like helical domain | 117 to 121, 162 to 166, 372 to 376, 489 to 493 |
| | >Os03t0565200-01 | 4 | LOC_Os03g36750 | Haloacid dehalogenase-like hydrolase domain. cbbY | 237 to 241, 337 to 341, 370 to 374, 515 to 519 |
| | >Os03t0576600-00 | 4 | LOC_Os03g37960 | Acyl-CoA-binding protein (ACBP) | 31 to 35, 82 to 86, 265 to 269, 338 to 342 |
| | >Os03t0583900-03 | 4 | LOC_Os03g38740 | DEAD-like helicase, N-terminal domain | 52 to 56, 344 to 348, 416 to 420, 561 to 565 |
| | >Os03t0599400-01 | 4 | LOC_Os03g40250 | NB-ARC domain. Leucine Rich Repeat family protein | 111 to 115, 127 to 131, 242 to 246, 670 to 674 |

| | | | | | |
|---|---|---|---|---|---|
| | >Os03t0645100-02 | 4 | LOC_Os03g44300 | Pyruvate dehydrogenase E1 component subunit beta. transketolase, | 36 to 40, 253 to 257, 341 to 345, 777 to 781 |
| | >Os03t0675000-01 | 4 | LOC_Os03g47169 | Peptidase family C50 | 6 to 10, 121 to 125, 137 to 141, 161 to 165 |
| | >Os03t0697200-00 | 4 | LOC_Os03g49050 | Carboxylyase. possible lysine decarboxylase domain | 6 to 10, 121 to 125, 137 to 141, 161 to 165 |
| | >Os03t0705800-00 | 4 | LOC_Os03g49830 | Protein of unknown function DUF3049 domain | 103 to 107, 689 to 693, 873 to 877, 917 to 921 |
| | >Os03t0712700-03 | 4 | LOC_Os03g50480 | Phosphoglucomutase, cytoplasmic 2 | 161 to 165, 524 to 528, 675 to 679, 863 to 867 |
| | >Os03t0722400-00 | 4 | LOC_Os03g51230 | Chromatin complex subunit A101 | 86 to 90, 580 to 584, 709 to 713, 808 to 812 |
| | >Os03t0737000-01 | 4 | LOC_Os03g52690 | Cystathionine beta-synthase, core domain | 96 to 100, 559 to 563, 850 to 854, 986 to 990 |
| | >Os03t0737000-02 | 4 | LOC_Os03g52690 | Cystathionine beta-synthase, core domain | 39 to 43, 502 to 506, 793 to 797, 929 to 933 |
| | >Os03t0737000-03 | 4 | LOC_Os03g52690 | Cystathionine beta-synthase, core domain | 37 to 41, 500 to 504, 791 to 795, 927 to 931 |
| | >Os03t0741400-01 | 4 | LOC_Os03g53050 | SUSIBA2. WRKY121 | 51 to 55, 129 to 133, 279 to 283, 347 to 351 |
| | >Os03t0743500-01 | 5 | LOC_Os03g53200 | Calcium-binding EF-hand domain. OsCML4 - Calmodulin-related calcium sensor protein | 147 to 151, 222 to 226, 248 to 252, 312 to 316, 904 to 908 |
| | >Os03t0743500-02 | 6 | LOC_Os03g53200 | Calcium-binding EF-hand domain. OsCML4 - Calmodulin-related calcium sensor protein | 56 to 60, 131 to 135, 157 to 161, 221 to 225, 813 to 817, 985 to 989 |
| | >Os03t0767500-01 | 4 | LOC_Os03g55820 | Thioredoxin domain 2 | 430 to 434, 590 to 594, 944 to 948, 967 to 971 |
| | >Os03t0788000-00 | 5 | LOC_Os03g57400 | OSIGBa0116M22.8 protein. | 45 to 49, 71 to 75, 655 to 659, 714 to 718, 805 to 809 |
| | >Os03t0804900-00 | 4 | LOC_Os03g59030 | UDP-glucuronosyl/UDP-glucosyltransferase family protein. UDP-rhamnose rhamnosyltransferase | 83 to 87, 228 to 232, 345 to 349, 594 to 598 |
| | >Os03t0808200-00 | 4 | LOC_Os03g59350 | UDP-glucuronosyl/UDP-glucosyltransferase family protein. anthocyanin 3-O- | 90 to 94, 122 to 126, 270 to 274, 343 to 347 |

| | | | | | |
|---|---|---|---|---|---|
| | | | | beta-glucosyltransferase | |
| | >Os03t0812400-01 | 4 | LOC_Os03g59770 | EF-Hand type domain. EF hand family protein | 65 to 69, 146 to 150, 402 to 406, 407 to 411 |
| | >Os03t0818200-01 | 4 | LOC_Os03g60380 | NAD(P)-binding domain. cinnamoyl CoA reductase | 293 to 297, 518 to 522, 689 to 693, 860 to 864 |
| | >Os03t0821100-01 | 5 | LOC_Os03g60620 | Non-cell-autonomous heat shock cognate protein 70. DnaK family protein | 32 to 36, 64 to 68, 213 to 217, 219 to 223, 367 to 371 |
| | >Os03t0823100-01 | 4 | LOC_Os03g60820 | Major facilitator superfamily protein. transporter, major facilitator superfamily domain | 175 to 179, 801 to 805, 938 to 942, 946 to 950 |
| | >Os03t0828800-01 | 4 | LOC_Os03g61310 | Serine/threonine protein kinase-related domain. receptor-like protein kinase | 404 to 408, 696 to 700, 738 to 742, 785 to 789 |
| | >Os03t0851900-02 | 6 | LOC_Os03g63500 | ATPase, AFG1-like domain. | 79 to 83, 618 to 622, 625 to 629, 704 to 708, 711 to 715, 852 to 856 |
| | >Os03t0858600-01 | 4 | LOC_Os03g64130 | Protein of unknown function DUF668 family protein. | 41 to 45, 45 to 49, 86 to 90, 831 to 835 |
| **TTGAC** | >Os03t0101000-01 | 5 | LOC_Os03g01140 | acyl-activating enzyme 11 | 194 to 198, 404 to 408, 438 to 442, 450 to 454, |
| | >Os03t0107300-01 | 4 | LOC_Os03g01700 | Anion transporter, Silicon efflux transporter, Arsenic species (As) uptake | 558 to 562, 645 to 649, 654 to 658, 673 to 677 |
| | >Os03t0111100-01 | 4 | LOC_Os03g02030 | Dihydrofolate synthetase /folylpolyglutamate synthetase. | 103 to 107, 231 to 235 , 362 to 366, 730 to 734 |
| | >Os03t0124300-01 | 4 | LOC_Os03g03290 | ATP binding protein. receptor-like protein kinase At3g46290 precursor | 45 to 49, 471 to 475, 646 to 650, 660 to 664 |
| | >Os03t0124300-02 | 4 | LOC_Os03g03290 | ATP binding protein. receptor-like protein kinase At3g46290 precursor | 2 to 6, 428 to 432, 603 to 607, 617 to 621 |
| | >Os03t0127600-01 | 4 | LOC_Os03g03560 | Forkhead-associated domain | 281 to 285, 633 to 637, 638 to 642, 862 to 866 |
| | >Os03t0127700-01 | 4 | LOC_Os03g03570 | Serine/threonine protein kinase-related domain. leucine-rich repeat transmembrane protein kinase | 8 to 12, 569 to 573, 601 to 605, 818 to 822 |

| | | | | | |
|---|---|---|---|---|---|
| | >Os03t0127700-02 | 4 | LOC_Os03g03570 | Serine/threonine protein kinase-related domain. leucine-rich repeat transmembrane protein kinase | 5 to 9, 566 to 570, 598 to 602, 815 to 819 |
| | >Os03t0128500-01 | 5 | LOC_Os03g03650 | DNA polymerase delta small subunit POLD2 - Putative DNA polymerase delta complex subunit | 84 to 88, 545 to 549, 617 to 621, 658 to 662,<br>686 to 690 |
| | >Os03t0131000-01 | 5 | LOC_Os03g03890 | H0409D10.5 protein. protein kinase family protein | 62 to 66, 110 to 114, 268 to 272, 666 to 670,<br>742 to 746 |
| | >Os03t0150500-01 | 6 | LOC_Os03g05610 | Phosphate transporter 6. inorganic | 215 to 219, 223 to 227, 298 to 302, 306 to 310,<br>411 to 415, 416 to 420 |
| | >Os03t0157800-01 | 4 | LOC_Os03g06190 | 3-5 exonuclease family protein | 136 to 140, 202 to 206, 412 to 416, 605 to 609 |
| | >Os03t0157800-02 | 4 | LOC_Os03g06190 | 3-5 exonuclease family protein | 75 to 79, 141 to 145, 351 to 355, 544 to 548 |
| | >Os03t0159200-01 | 7 | LOC_Os03g06340 | Region of unknown function XS domain | 383 to 387, 401 to 405, 575 to 579, 598 to 602, 615 to 619, 651 to 655, 853 to 857 |
| | >Os03t0164700-01 | 4 | LOC_Os03g06880 | Carbohydrate/purine kinase domain. kinase, pfkB family | 103 to 107, 119 to 123, 438 to 442, 823 to 827 |
| | >Os03t0165375-00 | 4 | | Chlorophyll a-b binding protein 2, chloroplastic | 716 to 720, 746 to 750, 805 to 809, 890 to 894 |
| | >Os03t0170800-00 | 5 | LOC_Os03g07470 | Glutaredoxin | 235 to 239, 264 to 268, 509 to 513, 791 to 795, 921 to 925 |
| | >Os03t0184550-01 | 5 | LOC_Os03g08624 | Dihydroflavonol-4-reductase. | 14 to 18, 299 to 303, 353 to 357, 648 to 652, 877 to 881 |
| | >Os03t0205400-00 | 4 | LOC_Os03g10780 | XPG I-region family protein. flap endonuclease | 336 to 340, 345 to 349, 364 to 368, 877 to 881 |
| | >Os03t0213100-01 | 4 | LOC_Os03g11440 | Sec61p. protein transport protein Sec61 subunit alpha | 252 to 256, 365 to 369, 458 to 462, 501 to 505 |
| | >Os03t0215000-03 | 4 | LOC_Os03g11590 | Integral membrane family protein. | 32 to 36, 109 to 113, 844 to 848, 954 to 958 |
| | >Os03t0215700-01 | 4 | LOC_Os03g11650 | Myosin II heavy chain-like family protein. ATMAP70 protein | 280 to 284, 389 to 393, 434 to 438, 592 to 596 |

| | | | | | |
|---|---|---|---|---|---|
| | >Os03t0227400-01 | 4 | LOC_Os03g12620 | glucan endo-1,3-beta-glucosidase 7. glycosyl hydrolases family 17 | 437 to 441, 472 to 476, 757 to 761, 832 to 836 |
| | >Os03t0235700-01 | 4 | LOC_Os03g13250 | Peptide transporter 1. peptide transporter PTR2 | 82 to 86, 389 to 393, 727 to 731, 746 to 750 |
| | >Os03t0236200-01 | 5 | LOC_Os03g13300 | Glutamate decarboxylase isozyme 3. | 243 to 247, 352 to 356, 499 to 503, 667 to 671, 817 to 821 |
| | >Os03t0258500-00 | 4 | LOC_Os03g15290 | Beclin 1 protein. beclin-1 | 439 to 443, 502 to 506, 571 to 575, 792 to 796 |
| | >Os03t0261900-01 | 4 | LOC_Os03g15540 | HEAT repeat family protein | 370 to 374, 608 to 612, 634 to 638, 906 to 910 |
| | >Os03t0261900-02 | 5 | LOC_Os03g15540 | HEAT repeat family protein | 86 to 90, 112 to 116, 384 to 388, 680 to 684, 898 to 902 |
| | >Os03t0264400-01 | 5 | LOC_Os03g15780 | Anthranilate synthase alpha 2 subunit. anthranilate synthase component I-1, chloroplast precursor | 25 to 29, 80 to 84, 129 to 133, 217 to 221, 991 to 995 |
| | >Os03t0268200-05 | 4 | LOC_Os03g16130 | Serine/threonine protein kinase domain. CAMK_CAMK_like_ ULKh_APGy.2 - CAMK includes calcium/calmodulin depedent protein kinases | 91 to 95, 658 to 662, 957 to 961, 978 to 982 |
| | >Os03t0284100-02 | 4 | LOC_Os03g17570 | Two-component response regulator-like PRR73. response regulator receiver domain | 359 to 363, 738 to 742, 749 to 753, 921 to 925 |
| | >Os03t0310500-01 | 4 | LOC_Os03g19690 | CAP, conserved site domain | 548 to 552, 635 to 639, 668 to 672, 892 to 896 |
| | >Os03t0336600-02 | 4 | LOC_Os03g21840 | Reticulon family protein. reticulon domain | 436 to 440, 609 to 613, 751 to 755, 792 to 796 |
| | >Os03t0336700-02 | 5 | LOC_Os03g21850 | PAP/25A core domain. nucleotidyltransferase | 21 to 25, 279 to 283, 348 to 352, 441 to 445, 539 to 543 |
| | >Os03t0346800-00 | 4 | LOC_Os03g22550 | metal tolerance protein. cation efflux family protein | 360 to 364, 687 to 691, 760 to 764, 800 to 804 |
| | >Os03t0353900-01 | 6 | LOC_Os03g23050 | Argonaute and Dicer protein, PAZ domain | 92 to 96, 215 to 219, 227 to 231, 758 to 762,<br>959 to 963, 981 to 985 |
| | >Os03t0356596-01 | 4 | LOC_Os03g24170 | Phosphatidylinositol-4-phosphate 5-kinase-like protein | 23 to 27, 184 to 188, 521 to 525, 540 to 544 |

| | | | | | |
|---|---|---|---|---|---|
| | >Os03t0365000-00 | 6 | LOC_Os03g24960 | GDSL-like Lipase/Acylhydrolase family protein. GDSL-like lipase/acylhydrolase | 271 to 275, 647 to 651, 702 to 706, 778 to 782,<br>806 to 810, 868 to 872 |
| | >Os03t0370800-00 | 4 | LOC_Os03g25470 | Ctr copper transporter family protein. | 470 to 474, 637 to 641, 646 to 650, 665 to 669 |
| | >Os03t0386800-01 | 4 | LOC_Os03g26930 | Peptidase S10, serine carboxypeptidase family protein. OsSCP13 - Putative Serine Carboxypeptidase homologue | 557 to 561, 752 to 756, 790 to 794, 910 to 914 |
| | >Os03t0401300-03 | 5 | LOC_Os03g28330 | Sucrose synthase 2 (EC 2.4.1.13) | 324 to 328, 447 to 451, 602 to 606, 619 to 623,<br>749 to 753 |
| | >Os03t0429800-01 | 5 | LOC_Os03g31550 | Xanthine dehydrogenase 1 aldehyde oxidase | 44 to 48, 402 to 406, 617 to 621, 708 to 712,<br>732 to 736 |
| | >Os03t0437100-01 | 4 | LOC_Os03g32220 | Zinc finger, C2H2-type domain. ZOS3-11 | 70 to 74, 445 to 449, 550 to 554, 894 to 898 |
| | >Os03t0565200-01 | 4 | LOC_Os03g36750 | Haloacid dehalogenase-like hydrolase domain. cbbY, | 220 to 224, 236 to 240, 336 to 340, 369 to 373 |
| | >Os03t0568500-01 | 4 | LOC_Os03g37090 | Uncharacterised protein family UPF0136, Transmembrane domain | 29 to 33, 151 to 155, 492 to 496, 519 to 523 |
| | >Os03t0577500-02 | 4 | LOC_Os03g38050 | Avr9 elicitor response-like protein. galactosyltransferase | 275 to 279, 280 to 284, 719 to 723, 787 to 791 |
| | >Os03t0581400-00 | 4 | LOC_Os03g38470 | Anther-specific proline-rich protein APG. GDSL-like lipase/acylhydrolase | 215 to 219, 224 to 228, 243 to 247, 452 to 456 |
| | >Os03t0607500-01 | 4 | LOC_Os03g41080 | Seed maturation protein PM23 | 153 to 157, 160 to 164, 236 to 240, 243 to 247 |
| | >Os03t0650800-00 | 4 | LOC_Os03g44800 | Protein of unknown function DUF593 family protein. | 114 to 118, 122 to 126, 428 to 432, 452 to 456 |
| | >Os03t0659400-02 | 6 | LOC_Os03g45730 | DWNN domain domain. DNA-binding protein-like | 20 to 24, 234 to 238, 336 to 340, 398 to 402, 561 to 565, 573 to 577 |
| | >Os03t0662000-01 | 4 | LOC_Os03g45990 | Dihydrolipoyl dehydrogenase, mitochondrial precursor | 179 to 183, 322 to 326, 424 to 428, 760 to 764 |
| | >Os03t0664700-01 | 4 | LOC_Os03g46190 | RNA polymerase II accessory factor, Cdc73 domain. parafibromin | 487 to 491, 787 to 791, 858 to 862, 863 to 867 |

| | | | | | |
|---|---|---|---|---|---|
| | >Os03t0713400-01 | 4 | LOC_Os03g50540 | NADH-ubiquinone oxidoreductase 75 kDa subunit, mitochondrial precursor 2Fe-2S iron-sulfur cluster binding domain | 96 to 100, 139 to 143, 668 to 672, 775 to 779 |
| | >Os03t0737000-01 | 4 | LOC_Os03g52690 | Cystathionine beta-synthase, core domain | 95 to 99, 558 to 562, 659 to 663, 849 to 853 |
| | >Os03t0737000-02 | 4 | LOC_Os03g52690 | Cystathionine beta-synthase, core domain | 38 to 42, 501 to 505, 602 to 606, 792 to 796 |
| | >Os03t0737000-03 | 4 | LOC_Os03g52690 | Cystathionine beta-synthase, core domain | 36 to 40, 499 to 503, 600 to 604, 790 to 794 |
| | >Os03t0746900-01 | 4 | LOC_Os03g53540 | Protein of unknown function DUF1677, plant family protein | 136 to 140, 410 to 414, 528 to 532, 559 to 563 |
| | >Os03t0758000-01 | 4 | LOC_Os03g55080 | WRKY3 | 305 to 309, 427 to 431, 436 to 440, 455 to 459 |
| | >Os03t0788000-00 | 5 | LOC_Os03g57400 | OSIGBa0116M22.8 protein | 44 to 48, 70 to 74, 200 to 204, 713 to 717, 804 to 808 |
| | >Os03t0788100-01 | 4 | LOC_Os03g57410 | RING-H2 finger protein ATL1P RING-H2 finger protein ATL5D, | 338 to 342, 722 to 726, 875 to 879, 954 to 958 |
| | >Os03t0788800-01 | 4 | LOC_Os03g57500 | Zinc finger, RING/FYVE/PHD-type domain. zinc finger, C3HC4 type domain | 84 to 88, 678 to 682, 706 to 710, 753 to 757 |
| | >Os03t0790900-01 | 4 | LOC_Os03g57690 | Aldehyde oxidase-2. aldehyde oxidase | 5 to 9, 407 to 411, 574 to 578, 931 to 935 |
| | >Os03t0792400-02 | 4 | LOC_Os03g57840 | mRNA, clone: RTFL01-01-G24. peptidase M50, mammalian sterol-regulatory element binding protein | 72 to 76, 311 to 315, 522 to 526, 733 to 737 |
| | >Os03t0800100-01 | 4 | LOC_Os03g58590 | Cactin, domain | 234 to 238, 242 to 246, 625 to 629, 659 to 663 |
| | >Os03t0804900-00 | 4 | LOC_Os03g59030 | UDP-glucuronosyl/UDP-glucosyltransferase family protein. UDP-rhamnose rhamnosyltransferase, | 82 to 86, 227 to 231, 344 to 348, 558 to 562 |
| | >Os03t0808100-01 | 4 | LOC_Os03g59340 | Cellulose synthase-5. CESA2 - cellulose synthase | 64 to 68, 96 to 100, 244 to 248, 250 to 254 |
| | >Os03t0808200-00 | 4 | LOC_Os03g59350 | UDP-glucuronosyl/UDP-glucosyltransferase family protein. anthocyanin 3-O- | 89 to 93, 121 to 125, 269 to 273, 342 to 346 |

| | | | | | |
|---|---|---|---|---|---|
| | | | | beta-glucosyltransferase | |
| | >Os03t0812400-01 | 4 | LOC_Os03g59770 | EF-Hand type domain | 21 to 25, 406 to 410, 421 to 425, 803 to 807 |
| | >Os03t0821100-01 | 7 | LOC_Os03g60620 | Non-cell-autonomous heat shock cognate protein 70. DnaK family protein | 31 to 35, 63 to 67, 212 to 216, 218 to 222 , 366 to 370, 753 to 757, 853 to 857 |
| | >Os03t0822100-01 | 6 | LOC_Os03g60730 | Transposase | 194 to 198, 271 to 275, 461 to 465, 598 to 602,<br>678 to 682, 991 to 995 |
| | >Os03t0822100-02 | 4 | LOC_Os03g60730 | Transposase | 106 to 110, 155 to 159, 430 to 434, 494 to 498 |
| | >Os03t0826300-00 | 4 | LOC_Os03g60730 | GDP-mannose transporter | 96 to 100, 148 to 152, 958 to 962,<br>969 to 973 |
| | >Os03t0828800-01 | 4 | LOC_Os03g61310 | Serine/threonine protein kinase-related domain. receptor-like protein kinase | 695 to 699, 737 to 741, 775 to 779, 784 to 788 |
| | >Os03t0833300-01 | 4 | LOC_Os03g61760 | Squamosa promoter-binding-like protein 6. OsSPL6 - SBP-box gene family member | 75 to 79, 266 to 270, 663 to 667, 695 to 699 |
| | >Os03t0837000-01 | 4 | LOC_Os03g62080 | Dcp1-like decapping domain. mRNA-decapping enzyme | 102 to 106, 253 to 257, 447 to 451, 605 to 609 |
| | >Os03t0837100-02 | 4 | LOC_Os03g62090 | Cellulose synthase-6. CESA5 - cellulose synthase | 109 to 113, 726 to 730, 734 to 738, 829 to 833 |
| | >Os03t0839200-01 | 4 | LOC_Os03g62270 | Multi antimicrobial extrusion protein MatE family protein. MATE efflux family protein | 200 to 204, 223 to 227, 263 to 267, 468 to 472 |
| | >Os03t0844600-01 | 4 | LOC_Os03g62740 | Esterase, SGNH hydrolase-type domain. GDSL-like lipase/ acylhydrolase | 99 to 103, 164 to 168, 175 to 179, 571 to 575 |
| | >Os03t0848300-01 | 5 | LOC_Os03g63090 | Actin-related protein 2/3 complex subunit 4 actin-related protein 2/3 complex subunit 4 | 64 to 68, 159 to 163, 641 to 645, 650 to 654, 669 to 673 |
| | >Os03t0850900-01 | 4 | LOC_Os03g63390 | Chemocyanin precursor (Basic blue protein) (Plantacyanin). plastocyanin-like domain | 240 to 244, 291 to 295, 502 to 506, 565 to 569 |

| | | | | | |
|---|---|---|---|---|---|
| **TTGACA** | >Os03t0101000-01 | 4 | LOC_Os03g01140 | AMP binding protein. acyl-activating enzyme 11 | 404 to 409, 438 to 443, 450 to 455, 498 to 503 |
| | >Os03t0150500-01 | 4 | LOC_Os03g05610 | Probable inorganic phosphate transporter 1-12 | 215 to 220, 223 to 228, 298 to 303, 306 to 311 |
| | >Os03t0159200-01 | 6 | LOC_Os03g06340 | Region of unknown function XS domain | 383 to 388, 401 to 406, 575 to 580, 598 to 603, 615 to 620, 651 to 656 |
| | >Os03t0170800-00 | 4 | LOC_Os03g07470 | GRX-like protein 5, glutaredoxin-like protein 5 | 235 to 240, 264 to 269, 791 to 796, 921 to 926 |
| | >Os03t0604600-01 | 4 | LOC_Os03g40780 | Transport protein-related | 16 to 21, 760to765, 834 to 839, 890 to 895 |
| **TTGACC** | >Os03t0258500-00 | 4 | LOC_Os03g15290 | Beclin 1 protein | 439 to 444, 502 to 507, 571 to 576, 792 to 797 |
| | >Os03t0822100-01 | 5 | LOC_Os03g60730 | Transposase | 194 to 199, 271 to 276, 461 to 466, 598 to 603, 991 to 996 |
| **TTGACT** | >Os03t0215000-03 | 4 | LOC_Os03g11590 | integral membrane family protein | 32 to 37, 109 to 114, 844 to 849, 954 to 959 |
| | >Os03t0353900-01 | 4 | LOC_Os03g23050 | Argonaute and Dicer protein, PAZ domain | 92 to 97, 215 to 220, 227 to 232, 758 to 763 |
| | >Os03t0788000-00 | 4 | LOC_Os03g57400 | OSIGBa0116M22.8 protein | 44 to 49, 70to75, 713 to 718, 804 to 809 |
| | >Os03t0808200-00 | 4 | LOC_Os03g59350 | UDP-glucuronosyl/UDP-glucosyltransferase family protein. anthocyanin 3-O-beta-glucosyltransferase | 89 to 94, 121 to 126, 269 to 274, 342 to 347 |
| | >Os03t0821100-01 | 5 | LOC_Os03g60620 | Non-cell-autonomous heat shock cognate protein 70. DnaK family protein | 31 to 36, 63 to 68, 212 to 217, 218 to 223, 366 to 371 |
| **Chromosome 4** | | | | | |
| **Motif** | **ID** | **Frequency** | **Locus ID** | **Protein name** | **Positions in 1k promoter region** |
| **CTGACC** | >Os04t0498600-02 | 4 | LOC_Os04g42090 | cDNA clone:J013002M21 | 630 to 635, 725 to 730, 761 to 766, 845 to 850 |
| **TTGACG** | >Os04t0508900-00 | 4 | LOC_Os04g42990 | Suppressor of stem-loop protein 1 | 470 to 475, 514 to 519, 684 to 689, 963 to 968 |
| **TGACC** | >Os04t0174200-01 | 4 | LOC_Os04g09450 | OsRAD23-like | 222 to 226, 484 to 488, 505 to 509, 945 to 949 |

| | | | | | |
|---|---|---|---|---|---|
| | >Os04t0202800-00 | 4 | LOC_Os04g12600 | H0512B01.12 protein. receptor-like protein kinase | 629 to 633, 642 to 646, 845 to 849, 928 to 932 |
| | >Os04t0287400-01 | 6 | LOC_Os04g21950 | WRKY51 | 94 to 98, 582 to 586, 592 to 596, 728 to 732, 930 to 934, 953 to 957 |
| | >Os04t0303100-00 | 4 | LOC_Os04g23700 | H0215E01.10 protein. lectin protein kinase family protein | 98 to 102, 374 to 378, 890 to 894, 951 to 955 |
| | >Os04t0398900-01 | 7 | LOC_Os04g32700 | H0209H04.6 protein. LEML3 - Anther-specific LEM1 family protein precursor | 333 to 337, 374 to 378, 413 to 417, 461 to 465, 508 to 512, 557 to 561, 637 to 641 |
| | >Os04t0408900-02 | 5 | LOC_Os04g33420 | DNA-binding protein S1FA | 260 to 264, 677 to 681, 684 to 688, 703 to 707, 745 to 749 |
| | >Os04t0408900-01 | 5 | LOC_Os04g33480 | Histone deacetylase. | 370 to 374, 787 to 791, 794 to 798, 813 to 817, 855 to 859 |
| | >Os04t0409600-01 | 4 | LOC_Os04g33480 | Histone deacetylase | 275 to 279, 349 to 353, 615 to 619, 941t o 945 |
| | >Os04t0449400-01 | 5 | LOC_Os04g37660 | H0818E04.11 protein. Protein phosphotase protein | 125 to 129, 253 to 257, 323 to 327, 752 to 756, 963 to 967 |
| | >Os04t0460200-00 | 4 | LOC_Os04g38660 | H0219H12.6 protein. transmembrane amino acid transporter protein | 168 to 172, 628 to 632, 663 to 667, 822 to 826 |
| | >Os04t0492600-01 | 4 | LOC_Os04g41510 | Anticodon-binding domain. serine/threonine-protein kinase GCN2 | 75 to 79, 152 to 156, 584 to 588, 852 to 856 |
| | >Os04t0498600-02 | 5 | LOC_Os04g42090 | S-adenosylmethionine decarboxylase proenzyme CPuORF7 - conserved peptide | 65 to 69, 631 to 635, 726 to 730, 762 to 766, 846 to 850 |
| | >Os04t0545700-00 | 5 | LOC_Os04g46140 | Transferase family protein. | 132 to 136, 147 to 151, 578 to 582, 745 to 749, 993 to 997 |
| | >Os04t0548300-02 | 5 | LOC_Os04g46310 | Armadillo-type fold domain. HEAT repeat family protein | 356 to 360, 395 to 399, 423 to 427, 511 to 515, 519 to 523 |
| | >Os04t0606700-01 | 4 | LOC_Os04g51770 | OSIGBa0113I13.5 protein | 596 to 600, 713 to 717, 738 to 742, 910 to 914 |
| | >Os04t0634000-01 | 4 | LOC_Os04g54140 | Serine/threonine protein kinase-related | 396 to 400, 408 to 412, 452 to 456, 466 to 470 |

| | | | | | |
|---|---|---|---|---|---|
| | >Os04t0634400-00 | 4 | LOC_Os04g54180 | H0315F07.5 protein. serine/threonine-protein kinase receptor precursor | 457 to 461, 698 to 702, 922 to 926, 991 to 995 |
| | >Os04t0691400-00 | 4 | LOC_Os04g59480 | POT family protein. POT family | 564 to 568, 616 to 620, 645 to 649, 653 to 657 |
| **TGACT** | >Os04t0111900-01 | 5 | LOC_Os04g02110 | Resistance gene analog PIC22 disease resistance protein | 760 to 764, 928 to 932, 932 to 936, 936 to 940, 946 to 950 |
| | >Os04t0111900-02 | 5 | LOC_Os04g02110 | Resistance gene analog PIC22 disease resistance protein | 755 to 759, 923 to 927, 927 to 931, 931 to 935, 941 to 945 |
| | >Os04t0188433-01 | 4 | LOC_Os04g10940 | OSIGBa0102N07.2 protein | 255 to 259, 700 to 704, 762 to 766, 835 to 839 |
| | >Os04t0194000-01 | 4 | LOC_Os04g11790 | Cyclin-like F-box domain. OsFBX120 - F-box domain | 198 to 202, 282 to 286, 557 to 561, 715 to 719 |
| | >Os04t0196200-00 | 4 | LOC_Os04g11970 | Winged helix repressor DNA-binding domain. O-methyltransferase | 426 to 430, 626 to 630, 658 to 662, 910 to 914 |
| | >Os04t0197500-00 | 4 | | UDP-glucuronosyl/UDP-glucosyltransferase family protein | 195 to 199, 307 to 311, 593 to 597, 986 to 990 |
| | >Os04t0204000-00 | 4 | LOC_Os04g12690 | N-hydroxythioamide S-beta-glucosyltransferase | 102 to 106, 125 to 129, 587 to 591, 635 to 639 |
| | >Os04t0208600-01 | 4 | LOC_Os04g13170 | Cyclin-like F-box domain. OsFBD10 - F-box and FBD domain | 224 to 228, 259 to 263, 670 to 674, 710 to 714 |
| | >Os04t0287400-01 | 4 | LOC_Os04g21950 | WRKY transcription factor 51 | 296 to 300, 413 to 417, 639 to 643, 908 to 912 |
| | >Os04t0291100-01 | 4 | LOC_Os04g22390 | Short-chain dehydrogenase Tic32 | 75 to 79, 238 to 242, 669 to 673, 822 to 826 |
| | >Os04t0291100-02 | 4 | LOC_Os04g22390 | Short-chain dehydrogenase Tic32. short chain dehydrogenase/ reductase protein, | 75 to 79, 238 to 242, 669 to 673, 822 to 826 |
| | >Os04t0369600-00 | 4 | LOC_Os04g30110 | OSIGBa0107E14.3 protein. wall-associated receptor kinase 3 precursor | 373 to 377, 484 to 488, 795 to 799, 823 to 827 |
| | >Os04t0373400-01 | 4 | LOC_Os04g30490 | MATE efflux family protein | 199 to 203, 285 to 289, 379 to 383, 991 to 995 |

| | | | | | |
|---|---|---|---|---|---|
| | >Os04t0379300-02 | 4 | LOC_Os04g31000 | Methyltransferase type 11 domain | 269 to 273, 302 to 306, 308 to 312, 590 to 594 |
| | >Os04t0421800-01 | 4 | LOC_Os04g34440 | EF-Hand type domain. ubiquitin interaction motif | 97 to 101, 244 to 248, 341 to 345, 622 to 626 |
| | >Os04t0432200-00 | 5 | LOC_Os04g35260 | DEAD/DEAH box helicase domain | 100 to 104, 166 to 170, 280 to 284, 315 to 319, 698 to 702 |
| | >Os04t0449500-01 | 4 | LOC_Os04g37670 | H0818E04.12 protein. steroid nuclear receptor, ligand-binding, | 167 to 171, 417 to 421, 762 to 766, 952 to 956 |
| | >Os04t0465800-01 | 4 | LOC_Os04g39170 | OSIGBa0115M15.1 protein. | 241 to 245, 630 to 634, 638 to 642, 717 to 721 |
| | >Os04t0465800-02 | 4 | LOC_Os04g39170 | OSIGBa0115M15.1 protein. | 153 to 157, 542 to 546, 550 to 554, 629 to 633 |
| | >Os04t0468000-01 | 4 | LOC_Os04g39300 | OSIGBa0128P10.7 protein. heavy metal transport/ detoxification protein | 863 to 867, 867 to 871, 871 to 875, 875 to 879 |
| | >Os04t0475100-00 | 4 | LOC_Os04g39910 | OSIGBa0106G07.2 protein. receptor-like protein kinase | 46 to 50, 50 to 54, 65 to 69, 411 to 415 |
| | >Os04t0491200-00 | 4 | LOC_Os04g41400 | Oligopeptide transporter domain. peptide transporter PTR2 | 592 to 596, 607 to 611, 650 to 654, 870 to 874 |
| | >Os04t0497200-01 | 4 | LOC_Os04g41970 | Cellulase precursor (Cellulase homolog OR16pep) | 260 to 264, 632 to 636, 651 to 655, 702 to 706 |
| | >Os04t0501700-01 | 4 | LOC_Os04g42330 | Spc97/Spc98 family protein. | 172 to 176, 178 to 182, 333 to 337, 558 to 562 |
| | >Os04t0513100-01 | 5 | LOC_Os04g43360 | Beta-glucosidase | 92 to 96, 237 to 241, 340 to 344, 552 to 556, 614 to 618 |
| | >Os04t0513400-01 | 4 | LOC_Os04g43390 | Beta-glucosidase | 235 to 239, 402 to 406, 551 to 555, 659 to 663 |
| | >Os04t0513400-02 | 4 | LOC_Os04g43390 | Beta-glucosidase | 235 to 239, 402 to 406, 551 to 555, 659 to 663 |
| | >Os04t0543000-01 | 4 | LOC_Os04g45920 | Protein kinase. protein kinase domain | 568 to 572, 903 to 907, 914 to 918, 988 to 992 |
| | >Os04t0589700-01 | 4 | LOC_Os04g49970 | Zinc finger, RING/FYVE/PHD-type domain. U-box | 135 to 139, 142 to 146, 260 to 264, 927 to 931 |
| | >Os04t0612500-01 | 4 | LOC_Os04g52260 | 36.4 kDa proline-rich protein. LTPL124 - Protease inhibitor/seed storage/LTP family protein precursor | 624 to 628, 664 to 668, 671 to 675, 803 to 807 |

| | | | | | |
|---|---|---|---|---|---|
| | >Os04t0612500-02 | 4 | LOC_Os04g52260 | 36.4 kDa proline-rich protein. LTPL124 - Protease inhibitor/seed storage/LTP family protein precursor | 345 to 349, 385 to 389, 392 to 396, 524 to 528 |
| | >Os04t0616300-01 | 4 | LOC_Os04g52600 | SHR5-receptor-like kinase | 46 to 50, 367 to 371, 697 to 701, 890 to 894 |
| | >Os04t0619300-02 | 4 | LOC_Os04g52830 | OsFBK15 - F-box domain and kelch repeat | 65 to 69, 239 to 243, 350 to 354, 994 to 998 |
| | >Os04t0620400-01 | 4 | LOC_Os04g52940 | SIT4 phosphatase-associated protein family protein. SIT4 phosphatase-associated protein domain | 160 to 164, 194 to 198, 210 to 214, 402 to 406 |
| | >Os04t0626500-01 | 5 | LOC_Os04g53496 | Disease resistance protein RGA2 (RGA2-blb) (Blight resistance protein RPI). NBS-LRR disease resistance protein | 166 to 170, 233 to 237, 712 to 716, 716 to 720, 937 to 941 |
| | >Os04t0636500-01 | 4 | LOC_Os04g54400 | BTB domain. BTBN12 - Bric-a-Brac, Tramtrack, Broad Complex BTB domain with non-phototropic hypocotyl 3 NPH3 and coiled-coil domains | 152 to 156, 329 to 333, 778 to 782, 860 to 864 |
| | >Os04t0644300-01 | 5 | LOC_Os04g55150 | UBA/TS-N domain | 74 to 78, 188 to 192, 639 to 643, 832 to 836, 967 to 971 |
| | >Os04t0647800-01 | 4 | LOC_Os04g55410 | Glycerol kinase-like protein. FGGY family of carbohydrate kinases | 36 to 40, 336 to 340, 342 to 346, 445 to 449 |
| | >Os04t0647800-02 | 4 | LOC_Os04g55410 | Glycerol kinase-like protein. FGGY family of carbohydrate kinases | 198 to 202, 204 to 208, 307 to 311, 888 to 892 |
| | >Os04t0652600-01 | 4 | LOC_Os04g55840 | OSIGBa0113E10.15 protein | 232 to 236, 295 to 299, 379 to 383, 778 to 782 |
| | >Os04t0655000-00 | 4 | LOC_Os04g56090 | Curculin-like (mannose-binding) lectin domain. S-locus-like receptor protein kinase | 558 to 562, 650 to 654, 812 to 816, 879 to 883 |
| | >Os04t0682000-01 | 4 | LOC_Os04g58560 | Autophagy 4a. cysteine protease ATG4 | 113 to 117, 302 to 306, 574 to 578, 926 to 930 |
| **TTGAC** | >Os04t0106000-01 | 4 | LOC_Os04g01570 | Pectinesterase inhibitor domain. Invertase/pectin methylesterase inhibitor family protein | 272 to 276, 660 to 664, 734 to 738, 743 to 747 |

| | | | | | |
|---|---|---|---|---|---|
| | >Os04t0109500-01 | 4 | LOC_Os04g01910 | Legume lectin, beta domain | 4 to 8, 61 to 65, 284 to 288, 528 to 532 |
| | >Os04t0110100-01 | 4 | LOC_Os04g01950 | Concanavalin A-like lectin/glucanase domain | 745 to 749, 860 to 864, 878 to 882, 928 to 932 |
| | >Os04t0111900-01 | 4 | LOC_Os04g02110 | Resistance gene analog PIC22 disease resistance protein RGA3 | 164 to 168, 719 to 723, 927 to 931, 945 to 949 |
| | >Os04t0111900-02 | 4 | LOC_Os04g02110 | Resistance gene analog PIC22 disease resistance protein RGA3 | 159 to 163, 714 to 718, 922 to 926, 940 to 944 |
| | >Os04t0163425-00 | 4 | LOC_Os04g08170 | NAF domain domain. CBL-interacting protein kinase | 778 to 782, 840 to 844, 890 to 894, 917 to 921 |
| | >Os04t0165600-01 | 4 | LOC_Os04g08340 | Peptidase S26A, signal peptidase I family protein. OsSigP3 - Putative Type I Signal Peptidase homologue; employs a putative Ser/Lys catalytic dyad | 433 to 437, 465 to 469, 741 to 745, 930 to 934 |
| | >Os04t0168400-01 | 5 | LOC_Os04g08640 | Embryogenesis transmembrane protein. cadmium tolerance factor | 182 to 186, 257 to 261, 644 to 648, 885 to 889, 984 to 988 |
| | >Os04t0188433-01 | 4 | LOC_Os04g10940 | OSIGBa0102N07.2 protein | 634 to 638, 671 to 675, 834 to 838, 965 to 969 |
| | >Os04t0197200-01 | 4 | LOC_Os04g12080 | Protein kinase, core domain containing protein. TKL_IRAK_DUF26-lc.7 | 155 to 159, 180 to 184, 328 to 332, 565 to 569 |
| | >Os04t0212700-00 | 4 | LOC_Os04g13530 | Histone H2A. Core histone H2A/H2B/H3/H4 domain | 151 to 155, 476 to 480, 566 to 570, 625 to 629 |
| | >Os04t0282200-01 | 4 | LOC_Os04g21340 | Protein of unknown function DUF1685 family protein | 168 to 172, 231 to 235, 631 to 635, 985 to 989 |
| | >Os04t0319600-01 | 4 | LOC_Os04g25360 | tRNA methyltransferase complex GCD14 subunit domain. tRNA methyltransferase complex GCD14 subunit | 178 to 182, 669 to 673, 755 to 759, 763 to 767 |
| | >Os04t0341100-00 | 4 | LOC_Os04g27300 | OSIGBa0159H11-OSIGBa0137A07.5 protein | 375 to 379, 638 to 642, 657 to 661, 888 to 892 |
| | >Os04t0356600-01 | 4 | LOC_Os04g28780 | OSIGBa0105P02.3 protein. serine/threonine-protein kinase receptor precursor | 195 to 199, 514 to 518, 617 to 621, 799 to 803 |
| | >Os04t0356600-01 | 5 | LOC_Os04g28780 | OSIGBa0105P02.3 protein. serine/threonine-protein kinase receptor precursor | 195 to 199, 514 to 518, 617 to 621, 799 to 803 |
| | >Os04t0371000-00 | 4 | LOC_Os04g30250 | Four-helical cytokine family | 254 to 258, 669 to 673, 677 to 681, 688 |

| | | | | | |
|---|---|---|---|---|---|
| | | | | protein. wall-associated receptor kinase-like 5 precursor | to 692 |
| | >Os04t0373400-01 | 5 | LOC_Os04g30490 | MATE efflux family protein | 198 to 202, 284 to 288, 378 to 382, 668 to 672, 990 to 994 |
| | >Os04t0382200-01 | 4 | LOC_Os04g31330 | Exo70 exocyst complex subunit family protein | 259 to 263, 665 to 669, 708 to 712, 714 to 718 |
| | >Os04t0382300-01 | 4 | LOC_Os04g31340 | SNF1-related protein kinase regulatory gamma subunit 1 (AKIN gamma1) (AKING1). CBS domain containing membrane protein | 69 to 73, 106 to 110, 147 to 151, 186 to 190 |
| | >Os04t0388000-00 | 4 | LOC_Os04g31880 | OSIGBa0075F02.5 protein | 211 to 215, 355 to 359, 729 to 733, 909 to 913 |
| | >Os04t0393500-00 | 4 | LOC_Os04g32300 | H0718E12.1 protein. OsGrx_C9 - glutaredoxin subgroup III | 202 to 206, 280 to 284, 742 to 746, 767 to 771 |
| | >Os04t0399800-00 | 4 | LOC_Os04g32790 | Ethylene response factor 82 | 70 to 74, 139 to 143, 440 to 444, 513 to 517 |
| | >Os04t0400800-01 | 4 | LOC_Os04g32840 | Heavy metal transport/ detoxification protein domain. formin-like protein 20 | 39 to 43, 240 to 244, 474 to 478, 483 to 487 |
| | >Os04t0420300-00 | 4 | LOC_Os04g34300 | H0525E10.1 protein. serine/ threonine-protein kinase receptor precursor | 359 to 363, 746 to 750, 893 to 897, 899 to 903 |
| | >Os04t0420600-00 | 4 | LOC_Os04g34330 | H0525E10.2 protein. serine/threonine-protein kinase receptor precursor | 36 to 40, 293 to 297, 586 to 590, 594 to 598 |
| | >Os04t0420900-01 | 4 | LOC_Os04g34370 | H0525E10.7 protein. serine/threonine-protein kinase receptor precursor | 228 to 232, 241 to 245, 778 to 782, 914 to 918 |
| | >Os04t0420900-02 | 4 | LOC_Os04g34370 | H0525E10.7 protein. serine/threonine-protein kinase receptor precursor | 200 to 204, 213 to 217, 750 to 754, 886 to 890 |
| | >Os04t0430200-01 | 5 | LOC_Os04g35100 | PLC-like phosphodiesterase, TIM beta/alpha-barrel domain domain. phospholipase C | 228 to 232, 241 to 245, 778 to 782, 914 to 918 |
| | >Os04t0440100-01 | 4 | LOC_Os04g35920 | H0525C06.10 protein. nonsense-mediated decay UPF3 | 98 to 102, 107 to 111, 449 to 453, 865 to 869 |

| | | | | | |
|---|---|---|---|---|---|
| | >Os04t0440100-02 | 4 | LOC_Os04g35920 | H0525C06.10 protein. nonsense-mediated decay UPF3 | 75 to 79, 84 to 88, 426 to 430, 842 to 846 |
| | >Os04t0455401-03 | 4 | LOC_Os04g38290 | OSIGBa0093K19.10 protein | 37 to 41, 309 to 313, 320 to 324, 372 to 376 |
| | >Os04t0469300-01 | 4 | LOC_Os04g39380 | ATFP4. heavy metal transport/detoxification protein | 198 to 202, 310 to 314, 338 to 342, 386 to 390 |
| | >Os04t0472900-01 | 4 | LOC_Os04g39680 | C2 calcium-dependent membrane targeting domain. anthranilate phosphoribosyltransferase, | 80 to 84, 378 to 382, 465 to 469, 966 to 970 |
| | >Os04t0480500-01 | 6 | LOC_Os04g40440 | Leucine-rich repeat, N-terminal domain | 136 to 140, 531 to 535, 716 to 720, 721 to 725, 888 to 892, 928 to 932 |
| | >Os04t0494400-01 | 4 | LOC_Os04g41700 | R4 - Corn type-A response regulator | 145 to 149, 190 to 194, 257 to 261, 848 to 852 |
| | >Os04t0495500-01 | 4 | LOC_Os04g41820 | OSIGBa0159F11.9 protein. transcription factor RF2a | 15 to 19, 621 to 625, 809 to 813, 881 to 885 |
| | >Os04t0501600-02 | 4 | LOC_Os04g42320 | AT hook, DNA-binding, conserved site domain | 47 to 51, 76 to 80, 274 to 278, 611 to 615 |
| | >Os04t0508900-00 | 6 | LOC_Os04g42990 | OSIGBa0101P20.13 protein. suppressor of stem-loop protein 1 | 363 to 367, 470 to 474, 514 to 518, 684 to 688<br>942 to 946, 963 to 967 |
| | >Os04t0513100-01 | 5 | LOC_Os04g43360 | Beta-glucosidase. Os4bglu14 - monolignol beta-glucoside homologue without catalytic acid/base | 91 to 95, 236 to 240, 339 to 343, 613 to 617<br>673 to 677 |
| | >Os04t0513400-01 | 4 | LOC_Os04g43390 | Beta-glucosidase. Os4bglu16 - monolignol beta-glucoside homologue | 150 to 154, 401 to 405, 550 to 554, 658 to 662 |
| | >Os04t0513400-02 | 4 | LOC_Os04g43390 | Beta-glucosidase. Os4bglu16 - monolignol beta-glucoside homologue | 150 to 154, 401 to 405, 550 to 554, 658 to 662 |
| | >Os04t0525200-01 | 4 | LOC_Os04g44354 | UDP-glucuronosyl/UDP-glucosyltransferase family protein | 287 to 291, 324 to 328, 371 to 375, 782 to 786 |
| | >Os04t0548100-01 | 4 | LOC_Os04g46300 | OSIGBa0106P14.3 protein. disease resistance protein RPM1 | 21 to 25, 926 to 930, 934 to 938, 969 to 973 |
| | >Os04t0552000-00 | 5 | LOC_Os04g46630 | Barwin-related endoglucanase domain. expansin precursor | 67 to 71, 126 to 130, 285 to 289, 354 to 358, 735 to 739 |

| | | | | | |
|---|---|---|---|---|---|
| | >Os04t0566100-00 | 4 | LOC_Os04g47830 | OSIGBa0158F05.11 protein.SNF2 family N-terminal domain | 151 to 155, 298 to 302, 379 to 383, 649 to 653 |
| | >Os04t0574100-01 | 4 | LOC_Os04g48480 | Exostosin-like family protein | 342 to 346, 351 to 355, 370 to 374, 823 to 827 |
| | >Os04t0574100-02 | 4 | LOC_Os04g48480 | Exostosin-like family protein | 130 to 134, 139 to 143, 158 to 162, 611 to 615 |
| | >Os04t0585600-01 | 4 | LOC_Os04g49610 | H0307D04.4 protein | 44 to 48, 210 to 214, 334 to 338, 788 to 792 |
| | >Os04t0588350-00 | 4 | LOC_Os04g49870 | Nucleotidylntransferase family protein | 22 to 26, 144 to 148, 367 to 371, 818 to 822 |
| | >Os04t0589700-01 | 4 | LOC_Os04g49970 | Zinc finger, RING/FYVE/PHD-type domain | 141 to 145, 259 to 263, 622 to 626, 926 to 930 |
| | >Os04t0590100-00 | 4 | LOC_Os04g50020 | Heavy metal-associated domain, HMA domain | 531 to 535, 622 to 626, 695 to 699, 726 to 730 |
| | >Os04t0603000-00 | 4 | LOC_Os04g51390 | OSIGBa0118P15.1 protein. aldose 1-epimerase | 134 to 138, 210 to 214, 520 to 524, 528 to 532 |
| | >Os04t0608300-01 | 4 | LOC_Os04g51890 | Auxin responsive SAUR protein domain. OsSAUR20 | 83 to 87, 236 to 240, 367 to 371, 481 to 485 |
| | >Os04t0609200-01 | 4 | LOC_Os04g51970 | Major facilitator superfamily protein. synaptic vesicle 2-related protein | 263 to 267, 294 to 298, 328 to 332, 960 to 964 |
| | >Os04t0610800-01 | 4 | LOC_Os04g52130 | Coproporphyrinogen III oxidase (Fragment) | 87 to 91, 238 to 242, 243 to 247, 598 to 602 |
| | >Os04t0612500-01 | 4 | LOC_Os04g52260 | 36.4 kD proline-rich protein. LTPL124 - Protease inhibitor/seed storage/LTP family protein precursor | 13 to 17, 623 to 627, 663 to 667, 670 to 674 |
| | >Os04t0613700-01 | 5 | LOC_Os04g52370 | UDP-N-acetylglucosamine pyrophosphorylase. UTP-glucose-1-phosphate uridylyltransferase | 67 to 71, 76 to 80, 550 to 554, 757 to 761, 847 to 851 |
| | >Os04t0616300-01 | 4 | LOC_Os04g52600 | H0525G02.10 protein. SHR5-receptor-like kinase | 366 to 370, 622 to 626, 696 to 700, 831 to 835 |
| | >Os04t0618700-01 | 4 | LOC_Os04g52780 | Protein kinase, core domain. leucine-rich repeat receptor protein kinase EXS precursor | 25 to 29, 891 to 895, 905 to 909, 926 to 930 |
| | >Os04t0620500-01 | 4 | LOC_Os04g52940 | H0714H04.7 protein. SIT4 phosphatase-associated protein domain | 259 to 263, 378 to 382, 450 to 454, 833 to 837 |

| | | | | | |
|---|---|---|---|---|---|
| | >Os04t0629100-01 | 5 | LOC_Os04g53700 | H0303G06.16 protein. zinc finger protein | 29 to 33, 174 to 178, 581 to 585, 586 to 590, 828 to 832 |
| | >Os04t0629100-02 | 4 | LOC_Os04g53700 | H0303G06.16 protein. zinc finger protein | 123 to 127, 530 to 534, 535 to 539, 777 to 781 |
| | >Os04t0647800-02 | 4 | LOC_Os04g55410 | Glycerol kinase-like protein.FGGY family of carbohydrate kinases | 197 to 201, 203 to 207, 306 to 310, 887 to 891 |
| | >Os04t0648600-01 | 6 | LOC_Os04g55490 | H0821G03.14 protein | 199 to 203, 667 to 671, 703 to 707, 753 to 757, 781 to 785, 878 to 882 |
| | >Os04t0649051-01 | 4 | LOC_Os04g55555 | H0211B05.9 protein. retrotransposon protein, | 189 to 193, 476 to 480, 713 to 717, 890 to 894 |
| | >Os04t0652700-01 | 4 | LOC_Os04g55850 | Nuclease PA3. nuclease PA3 | 57 to 61, 68 to 72, 518 to 522, 537 to 541 |
| | >Os04t0659900-01 | 4 | LOC_Os04g56480 | OSIGBa0132E09-OSIGBa0108L24.21 protein | 180 to 184, 207 to 211, 507 to 511, 641 to 645 |
| | >Os04t0659900-02 | 4 | LOC_Os04g56480 | OSIGBa0132E09-OSIGBa0108L24.21 protein | 170 to 174, 197 to 201, 497 to 501, 631 to 635 |
| | >Os04t0685100-01 | 4 | LOC_Os04g58830 | Ribosomal biogenesis regulatory protein family protein | 95 to 99, 114 to 118, 337 to 341, 569 to 573 |
| **TTGACA** | >Os04t0692400-01 | 5 | LOC_Os04g59590 | Ankyrin-like protein | 133 to 137, 165 to 169, 291 to 295, 297 to 301, 595 to 599 |
| | >Os04t0282200-01 | 4 | LOC_Os04g21340 | Protein of unknown function DUF1685 family protein | 168 to 173, 231 to 236, 631 to 636, 985 to 990 |
| | >Os04t0525200-01 | 4 | LOC_Os04g44354 | UDP-glucuronosyl/UDP-glucosyltransferase family protein. | 287 to 292, 324 to 329, 371 to 376, 782 to 787 |
| | >Os04t0648600-01 | 5 | LOC_Os04g55490 | H0821G03.14 protein. | 199 to 204, 667 to 672, 703 to 708, 753 to 758, 781 to 786 |
| **TTGACT** | >Os04t0373400-01 | 4 | LOC_Os04g30490 | MATE efflux family protein | 198 to 203, 284 to 289, 378 to 383, 990 to 995 |
| | >Os04t0513100-01 | 4 | LOC_Os04g43360 | Beta-glucosidase14 | 91 to 962, 36 to 241, 339 to 344, 613 to 618 |
| | >Os04t0647800-02 | 4 | LOC_Os04g55410 | Glycerol kinase-like protein. FGGY family of carbohydrate kinases | 197 to 202, 203 to 208, 306 to 311, 887 to 892 |
| **Chromosome 5** | | | | | |
| **Motif** | **ID** | **Frequency** | **Locus ID** | **Protein name** | **Positions in 1k promoter region** |

| | | | | | |
|---|---|---|---|---|---|
| **TTTGACT** | >Os05t0155100-01 | 4 | LOC_Os05g06310 | 60S ribosomal protein L18-3 | 196 to 202, 363 to 369, 445 to 451, 545 to 551 |
| | >Os05t0241100-01 | 4 | LOC_Os05g15150; | leucyl-tRNA synthetase | 129 to 351, 66to172, 283 to 289, 401 to 407 |
| **TTGACTT** | >Os05t0241100-01 | 4 | LOC_Os05g15150 | leucyl-tRNA synthetase | 30 to 36, 167 to 173, 284 to 290, 402 to 408 |
| **TGACC** | >Os05t0117798-01 | 5 | LOC_Os05g02670 | Kinesin, motor region domain | 223 to 227, 800 to 804, 883 to 887, 891 to 895, 978 to 982 |
| | >Os05t0144200-01 | 4 | LOC_Os05g05220 | Glioma tumor suppressor-like protein | 93 to 97, 347 to 351, 444 to 448, 915 to 919 |
| | >Os05t0201700-00 | 5 | LOC_Os05g11210 | Lachrymatory factor synthase, | 152 to 156, 196 to 200, 387 to 391, 474 to 478, 693 to 697 |
| | >Os05t0295300-01 | 4 | LOC_Os05g22940 | Acetyl-coenzyme A carboxylase acetyl-CoA carboxylase | 87 to 91, 222 to 226, 549 to 553, 972 to 976 |
| | >Os05t0357600-01 | 7 | LOC_Os05g28940 | Developmentally regulated GTP binding protein | 45 to 49, 56 to 60, 115 to 119, 141 to 145, 208 to 212, 217 to 221, 740 to 744 |
| | >Os05t0357600-02 | 7 | LOC_Os05g28940 | Developmentally regulated GTP binding protein | 17 to 21, 28 to 32, 87 to 91, 113 to 117, 180 to 184, 189 to 193, 712 to 716 |
| | >Os05t0403400-01 | 4 | LOC_Os05g33440 | Mitochodrial transcription termination factor-related family protein | 130 to 134, 268 to 272, 396 to 400, 472 to 476 |
| | >Os05t0454200-01 | 4 | LOC_Os05g38000 | Pollen-specific kinase partner protein, ATROPGEF7/ROPGEF7 | 349 to 353, 502 to 506, 524 to 528, 610 to 614 |
| | >Os05t0461300-01 | 4 | LOC_Os05g38630 | RAB8C. ras-related protein | 692 to 696, 918 to 922, 923 to 927, 929 to 933 |
| | >Os05t0461300-02 | 4 | LOC_Os05g38630 | RAB8C. ras-related protein | 616 to 620, 842 to 846, 847 to 851, 853 to 857 |
| | >Os05t0489600-01 | 4 | LOC_Os05g41060 | ADP-ribosylation factor 1. ADP-ribosylation factor | 435 to 439, 759 to 763, 836 to 840, 943 to 947 |
| | >Os05t0489600-03 | 4 | LOC_Os05g41060 | ADP-ribosylation factor 1. ADP-ribosylation factor | 395 to 399 719 to 723, 796 to 800, 903 to 907 |
| | >Os05t0491000-01 | 5 | LOC_Os05g41200 | EF-Hand type domain. OsCML9 Calmodulin-related calcium sensor protein | 389 to 393, 793 to 797, 799 to 803, 812 to 816, 961 to 965 |
| | >Os05t0499450-00 | 4 | LOC_Os05g42010 | TraB family protein | 380 to 384, 395 to 399, 732 to 736, 777to781 |

| | | | | | |
|---|---|---|---|---|---|
| | >Os05t0513800-02 | 4 | LOC_Os05g43820 | GTP-binding protein OsRac2. ras-related protein | 446 to 450, 511 to 515, 801 to 805, 922 to 926 |
| | >Os05t0522600-01 | 5 | LOC_Os05g44770 | Leucine-rich repeat, plant specific. receptor-like protein kinase 5 precursor | 16 to 20, 168 to 172, 291, to 295, 547 to 551, 776 to 780 |
| | >Os05t0596000-01 | 5 | LOC_Os05g51750 | Peptidase A1 domain. aspartyl protease family protein | 104 to 108, 137 to 141, 177 to 181, 211 to 215, 475 to 479 |
| **TGACT** | >Os05t0125500-01 | 4 | LOC_Os05g03480 | Isovaleryl-CoA dehydrogenase, mitochondrial precursor | 61 to 65, 163 to 167, 476 to 480, 672 to 676 |
| | >Os05t0128000-00 | 5 | LOC_Os05g03740 | Homeodomain-like. transcription factor TF2 | 165 to 169, 506 to 510, 557 to 561, 782 to 786, 981 to 985 |
| | >Os05t0135700-01 | 4 | LOC_Os05g04510 | S-adenosylmethionine synthetase | 49 to 53, 148 to 152, 282 to 286, 296 to 300 |
| | >Os05t0155100-01 | 4 | LOC_Os05g06310 | 60S ribosomal protein L18. | 198 to 202, 365 to 369, 447 to 451, 547 to 551 |
| | >Os05t0157200-01 | 4 | LOC_Os05g06500 | UspA domain. universal stress protein domain | 20 to 24, 135 to 139, 376 to 380, 730 to 734 |
| | >Os05t0199800-01 | 4 | LOC_Os05g11070 | BHLH transcription activator Ivory seed. helix-loop-helix DNA-binding domain | 245 to 249, 339 to 343, 509 to 513, 677 to 681 |
| | >Os05t0241100-01 | 4 | LOC_Os05g15150 | leucyl-tRNA synthetase, cytoplasmic | 31 to 35, 168 to 172, 285 to 289, 403 to 407 |
| | >Os05t0244900-00 | 4 | LOC_Os05g15600 | Transcription elongation factor, TFIIS/CRSP70, N-terminal, sub-type domain | 482 to 486, 613 to 617, 633 to 637, 684 to 688 |
| | >Os05t0349200-01 | 5 | LOC_Os05g28180 | AMP deaminase 1 AMP deaminase | 291 to 295, 368 to 372, 501 to 505, 751 to 755, 855 to 859 |
| | >Os05t0356700-01 | 4 | LOC_Os05g28830 | Protein of unknown function DUF231, plant. PMR5 | 144 to 148, 699 to 703,818 to 822, 870 to 874 |
| | >Os05t0363566-00 | 4 | LOC_Os05g30010 | WD domain, G-beta repeat domain | 545 to 549, 567 to 571, 649 to 653, 720 to 724 |
| | >Os05t0371600-01 | 4 | LOC_Os05g30820 | BRASSINOSTEROID INSENSITIVE 1-associated receptor kinase 1 | 36 to 40, 83 to 87, 159 to 163, 397 to 401 |
| | >Os05t0378800-03 | 4 | LOC_Os05g31480 | Ankyrin like protein | 579 to 583, 647 to 651, 657 to 661, 864 to 868 |

| | | | | | |
|---|---|---|---|---|---|
| | >Os05t0436100-00 | 4 | LOC_Os05g36050 | Serine/threonine-protein kinase | 202 to 206, 528 to 532, 730 to 734, 917 to 921 |
| | >Os05t0458500-00 | 4 | LOC_Os05g38410 | Laccase (EC 1.10.3.2). laccase precursor protein | 85 to 89, 238 to 242, 318 to 322, 556 to 560 |
| | >Os05t0468400-01 | 5 | LOC_Os05g39210 | Domain of unknown function DUF1618 domain | 136 to 140, 319 to 323, 529 to 533, 856 to 860, 895 to 899 |
| | >Os05t0480000-01 | 4 | LOC_Os05g40180 | Serine/threonine protein kinase-related domain | 11 to 15, 279 to 283, 348 to 352, 622 to 626 |
| | >Os05t0486700-01 | 4 | LOC_Os05g40820 | Ribosomal protein L24e domain. ribosomal protein L24 | 219 to 223, 272 to 276, 330 to 334, 643 to 647 |
| | >Os05t0493100-01 | 5 | LOC_Os05g41370 | KI domain interacting kinase 1, TKL_IRAK_DUF26-la.1 - DUF26 kinases | 272 to 276, 303 to 307, 471 to 475, 555 to 559, 908 to 912 |
| | >Os05t0497700-01 | 4 | LOC_Os05g41810 | Peptidase. OsDegp8 - Putative Deg protease homologue | 171 to 175, 316 to 320, 419 to 423, 782 to 786 |
| | >Os05t0500400-01 | 4 | LOC_Os05g42100 | Domain of unknown function DUF1995 domain | 207 to 211, 393 to 397, 721 to 725, 749 to 753 |
| | >Os05t0519700-01 | 4 | LOC_Os05g44340 | Heat shock protein 101 | 702 to 706, 766 to 770, 771 to 775, 801 to 805 |
| | >Os05t0519700-02 | 4 | LOC_Os05g44340 | Heat shock protein 101 | 648 to 652, 712 to 716, 717 to 721, 747 to 751 |
| | >Os05t0527100-00 | 4 | LOC_Os05g45110 | UDP-glucuronosyl/UDP-glucosyltransferase family protein. anthocyanidin 5,3-O-glucosyltransferase | 277 to 281, 318 to 322, 696 to 700, 779 to 783 |
| | >Os05t0555900-01 | 4 | LOC_Os05g48220 | 60S ribosomal protein L35a-3 | 504 to 508, 656 to 660, 707 to 711, 796 to 800 |
| | >Os05t0556000-01 | 4 | LOC_Os05g48230 | Oxoglutarate/iron-dependent oxygenase domain | 29 to 33, 118 to 122, 517 to 521, 789 to 793 |
| | >Os05t0582600-01 | 4 | LOC_Os05g50580 | Peptidase S10, serine carboxypeptidase family protein. OsSCP30 - Putative Serine Carboxypeptidase homologue | 51 to 55, 559 to 563, 768 to 772, 813 to 817 |
| **TTGAC** | >Os05t0148600-01 | 4 | LOC_Os05g05590 | Na /H antiporter. transporter, monovalent cation: proton antiporter-2 family | 60 to 64, 81 to 85, 939 to 943, 947 to 951 |
| | >Os05t0152900-00 | 4 | LOC_Os05g06100 | Seven-in-absentia protein, | 38 to 42, 164 to 168, 252 to 256, 992 to |

| | | | | | |
|---|---|---|---|---|---|
| | | | | sina domain. E3 ubiquitin-protein ligase SINA-like 2 | 996 |
| | >Os05t0153000-01 | 5 | LOC_Os05g06110 | Gelsolin family protein. villin, | 23 to 27, 339 to 343, 350 to 354, 550 to 554, 673 to 677 |
| | >Os05t0153000-02 | 4 | LOC_Os05g06110 | Gelsolin family protein. villin | 203 to 207, 214 to 218, 414 to 418, 537 to 541 |
| | >Os05t0155100-01 | 6 | LOC_Os05g06310 | 60S ribosomal protein L18. 60S ribosomal protein L18-3 | 153 to 157, 197 to 201, 281 to 285, 364 to 368, 446 to 450, 546 to 550 |
| | >Os05t0158400-01 | 4 | LOC_Os05g06650 | Peptidase, trypsin-like serine and cysteine domain | 473 to 477, 492 to 496, 781 to 785, 800 to 804 |
| | >Os05t0165900-01 | 4 | LOC_Os05g07300 | Serine/threonine protein kinase-related domain. serine/threonine-protein kinase receptor precursor | 241 to 245, 695 to 699, 714 to 718, 861 to 865 |
| | >Os05t0166300-01 | 5 | LOC_Os05g07420 | Serine/threonine protein kinase-related domain. S-domain receptor-like protein kinase | 366 to 370, 548 to 552, 567 to 571, 858 to 862, 882 to 886 |
| | >Os05t0170200-01 | 4 | LOC_Os05g07810 | Universal stress protein | 47 to 51, 437 to 441, 465 to 469, 693 to 697 |
| | >Os05t0170200-02 | 4 | LOC_Os05g07810 | Universal stress protein | 47 to 51, 437 to 441, 465 to 469, 693 to 697 |
| | >Os05t0178100-01 | 4 | LOC_Os05g08540 | GA 3beta-hydroxylase1, GA metabolism gibberellin 3-beta-dioxygenase 2-2 | 48 to 52, 392 to 396, 516 to 520, 535 to 539 |
| | >Os05t0199800-01 | 5 | LOC_Os05g11070 | BHLH transcription activator Ivory seed. helix-loop-helix DNA-binding domain | 244 to 248, 508 to 512, 667 to 671, 676 to 680, 695 to 699 |
| | >Os05t0213900-01 | 4 | LOC_Os05g12280 | Virulence factor, pectin lyase fold family protein. clumping factor A precursor | 110 to 114, 673 to 677, 682 to 686, 700 to 704 |
| | >Os05t0237100-01 | 4 | LOC_Os05g14730 | Cyclin, C-terminal domain. cyclin-A1 | 434 to 438, 479 to 483, 781 to 785, 981 to 985 |
| | >Os05t0237400-01 | 5 | LOC_Os05g14750 | Viroid symptom modulation protein. AGC_PVPK_like_kin82y.12 - ACG kinases include homologs to PKA, PKG and PKC | 186 to 190, 257 to 261, 415 to 419, 636 to 640, 765 to 769 |

| | | | | | |
|---|---|---|---|---|---|
| | >Os05t0241100-01 | 4 | LOC_Os05g15150 | Predicted protein. leucyl-tRNA synthetase, cytoplasmic | 30 to 34, 167 to 171, 284 to 288, 402 to 406 |
| | >Os05t0325000-00 | 5 | LOC_Os05g25930 | OSIGBa0140J09.3 protein | 248 to 252, 596 to 600, 688 to 692, 748 to 752, 775 to 779 |
| | >Os05t0341900-00 | 4 | LOC_Os05g27570 | SAG20 | 124 to 128, 714 to 718, 726 to 730, 792 to 796 |
| | >Os05t0354400-02 | 5 | LOC_Os05g28630 | ESK1 (ESKIMO 1). leaf senescence related protein | 3 to 7, 23 to 27, 524 to 528, 766 to 770, 785 to 789 |
| | >Os05t0358200-01 | 5 | LOC_Os05g29010 | DNA primase. POLA4 - Putative DNA polymerase alpha complex subunit | 202 to 206, 509 to 513, 536 to 540, 770 to 774, 871 to 875 |
| | >Os05t0361500-01 | 4 | LOC_Os05g29790 | Pectin esterase | 223 to 227, 244 to 248, 638 to 642, 779 to 783 |
| | >Os05t0372200-00 | 4 | LOC_Os05g30880 | Surfeit locus 6 family protein. nucleolar matrix protein-related | 20 to 24, 64 to 68, 104 to 108, 533 to 537 |
| | >Os05t0376600-01 | 4 | LOC_Os05g31254 | BRCT domain. acetyltransferase, GNAT family | 213 to 217, 357 to 361, 604 to 608, 623 to 627 |
| | >Os05t0378800-03 | 4 | LOC_Os05g31480 | Ankyrin like protein | 273 to 277, 297 to 301, 578 to 582, 646 to 650 |
| | >Os05t0389000-01 | 4 | LOC_Os05g32270 | AP2-1 protein (Fragment) | 265 to 269, 315 to 319, 591 to 595, 752 to 756 |
| | >Os05t0389500-01 | 4 | LOC_Os05g32330 | Endonuclease/exonuclease/phosphatase domain | 261 to 265, 360 to 364, 508 to 512, 536 to 540 |
| | >Os05t0389500-02 | 4 | LOC_Os05g32330 | Endonuclease/exonuclease/phosphatase domain | 232 to 236, 331 to 335, 479 to 483, 507 to 511 |
| | >Os05t0389500-03 | 4 | LOC_Os05g32330 | Endonuclease/exonuclease/phosphatase domain | 219 to 223, 318 to 322, 466 to 470, 494 to 498 |
| | >Os05t0393100-01 | 5 | LOC_Os05g32660 | Leucine-rich repeat, N-terminal domain | 42 to 46, 91 to 95, 99 to 103, 650 to 654, 969 to 973 |
| | >Os05t0403000-01 | 4 | LOC_Os05g33410 | Peptidase aspartic, catalytic domain. Xylanase inhibitor | 50 to 54, 115 to 119, 364 to 368, 518 to 522 |
| | >Os05t0404300-01 | 4 | LOC_Os05g33514 | AlNc14C59G4370 protein | 145 to 149, 563 to 567, 647 to 651, 876 to 880 |
| | >Os05t0438600-01 | 4 | LOC_Os05g36270 | Fructose-1,6-bisphosphatase class 1/Sedoheputulose-1,7-bisphosphatase domain | 127 to 131, 310 to 314, 333 to 337, 834 to 838 |

| | | | | | |
|---|---|---|---|---|---|
| | >Os05t0439000-01 | 4 | LOC_Os05g36310 | RING-H2 finger protein ATL1O zinc finger, C3HC4 type domain | 152 to 156, 466 to 470, 737 to 741, 795 to 799 |
| | >Os05t0439400-01 | 4 | LOC_Os05g36360 | Arm repeat. U-box domain | 719 to 723, 925 to 929, 931 to 935, 938 to 942 |
| | >Os05t0442550-01 | 4 | LOC_Os05g37070 | H0801D08.16 protein. Retrotransposon, Ty1-copia subclass | 196 to 200, 207 to 211, 298 to 302, 888 to 892 |
| | >Os05t0447500-00 | 4 | LOC_Os05g37500 | Mediator complex, subunit Med13 domain | 561 to 565, 642 to 646, 675 to 679, 821 to 825 |
| | >Os05t0452600-01 | 4 | LOC_Os05g37884 | 50S ribosomal protein L33. 50S ribosomal protein L33 | 53 to 57, 208 to 212, 336 to 340, 631 to 635 |
| | >Os05t0468200-01 | 5 | LOC_Os05g39190 | Domain of unknown function DUF1618 domain | 341 to 345, 431 to 435, 582 to 586, 768 to 772, 823 to 827 |
| | >Os05t0480000-01 | 4 | LOC_Os05g40180 | Serine/threonine protein kinase-related domain | 10 to 14, 42 to 46, 278 to 282, 621 to 625 |
| | >Os05t0486700-01 | 5 | LOC_Os05g40820 | Ribosomal protein L24e domain | 139 to 143, 317 to 321, 525 to 529, 593 to 597, 642 to 646 |
| | >Os05t0492600-00 | 4 | LOC_Os05g41310 | NBS-LRR type resistance protein disease resistance protein RGA2 | 715 to 719, 863 to 867, 891 to 895, 898 to 902 |
| | >Os05t0496500-01 | 4 | LOC_Os05g41670 | Latex-abundant protein. ICE-like protease p20 domain | 351 to 355, 612 to 616, 648 to 652, 665 to 669 |
| | >Os05t0501350-00 | 4 | LOC_Os05g42200 | Cyclin-B1-5 | 479 to 483, 508 to 512, 535 to 539 , 935 to 939 |
| | >Os05t0508700-03 | 4 | LOC_Os05g43280 | TRAF-type domain, MATH domain | 412 to 416, 623 to 627, 746 to 750, 959 to 963 |
| | >Os05t0519400-03 | 4 | LOC_Os05g44310 | N-ethylmaleimide sensitive factor NSF (Fragment). vesicle-fusing ATPase | 46 to 50, 194 to 198, 881 to 885, 888 to 892 |
| | >Os05t0533100-01 | 4 | LOC_Os05g45660 | Plasminogen activator inhibitor 1 RNA-binding protein | 93 to 97, 145 to 149, 272 to 276, 298 to 302 |
| | >Os05t0533600-01 | 4 | LOC_Os05g45720 | Starch synthase IVa (Glycogen (Starch) synthase-like). starch synthase | 60 to 64, 226 to 230, 255 to 259, 299 to 303 |
| | >Os05t0579100-01 | 4 | LOC_Os05g50290 | HhH-GPD domain domain, hhH-GPD superfamily base excision DNA repair protein | 396 to 400, 486 to 490, 494 to 498, 769 to 773 |

| | | | | | |
|---|---|---|---|---|---|
| | >Os05t0582600-01 | 5 | LOC_Os05g50580 | Peptidase S10, serine carboxypeptidase family protein. OsSCP30 - Putative Serine Carboxypeptidase homologue | 586 to 590, 611 to 615, 625 to 629, 767 to 771, 812 to 816 |
| **TTGACA** | >Os05t0468200-01 | 4 | LOC_Os05g39190 | Domain of unknown function DUF1618 domain | 341 to 346, 431 to 436, 582 to 587, 823 to 828 |
| **TTGACTT** | >Os05t0155100-01 | 4 | LOC_Os05g06310 | 60S ribosomal protein L18-3 | 197 to 202, 364 to 369, 446 to 451, 546 to 551 |
| | >Os05t0241100-01 | 4 | LOC_Os05g15150 | Leucyl-tRNA synthetase, cytoplasmic | 30 to 35, 167 to 172, 284 to 289, 402 to 407 |
| | >Os05t0346901-00 | 6 | LOC_Os05g28010 | Non-protein coding transcript. TATA-binding protein-associated factor TAFII55 family protein | 148 to 153, 239 to 244, 271 to 276, 495 to 500, 654 to 659, 884 to 889 |
| **Chromosome 6** | | | | | |
| **Motif** | **ID** | **Frequency** | **Locus ID** | **Protein name** | **Positions in 1k promoter region** |
| **CTGACC** | >Os06t0597250-00 | 4 | LOC_Os06g39624 | B protein | 715 to 720, 852 to 857, 884 to 889, 928 to 933 |
| **TTTTCCAC** | >Os01t0730100-01 | 6 | LOC_Os01g52970 | Cyclin-like F-box domain | 112 to 197, 5 to 82, 277 to 284, 362 to 369, 425 to 432, 627 to 634 |
| **TTTGACT** | >Os01t0633000-01 | 4 | LOC_Os01g44210 | 50S ribosomal protein L31 | 408 to 414, 429 to 435, 520 to 526, 587 to 593 |
| **GTACGTAC** | >Os06t0107800-01 | 4 | LOC_Os06g01860 | RAV-like protein | 501 to 508, 505 to 512, 509 to 516, 513 to 520 |
| **TGACC** | >Os06t0136600-01 | 4 | LOC_Os06g04510 | Enolase 1 | 119 to 123, 148 to 152, 173 to 177, 202 to 206 |
| | >Os06t0172800-00 | 4 | LOC_Os06g07600 | Alkaline alpha galactosidase 2. uncharacterized glycosyltransferase | 10 to 14, 521 to 525, 606 to 610, 976 to 980 |
| | >Os06t0231350-00 | 4 | LOC_Os06g13470 | SAM dependent carboxyl methyltransferase domain. SAM dependent carboxyl methyltransferase | 176 to 180, 321 to 325, 747 to 751, 980 to 984 |
| | >Os06t0312950-00 | 4 | LOC_Os06g22600 | UPF0005 domain, aluminum-activated malate transporter | 545 to 549, 575 to 579, 603 to 607, 635 to 639 |
| | >Os06t0332450-00 | 4 | LOC_Os06g22700 | Zn-finger, RanBP-type. zinc finger family protein, | 31 to 35, 55 to 59, 415 to 419, 421 to 425 |

| | | | | | |
|---|---|---|---|---|---|
| | >Os06t0531600-01 | 4 | LOC_Os06g34070 | Lipase, GDSL domain | 379 to 383, 712 to 716, 723 to 727, 943 to 947 |
| | >Os06t0542200-01 | 4 | LOC_Os06g35050 | Prephenate dehydrogenase domain. arogenate dehydrogenase 1, chloroplast precursor | 750 to 754, 907 to 911, 916 to 920, 970 to 974 |
| | >Os06t0561800-00 | 4 | LOC_Os06g36650 | ABC transporter, transmembrane domain domain. ABC transporter family protein | 190 to 194, 247 to 251, 270 to 274, 293 to 297 |
| | >Os06t0594100-01 | 4 | LOC_Os06g39344 | Crotonase, core domain. enoyl-CoA hydratase/ isomerase family protein | 294 to 298, 352 to 356, 384 to 388, 456 to 460 |
| | >Os06t0594100-02 | 4 | LOC_Os06g39344 | Crotonase, core domain. enoyl-CoA hydratase/ isomerase family protein | 234 to 238, 292 to 296, 324 to 328, 396 to 400 |
| | >Os06t0597250-00 | 4 | LOC_Os06g39624 | B protein. transposon protein, Pong sub-class | 716 to 720, 853 to 857, 885 to 889, 929 to 933 |
| | >Os06t0691200-01 | 4 | LOC_Os06g47600 | Thaumatin-like protein precursor | 88 to 92, 163 to 167, 639 to 643, 749 to 753 |
| **TGACT** | >Os05t0125500-01 | 4 | LOC_Os05g03480 | Isovaleryl-CoA dehydrogenase, mitochondrial precursor | 61 to 65, 163 to 167, 476 to 480, 672 to 676 |
| | >Os05t0128000-00 | 5 | LOC_Os05g03740 | Homeodomain-like. transcription factor TF2 | 165 to 169, 506 to 510, 557 to 561, 782 to 786, 981 to 985 |
| | >Os05t0135700-01 | 4 | LOC_Os05g04510 | S-adenosylmethionine synthetase | 49 to 53, 148 to 152, 282 to 286, 296 to 300 |
| | >Os05t0155100-01 | 4 | LOC_Os05g06310 | 60S ribosomal protein L18. | 198 to 202, 365 to 369, 447 to 451, 547 to 551 |
| | >Os05t0157200-01 | 4 | LOC_Os05g06500 | UspA domain containing protein. universal stress protein domain | 20 to 24, 135 to 139, 376 to 380, 730 to 734 |
| | >Os05t0199800-01 | 4 | LOC_Os05g11070 | BHLH transcription activator Ivory seed. helix-loop-helix DNA-binding domain | 245 to 249, 339 to 343, 509 to 513, 677 to 681 |
| | >Os05t0241100-01 | 4 | LOC_Os05g15150 | Leucyl-tRNA synthetase, cytoplasmic | 31 to 35, 168 to 172, 285 to 289, 403 to 407 |

| | | | | | |
|---|---|---|---|---|---|
| | >Os05t0244900-00 | 4 | LOC_Os05g15600 | Transcription elongation factor, TFIIS/CRSP70, N-terminal, sub-type domain | 482 to 486, 613 to 617, 633 to 637, 684 to 688 |
| | >Os05t0346901-00 | 7 | LOC_Os05g28010 | Non-protein coding transcript. TATA-binding protein-associated factor TAFII55 family protein | 149 to 153, 240 to 244, 272 to 276, 496 to 500, 655 to 659, 800 to 804, 885 to 889 |
| | >Os05t0349200-01 | 5 | LOC_Os05g28180 | AMP deaminase 1 AMP deaminase | 291 to 295, 368 to 372, 501 to 505, 751 to 755, 855 to 859 |
| | >Os05t0356700-01 | 4 | LOC_Os05g28830 | Protein of unknown function DUF231, plant. PMR5 | 144 to 148, 699 to 703,818 to 822, 870 to 874 |
| | >Os05t0363566-00 | 4 | LOC_Os05g30010 | WD domain, G-beta repeat domain | 545 to 549, 567 to 571, 649 to 653, 720 to 724 |
| | >Os05t0371600-01 | 4 | LOC_Os05g30820 | BRASSINOSTEROID INSENSITIVE 1-associated receptor kinase 1 | 36 to 40, 83 to 87, 159 to 163, 397 to 401 |
| | >Os05t0378800-03 | 4 | LOC_Os05g31480 | Ankyrin like protein | 579 to 583, 647 to 651, 657 to 661, 864 to 868 |
| | >Os05t0436100-00 | 4 | LOC_Os05g36050 | Serine/threonine-protein kinase | 202 to 206, 528 to 532, 730 to 734, 917 to 921 |
| | >Os05t0458500-00 | 4 | LOC_Os05g38410 | Laccase (EC 1.10.3.2). laccase precursor protein | 85 to 89, 238 to 242, 318 to 322, 556 to 560 |
| | >Os05t0468400-01 | 5 | LOC_Os05g39210 | Domain of unknown function DUF1618 domain | 136 to 140, 319 to 323, 529 to 533, 856 to 860, 895 to 899 |
| | >Os05t0480000-01 | 4 | LOC_Os05g40180 | Serine/threonine protein kinase-related domain | 11 to 15, 279 to 283, 348 to 352, 622 to 626 |
| | >Os05t0486700-01 | 4 | LOC_Os05g40820 | Ribosomal protein L24e domain | 219 to 223, 272 to 276, 330 to 334, 643 to 647 |
| | >Os05t0493100-01 | 5 | LOC_Os05g41370 | KI domain interacting kinase 1, TKL_IRAK_DUF26-la.1 - DUF26 kinases | 272 to 276, 303 to 307, 471 to 475, 555 to 559, 908 to 912 |
| | >Os05t0497700-01 | 4 | LOC_Os05g41810 | Peptidase. OsDegp8 - Putative Deg protease homologue | 171 to 175, 316 to 320, 419 to 423, 782 to 786 |
| | >Os05t0500400-01 | 4 | LOC_Os05g42100 | Domain of unknown function DUF1995 domain | 207 to 211, 393 to 397, 721 to 725, 749 to 753 |

| | | | | | |
|---|---|---|---|---|---|
| | >Os05t0519700-01 | 4 | LOC_Os05g44340 | Heat shock protein 101 | 702 to 706, 766 to 770, 771 to 775, 801 to 805 |
| | >Os05t0519700-02 | 4 | LOC_Os05g44340 | Heat shock protein 101 | 648 to 652, 712 to 716, 717 to 721, 747 to 751 |
| | >Os05t0527100-00 | 4 | LOC_Os05g45110 | UDP-glucuronosyl/UDP-glucosyltransferase family protein. anthocyanidin 5,3-O-glucosyltransferase | 277 to 281, 318 to 322, 696 to 700, 779 to 783 |
| | >Os05t0555900-01 | 4 | LOC_Os05g48220 | 60S ribosomal protein L35a-3 | 504 to 508, 656 to 660, 707 to 711, 796 to 800 |
| | >Os05t0556000-01 | 4 | LOC_Os05g48230 | Oxoglutarate/iron-dependent oxygenase domain | 29 to 33, 118 to 122, 517 to 521, 789 to 793 |
| | >Os05t0582600-01 | 4 | LOC_Os05g50580 | Peptidase S10, serine carboxypeptidase family protein. OsSCP30 - Putative Serine Carboxypeptidase homologue | 51 to 55, 559 to 563, 768 to 772, 813 to 817 |
| **TTGAC** | >Os06t0102700-01 | 4 | LOC_Os06g01304 | Predicted protein. spotted leaf 11 | 278 to 282, 525 to 529, 711 to 715, 970 to 974 |
| | >Os06t0103200-01 | 4 | LOC_Os06g01350 | Transferase family protein. transferase family protein | 323 to 327, 772 to 776, 802 to 806, 864 to 868 |
| | >Os06t0104800-00 | 4 | LOC_Os06g01580 | Proline-rich protein. LTPL127 - Protease inhibitor/seed storage/LTP family protein precursor | 139 to 143, 175 to 179, 292 to 296, 629 to 633 |
| | >Os06t0115300-01 | 4 | LOC_Os06g02490 | Acyl-CoA-binding protein 2 (ACBP 2) acyl CoA binding protein | 129 to 133, 331 to 335, 350 to 354, 736 to 740 |
| | >Os06t0120200-01 | 4 | LOC_Os06g02960 | Protein of unknown function DUF594 family protein. | 171 to 175, 186 to 190, 195 to 199, 432 to 436 |
| | >Os06t0128700-01 | 4 | LOC_Os06g03800 | Ankyrin repeat domain | 269 to 273, 378 to 382, 526 to 530, 681 to 685 |
| | >Os06t0136500-01 | 7 | LOC_Os06g04500 | Cornichon family protein | 69 to 73, 261 to 265, 303 to 307, 414 to 418, 671 to 675, 680 to 684, 699 to 703 |
| | >Os06t0137600-01 | 5 | LOC_Os06g04610 | K homology-like, alpha/beta domain. ribosome-binding factor A, chloroplast precursor | 204 to 208, 295 to 299, 304 to 308, 323 to 327, 735 to 739 |

| | | | | | |
|---|---|---|---|---|---|
| | >Os06t0137600-02 | 5 | LOC_Os06g04610 | K homology-like, alpha/beta domain. ribosome-binding factor A, chloroplast precursor | 202 to 206, 293 to 297, 302 to 306, 321 to 325, 733 to 737 |
| | >Os06t0140200-01 | 7 | LOC_Os06g04830 | Leucine-rich repeat, plant specific containing protein | 135 to 139, 221 to 225, 240 to 244, 622 to 626, 653 to 657, 671 to 675, 682 to 686 |
| | >Os06t0142400-01 | 4 | LOC_Os06g05020 | Early nodulin. early nodulin 93 ENOD93 protein | 455 to 459, 512 to 516, 521 to 525, 540 to 544 |
| | >Os06t0179200-01 | 5 | LOC_Os06g08110 | Nodulin-like protein. nodulin | 476 to 480, 516 to 520, 654 to 658, 663 to 667, 682 to 686 |
| | >Os06t0188800-01 | 7 | LOC_Os06g08910 | Extracellular solute-binding protein, family 3 domain. glutamate receptor 2.8 precursor | 494 to 498, 523 to 527, 563 to 567, 745 to 749, 754 to 758, 774 to 778, 878 to 882 |
| | >Os06t0188800-02 | 5 | LOC_Os06g08910 | Extracellular solute-binding protein, family 3 domain. glutamate receptor 2.8 precursor | 191 to 195, 361 to 365, 762 to 766, 911 to 915, 918 to 922 |
| | >Os06t0193000-02 | 4 | LOC_Os06g09330 | Ubiquitin conjugating enzyme 2 | 245 to 249, 301 to 305, 329 to 333, 524 to 528 |
| | >Os06t0208151-00 | 8 | | Quinone oxidoreductase | 163 to 167, 266 to 270, 444 to 448, 545 to 549, 605 to 609, 693 to 697, 727 to 731, 752 to 756 |
| | >Os06t0210400-01 | 6 | LOC_Os06g10790 | Concanavalin A-like lectin/glucanase, subgroup domain containing protein. lectin-like receptor kinase | 32 to 36, 229 to 233, 561 to 565, 625 to 629, 667 to 671, 766 to 770 |
| | >Os06t0225900-01 | 4 | LOC_Os06g12160 | ATP binding / ATPase/ nucleoside-triphosphatase/ nucleotide binding | 20 to 24, 79 to 83, 528 to 532, 674 to 678 |
| | >Os06t0226700-01 | 4 | LOC_Os06g12230 | PCF1. TCP-domain protein | 17 to 21, 43 to 47, 228 to 232, 408 to 412 |
| | >Os06t0230100-00 | 5 | LOC_Os06g12460 | Cellulose synthase-like A1. CSLA3 - cellulose synthase-like family A; mannan synthase | 16 to 20, 107 to 111, 189 to 193, 828 to 832, 981 to 985 |

| | | | | | |
|---|---|---|---|---|---|
| | >Os06t0242900-01 | 4 | LOC_Os06g13470) | SAM dependent carboxyl methyltransferase domain | 198 to 202, 531 to 535, 548 to 552, 559 to 563 |
| | >Os06t0264700-01 | 4 | LOC_Os06g15390 | Acylphosphatase domain | 42 to 46, 250 to 254, 340 to 344, 816 to 820 |
| | >Os06t0266266-00 | 4 | LOC_Os06g15590 | Cell number regulator 11. uncharacterized Cys-rich domain | 32 to 36, 469 to 473, 607 to 611, 996 to 1000 |
| | >Os06t0295000-01 | 4 | LOC_Os06g19110 | Cadmium tolerance factor | 4 to 8, 258 to 262, 410 to 414, 499 to 503 |
| | >Os06t0296900-00 | 4 | LOC_Os06g19300 | Cadmium tolerance factor | 177 to 181, 198 to 202, 680 to 684, 847 to 851 |
| | >Os06t0301100-01 | 4 | LOC_Os06g19690 | OSIGBa0124N08.6 protein | 407 to 411, 496 to 500, 610 to 614, 695 to 699 |
| | >Os06t0308900-01 | 7 | LOC_Os06g20400 | DHHC zinc finger domain | 18 to 22, 108 to 112, 167 to 171, 194 to 198, 552 to 556, 561 to 565, 580 to 584 |
| | >Os06t0313440-00 | 7 | LOC_Os06g20790 | SAM dependent carboxyl methyltransferase domain | 69 to 73, 279 to 283, 317 to 321, 463 to 467, 501 to 505, 633 to 637, 671 to 675 |
| | >Os06t0320300-01 | 4 | LOC_Os06g21580 | HMG-I and HMG-Y, DNA-binding domain | 102 to 106, 110 to 114, 119 to 123, 290 to 294 |
| | >Os06t0331900-01 | 5 | LOC_Os06g22600 | Aluminum-activated malate transporter | 122 to 126, 130 to 134, 639 to 643, 740 to 744, 751 to 755 |
| | >Os06t0332900-01 | 4 | LOC_Os06g22700 | Zn-finger, RanBP-type. zinc finger family protein | 6 to 10, 46 to 50, 652 to 656, 680 to 684 |
| | >Os06t0335950-00 | 5 | LOC_Os06g22919 | DEFL9 - Defensin and Defensin-like DEFL family | 27 to 31, 77 to 81, 123 to 127, 351 to 355, 713 to 717 |
| | >Os06t0484450-00 | 4 | LOC_Os06g28960 | Chlorophyll a-b binding protein 2 | 555 to 559, 586 to 590, 666 to 670, 725 to 729 |
| | >Os06t0493700-01 | 4 | | Pi homeostasis | 31 to 35, 360 to 364, 451 to 455, 796 to 800 |
| | >Os06t0500700-01 | 4 | LOC_Os06g30500 | Cytochrome P450 family protein | 121 to 125, 347 to 351, 636 to 640, 645 to 649 |
| | >Os06t0536100-00 | 4 | LOC_Os06g34530 | zinc finger, C3HC4 type domain | 108 to 112, 225 to 229, 507 to 511, 915 to 919 |
| | >Os06t0543400-01 | 4 | LOC_Os06g35160 | CAMK_KIN1/SNF1/Nim1_like.26 - CAMK includes calcium/calmodulin depedent protein kinases | 270 to 274, 279 to 283, 298 to 302, 889 to 893 |

| | | | | | |
|---|---|---|---|---|---|
| | >Os06t0549600-01 | 5 | LOC_Os06g35650 | FAD-linked oxidase, FAD-binding, subdomain 2 domain. Reticuline oxidase-like protein precursor | 11 to 15, 226 to 230, 246 to 250, 513 to 517, 841 to 845 |
| | >Os06t0552400-01 | 4 | LOC_Os06g35910 | Zinc ion binding protein. FYVE zinc finger domain | 46 to 50, 296 to 300, 505 to 509, 632 to 636 |
| | >Os06t0560000-01 | 4 | LOC_Os06g36450 | Ferroportin1 family protein | 63 to 67, 101 to 105, 482 to 486, 503 to 507 |
| | >Os06t0561800-00 | 4 | LOC_Os06g36650 | ABC transporter, transmembrane domain | 99 to 103, 189 to 193, 246 to 250, 269 to 273 |
| | >Os06t0594100-01 | 5 | LOC_Os06g39344 | Crotonase, core domain. enoyl-CoA hydratase/isomerase family protein | 231 to 235, 293 to 297, 351 to 355, 455 to 459, 837 to 841 |
| | >Os06t0594100-02 | 5 | LOC_Os06g39344 | Crotonase, core domain. enoyl-CoA hydratase/isomerase family protein | 171 to 175, 233 to 237, 291 to 295, 395 to 399, 777 to 781 |
| | >Os06t0602500-01 | 4 | LOC_Os06g40030 | Serine/threonine protein kinase-related domain containing protein. S-locus-like receptor protein kinase | 122 to 126, 453 to 457, 761 to 765, 867 to 871 |
| | >Os06t0605750-00 | 4 | LOC_Os06g40330 | MYB family transcription factor | 152 to 156, 241 to 245, 338 to 342, 977 to 981 |
| | >Os06t0613400-01 | 4 | LOC_Os06g41050 | DNA repair protein Sae2/CtIP domain | 24 to 28, 334 to 338, 412 to 416, 475 to 479 |
| | >Os06t0626700-00 | 4 | LOC_Os06g42130 | Isopenicillin N synthase family protein. leucoanthocyanidin dioxygenase | 82 to 86, 91 to 95, 110 to 114, 538 to 542 |
| | >Os06t0632400-00 | 4 | LOC_Os06g42620 | OSIGBa0115M15.3 protein | 291 to 295, 300 to 304, 319 to 323, 948 to 952 |
| | >Os06t0633300-01 | 5 | LOC_Os06g42680 | Phytosulfokines 1 | 12 to 16, 19 to 23, 389 to 393, 568 to 572, 732 to 736 |
| | >Os06t0642600-00 | 4 | LOC_Os06g43520 | Cytochrome P450 CYP71Y10 | 207 to 211, 353 to 357, 485 to 489, 650 to 654 |

| | | | | | |
|---|---|---|---|---|---|
| | >Os06t0646400-01 | 4 | LOC_Os06g43840 | Tyrosine protein kinase domain. protein kinase domain | 380 to 384, 396 to 400, 431 to 435, 584 to 588 |
| | >Os06t0676000-01 | 4 | LOC_Os06g46310 | Integral membrane protein OsNramp3 metal transporter Nramp6 | 127 to 131, 396 to 400, 532 to 536, 752 to 756 |
| | >Os06t0690900-00 | 5 | LOC_Os06g47570 | Pentatricopeptide repeat domain. PPR repeat | 100 to 104, 249 to 253, 532 to 536, 541 to 545, 560 to 564 |
| | >Os06t0692600-01 | 5 | LOC_Os06g47750 | Protein kinase, core domain. phytosulfokine receptor precursor | 4 to 8, 433 to 437, 660 to 664, 876 to 880, 978 to 982 |
| | >Os06t0692700-01 | 4 | LOC_Os06g47760 | Phytosulfokine receptor precursor | 244 to 248 , 629 to 633, 934 to 938, 954 to 958 |
| | >Os06t0699700-01 | 4 | LOC_Os06g48620 | Aminodeoxychorismate synthase/glutamine amidotransferase. 4-amino-4-deoxychorismate synthase | 35 to 39 , 258 to 262, 520 to 524, 738 to 742 |
| | >Os06t0705700-01 | 4 | LOC_Os06g49220 | TGF-beta receptor, type I/II extracellular region family protein. peptide transporter | 438 to 442, 478 to 482, 627 to 631, 814 to 818 |
| | >Os06t0714900-01 | 4 | LOC_Os06g50100 | Serine/threonine protein kinase domain. tyrosine protein kinase domain | 523 to 527, 558 to 562, 707 to 711, 893 to 897 |
| | >Os06t0714900-02 | 4 | LOC_Os06g50100 | Serine/threonine protein kinase domain | 449 to 453, 484 to 488, 633 to 637, 819 to 823 |
| | >Os06t0715000-01 | 4 | LOC_Os06g50110 | UV radiation resistance protein/autophagy-related protein 14 domain | 171 to 175 , 361 to 365, 778 to 782, 792 to 796 |
| | >Os06t0716200-01 | 4 | LOC_Os06g50240 | Thaumatin, pathogenesis-related family protein | 585 to 589, 621 to 625, 790 to 794, 852 to 856 |
| **TTGACA** | >Os06t0188800-02 | 4 | LOC_Os06g08910 | Extracellular solute-binding protein, family 3 domain. glutamate receptor 2.8 | 191 to 196, 361 to 366, 911 to 916, 918 to 923 |
| | >Os06t0230100-00 | 4 | LOC_Os06g12460 | Cellulose synthase-like A3 | 368 to 373, 425 to 430, 486 to 491, 926 to 931 |
| | >Os06t0230100-01 | 4 | LOC_Os06g12460 | Cellulose synthase-like A3 | 16 to 21, 107 to 112, 189 to 194, 981 to 986 |

| | | | | | |
|---|---|---|---|---|---|
| | >Os06t0552400-01 | 4 | LOC_Os06g35910 | Zinc ion binding protein. FYVE zinc finger domain | 46 to 51, 296 to 301, 505 to 510, 632 to 637 |
| **TTGACC** | >Os06t0332900-01 | 4 | LOC_Os06g22700 | zinc finger, RanBP-type | 6 to 11, 46 to 51, 652 to 657, 680 to 685 |
| **TTGACT** | >Os06t0208151-00 | 4 | | Quinone oxidoreductase | 163 to 168, 605 to 610, 693 to 698, 752 to 757 |
| | >Os06t0308900-01 | 4 | LOC_Os06g20400 | DHHC zinc finger domain | 18 to 23, 108 to 113, 167 to 172, 561 to 566 |
| **Chromosome 7** | | | | | |
| **Motif** | **ID** | **Frequency** | **Locus ID** | **Protein name** | **Positions in 1k promoter region** |
| **TGACC** | >Os07t0130800-01 | 4 | LOC_Os07g03870 | Protein kinase, catalytic domain domain. receptor like protein kinase | 37 to 41, 182 to 186, 719 to 723, 853 to 857 |
| | >Os07t0134500-01 | 4 | LOC_Os07g04210 | Hydrolase/ protein serine/threonine phosphatase. Ser/Thr protein phosphatase family protein | 184 to 188, 247 to 251, 495 to 499, 540 to 544 |
| | >Os07t0137000-01 | 4 | LOC_Os07g04430 | Myb transcription factor | 122 to 126, 477 to 481, 628 to 632, 849 to 853 |
| | >Os07t0160500-01 | 4 | LOC_Os07g06670 | F-box domain, Skp2-like domain. OsFBL35 - F-box domain and LRR | 303 to 307, 494 to 498, 918 to 922, 988 to 992 |
| | >Os07t0237100-00 | 4 | LOC_Os07g13280 | RNA recognition motif domain | 144 to 148, 177 to 181, 317 to 321, 730 to 734 |
| | >Os07t0250900-01 | 4 | LOC_Os07g14700 | Harpin-induced 1 domain | 71 to 75, 79 to 83, 264 to 268, 821 to 825 |
| | >Os07t0434700-02 | 4 | LOC_Os07g25410 | Peptidase M24, methionine aminopeptidase family protein. peptidase, M24 family protein | 18 to 22, 358 to 362, 511 to 515, 734 to 738 |
| | >Os07t0443700-00 | 4 | LOC_Os07g26170 | Serine-threonine/tyrosine-protein kinase domain | 571 to 575, 695 to 699, 781 to 785, 882 to 886 |

| | | | | | |
|---|---|---|---|---|---|
| | >Os07t0447200-01 | 4 | LOC_Os07g26595 | OSIGBa0102I15.5 protein | 499 to 503, 534 to 538, 546 to 550, 663 to 667 |
| | >Os07t0451300-01 | 4 | LOC_Os07g26870 | Cytochrome P450 family protein | 74 to 78, 164 to 168, 184 to 188, 195 to 199 |
| | >Os07t0490600-02 | 4 | LOC_Os07g30820 | Type II membrane protein. zinc finger family protein | 73 to 77, 230 to 234, 241 to 245, 264 to 268 |
| | >Os07t0499500-01 | 4 | LOC_Os07g31610 | Peroxidase 7 precursor | 276 to 280, 498 to 502, 597 to 601, 702 to 706 |
| | >Os07t0564800-01 | 4 | LOC_Os07g37760 | Protein of unknown function DUF707 family protein. lysine ketoglutarate reductase trans-splicing related 1 | 239 to 243, 582 to 586, 830 to 834, 970 to 974 |
| | >Os07t0613300-01 | 4 | LOC_Os07g42180 | PAUSED. exportin 1 | 55 to 59, 341 to 345, 397 to 401, 468 to 472 |
| | >Os07t0639100-00 | 4 | LOC_Os07g44560 | AMP-dependent synthetase/ligase domain | 714 to 718, 724 to 728, 772 to 776, 782 to 786 |
| | >Os07t0650100-00 | 4 | LOC_Os07g45550 | Armadillo-like helical domain | 170 to 174, 192 to 196, 244 to 248, 733 to 737 |
| **TGACT** | >Os07t0108100-01 | 5 | LOC_Os07g01740 | OSIGBa0147O06.5 protein. cysteine protease | 319 to 323, 323 to 327, 327 to 331, 385 to 389, 890 to 894 |
| | >Os07t0120100-01 | 4 | LOC_Os07g02880 | Protein of unknown function DUF538 family protein | 223 to 227, 337 to 341, 345 to 349, 463 to 467 |
| | >Os07t0122100-01 | 5 | LOC_Os07g03040 | Protein of unknown function DUF1719, Oryza sativa family protein | 236 to 240, 392 to 396, 396 to 400, 411 to 415, 415 to 419 |
| | >Os07t0131375-00 | 4 | LOC_Os07g03920 | Protein kinase, catalytic domain domain. lectin-like receptor kinase 7 | 102 to 106, 566 to 570, 813 to 817, 912 to 916 |
| | >Os07t0133100-01 | 4 | LOC_Os07g04130 | Concanavalin A-like lectin/glucanase, subgroup domain | 251 to 255, 755 to 759, 907 to 911, 911 to 915 |
| | >Os07t0145800-00 | 4 | LOC_Os07g05210 | dof zinc finger protein 2 | 37 to 41, 148 to 152, 470 to 474, 479 to 483 |
| | >Os07t0150700-01 | 4 | LOC_Os07g05620 | Serine/threonine protein kinase, Pollination and drought stress responses CAMK_KIN1 /SNF1/Nim1_like.28 - CAMK includes calcium/calmodulin depedent protein kinases | 204 to 208, 307 to 311, 339 to 343, 437 to 441 |

| | | | | | |
|---|---|---|---|---|---|
| | >Os07t0172900-01 | 4 | LOC_Os07g07646 | Biopterin transport-related protein BT1 protein | 181 to 185, 301 to 305, 383 to 387, 785 to 789 |
| | >Os07t0172900-02 | 4 | LOC_Os07g07646 | Biopterin transport-related protein BT1 protein | 107 to 111, 189 to 193, 591 to 595, 845 to 849 |
| | >Os07t0244800-00 | 4 | LOC_Os07g14130 | Pectinesterase inhibitor domain | 663 to 667, 685 to 689, 767 to 771, 838 to 842 |
| | >Os07t0249800-01 | 4 | LOC_Os07g14600 | IAA-amino acid hydrolase ILR1-like 8. hydrolase | 81 to 85, 295 to 299, 851 to 855, 874 to 878 |
| | >Os07t0280200-01 | 4 | LOC_Os07g17970 | AMP-dependent synthetase and ligase domain | 422 to 426, 524 to 528, 608 to 612, 831 to 835 |
| | >Os07t0462000-01 | 4 | LOC_Os07g27790 | Glutamate--cysteine ligase, GCS2 family protein, chloroplast precursor | 84 to 88, 439 to 443, 677 to 681, 872 to 876 |
| | >Os07t0519450-00 | 5 | | MYB transcription factor | 47 to 51, 539 to 543, 659 to 663, 674 to 678, 751 to 755 |
| | >Os07t0549700-01 | 4 | LOC_Os07g36465 | Armadillo-like helical domain. vacuolar ATP synthase subunit H | 238 to 242, 257 to 261, 347 to 351, 355 to 359 |
| | >Os07t0570550-00 | 4 | LOC_Os07g38290 | Cupredoxin domain, plastocyanin-like domain | 123 to 127, 550 to 554, 918 to 922, 922 to 926 |
| | >Os07t0599100-00 | 5 | LOC_Os07g40810 | NBS-LRR type disease resistance protein | 166 to 170, 346 to 350, 617 to 621, 864 to 868, 910 to 914 |
| | >Os07t0599500-01 | 4 | LOC_Os07g40850 | Retrotransposon protein | 228 to 232, 282 to 286, 822 to 826, 826 to 830 |
| | >Os07t0601900-01 | 4 | LOC_Os07g41060 | NADPH HC toxin reductase dihydroflavonol-4-reductase | 82 to 86, 241 to 245, 335 to 339, 504 to 508 |
| | >Os07t0620500-01 | 5 | | Oligosaccaryltransferase domain | 298 to 302, 498 to 502, 723 to 727, 769 to 773, 933 to 937 |
| | >Os07t0658300-02 | 4 | LOC_Os07g46450 | RhoGAP domain. pleckstrin homology domain | 178 to 182, 279 to 283, 659 to 663, 831 to 835 |
| | >Os07t0676900-01 | 4 | LOC_Os07g47990 | Peroxidase | 34 to 38, 49 to 53, 102 to 106, 118 to 122 |
| | >Os07t0677100-01 | 4 | LOC_Os07g48010 | Peroxidase precursor | 68 to 72, 94 to 98, 225 to 229, 859 to 863 |
| **TTGAC** | >Os07t0103000-01 | 4 | LOC_Os07g01300 | GRF zinc finger family protein | 498 to 502, 508 to 512, 720 to 724, 860 to 864 |

| | | | | | |
|---|---|---|---|---|---|
| | >Os07t0119000-01 | 4 | LOC_Os07g02780 | MAP3K gamma protein kinase (Fragment).STE_MEKK_ste 11_MAP3K.20 - STE kinases include homologs to sterile 7, sterile 11 and sterile 20 from yeast | 211 to 215, 218 to 222, 296 to 300, 651 to 655 |
| | >Os07t0120100-01 | 4 | LOC_Os07g02880 | Protein of unknown function DUF538 family protein. DUF538 domain | 87 to 91, 222 to 226, 336 to 340, 462 to 466 |
| | >Os07t0120900-01 | 4 | LOC_Os07g02980 | Protein of unknown function DUF1719, Oryza sativa family protein | 471 to 475, 478 to 482, 689 to 693, 755 to 759 |
| | >Os07t0122100-01 | 4 | LOC_Os07g03040 | Protein of unknown function DUF1719, Oryza sativa family protein | 73 to 77, 410 to 414, 755 to 759, 808 to 812 |
| | >Os07t0127700-01 | 4 | LOC_Os07g03600 | PR1b | 51 to 55, 268 to 272, 277 to 281, 296 to 300 |
| | >Os07t0155100-01 | 4 | LOC_Os07g06080 | FAD dependent oxidoreductase family protein | 304 to 308, 563 to 567, 659 to 663, 968 to 972 |
| | >Os07t0188800-01 | 4 | LOC_Os07g09060 | Methylmalonate-semialdehyde dehydrogenase | 458 to 462, 764 to 768, 805 to 809, 812 to 816 |
| | >Os07t0244800-00 | 4 | LOC_Os07g14130 | Pectinesterase inhibitor domain | 33 to 37, 93 to 97, 198 to 202, 394 to 398 |
| | >Os07t0249800-01 | 4 | LOC_Os07g14600 | IAA-amino acid hydrolase ILR1-like 8. hydrolase | 32 to 36, 158 to 162, 294 to 298, 873 to 877 |
| | >Os07t0250900-01 | 4 | LOC_Os07g14700 | Harpin-induced 1 domain | 70 to 74, 78 to 82, 835 to 839, 931 to 935 |
| | >Os07t0271600-00 | 4 | LOC_Os07g17040 | MAK16 protein-related | 63 to 67, 111 to 115, 141 to 145, 394 to 398 |
| | >Os07t0290500-00 | 4 | LOC_Os07g19000 | Plant lipid transfer/seed storage/trypsin-alpha amylase inhibitor domain. LTPL41 - Protease inhibitor/seed storage/LTP family protein precursor | 308 to 312, 426 to 430, 632 to 636, 818 to 822 |

| | | | | | |
|---|---|---|---|---|---|
| | >Os07t0290800-01 | 6 | LOC_Os07g19030 | Tic22-like family protein. tic22-like family domain | 123 to 127, 205 to 209, 447 to 451, 478 to 482, 526 to 530, 553 to 557 |
| | >Os07t0407900-01 | 4 | LOC_Os07g22510 | Protein of unknown function DUF679 family protein | 436 to 440, 593 to 597, 673 to 677, 890 to 894 |
| | >Os07t0414700-00 | 4 | LOC_Os07g23190 | Protein of unknown function DUF707 domain. lysine ketoglutarate reductase trans-splicing related 1 | 53 to 57, 162 to 166, 701 to 705, 869 to 873 |
| | >Os07t0415200-01 | 4 | LOC_Os07g23244 | Ribosomal protein L25/L23 domain | 156 to 160, 359 to 363, 451 to 455, 596 to 600 |
| | >Os07t0443500-00 | 5 | LOC_Os07g26150 | Molecular chaperone, heat shock protein, Hsp40 | 224 to 228, 245 to 249, 328 to 332, 713 to 717, 749 to 753 |
| | >Os07t0447200-01 | 6 | LOC_Os07g26595 | OSIGBa0102I15.5 protein | 179 to 183, 498 to 502, 533 to 537, 545 to 549, 594 to 598, 662 to 666 |
| | >Os07t0451300-01 | 4 | LOC_Os07g26870 | Cytochrome P450 family protein. | 152 to 156, 163 to 167, 183 to 187, 194 to 198 |
| | >Os07t0476401-00 | 4 | LOC_Os07g29380 | K-exchanger-like protein | 249 to 253, 265 to 269, 276 to 280, 915 to 919 |
| | >Os07t0476401-00 | 5 | LOC_Os07g29380 | K-exchanger-like protein | 249 to 253, 265 to 269, 276 to 280, 915 to 919 |
| | >Os07t0495900-04 | 4 | LOC_Os07g31340 | Low-level beta-amylase 1. regulator of nonsense transcripts 1 | 14 to 18, 30 to 34, 444 to 448, 656 to 660 |
| | >Os07t0497500-00 | 4 | LOC_Os07g31470 | MYB transcription factor | 26 to 30, 86 to 90, 197 to 201, 638 to 642 |
| | >Os07t0499500-01 | 4 | LOC_Os07g31610 | Peroxidase 7 precursor | 275 to 279, 497 to 501, 701 to 705, 742 to 746 |
| | >Os07t0504200-00 | 4 | LOC_Os07g32080 | Transcription initiation factor IIB | 867 to 871, 876 to 880, 976 to 980, 987 to 991 |
| | >Os07t0519450-00 | 5 | | MYB transcription factor | 658 to 662, 719 to 723, 750 to 754, 798 to 802, 816 to 820 |

| | | | | | |
|---|---|---|---|---|---|
| | >Os07t0528900-00 | 4 | LOC_Os07g34510 | SYP132.retrotransposon protein, | 149 to 153, 162 to 166, 618 to 622, 843 to 847 |
| | >Os07t0534000-01 | 4 | LOC_Os07g34950 | Permeases of the major facilitator superfamily. uncharacterized membrane protein | 80 to 84, 112 to 116, 414 to 418, 446 to 450 |
| | >Os07t0535600-00 | 4 | LOC_Os07g35110 | F-box associated interaction domain domain. Leucine Rich Repeat family | 234 to 238, 618 to 622, 758 to 762, 768 to 772 |
| | >Os07t0564800-01 | 4 | LOC_Os07g37760 | Protein of unknown function DUF707 family protein. lysine ketoglutarate reductase trans-splicing related 1 | 294 to 298, 581 to 585, 731 to 735, 848 to 852 |
| | >Os07t0572000-01 | 4 | LOC_Os07g38430 | WD40/YVTN repeat-like domain. WD domain, G-beta repeat domain | 500 to 504, 594 to 598, 634 to 638, 897 to 901 |
| | >Os07t0573600-01 | 4 | LOC_Os07g38600 | Nucleotide excision repair, TFIIH, subunit TTDA domain containing protein. REX1 DNA Repair family | 183 to 187, 291 to 295, 299 to 303, 845 to 849 |
| | >Os07t0575000-00 | 5 | LOC_Os07g38750 | Ethylene response factor 6 | 139 to 143, 258 to 262, 303 to 307, 544 to 548, 757 to 761 |
| | >Os07t0577700-01 | 4 | LOC_Os07g38970 | Succinyl-CoA synthetase-like domain containing protein. succinyl-CoA ligase subunit alpha-2, mitochondrial precursor | 328 to 332, 509 to 513, 579 to 583, 890 to 894 |
| | >Os07t0584750-00 | 5 | LOC_Os07g39570 | OSIGBa0147H17.4 protein | 9 to 13, 130 to 134, 175 to 179, 823 to 827, 850 to 854 |
| | >Os07t0590100-01 | 4 | LOC_Os07g40080 | Zinc finger, C2H2 domain. ZOS7-09 - C2H2 zinc finger protein | 377 to 381, 540 to 544, 572 to 576, 652 to 656 |
| | >Os07t0599100-00 | 4 | LOC_Os07g40810 | NBS-LRR type disease resistance protein | 165 to 169, 197 to 201, 345 to 349, 863 to 867 |
| | >Os07t0623200-01 | 5 | LOC_Os07g43040 | ATPase, P-type, K/Mg/Cd/Cu/Zn/Na/Ca/Na/H-transporter domain. heavy metal-associated domain | 42 to 46, 416 to 420, 447 to 451, 454 to 458, 659 to 663 |

| | | | | | |
|---|---|---|---|---|---|
| | >Os07t0623200-02 | 6 | LOC_Os07g43040 | ATPase, P-type, K/Mg/Cd/Cu/Zn/Na/Ca/Na/ H-transporter domain. heavy metal-associated domain | 15 to 19, 389 to 393, 420 to 424, 427 to 431, 632 to 636, 976 to 980 |
| | >Os07t0623200-03 | 5 | LOC_Os07g43040 | ATPase, P-type, K/Mg/Cd/Cu/Zn/Na/Ca/Na/ H-transporter domain containing protein. heavy metal-associated domain | 310 to 314, 341 to 345, 348 to 352, 553 to 557, 897 to 901 |
| | >Os07t0623200-02 | 4 | LOC_Os07g43990 | SUB1; calcium ion binding | 15 to 19, 389 to 393, 420 to 424, 427 to 431, 632 to 636, 976 to 980 |
| | >Os07t0635150-01 | 4 | LOC_Os07g44200 | Protein of unknown function DUF573 family protein. transcription regulator | 365 to 369, 399 to 403, 708 to 712, 865 to 869 |
| | >Os07t0636100-01 | 4 | LOC_Os07g44840 | Bacterial transferase hexapeptide repeat domain | 198 to 202, 207 to 211, 675 to 679, 694 to 698 |
| | >Os07t0669750-02 | 4 | LOC_Os07g48010 | Peroxidase | 215 to 219, 284 to 288, 466 to 470, 531 to 535 |
| | >Os07t0681500-02 | 4 | LOC_Os07g48420 | Ankyrin repeat | 79 to 83, 174 to 178, 349 to 353, 990 to 994 |
| | >Os07t0682700-01 | 5 | | Chitin-binding lectin 1 precursor | 39 to 43, 157 to 161, 390 to 394, 421 to 425, 525 to 529 |
| | >Os07t0684500-01 | 5 | LOC_Os07g48810 | Rhodanese-like domain. parvulin-type peptidyl prolyl cis/trans isomerase | 43 to 47, 325 to 329, 487 to 491, 630 to 634, 930 to 934 |
| | >Os07t0688200-01 | 4 | | MYB transcription factor | 76 to 80, 270 to 274, 352 to 356, 748 to 752 |
| | >Os07t0692950-00 | 4 | LOC_Os07g49370 | Glycosyl transferase, family 43 protein. glycosyltransferase family 43 protein | 571 to 575, 617 to 621, 666 to 670, 993 to 997 |
| | >Os07t0694400-01 | 4 | LOC_Os07g29380 | K-exchanger-like protein. | 194 to 198, 269 to 273, 459 to 463, 478 to 482 |
| **TTGACA** | >Os07t0623200-02 | 4 | LOC_Os07g43040 | ATPase, P-type, K/Mg/Cd/Cu/Zn/Na/Ca/Na/ H-transporter domain. heavy metal-associated domain | 15 to 203, 89 to 394, 420 to 425, 976 to 981 |
| **TTGACC** | >Os07t0447200-01 | 4 | LOC_Os07g26595 | OSIGBa0102I15.5 protein | 498 to 503, 533 to 538, 545 to 550, 662 to 667 |

| Chromosome 8 | | | | | |
|---|---|---|---|---|---|
| **Motif** | **ID** | **Frequency** | **Locus ID** | **Protein name** | **Positions in 1k promoter region** |
| **TTTGACT** | >Os08t0173700-01 | 5 | LOC_Os08g07700 | Ethylene-responsive transcription factor 80 | 122 to 285, 4 to 60, 201 to 207, 380 to 386, 744 to 750 |
| | >Os08t0526500-01 | 4 | LOC_Os08g41480 | SOX-1 protein. SAM domain | 500 to 506, 532 to 538, 681 to 687, 735 to 741 |
| **TTGACG** | >Os08t0377100-00 | 4 | LOC_Os08g28940 | OsFBX289 - F-box domain | 11 to 16, 82to87, 262 to 267, 442 to 447 |
| **GTTGAC** | >Os08t0412600-01 | 4 | LOC_Os08g31840 | RabGAP/TBC domain | 262 to 267, 414 to 419, 620 to 625, 642 to 647 |
| **GTACGTAC** | >Os08t0454000-00 | 4 | LOC_Os08g35240 | Ethylene Response Factor 12 | 590 to 597, 594 to 601, 598 to 605, 602 to 609 |
| **TGACC** | >Os08t0299200-01 | 6 | LOC_Os08g20400 | Adenylate cyclase domain | 164 to 168, 249 to 253, 515 to 519, 526 to 530, 543 to 547, 798 to 802 |
| | >Os08t0378000-02 | 4 | LOC_Os08g29020 | Wall-associated kinase-like 2 | 734 to 738, 858 to 862, 877 to 881, 898 to 902 |
| | >Os08t0412600-01 | 6 | LOC_Os08g31840 | RabGAP/TBC domain | 416 to 420, 456 to 460, 515 to 519, 607 to 611, 622 to 626, 644 to 648 |
| | >Os08t0465300-01 | 5 | LOC_Os08g36250 | Sodium/hydrogen exchanger family protein. | 297 to 301, 554 to 558, 568 to 572, 616 to 620, 702 to 706 |
| **TGACT** | >Os08t0100800-01 | 4 | LOC_Os08g01080 | Homeodomain-like | 511 to 515, 589 to 593, 854 to 858, 858 to 862 |
| | >Os08t0137300-01 | 4 | LOC_Os08g04300 | Protein of unknown function DUF604 family protein. fringe-related protein | 145 to 149, 199 to 203, 469 to 473, 922 to 926 |
| | >Os08t0162800-01 | 4 | LOC_Os08g06550 | Acyl-CoA-binding protein | 173 to 177, 240 to 244, 343 to 347, 435 to 439 |
| | >Os08t0163500-01 | 6 | LOC_Os08g06640 | Protein of unknown function DUF1005 family protein. stress-induced protein | 315 to 319, 445 to 449, 467 to 471, 549 to 553, 610 to 614, 882 to 886 |
| | >Os08t0169700-01 | 5 | LOC_Os08g07290 | Predicted protein. HEAT repeat family protein | 155 to 159, 339 to 343, 410 to 414, 560 to 564, 612 to 616 |
| | >Os08t0170100-01 | 4 | LOC_Os08g07330 | NB-ARC domain. RGH1A | 448 to 452, 480 to 484, 801 to 805, 866 to 870 |

| | | | | | |
|---|---|---|---|---|---|
| | >Os08t0173700-01 | 5 | LOC_Os08g07700 | Ethylene-responsive transcription factor 4 | 24 to 28, 56 to 60, 203 to 207, 382 to 386, 746 to 750 |
| | >Os08t0174700-01 | 6 | LOC_Os08g07760 | SERK1 BRASSINOSTEROID INSENSITIVE 1-associated receptor kinase 1 precursor | 229 to 233, 259 to 263, 567 to 571, 626 to 630, 790 to 794, 854 to 858 |
| | >Os08t0189600-01 | 5 | LOC_Os08g09010 | Germin-like protein 8-7, Disease resistance Cupin domain | 162 to 166, 320 to 324, 352 to 356, 735 to 739, 921 to 925 |
| | >Os08t0189900-00 | 4 | LOC_Os08g09060 | Oxalate oxidase-like protein or germin-like protein (Germin-like 8) (Germin-like 12). Cupin domain | 101 to 105, 384 to 388, 566 to 570, 834 to 838 |
| | >Os08t0192900-02 | 4 | LOC_Os08g09350 | Nucleotide-binding, alpha-beta plait domain. Gar2 | 291 to 295, 377 to 381, 582 to 586, 925 to 929 |
| | >Os08t0198100-00 | 5 | LOC_Os08g09810 | WRKY106 | 94 to 98, 218 to 222, 302 to 306, 612 to 616, 867 to 871 |
| | >Os08t0206900-01 | 4 | LOC_Os08g10600 | UTP--glucose-1-phosphate uridylyltransferase family protein. | 115 to 119, 487 to 491, 499 to 503, 569 to 573 |
| | >Os08t0236400-00 | 4 | LOC_Os08g13870 | Epidermal growth factor-like, type 3 domain. S-locus lectin protein kinase family protein | 201 to 205, 288 to 292, 670 to 674, 805 to 809 |
| | >Os08t0242900-01 | 4 | LOC_Os08g14460 | Afadin/alpha-actinin-binding domain | 207 to 211, 391 to 395, 497 to 501, 626 to 630 |
| | >Os08t0366100-01 | 4 | LOC_Os08g27850 | Endothelial differentiation-related factor 1 (EDF-1) | 240 to 244, 366 to 370, 384 to 388, 438 to 442 |
| | >Os08t0374100-01 | 4 | LOC_Os08g28680 | Ubiquitin carrier protein | 222 to 226, 318 to 322, 360 to 364, 712 to 716 |
| | >Os08t0374100-02 | 4 | LOC_Os08g28680 | Ubiquitin carrier protein | 197 to 201, 293 to 297, 335 to 339, 687 to 691 |
| | >Os08t0434500-00 | 4 | LOC_Os08g33740 | Hypothetical protein. CSLA11 - cellulose synthase-like family A | 5 to 9, 156 to 160, 660 to 664, 667 to 671 |
| | >Os08t0456200-00 | 5 | LOC_Os08g35510 | Flavonoid 3-monooxygenase. cytochrome P450 | 233 to 237, 552 to 556, 811 to 815, 865 to 869, 871 to 875 |

| | | | | | |
|---|---|---|---|---|---|
| | >Os08t0460800-01 | 4 | LOC_Os08g35870 | OsFBL48 - F-box domain and LRR | 35 to 39, 229 to 233, 473 to 477, 703 to 707 |
| | >Os08t0465300-01 | 5 | LOC_Os08g36250 | Sodium/hydrogen exchanger family protein | 115 to 119, 538 to 542, 638 to 642, 910 to 914, 938 to 942 |
| | >Os08t0481800-01 | 4 | LOC_Os08g37600 | Plastidic general dicarboxylate transporter. citrate transporter | 39 to 43, 187 to 191, 300 to 304, 872 to 876 |
| | >Os08t0515800-01 | 5 | LOC_Os08g40430 | Mitochodrial transcription termination factor-related family protein. mTERF domain | 411 to 415, 539 to 543, 697 to 701, 710 to 714, 873 to 877 |
| | >Os08t0519600-01 | 4 | LOC_Os08g40820 | Protein of unknown function DUF1666 family protein | 244 to 248, 270 to 274, 326 to 330, 397 to 401 |
| | >Os08t0519600-02 | 4 | LOC_Os08g40820 | Protein of unknown function DUF1666 family protein | 175 to 179, 201 to 205, 257 to 261, 328 to 332 |
| | >Os08t0526500-01 | 4 | LOC_Os08g41480 | SOX-1 protein. SAM domain | 502 to 506, 534 to 538, 683 to 687, 737 to 741 |
| | >Os08t0532600-01 | 4 | | Peroxidase | 297 to 301, 343 to 347, 520 to 524, 534 to 538 |
| | >Os08t0543400-03 | 5 | LOC_Os08g43040 | Transferase family protein | 26 to 30, 191 to 195, 229 to 233, 274 to 278, 326 to 330 |
| **TGACT** | >Os08t0100800-01 | 4 | LOC_Os08g01080 | Homeodomain-like | 511 to 515, 589 to 593, 854 to 858, 858 to 862 |
| | >Os08t0137300-01 | 4 | LOC_Os08g04300 | Protein of unknown function DUF604 family protein. fringe-related protein | 145 to 149, 199 to 203, 469 to 473, 922 to 926 |
| | >Os08t0162800-01 | 4 | LOC_Os08g06550 | Acyl-CoA-binding protein | 173 to 177, 240 to 244, 343 to 347, 435 to 439 |
| | >Os08t0163500-01 | 6 | LOC_Os08g06640 | Protein of unknown function DUF1005 family protein. stress-induced protein | 315 to 319, 445 to 449, 467 to 471, 549 to 553, 610 to 614, 882 to 886 |
| | >Os08t0169700-01 | 5 | LOC_Os08g07290 | Predicted protein. HEAT repeat family protein | 155 to 159, 339 to 343, 410 to 414, 560 to 564, 612 to 616 |
| | >Os08t0170100-01 | 4 | LOC_Os08g07330 | NB-ARC domain. RGH1A, | 448 to 452, 480 to 484, 801 to 805, 866 to 870 |
| | >Os08t0173700-01 | 5 | LOC_Os08g07700 | Ethylene-responsive transcription factor 4 | 24 to 28, 56 to 60, 203 to 207, 382 to 386, 746 to 750 |

| | | | | | |
|---|---|---|---|---|---|
| | >Os08t0174700-01 | 6 | LOC_Os08g07760 | SERK1 BRASSINOSTEROID INSENSITIVE 1-associated receptor kinase 1 precursor | 229 to 233, 259 to 263, 567 to 571, 626 to 630, 790 to 794, 854 to 858 |
| | >Os08t0189600-01 | 5 | LOC_Os08g09010 | Germin-like protein 8-7, Disease resistance Cupin domain | 162 to 166, 320 to 324, 352 to 356, 735 to 739, 921 to 925 |
| | >Os08t0189900-00 | 4 | LOC_Os08g09060 | Oxalate oxidase-like protein or germin-like protein (Germin-like 8) (Germin-like 12). Cupin domain | 101 to 105, 384 to 388, 566 to 570, 834 to 838 |
| | >Os08t0192900-02 | 4 | LOC_Os08g09350 | Nucleotide-binding, alpha-beta plait domain. gar2 | 291 to 295, 377 to 381, 582 to 586, 925 to 929 |
| | >Os08t0198100-00 | 5 | LOC_Os08g09810 | WRKY106 | 94 to 98, 218 to 222, 302 to 306, 612 to 616, 867 to 871 |
| | >Os08t0206900-01 | 4 | LOC_Os08g10600 | UTP--glucose-1-phosphate uridylyltransferase family protein. | 115 to 119, 487 to 491, 499 to 503, 569 to 573 |
| | >Os08t0236400-00 | 4 | LOC_Os08g13870 | Epidermal growth factor-like, type 3 domain. S-locus lectin protein kinase family protein, | 201 to 205, 288 to 292, 670 to 674, 805 to 809 |
| | >Os08t0242900-01 | 4 | LOC_Os08g14460 | Afadin/alpha-actinin-binding domain | 207 to 211, 391 to 395, 497 to 501, 626 to 630 |
| | >Os08t0366100-01 | 4 | LOC_Os08g27850 | Endothelial differentiation-related factor 1 (EDF-1) | 240 to 244, 366 to 370, 384 to 388, 438 to 442 |
| | >Os08t0374100-01 | 4 | LOC_Os08g28680 | Ubiquitin carrier protein | 222 to 226, 318 to 322, 360 to 364, 712 to 716 |
| | >Os08t0374100-02 | 4 | LOC_Os08g28680 | Ubiquitin carrier protein | 197 to 201, 293 to 297, 335 to 339, 687 to 691 |
| | >Os08t0434500-00 | 4 | LOC_Os08g33740 | Hypothetical protein. CSLA11 - cellulose synthase-like family A, | 5 to 9, 156 to 160, 660 to 664, 667 to 671 |
| | >Os08t0456200-00 | 5 | LOC_Os08g35510 | Flavonoid 3-monooxygenase. cytochrome P450 | 233 to 237, 552 to 556, 811 to 815, 865 to 869, 871 to 875 |
| | >Os08t0460800-01 | 4 | LOC_Os08g35870 | OsFBL48 - F-box domain and LRR | 35 to 39, 229 to 233, 473 to 477, 703 to 707 |

| | | | | | |
|---|---|---|---|---|---|
| | >Os08t0465300-01 | 5 | LOC_Os08g36250 | Sodium/hydrogen exchanger family protein | 115 to 119, 538 to 542, 638 to 642, 910 to 914, 938 to 942 |
| | >Os08t0481800-01 | 4 | LOC_Os08g37600 | Plastidic general dicarboxylate transporter. citrate transporter | 39 to 43, 187 to 191, 300 to 304, 872 to 876 |
| | >Os08t0515800-01 | 5 | LOC_Os08g40430 | Mitochodrial transcription termination factor-related family protein. mTERF domain | 411 to 415, 539 to 543, 697 to 701, 710 to 714, 873 to 877 |
| | >Os08t0519600-01 | 4 | LOC_Os08g40820 | Protein of unknown function DUF1666 family protein | 244 to 248, 270 to 274, 326 to 330, 397 to 401 |
| | >Os08t0519600-02 | 4 | LOC_Os08g40820 | Protein of unknown function DUF1666 family protein | 175 to 179, 201 to 205, 257 to 261, 328 to 332 |
| | >Os08t0526500-01 | 4 | LOC_Os08g41480 | SOX-1 protein. SAM domain | 502 to 506, 534 to 538, 683 to 687, 737 to 741 |
| | >Os08t0532600-01 | 4 | | Peroxidase | 297 to 301, 343 to 347, 520 to 524, 534 to 538 |
| | >Os08t0543400-03 | 5 | LOC_Os08g43040 | Transferase family protein | 26 to 30, 191 to 195, 229 to 233, 274 to 278, 326 to 330 |
| **TTGAC** | >Os08t0101500-01 | 4 | LOC_Os08g01120 | sulfate transporter | 232 to 236, 381 to 385, 884 to 888, 909 to 913 |
| | >Os08t0114300-01 | 4 | LOC_Os08g02230 | D-arabinono-1,4-lactone oxidase domain. FAD-binding and arabino-lactone oxidase domains | 248 to 252, 599 to 603, 804 to 808, 980 to 984 |
| | >Os08t0138700-01 | 4 | LOC_Os08g04420 | Serine/threonine protein kinase-related domain. tyrosine protein kinase domain | 379 to 383, 859 to 863, 875 to 879, 964 to 968 |
| | >Os08t0154300-01 | 4 | LOC_Os08g05830 | Aldolase-type TIM barrel domain. transaldolase, | 281 to 285, 526 to 530, 534 to 538, 545 to 549 |
| | >Os08t0154300-02 | 4 | LOC_Os08g05830 | Aldolase-type TIM barrel domain. transaldolase, | 253 to 257, 498 to 502, 506 to 510, 517 to 521 |
| | >Os08t0157600-01 | 5 | LOC_Os08g06110 | MYB transcription factor | 259 to 263, 310 to 314, 589 to 593, 598 to 602, 616 to 620 |
| | >Os08t0159100-01 | 4 | LOC_Os08g06250 | Tetraspanin domain | 294 to 298, 303 to 307, 608 to 612, 627 to 631 |
| | >Os08t0162000-01 | 5 | LOC_Os08g06470 | Transmembrane 9 superfamily protein member 1. transmembrane 9 superfamily member | 26 to 30, 44 to 48, 57 to 61, 473 to 477, 750 to 754 |

| | | | | | |
|---|---|---|---|---|---|
| | >Os08t0162600-01 | 4 | LOC_Os08g06530 | Rubredoxin-type Fe(Cys)4 protein family protein | 359 to 363, 368 to 372, 387 to 391 |
| | >Os08t0162600-02 | 4 | LOC_Os08g06530 | Rubredoxin-type Fe(Cys)4 protein family protein | 32 to 36, 462 to 466, 602 to 606, 754 to 758 |
| | >Os08t0168700-00 | 4 | LOC_Os08g07200 | Cytokinin-O-glucosyltransferase 2. UDP-glucoronosyl/UDP-glucosyl transferase | 285 to 289, 622 to 626, 715 to 719, 774 to 778 |
| | >Os08t0173700-01 | 6 | LOC_Os08g07700 | ETHYLENE RESPONSE FACTOR 80 | 23 to 27, 55 to 59, 202 to 206, 208 to 212, 381 to 385, 745 to 749 |
| | >Os08t0174700-01 | 4 | LOC_Os08g07760 | SERK1 (Fragment). BRASSINOSTEROID INSENSITIVE 1-associated receptor kinase 1 precursor | 228 to 232, 258 to 262, 678 to 682, 789 to 793 |
| | >Os08t0178700-01 | 4 | LOC_Os08g08110 | Calmodulin-binding diacylglycerol kinase. | 263 to 267, 404 to 408, 673 to 677, 882 to 886 |
| | >Os08t0189600-01 | 4 | LOC_Os08g09010 | GERMIN-LIKE PROTEIN 8-7 | 161 to 165, 319 to 323, 351 to 355, 920 to 924 |
| | >Os08t0191000-01 | 4 | LOC_Os08g09190 | Auxin efflux carrier domain | 249 to 253, 443 to 447, 502 to 506, 784 to 788 |
| | >Os08t0195000-00 | 4 | LOC_Os08g09590 | F-box domain, Skp2-like domain. OsFBL42 - F-box domain and LRR | 305 to 309, 678 to 682, 686 to 690, 697 to 701 |
| | >Os08t0203100-01 | 4 | LOC_Os08g10290 | SHR5-receptor-like kinase | 129 to 133, 138 to 142, 158 to 162, 703 to 707 |
| | >Os08t0203600-04 | 4 | LOC_Os08g10320; | SHR5-receptor-like kinase | 283 to 287, 292 to 296, 838 to 842, 928 to 932 |
| | >Os08t0230100-00 | 4 | LOC_Os08g13330) | retrotransposon protein, putative, Ty1-copia subclass | 129 to 133, 138 to 142, 158 to 162, 703 to 707 |
| | >Os08t0263000-00 | 5 | | Cytochrome P450-like protein (CYP86B1). cytochrome P450 protein | 475 to 479, 552 to 556, 563 to 567, 583 to 587, 594 to 598 |
| | >Os08t0275200-00 | 4 | LOC_Os08g17320 | Protein kinase | 182 to 186, 291 to 295, 552 to 556, 977 to 981 |
| | >Os08t0277300-00 | 5 | LOC_Os08g17510 | Flavonol 4-sulfotransferase. sulfotransferase domain | 488 to 492, 753 to 757, 765 to 769, 773 to 777, 784 to 788 |
| | >Os08t0289266-00 | 4 | LOC_Os08g19240 | Histone H2A. histone H2A.6 | 481 to 485, 487 to 491, 670 to 674, 697 to 701 |
| | >Os08t0323700-01 | 4 | LOC_Os08g23440 | Cation-chloride cotransporter. amino acid permease family protein | 17 to 21, 243 to 247, 359 to 363, 983 to 987 |

| | | | | | |
|---|---|---|---|---|---|
| | >Os08t0327200-01 | 4 | LOC_Os08g23790 | Pectin lyase fold/virulence factor domain. polygalacturonase | 74 to 78305to309532to536740to744 |
| | >Os08t0337700-01 | 4 | LOC_Os08g25010 | RabGAP/TBC domain | 257 to 261, 266 to 270, 285 to 289, 853 to 857 |
| | >Os08t0343300-01 | 4 | LOC_Os08g25460 | Zinc finger, RING/FYVE/PHD-type domain. transcription initiation factor | 257 to 261, 313 to 317, 336 to 340, 917 to 921 |
| | >Os08t0356700-01 | 4 | LOC_Os08g26840 | Protein of unknown function DUF247, plant family protein | 257 to 261, 348 to 352, 532 to 536, 746 to 750 |
| | >Os08t0376300-01 | 4 | LOC_Os08g28870 | Leucine-rich receptor-like protein kinase. receptor-like protein kinase 5 precursor | 31 to 35, 301 to 305, 395 to 399, 732 to 736 |
| | >Os08t0376500-00 | 4 | LOC_Os08g28880 | Patatin-related phospholipase A II gamma | 127 to 131, 133 to 137, 472 to 476, 491 to 495 |
| | >Os08t0376600-01 | 4 | LOC_Os08g28890 | Serine/threonine protein kinase domain. protein kinase family protein | 184 to 188, 674 to 678, 859 to 863, 909 to 913 |
| | >Os08t0377100-00 | 5 | LOC_Os08g28940 | OsFBX289 - F-box domain | 11 to 15, 82 to 86, 262 to 266, 442 to 446, 597 to 601 |
| | >Os08t0377200-01 | 5 | LOC_Os08g28950 | A-type response regulator, Cytokinin signaling response regulator receiver domain | 128 to 132, 355 to 359, 392 to 396, 424 to 428, 574 to 578 |
| | >Os08t0389300-00 | 4 | LOC_Os08g29950 | UbiA prenyltransferase family domain | 155 to 159, 414 to 418, 484 to 488, 489 to 493 |
| | >Os08t0396401-01 | 4 | LOC_Os08g30800 | KED-like protein. KED | 11 to 15, 30 to 34, 245 to 249, 388 to 392 |
| | >Os08t0398500-01 | 4 | LOC_Os08g31420 | Kelch related domain | 134 to 138, 360 to 364, 372 to 376, 658 to 662 |
| | >Os08t0412600-01 | 6 | LOC_Os08g31840 | RabGAP/TBC domain | 263 to 267, 415 to 419, 455 to 459, 621 to 625, 643 to 647, 787 to 791 |
| | >Os08t0421800-01 | 4 | LOC_Os08g32600 | Mitogen-activated protein kinase kinase kinase 1 | 453 to 457, 569 to 573, 605 to 609, 761 to 765 |
| | >Os08t0480200-01 | 4 | LOC_Os08g37456 | 2OG-Fe(II) oxygenase domain. flavonol synthase/flavanone 3-hydroxylase | 177 to 181, 722 to 726, 781 to 785, 808 to 812 |
| | >Os08t0480800-01 | 4 | LOC_Os08g37490 | TaWIN2. 14-3-3 protein | 112 to 116, 225 to 229, 472 to 476, 653 to 657 |

| | | | | | |
|---|---|---|---|---|---|
| | >Os08t0482100-01 | 5 | LOC_Os08g37610 | LETM1-like domain | 223 to 227, 364 to 368, 421 to 425, 848 to 852, 986 to 990 |
| | >Os08t0487800-01 | 5 | LOC_Os08g38086 | Heat-shock protein | 57 to 61, 66 to 70, 303 to 307, 655 to 659, 689 to 693 |
| | >Os08t0489100-00 | 4 | LOC_Os08g38160 | Cytokinin-O-glucosyltransferase, | 711 to 715, 815 to 819, 845 to 849, 884 to 888 |
| | >Os08t0501700-01 | 4 | LOC_Os08g39240 | OsWAK76 - WALL-ASSOCIATED KINASE GENE 76 | 380 to 384, 418 to 422, 777 to 781, 782 to 786 |
| | >Os08t0526500-01 | 7 | LOC_Os08g41480 | SOX-1 protein. SAM domain containing protein | 228 to 232, 263 to 267, 501 to 505, 533 to 537, 682 to 686, 688 to 692, 736 to 740 |
| | >Os08t0527300-01 | 5 | LOC_Os08g41550 | Peptidase C19, ubiquitin carboxyl-terminal hydrolase 2 domain. ubiquitin carboxyl-terminal hydrolase | 8 to 12, 113 to 117, 192 to 196 201 to 205, 220 to 224 |
| | >Os08t0529300-00 | 4 | LOC_Os08g41750 | OsFBX303 - F-box domain | 308 to 312, 317 to 321, 336 to 340, 657 to 661 |
| | >Os08t0531000-01 | 6 | LOC_Os08g41880 | Diphosphonucleotide phosphatase 1 precursor. nucleotide pyrophosphatase/ phosphodiesterase, | 133 to 137, 142 to 146, 161 to 165, 462 to 466, 603 to 607, 622 to 626 |
| | >Os08t0533600-01 | 4 | LOC_Os08g42100 | ACR4. ACT domain | 435 to 439, 546 to 550, 657 to 661, 754 to 758 |
| | >Os08t0533600-02 | 4 | LOC_Os08g42100 | ACR4. ACT domain | 372 to 376, 483 to 487, 594 to 598, 691 to 695 |
| | >Os08t0538200-01 | 4 | LOC_Os08g42570 | Protein of unknown function DUF247, plant family protein. plant protein of unknown function domain | 19 to 23, 237 to 241, 583 to 587, 726 to 730 |
| | >Os08t0543100-00 | 4 | LOC_Os08g43010 | NBS-LRR type resistance protein disease resistance RPP13-like protein 1 | 466 to 470, 604 to 608, 845 to 849, 856 to 860 |
| | >Os08t0544400-01 | 4 | LOC_Os08g43120 | ABC-2 type transporter domain. Plant PDR ABC transporter associated domain | 447 to 451, 469 to 473, 561 to 565, 910 to 914 |
| | >Os08t0558900-01 | 5 | LOC_Os08g44460 | F1F0-ATPase inhibitor protein | 309 to 313, 318 to 322, 337 to 341, 364 to 368, 636 to 640 |

| | | | | | |
|---|---|---|---|---|---|
| | >Os08t0558900-02 | 5 | LOC_Os08g44460 | F1F0-ATPase inhibitor protein | 261 to 265, 270 to 274, 289 to 293, 316 to 320, 588 to 592 |
| | >Os08t0566400-01 | 4 | LOC_Os08g45180 | Ribokinase family protein. kinase, pfkB family | 98 to 102, 412 to 416, 537 to 541, 650 to 654 |
| **TTGACA** | >Os08t0114300-01 | 4 | LOC_Os08g02230 | D-arabinono-1,4-lactone oxidase domain. FAD-binding and arabino-lactone oxidase domains | 248 to 253, 599 to 604, 804 to 809, 980 to 985 |
| | >Os08t0246800-01 | 4 | LOC_Os08g14880 | Transposon protein | 253 to 258, 465 to 470, 650 to 655, 849 to 854 |
| | >Os08t0487800-01 | 4 | LOC_Os08g38086 | Heat shock protein | 57 to 62, 66to71, 303 to 308, 655 to 660 |
| | >Os08t0533600-01 | 4 | LOC_Os08g42100 | ACR4. ACT domain | 435 to 440, 546 to 551, 657 to 662, 754 to 759 |
| | >Os08t0533600-02 | 4 | LOC_Os08g42100 | ACR4. ACT domain | 372 to 377, 483 to 488, 594 to 599, 691 to 696 |
| **TTGACC** | >Os08t0412600-01 | 4 | LOC_Os08g31840 | RabGAP/TBC domain | 415 to 420, 455 to 460, 621 to 626, 643 to 648 |
| **TTGACT** | >Os08t0173700-01 | 5 | LOC_Os08g07700 | Ethylene-responsive transcription factor 4 | 23 to 28, 55 to 60, 202 to 207, 381 to 386, 745 to 750 |
| | >Os08t0189600-01 | 4 | LOC_Os08g09010 | Germin-like protein 8-7 | 161 to 166, 319 to 324, 351 to 356, 920 to 925 |
| | >Os08t0526500-01 | 4 | LOC_Os08g41480 | SOX-1 protein. SAM domain | 501 to 506, 533 to 538, 682 to 687, 736 to 741 |
| **Chromosome 9** | | | | | |
| **Motif** | **ID** | **Frequency** | **Locus ID** | **Protein name** | **Positions in 1k promoter region** |
| **GTACGTAC** | >Os09t0571000-00 | 5 | LOC_Os09g39750 | DUF966-stress repressive gene 7 | 131 to 138, 459 to 466, 492 to 499, 496 to 503, 845 to 852 |
| **TGACC** | >Os09t0101200-00 | 4 | LOC_Os09g01300 | Transposon protein. | 599 to 603, 694 to 698, 702 to 706, 823 to 827 |
| | >Os09t0309200-01 | 4 | LOC_Os09g13890 | Calmodulin binding protein-like family protein | 601 to 605, 726 to 730, 763 to 767, 888 to 892 |
| | >Os09t0371200-01 | 4 | LOC_Os09g20500 | Major facilitator superfamily protein | 253 to 257, 455 to 459, 809 to 813, 996 to 1000 |
| | >Os09t0390400-01 | 4 | LOC_Os09g22280 | OsRAD23-like | 307 to 311, 446 to 450, 601 to 605, 888 to 892 |
| | >Os09t0416900-00 | 5 | LOC_Os09g25000 | Armadillo-like helical domain. spotted leaf 11 | 411 to 415, 514 to 518, 613 to 617, 732 to 736, 840 to 844 |
| | >Os09t0448150-00 | 5 | | Carboxyl-terminal proteinase | 48 to 52, 85 to 89, 97 to 101, 342 to 346, 923 to 927 |

| | | | | | |
|---|---|---|---|---|---|
| | >Os09t0457400-01 | 5 | LOC_Os09g28400 | Alpha-amylase isozyme 3A precursor alpha-amylase precursor | 63 to 67, 73 to 77, 181 to 185, 245 to 249, 583 to 587 |
| | >Os09t0520100-01 | 5 | LOC_Os09g34850 | DNA polymerase delta, subunit 4 family protein. POLD4A - Putative DNA polymerase delta complex subunit | 23 to 27, 40 to 44, 45 to 49, 642 to 646, 722 to 726 |
| | >Os09t0520100-02 | 5 | LOC_Os09g34850 | DNA polymerase delta, subunit 4 family protein. POLD4A - Putative DNA polymerase delta complex subunit | 4 to 8, 21 to 25, 26 to 30, 623 to 627, 703 to 707 |
| | >Os09t0529700-01 | 4 | LOC_Os09g36020 | EAP30 family protein. EAP30/Vps36 family domain | 201 to 205, 288 to 292, 347 to 351, 389 to 393 |
| | >Os09t0560900-01 | 4 | LOC_Os09g38790 | Zinc finger, C2H2-like domain. ZOS9-19 - C2H2 zinc finger protein | 817 to 821, 822 to 826, 840 to 844, 873 to 877 |
| **TGACT** | >Os09t0115400-03 | 4 | LOC_Os09g02700 | Poly(A)-binding protein. polyadenylate-binding protein | 192 to 196, 358 to 362, 588 to 592, 905 to 909 |
| | >Os09t0115400-04 | 4 | LOC_Os09g02700 | Poly(A)-binding protein. polyadenylate-binding protein | 2 to 6, 168 to 172, 398 to 402, 715 to 719 |
| | >Os09t0115600-01 | 4 | LOC_Os09g02729 | PLC-like phosphodiesterase, TIM beta/alpha-barrel domain | 402 to 406, 620 to 624, 762 to 766, 823 to 827 |
| | >Os09t0115600-02 | 4 | LOC_Os09g02729 | PLC-like phosphodiesterase, TIM beta/alpha-barrel domain | 355 to 359, 573 to 577, 715 to 719, 776 to 780 |
| | >Os09t0123300-01 | 4 | LOC_Os09g03620 | Calmodulin-binding receptor-like kinase. wall-associated receptor kinase-like 20 precursor | 212 to 216, 551 to 555, 730 to 734, 933 to 937 |
| | >Os09t0124100-00 | 4 | LOC_Os09g03680 | Ankyrin repeat | 242 to 246, 349 to 353, 759 to 763, 784 to 788 |
| | >Os09t0279400-01 | 4 | LOC_Os09g10750 | Rhodanese-like domain | 73 to 77, 273 to 277, 454 to 458, 658 to 662 |
| | >Os09t0314200-00 | 4 | LOC_Os09g14490 | Disease resistance protein domain. TIR-NBS type disease resistance protein | 94 to 98, 506 to 510, 572 to 576, 920 to 924 |

| | | | | | |
|---|---|---|---|---|---|
| | >Os09t0320400-01 | 4 | LOC_Os09g15170 | Permease. nucleobase-ascorbate transporter | 12 to 16, 420 to 424, 600 to 604, 907 to 911 |
| | >Os09t0333000-00 | 4 | LOC_Os09g16380 | ABC transporter-like domain. pleiotropic drug resistance protein | 103 to 107, 187 to 191, 634 to 638, 665 to 669 |
| | >Os09t0390400-01 | 4 | LOC_Os09g22280 | OsRAD23 | 33 to 37, 342 to 346, 498 to 502, 653 to 657 |
| | >Os09t0445100-00 | 5 | LOC_Os09g27270 | Ribosomal RNA methyltransferase RrmJ/FtsJ domain. ribosomal RNA large subunit methyltransferase J | 191 to 195, 326 to 330, 604 to 608, 608 to 612, 772 to 776 |
| | >Os09t0448200-02 | 4 | LOC_Os09g27580) | High-affinity potassium transporter | 69 to 73, 304 to 308, 926 to 930, 935 to 939 |
| | >Os09t0458200-01 | 4 | LOC_Os09g28470 | leucine-rich repeat family protein | 83 to 87, 175 to 179, 332 to 336, 858 to 862 |
| | >Os09t0481650-00 | 5 | LOC_Os09g30390 | Polyubiquitin-like protein | 3 to 7, 133 to 137, 342 to 346, 361 to 365, 495 to 499 |
| | >Os09t0482640-01 | 4 | LOC_Os09g30454 | Serine/threonine protein kinase-related domain. OsWAK87 | 22 to 26, 655 to 659, 847 to 851, 855 to 859 |
| | >Os09t0482660-01 | 4 | LOC_Os09g30458) | Subtilisin-type protease | 231 to 235, 451 to 455, 483 to 487, 726 to 730 |
| | >Os09t0484800-01 | 4 | LOC_Os09g31120 | Pirin-like protein. pirin | 376 to 380, 395 to 399, 463 to 467, 507 to 511 |
| | >Os09t0491100-02 | 4 | LOC_Os09g31430 | Beta-primeverosidase Os9bglu30 - beta-glucosidase | 487 to 491, 533 to 537, 569 to 573, 573 to 577 |
| | >Os09t0570300-02 | 5 | LOC_Os09g39670 | Short-chain dehydrogenase Tic32. oxidoreductase, short chain dehydrogenase/reductase family domain | 562 to 566, 720 to 724, 782 to 786, 937 to 941, 959 to 963 |
| **TTGAC** | >Os09t0123300-01 | 4 | LOC_Os09g03620 | Calmodulin-binding receptor-like kinase. wall-associated receptor kinase-like 20 precursor | 392 to 396, 550 to 554, 729 to 733, 895 to 899 |
| | >Os09t0246200-01 | 4 | LOC_Os09g07120 | DNA-directed RNA polymerase 1B, mitochondrial precursor | 243 to 247, 654 to 658, 907 to 911, 941 to 945 |
| | >Os09t0248300-01 | 5 | LOC_Os09g07380 | Tektin domain | 10 to 14, 159 to 163, 165 to 169, 427 to 431, 454 to 458 |

| | | | | | |
|---|---|---|---|---|---|
| | >Os09t0267400-01 | 4 | LOC_Os09g09460 | Harpin-induced 1 domain | 156 to 160, 161 to 165, 169 to 173, 534 to 538 |
| | >Os09t0267600-01 | 4 | LOC_Os09g09480 | Charged multivesicular body protein 4b. SNF7 domain | 250 to 254, 352 to 356, 405 to 409, 777 to 781 |
| | >Os09t0279400-01 | 5 | LOC_Os09g10750 | Rhodanese-like domain | 72 to 76, 272 to 276, 453 to 457, 657 to 661, 736 to 740 |
| | >Os09t0296800-01 | 4 | LOC_Os09g12540 | Chlorophyll A-B binding protein family protein. chlorophyll A-B binding protein | 285 to 289, 317 to 321, 511 to 515, 750 to 754 |
| | >Os09t0309200-01 | 4 | LOC_Os09g13890 | Calmodulin binding protein-like family protein | 53 to 57, 459 to 463, 600 to 604, 725 to 729 |
| | >Os09t0320400-01 | 4 | LOC_Os09g15170 | Permease. nucleobase-ascorbate transporter | 419 to 423, 451 to 455, 599 to 603, 906 to 910 |
| | >Os09t0324400-01 | 4 | LOC_Os09g15560 | Cyclin-like F-box domain. OsFBX314 - F-box domain | 40 to 44, 489 to 493, 594 to 598, 885 to 889 |
| | >Os09t0342000-01 | 4 | LOC_Os09g17190 | Cyclin-like F-box domain. OsFBX320 - F-box domain | 129 to 133, 245 to 249, 661 to 665, 859 to 863 |
| | >Os09t0400000-00 | 4 | LOC_Os09g23540 | Cinnamyl alcohol dehydrogenase | 220 to 224, 740 to 744, 824 to 828, 893 to 897 |
| | >Os09t0417600-01 | 4 | LOC_Os09g25060 | WRKY transcription factor 76 | 88 to 92, 551 to 555, 669 to 673, 876 to 880 |
| | >Os09t0417600-02 | 4 | LOC_Os09g25060 | WRKY transcription factor 76 | 15 to 19, 478 to 482, 596 to 600, 803 to 807 |
| | >Os09t0419600-01 | 5 | LOC_Os09g25200 | Zinc finger, RING-type domain. | 232 to 236, 310 to 314, 882 to 886, 888 to 892, 903 to 907 |
| | >Os09t0425300-00 | 4 | LOC_Os09g25700 | TsetseEP precursor | 17 to 21, 89 to 93, 234 to 238, 259 to 263 |
| | >Os09t0431600-00 | 4 | LOC_Os09g26190 | Cystathionine beta-synthase, core domain. CBS domain | 234 to 238, 639 to 643, 648 to 652, 667 to 671 |
| | >Os09t0433900-01 | 4 | LOC_Os09g26380 | ALANINE AMINOTRANSFERASE 5 | 45 to 49, 373 to 377, 758 to 762, 858 to 862 |
| | >Os09t0452900-01 | 4 | LOC_Os09g27950 | Glycosyl transferase, family 31 protein. galactosyltransferase | 216 to 220, 550 to 554, 699 to 703, 811 to 815 |

| | | | | | |
|---|---|---|---|---|---|
| | >Os09t0458900-01 | 4 | LOC_Os09g28510 | EF hand domain. EF hand family protein | 118 to 122, 182 to 186, 191 to 195, 210 to 214 |
| | >Os09t0473300-01 | 4 | LOC_Os09g29750 | Methylcytosine binding domain protein. ZOS9-15 - C2H2 zinc finger protein | 18 to 22, 202 to 206, 368 to 372, 641 to 645 |
| | >Os09t0474400-00 | 4 | LOC_Os09g29850 | methyltransferase small domain | 361 to 365, 453 to 457, 498 to 502, 512 to 516 |
| | >Os09t0482200-01 | 4 | LOC_Os09g30414 | Peptidase A1 domain. aspartic proteinase nepenthesin-2 precursor | 369 to 373, 473 to 477, 714 to 718, 894 to 898 |
| | >Os09t0482400-03 | 4 | | Heat shock protein 81-3 | 85 to 89, 441 to 445, 450 to 454, 470 to 474 |
| | >Os09t0482500-01 | 4 | | Peptidase A1 domain | 369 to 373, 473 to 477, 714 to 718, 894 to 898 |
| | >Os09t0482600-02 | 4 | LOC_Os09g30418 | Heat shock protein 81-3. heat shock protein | 85 to 89, 441 to 445, 450 to 454, 470 to 474 |
| | >Os09t0492700-01 | 4 | LOC_Os09g31970 | 3-hydroxy-3-methylglutaryl coenzyme A reductase | 11 to 15, 184 to 188, 364 to 368, 485 to 489 |
| | >Os09t0500151-00 | 5 | LOC_Os09g32410 | OsFBX333 - F-box domain | 505 to 509, 566 to 570, 613 to 617, 660 to 664, 754 to 758 |
| | >Os09t0507300-01 | 5 | LOC_Os09g32952 | L-ascorbate oxidase precursor | 456 to 460, 471 to 475, 542 to 546, 892 to 896, 953 to 957 |
| | >Os09t0511900-01 | 5 | LOC_Os09g33710 | Beta-glucosidase 32 | 51 to 55, 370 to 374, 395 to 399, 531 to 535, 581 to 585 |
| | >Os09t0520100-01 | 5 | LOC_Os09g34850 | DNA polymerase delta, subunit 4 family protein. POLD4A - Putative DNA polymerase delta complex subunit | 22 to 26, 476 to 480, 641 to 645, 689 to 693, 721 to 725 |
| | >Os09t0520100-02 | 5 | LOC_Os09g34850 | DNA polymerase delta, subunit 4 family protein. POLD4A - Putative DNA polymerase delta complex subunit | 3 to 7, 457 to 461, 622 to 626, 670 to 674, 702 to 706 |
| | >Os09t0521300-00 | 4 | LOC_Os09g34950 | TCP family transcription factor | 74 to 78, 81 to 85, 257 to 261, 391 to 395 |
| | >Os09t0530900-01 | 5 | LOC_Os09g36120 | Oxydoreductase-like protein | 225 to 229, 232 to 236, 247 to 251, 313 to 317, 397 to 401 |

| | | | | | |
|---|---|---|---|---|---|
| | >Os09t0531900-02 | 4 | LOC_Os09g36190 | Glycosyltransferase QUASIMODO1 glycosyl transferase family 8 | 450 to 454, 529 to 533, 844 to 848, 857 to 861 |
| | >Os09t0533400-01 | 4 | LOC_Os09g36300 | Lon protease homolog. OsLonP4 - Putative Lon protease homologue | 61 to 65, 118 to 122, 466 to 470, 938 to 942 |
| | >Os09t0555100-00 | 4 | LOC_Os09g38210 | Auxin Efflux Carrier family protein. auxin efflux carrier component | 106 to 110, 267 to 271, 690 to 694, 742 to 746 |
| | >Os09t0560900-01 | 5 | LOC_Os09g38790 | Zinc finger, C2H2-like domain. ZOS9-19 - C2H2 zinc finger protein | 68 to 72, 498 to 502, 674 to 678, 812 to 816, 839 to 843 |
| | >Os09t0570850-00 | 4 | LOC_Os09g39730 | Histone H2B. Core histone H2A/H2B/H3/H4 domain | 199 to 203, 511 to 515, 741 to 745, 799 to 803 |
| | >Os09t0572700-02 | 4 | LOC_Os09g39730 | Histone H2B. Core histone H2A/H2B/H3/H4 domain | 248 to 252, 417 to 421, 598 to 602, 604 to 608 |
| **TTGACA** | >Os09t0500151-00 | 5 | LOC_Os09g32410 | OsFBX333 - F-box domain | 505 to 510, 566 to 571, 613 to 618, 660 to 665, 754 to 759 |
| **TTGACT** | >Os09t0279400-01 | 4 | LOC_Os09g10750 | Sulfurtransferase 3 | 72 to 77, 272 to 277, 453 to 458, 657 to 662 |
| **Chromosome 10** | | | | | |
| **Motif** | **ID** | **Frequency** | **Locus ID** | **Protein name** | **Positions in 1k promoter region** |
| **CTGACC** | >Os10t0532300-01 | 4 | LOC_Os10g38870 | Heavy metal transport/ detoxification protein domain | 374 to 379, 397 to 402, 420 to 425, 896 to 901 |
| **CTGACT** | >Os10t0502500-04 | 4 | LOC_Os10g35850 | Cytochrome b5-like Heme/Steroid binding | 550 to 555, 713 to 718, 747 to 752, 872 to 877 |
| **TGACC** | >Os10t0170200-01 | 4 | LOC_Os10g08930 | 40S ribosomal protein S20, | 48 to 52, 314 to 318, 325 to 329, 580 to 584 |
| | >Os10t0172600-01 | 4 | LOC_Os10g09200 | zinc finger protein | 257 to 261, 456 to 460, 612 to 616, 900 to 904 |
| | >Os10t0174548-00 | 4 | LOC_Os10g09550 | Calcium binding EGF domain. OsWAK106 | 338 to 342, 397 to 401, 737 to 741, 773 to 777 |
| | >Os10t0175800-00 | 4 | LOC_Os10g09710 | Nodulin protein. auxin-induced protein 5NG4 | 280 to 284, 573 to 577, 737 to 741, 824 to 828 |

| | | | | | |
|---|---|---|---|---|---|
| | >Os10t0458700-01 | 4 | LOC_Os10g32060 | Ty3-gypsy subclass | 254 to 258, 575 to 579, 936 to 940, 963 to 967 |
| | >Os10t0458700-02 | 4 | LOC_Os10g32060 | Ty3-gypsy subclass | 228 to 232, 549 to 553, 910 to 914, 937 to 941 |
| | >Os10t0469000-01 | 4 | LOC_Os10g33080 | Leucine-rich repeat receptor protein kinase EXS precursor | 12 to 16, 140 to 144, 163 to 167, 974 to 978 |
| | >Os10t0519300-00 | 4 | LOC_Os10g37500 | AAA-type ATPase family protein | 616 to 620, 663 to 667, 695 to 699, 868 to 872 |
| | >Os10t0532300-01 | 4 | LOC_Os10g38870 | Heavy metal transport/ detoxification protein domain | 375 to 379, 398 to 402, 421 to 425, 897 to 901 |
| **TGACT** | >Os10t0116900-01 | 4 | LOC_Os10g02760 | Hydroxyproline-rich glycoprotein family protein | 131 to 135, 158 to 162, 729 to 733, 959 to 963 |
| | >Os10t0136150-00 | 4 | LOC_Os10g04690 | F-box domain | 189 to 193, 370 to 374, 435 to 439, 902 to 906 |
| | >Os10t0136200-01 | 4 | LOC_Os10g04700 | Cyclin-like F-box domain. OsFBX361 - F-box domain | 38 to 42, 178 to 182, 701 to 705, 785 to 789 |
| | >Os10t0163100-01 | 4 | LOC_Os10g07994 | Autophagy protein Apg9 | 50 to 54, 457 to 461, 603 to 607, 609 to 613 |
| | >Os10t0343400-01 | 4 | LOC_Os10g20260 | Cellulose synthase family protein. CSLF7 - cellulose synthase-like family F; beta1,3;1,4 glucan synthase | 116 to 120, 194 to 198, 517 to 521, 759 to 763 |
| | >Os10t0377800-01 | 4 | LOC_Os10g23120 | Pyridoxamine 5-phosphate oxidase. | 10 to 14, 225 to 229, 387 to 391, 722 to 726 |
| | >Os10t0397900-00 | 5 | LOC_Os10g25850 | CCAAT-binding transcription factor, subunit B domain. | 218 to 222, 531 to 535, 680 to 684, 736 to 740, 843 to 847 |
| | >Os10t0441900-01 | 4 | LOC_Os10g30530 | Resistance protein candidate (Fragment). lectin-like receptor kinase | 375 to 379, 380 to 384, 564 to 568, 709 to 713 |
| | >Os10t0469900-02 | 4 | LOC_Os10g33170 | TGF-beta receptor, type I/II extracellular region family protein. POT domain transporter | 184 to 188, 342 to 346, 374 to 378, 606 to 610 |
| | >Os10t0502500-04 | 9 | LOC_Os10g35850 | Cytochrome b5 domain | 113 to 117, 551 to 555, 581 to 585, 586 to 590, 591 to 595, 714 to 718, 748 to 752, 873 to 877, 908 to 912 |

| | | | | | |
|---|---|---|---|---|---|
| | >Os10t0542400-01 | 4 | LOC_Os10g39640 | Expansin/Lol pI family protein. | 164 to 168, 570 to 574, 747 to 751, 797 to 801 |
| | >Os10t0567500-01 | 4 | LOC_Os10g41790 | RNA-directed DNA polymerase (reverse transcriptase) type II intron maturase protein | 201 to 205, 223 to 227, 622 to 626, 663 to 667 |
| | >Os10t0577700-01 | 4 | LOC_Os10g42700 | CS domain | 110 to 114, 177 to 181, 181 to 185, 564 to 568 |
| **TTGAC** | >Os10t0100500-01 | 4 | LOC_Os10g01060 | Serine/threonine protein kinase-related domain. protein kinase family protein | 515 to 519, 814 to 818, 906 to 910, 971 to 975 |
| | >Os10t0110250-00 | 4 | | Oligopeptide transporter domain | 210 to 214, 420 to 424, 570 to 574, 812 to 816 |
| | >Os10t0111900-00 | 4 | LOC_Os10g02276 | Protein kinase domain. OsWAK96 - OsWAK receptor-like protein kinase | 162 to 166, 195 to 199, 344 to 348, 928 to 932 |
| | >Os10t0132700-01 | 4 | LOC_Os10g04342 | NB-ARC domain. stripe rust resistance protein Yr10 | 271 to 275, 396 to 400, 487 to 491, 604 to 608 |
| | >Os10t0134900-00 | 5 | LOC_Os10g04560 | NB-ARC domain | 22 to 26, 38 to 42, 97 to 101, 443 to 447, 877 to 881 |
| | >Os10t0136150-00 | 4 | LOC_Os10g04690 | F-box domain | 188 to 192, 369 to 373, 375 to 379, 883 to 887 |
| | >Os10t0210500-01 | 4 | LOC_Os10g14920 | Protein of unknown function DUF6, transmembrane domain | 192 to 196, 373 to 377, 627 to 631, 750 to 754 |
| | >Os10t0210500-02 | 4 | LOC_Os10g14920 | Protein of unknown function DUF6, transmembrane domain | 178 to 182, 359 to 363, 613 to 617, 736 to 740 |
| | >Os10t0336300-00 | 4 | LOC_Os10g18990 | Leucine-rich repeat domain. receptor kinase 2 | 27 to 31, 602 to 606, 643 to 647, 671 to 675 |
| | >Os10t0344800-00 | 5 | | Phosphoglycerate mutase family protein | 132 to 136, 447 to 451, 466 to 470, 792 to 796, 959 to 963 |
| | >Os10t0351500-01 | 4 | LOC_Os10g21090 | Tyrosine protein kinase domain. ATP binding protein | 136 to 140, 157 to 161, 207 to 211, 434 to 438 |
| | >Os10t0360800-00 | 5 | | Protein kinase, catalytic domain | 17 to 21, 279 to 283, 418 to 422, 423 to 427, 849 to 853 |
| | >Os10t0368200-01 | 5 | LOC_Os10g22310 | Glutathione S-transferase GST 26 | 225 to 229, 317 to 321, 728 to 732, 737 to 741, 756 to 760 |
| | >Os10t0370100-02 | 4 | LOC_Os10g22484 | NB-ARC domain | 45 to 49, 77 to 81, 225 to 229, 565 to 569 |

| | | | | | |
|---|---|---|---|---|---|
| | >Os10t0389300-01 | 5 | LOC_Os10g25040 | Red chlorophyll catabolite reductase | 84 to 88, 116 to 120, 195 to 199, 776 to 780, 874 to 878 |
| | >Os10t0394000-01 | 4 | LOC_Os10g25420 | GDSL-like Lipase/Acylhydrolase family protein | 615 to 619, 624 to 628, 643 to 647, 935 to 939 |
| | >Os10t0395100-01 | 4 | LOC_Os10g25560 | Dil domain | 13 to 17, 73 to 77, 624 to 628, 941 to 945 |
| | >Os10t0397800-00 | 5 | LOC_Os10g25830 | Mitochondrial carrier protein family protein. | 137 to 141, 346 to 350, 641 to 645, 926 to 930, 950 to 954 |
| | >Os10t0434900-00 | 4 | | Protein STAR1 | 205 to 209, 238 to 242, 342 to 346, 534 to 538 |
| | >Os10t0441900-01 | 4 | LOC_Os10g30530 | Resistance protein candidate lectin-like receptor kinase | 374 to 378, 379 to 383, 572 to 576, 708 to 712 |
| | >Os10t0452300-01 | 6 | LOC_Os10g31460 | Eggshell protein family protein. retrotransposon protein | 26 to 30, 285 to 289, 294 to 298, 313 to 317, 519 to 523, 910 to 914 |
| | >Os10t0464000-01 | 4 | LOC_Os10g32700 | Hypersensitive-induced response protein | 236 to 240, 457 to 461, 720 to 724, 935 to 939 |
| | >Os10t0469900-02 | 5 | LOC_Os10g33170 | TGF-beta receptor, type I/II extracellular region family protein. POT domain transporter | 89 to 93, 341 to 345, 373 to 377, 605 to 609, 771 to 775 |
| | >Os10t0470700-01 | 5 | LOC_Os10g33210 | Peptide transporter. peptide transporter PTR3-A | 484 to 488, 638 to 642, 647 to 651, 666 to 670, 926 to 930 |
| | >Os10t0481300-01 | 4 | LOC_Os10g34020 | Glutathione S-transferase, N-terminal domain | 402 to 406, 462 to 466, 471 to 475, 490 to 494 |
| | >Os10t0493100-02 | 4 | LOC_Os10g35090 | K Homology, type 1 domain | 92 to 96, 434 to 438, 681 to 685, 846 to 850 |
| | >Os10t0493100-03 | 4 | LOC_Os10g35090 | K Homology, type 1 domain | 64 to 68, 406 to 410, 653 to 657, 818 to 822 |
| | >Os10t0495466-00 | 4 | | BRO1 domain domain | 174 to 178, 369 to 373, 506 to 510, 764 to 768 |
| | >Os10t0502500-04 | 8 | LOC_Os10g35850 | Cytochrome b5 domain. cytochrome b5-like Heme/Steroid binding domain | 536 to 540, 580 to 584, 585 to 589, 590 to 594, 732 to 736, 858 to 862, 891 to 895, 907 to 911 |

| | | | | | |
|---|---|---|---|---|---|
| | >Os10t0516300-01 | 4 | LOC_Os10g37210 | ETFQO (electron-transfer flavoprotein:ubiquinone oxidoreductase); catalytic/ electron carrier/ electron-transferring-flavoprotein dehydrogenase. FAD dependent oxidoreductase domain | 143 to 147, 363 to 367, 496 to 500, 782 to 786 |
| | >Os10t0522500-01 | 4 | LOC_Os10g37840 | CCB2 | 222 to 226, 314 to 318, 768 to 772, 935 to 939 |
| | >Os10t0522500-02 | 4 | LOC_Os10g37840 | CCB2 | 216 to 220, 308 to 312, 762 to 766, 929 to 933 |
| | >Os10t0527800-01 | 4 | LOC_Os10g38360 | Tau class GST protein 3. glutathione S-transferase | 504 to 508, 551 to 555, 582 to 586, 800 to 804 |
| | >Os10t0528300-01 | 4 | | Tau class GST protein 4 | 180 to 184, 491 to 495, 599 to 603, 610 to 614 |
| | >Os10t0528400-01 | 4 | LOC_Os10g38489 | Glutathione S-transferase, C-terminal-like domain GSTU6 | 124 to 128, 168 to 172, 687 to 691, 820 to 824 |
| | >Os10t0531700-00 | 4 | LOC_Os10g38800 | Serine/threonine protein kinase-related domain. leucine-rich repeat transmembrane protein kinase | 195 to 199, 368 to 372, 679 to 683, 990 to 994 |
| | >Os10t0547200-01 | 4 | LOC_Os10g39970 | Harpin-induced 1 domain. harpin-induced protein 1 | 3 to 7, 152 to 156, 326 to 330, 608 to 612 |
| | >Os10t0548400-00 | 5 | LOC_Os10g40070 | F25C20.9 | 226 to 230, 233 to 237, 628 to 632,704 to 708, 711to715 |
| | >Os10t0550200-01 | 4 | | Protein of unknown function DUF3615 domain | 88 to 92, 97 to 101, 116 to 120, 726 to 730 |
| | >Os10t0559500-01 | 4 | LOC_Os10g41020 | 2OG-Fe(II) oxygenase domain. flavonol synthase/flavanone 3-hydroxylase | 205 to 209, 253 to 257, 470 to 474, 846 to 850 |
| | >Os10t0559500-02 | 4 | LOC_Os10g41020 | 2OG-Fe(II) oxygenase domain. flavonol synthase/flavanone 3-hydroxylase | 26 to 30, 74 to 78, 291 to 295, 667 to 671 |
| | >Os10t0567900-01 | 4 | LOC_Os10g41838 | F-box protein interaction domain | 148 to 152, 673 to 677, 808 to 812, 989 to 993 |

| | | | | | |
|---|---|---|---|---|---|
| | >Os10t0571900-00 | 5 | LOC_Os10g42160 | Malic enzyme transposon protein | 441 to 445, 703 to 707, 795 to 799, 843 to 847, 860 to 864 |
| | >Os10t0577800-01 | 4 | LOC_Os10g42710 | Poly(ADP-ribose) polymerase, catalytic region domain. RCD1, | 279 to 283, 470 to 474, 552 to 556, 777 to 781 |
| | >Os10t0577800-02 | 4 | LOC_Os10g42710 | Poly(ADP-ribose) polymerase, catalytic region domain. RCD1 | 224 to 228, 415 to 419, 497 to 501, 722 to 726 |
| | >Os10t0580400-01 | 4 | LOC_Os10g42960 | Urea active transporter-like protein. urea active transporter, | 337 to 341, 430 to 434, 509 to 513, 655 to 659 |
| **TTGACA** | >Os10t0567900-01 | 4 | LOC_Os10g41838 | F-box protein interaction domain | 148 to 153, 673 to 678, 808 to 813, 989 to 994 |
| **TTGACT** | >Os10t0502500-04 | 4 | LOC_Os10g35850 | Cytochrome b5 domain | 580 to 585, 585 to 590, 590 to 595, 907 to 912 |
| **Chromosome 11** | | | | | |
| **Motif** | **ID** | **Frequency** | **Locus ID** | **Protein name** | **Positions in 1k promoter region** |
| **TTTGACT** | >Os11t0244100-00 | 4 | LOC_Os11g13970; | transferase family protein, | 330 to 336, 343 to 349, 492 to 498, 620 to 626 |
| | >Os11t0306300-01 | 4 | LOC_Os11g20080 | Herbicide safener binding protein. O-methyltransferase, | 99 to 105, 297 to 303, 340 to 346, 459 to 465 |
| **TTGACTT** | >Os11t0306300-01 | 4 | LOC_Os11g20080 | Herbicide safener binding protein. O-methyltransferase | 100 to 106, 298 to 304, 341 to 347, 460 to 466 |
| **TTGACCA** | >Os11t0529900-00 | 4 | LOC_Os11g32620 | Chalcone/stilbene synthase, N-terminal domain | 88 to 94, 96 to 102, 119 to 125, 127 to 133 |
| **GTTGAC** | >Os11t0474500-00 | 4 | LOC_Os11g28470 | NB-ARC domain | 207 to 212, 243 to 248, 808 to 813, 895 to 900 |
| | >Os11t0669100-01 | 4 | LOC_Os11g44680 | Calmodulin binding protein-like family protein | 424 to 429, 434 to 439, 533 to 538, 572 to 577 |
| | >Os11t0474500-00 | 4 | LOC_Os11g28470 | NB-ARC domain | 207 to 212, 243 to 248, 808 to 813, 895 to 900 |

| | | | | | |
|---|---|---|---|---|---|
| **TGACC** | >Os11t0108932-01 | 4 | LOC_Os11g01780 | F-box protein 563 | 92 to 96, 175 to 179, 807 to 811, 881 to 885 |
| | >Os11t0120725-00 | 4 | LOC_Os11g02774 | ATPase-like protein. histidine triad family protein | 799 to 803, 941 to 945, 969 to 973, 977 to 981 |
| | >Os11t0145000-01 | 5 | LOC_Os11g04840 | phosphatidylinositol-4-phosphate 5-kinase family protein. | 20 to 24, 26 to 30, 334 to 338, 377 to 381, 664 to 668 |
| | >Os11t0233201-01 | 4 | LOC_Os11g12560 | Receptor-like protein kinase HAIKU2 precursor | 38 to 42, 192 to 196, 454 to 458, 612 to 616 |
| | >Os11t0249900-01 | 4 | LOC_Os11g14420 | Protein kinase-like domain | 21 to 25, 275 to 279, 321 to 325, 805 to 809 |
| | >Os11t0256300-01 | 4 | LOC_Os11g15000 | Zinc finger, GRF-type domain | 397 to 401, 496 to 500, 824 to 828, 873 to 877 |
| | >Os11t0430600-00 | 4 | LOC_Os11g24240 | Sinapoylglucose choline | 53 to 57, 337 to 341, 360 to 364, 620 to 624 |
| | >Os11t0492300-00 | 5 | LOC_Os11g29990 | NBS-LRR type disease resistance protein | 224 to 228, 472 to 476, 541 to 545, 622 to 626, 792 to 796 |
| | >Os11t0529900-00 | 6 | LOC_Os11g32620 | Chalcone/stilbene synthase, N-terminal domain | 89 to 93, 97 to 101, 107 to 111, 120 to 124, 128 to 132, 138 to 142 |
| | >Os11t0549670-01 | 4 | - | Aspartate-semialdehyde dehydrogenase | 88 to 92, 191 to 195, 629 to 633, 840 to 844 |
| | >Os11t0549680-01 | 5 | LOC_Os11g34820 | Zinc knuckle family protein, expressed. THUMP domain- | 4 to 8, 15 to 19, 537 to 541, 585 to 589, 773 to 777 |
| | >Os11t0594700-00 | 5 | LOC_Os11g38210 | Protein of unknown function DUF538 family protein. | 32 to 36, 632 to 636, 665 to 669, 751 to 755, 760 to 764 |
| | >Os11t0655300-01 | 4 | LOC_Os11g43420 | NB-ARC domain | 49 to 53, 150 to 154, 511 to 515, 522 to 526 |
| **TGACT** | >Os11t0109400-01 | 4 | LOC_Os11g01820 | Cation/H exchanger domain. ATCHX | 77 to 81, 568 to 572, 782 to 786, 802 to 806 |
| | >Os11t0116550-01 | 4 | LOC_Os11g02464 | Vacuolar-sorting receptor precursor | 17 to 21, 125 to 129, 223 to 227, 314 to 318 |
| | >Os11t0133500-00 | 5 | LOC_Os11g03880 | Serine/threonine protein kinase-related domain. D-mannose binding lectin family | 102 to 106, 117 to 121, 285 to 289, 585 to 589, 903 to 907 |

| | | | | | |
|---|---|---|---|---|---|
| | >Os11t0160900-00 | 5 | LOC_Os11g06210 | NB-ARC domain | 112 to 116, 406 to 410, 493 to 497, 890 to 894, 954 to 958 |
| | >Os11t0173550-01 | 4 | LOC_Os11g07240 | Serine/threonine-protein kinase BRI1-like 2 precursor | 357 to 361, 371 to 375, 949 to 953, 953 to 957 |
| | >Os11t0201360-01 | 4 | LOC_Os11g09478 | Cyclin-like F-box domain. OsFBX402 | 36 to 40, 95 to 99, 379 to 383, 383 to 387 |
| | >Os11t0224200-00 | 4 | LOC_Os11g11680 | LGC1. LGC1 | 115 to 119, 184 to 188, 339 to 343, 510 to 514 |
| | >Os11t0226201-00 | 4 | LOC_Os11g11890 | Protein kinase domain | 178 to 182, 340 to 344, 614 to 618, 732 to 736 |
| | >Os11t0238000-00 | 4 | LOC_Os11g13440 | NB-ARC domain. RGH1A | 102 to 106, 317 to 321, 606 to 610, 793 to 797 |
| | >Os11t0244100-00 | 4 | LOC_Os11g13970 | Transferase family protein | 332 to 336, 345 to 349, 494 to 498, 622 to 626 |
| | >Os11t0252400-01 | 4 | LOC_Os11g14570 | Ankyrin repeat | 415 to 419, 425 to 429, 472 to 476, 489 to 493 |
| | >Os11t0252400-02 | 4 | LOC_Os11g14570 | Ankyrin repeat | 352 to 356, 362 to 366, 409 to 413, 426 to 430 |
| | >Os11t0256000-01 | 4 | LOC_Os11g14950 | ACT domain | 145 to 149, 346 to 350, 557 to 561, 561 to 565 |
| | >Os11t0306300-01 | 5 | LOC_Os11g20080 | Herbicide safener binding protein. O-methyltransferase | 145 to 149, 346 to 350, 557 to 561, 561 to 565 |
| | >Os11t0432400-01 | 4 | LOC_Os11g24450 | 2-oxoglutarate/malate translocator. mitochondrial carrier protein | 145 to 149, 346 to 350, 557 to 561, 561 to 565 |
| | >Os11t0433500-01 | 4 | LOC_Os11g24560 | Protein transport protein Sec23A | 258 to 262, 312 to 316, 685 to 689, 938 to 942 |
| | >Os11t0435300-00 | 4 | LOC_Os11g24780 | Ankyrin repeat domain | 71 to 75, 498 to 502, 548 to 552, 877 to 881 |
| | >Os11t0467632-01 | 4 | LOC_Os11g27870 | Aminotransferase-like. retrotransposon protein, Ty3-gypsy subclass | 482 to 486, 708 to 712, 753 to 757, 799 to 803 |
| | >Os11t0474500-00 | 4 | LOC_Os11g28470 | NB-ARC domain | 209 to 213, 710 to 714, 810 to 814, 897 to 901 |
| | >Os11t0493000-00 | 5 | LOC_Os11g30060 | NB-ARC domain. NBS-LRR type disease resistance | 206 to 210, 485 to 489, 560 to 564, 641 to 645, 744 to 748 |

| | | | | | |
|---|---|---|---|---|---|
| | | | | protein Hom-B | |
| | >Os11t0540600-01 | 4 | LOC_Os11g33394 | Protein of unknown function DUF247 | 210 to 214, 222 to 226, 267 to 271, 541 to 545 |
| | >Os11t0557500-00 | 4 | LOC_Os11g35330 | Protein kinase, core domain. LYK | 285 to 289, 299 to 303, 548 to 552, 700 to 704 |
| | >Os11t0565300-00 | 4 | LOC_Os11g35860 | Protein kinase domain. OsWAK120 - OsWAK receptor-like protein kinase | 100 to 104, 275 to 279, 424 to 428, 445 to 449 |
| | >Os11t0573000-02 | 4 | LOC_Os11g36470 | Peptidase C19, ubiquitin carboxyl-terminal hydrolase 2 family protein. ubiquitin carboxyl-terminal hydrolase 21 | 408 to 412, 519 to 523, 691 to 695, 699 to 703 |
| | >Os11t0598500-00 | 4 | LOC_Os11g38580 | NB-ARC domain | 362 to 366, 390 to 394, 404 to 408, 466 to 470 |
| | >Os11t0600900-01 | 4 | LOC_Os11g38810 | Mannose-6-phosphate isomerase (ManA) | 500 to 504, 564 to 568, 599 to 603, 662 to 666 |
| | >Os11t0655300-01 | 4 | LOC_Os11g43420 | NB-ARC domain. LZ-NBS-LRR class RGA | 17 to 21, 109 to 113, 479 to 483, 498 to 502, 724 to 728 |
| | >Os11t0655900-01 | 5 | LOC_Os11g43520 | Thioredoxin fold domain. OsGrx_C17 - glutaredoxin subgroup III | 17 to 21, 109 to 113, 479 to 483, 498 to 502, 724 to 728 |
| | >Os11t0668100-00 | 4 | LOC_Os11g44580 | NB-ARC domain. go35 NBS-LRR | 177 to 181, 765 to 769, 829 to 833, 849 to 853 |
| | >Os11t0669100-01 | 4 | LOC_Os11g44680 | Calmodulin binding protein-like family protein | 426 to 430, 436 to 440, 542 to 546, 574 to 578 |
| | >Os11t0702200-01 | 4 | LOC_Os11g47610 | Glycoside hydrolase, family 18 protein | 202 to 206, 897 to 901, 979 to 983, 989 to 993 |
| **TTGAC** | >Os12+B53:F91t0117 100-01 | 6 | LOC_Os12g02500 | Alpha/beta hydrolase fold-1 domain. esterase | 5 to 9, 234 to 238, 290 to 294, 464 to 468, 845 to 849, 854 to 858 |
| | >Os12t0118900-01 | 5 | LOC_Os12g02630 | Cytochrome P450 family protein | 248 to 252, 275 to 279, 455 to 459, 579 to 583, 970 to 974 |
| | >Os12t0120400-01 | 4 | LOC_Os12g02750 | ATPase-like protein | 806 to 810, 811 to 815, 816 to 820, 844 to 848 |
| | >Os12t0124100-01 | 4 | LOC_Os12g03080 | rp3 protein | 280 to 284, 444 to 448, 561 to 565, 803 to 807 |

| | | | | | |
|---|---|---|---|---|---|
| | >Os12t0128000-01 | 4 | LOC_Os12g03440 | Cyclin-like F-box domain OsFBX436 - F-box domain | 94 to 98, 432 to 436, 631 to 635, 659 to 663 |
| | >Os12t0128000-01 | 4 | LOC_Os12g03480 | Arabinoxylan arabinofuranohydrolase isoenzyme AXAH-I. alpha-N-arabinofuranosidase | 184 to 188, 212 to 216, 425 to 429, 515 to 519 |
| | >Os12t0128800-02 | 4 | LOC_Os12g03480 | Arabinoxylan arabinofuranohydrolase isoenzyme AXAH-I. alpha-N-arabinofuranosidase | 115 to 119, 143 to 147, 356 to 360, 446 to 450 |
| | >Os12t0150200-01 | 4 | LOC_Os12g05440 | Cytochrome P450 CYP94C20 | 281 to 285, 290 to 294, 309 to 313, 978 to 982 |
| | >Os12t0153500-01 | 4 | LOC_Os12g05730 | Carbonic anhydrase, CAH1-like domain | 142 to 146, 183 to 187, 188 to 192, 311 to 315 |
| | >Os12t0158300-01 | 4 | LOC_Os12g06150 | Protein of unknown function DUF623, plant domain | 260 to 264, 457 to 461, 477 to 481, 709 to 713 |
| | >Os12t0169000-00 | 4 | LOC_Os12g07150 | Amidase family protein | 9 to 13, 53 to 57, 121 to 125, 467 to 471 |
| | >Os12t0185900-01 | 4 | LOC_Os12g08550 | Zinc knuckle family protein | 454 to 458, 508 to 512, 746 to 750, 769 to 773 |
| | >Os12t0189900-01 | 5 | LOC_Os12g08800 | Las1-like family protein | 117 to 121, 168 to 172, 178 to 182, 276 to 280, 836 to 840 |
| | >Os12t0209200-01 | 5 | LOC_Os12g10660 | CONSTANS-LIKE a. B-box zinc finger family protein | 15 to 19, 491 to 495, 693 to 697, 702 to 706, 721 to 725 |
| | >Os12t0222800-00 | 4 | LOC_Os12g12120 | Leucine-rich repeat, N-terminal domain | 135 to 139, 278 to 282, 308 to 312, 581 to 585 |
| | >Os12t0233200-00 | 4 | LOC_Os12g13110 | pentatricopeptide | 156 to 160, 188 to 192, 338 to 342, 964 to 968 |
| | >Os12t0237900-01 | 4 | LOC_Os12g13550 | NB-ARC domain | 390 to 394, 398 to 402, 528 to 532, 942 to 946 |
| | >Os12t0274700-01 | 4 | LOC_Os12g17600 | Petunia ribulose 1,5-bisphosphate carboxylase small subunit mRNA | 9 to 13, 41 to 451, 90 to 194, 196 to 200 |
| | >Os12t0297500-02 | 4 | LOC_Os12g20150 | Pyruvate phosphate dikinase, PEP/pyruvate-binding domain. phosphoglucan, water dikinase, chloroplast precursor | 18 to 22, 422 to 426, 706 to 710, 886 to 890 |

| | | | | | |
|---|---|---|---|---|---|
| | >Os12t0411700-02 | 4 | LOC_Os12g22284 | ABC transporter-like domain | 248 to 252, 333 to 337, 373 to 377, 864 to 868 |
| | >Os12t0420400-01 | 4 | LOC_Os12g23200 | Photosystem I reaction center subunit XI | 20 to 24, 215 to 219, 655 to 659, 707 to 711 |
| | >Os12t0421000-01 | 5 | LOC_Os12g23280 | Barley stem rust resistance protein | 136 to 140, 510 to 514, 521 to 525, 565 to 569, 825 to 829 |
| | >Os12t0442800-01 | 4 | LOC_Os12g25630 | Sulfite oxidase | 78 to 82, 230 to 234, 459 to 463, 706 to 710 |
| | >Os12t0443000-00 | 4 | LOC_Os12g25660 | Cytochrome P450 | 104 to 108, 501 to 505, 526 to 530, 787 to 791 |
| | >Os12t0443200-00 | 4 | LOC_Os12g25680 | Papain family cysteine protease domain | 126 to 130, 154 to 158, 882 to 886, 909 to 913 |
| | >Os12t0506500-01 | 4 | LOC_Os12g32190 | Calmodulin-binding, plant domain | 12 to 16, 342 to 346, 766 to 770, 853 to 857 |
| | >Os12t0511900-00 | 4 | LOC_Os12g32710 | NB-ARC domain | 643 to 647, 662 to 666, 897 to 901, 914 to 918 |
| | >Os12t0516300-02 | 4 | LOC_Os12g33160 | NB-ARC domain | 15 to 19, 37 to 41, 403 to 407, 739 to 743 |
| | >Os12t0541700-01 | 4 | LOC_Os12g35670 | Rapid alkalinization factor 2. RALFL4 - Rapid ALkalinization Factor RALF family protein | 63 to 67, 229 to 233, 303 to 307, 322 to 326 |
| | >Os12t0556200-02 | 4 | LOC_Os12g36910 | Calmodulin binding protein-like family protein | 278 to 282, 307 to 311, 536 to 540, 793 to 797 |
| | >Os12t0561750-00 | 4 | | Ferroportin protein family | 148 to 152, 247 to 251, 318 to 322, 410 to 414 |
| | >Os12t0561900-01 | 5 | LOC_Os12g37510 | UDP-glucoronosyl and UDP-glucosyl transferase family. UDP-glucoronosyl | 17 to 21, 120 to 124, 732 to 736, 741 to 745, 760 to 764 |
| | >Os12t0568200-01 | 4 | LOC_Os12g38051 | Metallothionein-like protein type 1. metallothionein | 289 to 293, 341 to 345, 608 to 612, 924 to 928 |
| | >Os12t0573000-01 | 4 | LOC_Os12g38480 | Domain of unknown function DUF304l domain | 120 to 124, 359 to 363, 389 to 393, 778 to 782 |
| | >Os12t0573000-02 | 4 | LOC_Os12g38480 | Domain of unknown function DUF304l domain | 83 to 87, 322 to 326, 352 to 356, 741 to 745 |
| | >Os12t0582600-01 | 4 | LOC_Os12g39290 | TFIIS N-terminal domain | 538 to 542, 747 to 751, 813 to 817, 838 to 842 |
| | >Os12t0594600-01 | 4 | LOC_Os12g40300 | BTB/POZ fold domain | 266 to 270, 436 to 440, 468 to 472, 477 to 481 |

| | | | | | |
|---|---|---|---|---|---|
| | >Os12t0596800-01 | 4 | LOC_Os12g40490 | Zinc finger, LIM-type domain | 395 to 399, 455 to 459, 598 to 602, 861 to 865 |
| | >Os12t0597700-01 | 4 | LOC_Os12g40570 | WRKY94 | 382 to 386, 808 to 812, 821 to 825, 830 to 834 |
| | >Os12t0600200-03 | 4 | LOC_Os12g40790 | Zinc finger, C3HC4 type family protein. zinc finger family protein | 189 to 193, 699 to 703, 884 to 888, 963 to 967 |
| | >Os12t0604600-01 | 4 | LOC_Os12g41170 | GTP-binding protein-like; root hair defective 3 protein-like. SEY1 | 419 to 423, 559 to 563, 591 to 595, 748 to 752 |
| | >Os12t0610500-01 | 5 | LOC_Os12g41670 | SAM binding motif domain | 197 to 201, 212 to 216, 270 to 274, 569 to 573, 855 to 859 |
| | >Os12t0610600-01 | 4 | LOC_Os12g41680 | NAC60 | 238 to 242, 415 to 419, 590 to 594, 655 to 659 |
| | >Os12t0614500-00 | 4 | LOC_Os12g42010 | Lipase class 3 family protein | 65 to 69, 74 to 78, 93 to 97, 529 to 533 |
| | >Os12t0614800-00 | 5 | LOC_Os12g42040 | Receptor-like Cytoplasmic Kinase 374 | 308 to 312, 440 to 444, 593 to 597, 632 to 636, 857 to 861 |
| | >Os12t0615400-01 | 4 | LOC_Os12g42090 | 37 kDa inner envelope membrane protein, chloroplast precursor (E37). methyltransferase domain | 175 to 179, 496 to 500, 505 to 509, 524 to 528 |
| | >Os12t0630800-00 | 6 | LOC_Os12g43530 | NAC140 | 65 to 69, 92b to 96, 221 to 225, 269 to 273, 372 to 376, 706 to 710 |
| **TTGACA** | >Os11t0311300-01 | 4 | LOC_Os11g20689 | Exoribonuclease domain | 86 to 91, 289 to 294, 314 to 319, 688 to 693 |
| | >Os11t0311300-02 | 4 | LOC_Os11g20689 | Exoribonuclease domain | 84 to 89, 287 to 292, 312 to 317, 686 to 691 |
| | >Os11t0311300-03 | 4 | LOC_Os11g20689 | Exoribonuclease domain | 51 to 56, 254 to 259, 279 to 284, 653 to 658 |
| | >Os11t0311300-04 | 4 | LOC_Os11g20689 | Exoribonuclease domain | 46 to 51, 249 to 254, 274 to 279, 648 to 653 |
| | >Os11t0583200-01 | 4 | LOC_Os11g37330 | Pentatricopeptide repeat domain | 459 to 464, 623 to 628, 736 to 741, 863 to 868 |
| **TTGACC** | >Os11t0529900-00 | 4 | LOC_Os11g32620 | Chalcone/stilbene synthase | 88 to 93, 96 to 101, 119 to 124, 127 to 132 |
| | >Os11t0549680-01 | 4 | LOC_Os11g34820 | Zinc knuckle family protein | 3 to 8, 14 to 19, 536 to 541, 772 to 777 |
| **TTGACT** | >Os11t0133500-00 | 4 | LOC_Os11g03880 | Receptor-like Cytoplasmic Kinase 310 | 116 to 121, 284 to 289, 584 to 589, 902 to 907 |
| | >Os11t0160900-00 | 4 | LOC_Os11g06210 | NB-ARC domain | 111 to 116, 492 to 497, 889 to 894, 953 to 958 |
| | >Os11t0244100-00 | 4 | LOC_Os11g13970 | Transferase family protein | 331 to 336, 344 to 349, 493 to 498, 621 |

| | | | | | |
|---|---|---|---|---|---|
| | | | | | to 626 |
| | >Os11t0306300-01 | 4 | LOC_Os11g20080 | Herbicide safener binding protein. O-methyltransferase | 100 to 105, 298 to 303, 341 to 346, 460 to 465 |
| | >Os11t0655900-01 | 4 | LOC_Os11g43520 | Glutaredoxin 23 | 16 to 21, 478 to 483, 497 to 502, 723 to 728 |
| **Chromosome 12** | | | | | |
| **Motif** | **ID** | **Frequency** | **Locus ID** | **Protein name** | **Positions in 1k promoter region** |
| **CTGACT** | >Os12t0242100-01 | 4 | LOC_Os12g13890.1 | Glycine-rich cell wall structural protein 1 precursor | 291 to 296, 295 to 300, 381 to 386, 385 to 390 |
| **TGACC** | >Os12t0158300-01 | 4 | LOC_Os12g06150 | Protein of unknown function DUF623 | 261 to 265, 478 to 482, 910 to 914, 977 to 981 |
| | >Os12t0189900-01 | 6 | LOC_Os12g08800 | Las1-like family protein. | 261 to 265, 478 to 482, 910 to 914, 977 to 981 |
| | >Os12t0203500-00 | 4 | LOC_Os12g10250 | Zinc finger, RING-type domain | 261 to 265, 478 to 482, 910 to 914, 977 to 981 |
| | >Os12t0233100-01 | 4 | LOC_Os12g13100 | WW/Rsp5/WWP domain | 261 to 265, 478 to 482, 910 to 914, 977 to 981 |
| | >Os12t0233100-01 | 4 | LOC_Os12g13100 | WW/Rsp5/WWP domain | 514 to 518, 681 to 685, 907 to 911, 940to944 |
| | >Os12t0506500-01 | 7 | LOC_Os12g32190 | Calmodulin-binding | 514 to 518, 681 to 685, 907 to 911, 940 to 944 |
| | >Os12t0158300-01 | 4 | LOC_Os12g06150 | Protein of unknown function DUF623 | 261 to 265, 478 to 482, 910 to 914, 977 to 981 |
| | >Os12t0189900-01 | 6 | LOC_Os12g08800 | Las1-like family protein | 118 to 122, 169 to 173, 179 to 183, 277 to 281, 287 to 291, 351 to 355 |
| | >Os12t0203500-00 | 4 | LOC_Os12g10250 | Zinc finger, RING-type domain | 171 to 175, 194 to 198, 522 to 526, 755 to 759 |
| | >Os12t0233100-01 | 4 | LOC_Os12g13100 | WW/Rsp5/WWP domain | 514 to 518, 681 to 685, 907 to 911, 940 to 944 |
| | >Os12t0506500-01 | 7 | LOC_Os12g32190 | Calmodulin-binding | 13 to 17, 249 to 253, 767 to 771, 854 to 858, 937 to 941, 942 to 946, 947 to 951 |
| **TGACT** | >Os12t0109200-01 | 4 | LOC_Os12g01830 | Ca (2 )-dependent nuclease. staphylococcal nuclease homologue | 2 to 6, 338 to 342, 514 to 518, 754 to 758 |
| | >Os12t0135800-01 | 4 | LOC_Os12g04150 | Alpha/beta hydrolase fold-3 domain | 123 to 127, 234 to 238, 524 to 528, 528 to 532 |
| | >Os12t0168100-01 | 4 | LOC_Os12g07030 | ETHYLENE RESPONSE FACTOR 124 | 43 to 47, 404 to 408, 500 to 504, 884 to 888 |

| | | | | | |
|---|---|---|---|---|---|
| | >Os12t0178200-01 | 5 | LOC_Os12g07830 | Thylakoid-bound ascorbate OsAPx5 - Stromal Ascorbate Peroxidase encoding gene 5,8 | 309 to 313, 361 to 365, 627 to 631, 693 to 697, 727 to 731 |
| | >Os12t0188566-01 | 4 | | Thioredoxin (TRX) | 386 to 390, 397 to 401, 730 to 734, 928 to 932 |
| | >Os12t0192500-01 | 4 | LOC_Os12g09000 | Phosphomethylpyrimidine kinase type-1 domain | 57 to 61, 246 to 250, 537 to 541, 994 to 998 |
| | >Os12t0192500-02 | 4 | LOC_Os12g09000 | Phosphomethylpyrimidine kinase type-1 domain | 50 to 54, 239 to 243, 530 to 534, 987 to 991 |
| | >Os12t0197500-03 | 4 | LOC_Os12g09580 | Region of unknown function, putative Zinc finger, XS and XH domain. leafbladeless1 | 445 to 449, 620 to 624, 782 to 786, 811 to 815 |
| | >Os12t0242100-01 | 5 | LOC_Os12g13890) | Glycine-rich cell wall structural protein 1 precursor. retrotransposon protein | 292 to 296, 296 to 300, 382 to 386, 386 to 390, 786 to 790 |
| | >Os12t0263100-01 | 4 | LOC_Os12g16210 | Zinc finger, DHHC domain containing 4. zinc finger family protein | 534 to 538, 556 to 560, 638 to 642, 709 to 713 |
| | >Os12t0272800-01 | 4 | LOC_Os12g17410 | NB-ARC domain | 45 to 49, 66 to 70, 127 to 131, 752 to 756 |
| | >Os12t0277500-02 | 4 | LOC_Os12g17910 | RuBisCO subunit binding-protein alpha subunit, chloroplast precursor) T-complex protein | 34 to 38, 394 to 398, 426 to 430, 845 to 849 |
| | >Os12t0438300-01 | 4 | LOC_Os12g25170 | NB-ARC domain | 142 to 146, 800 to 804, 833 to 837, 990 to 994 |
| | >Os12t0454800-01 | 5 | LOC_Os12g26940 | Receptor-like serine/threonine kinase, Cytokinin signaling CHASE domain | 336 to 340, 422 to 426, 562 to 566, 628 to 632, 770 to 774 |
| | >Os12t0486900-01 | 4 | LOC_Os12g30180 | Protein kinase domain | 6 to 10, 542 to 546, 636 to 640, 678 to 682 |
| | >Os12t0512800-01 | 4 | LOC_Os12g32850 | Cytochrome P450 | 397 to 401, 404 to 408, 659 to 663, 915 to 919 |
| | >Os12t0517200-01 | 6 | LOC_Os12g33240 | Mitochondrial ribosomal protein S10 | 186 to 190, 205 to 209, 522 to 526, 544 to 548, 626 to 630, 955 to 959 |

| | | | | | |
|---|---|---|---|---|---|
| | >Os12t0538900-01 | 4 | LOC_Os12g35360 | Armadillo-type fold domain. vesicle tethering family protein | 99 to 103, 256 to 260, 484 to 488, 786 to 790 |
| | >Os12t0572500-01 | 4 | LOC_Os12g38440 | Region of unknown function XH domain | 241 to 245, 494 to 498, 704 to 708, 717 to 721 |
| | >Os12t0576700-01 | 4 | LOC_Os12g38760 | Diphosphonucleotide phosphatase 1 precursor. nucleotide pyrophosphatase/phosphodiesterase | 241 to 245, 617 to 621, 765 to 769, 769 to 773 |
| | >Os12t0596800-01 | 5 | | Zinc finger, LIM-type domain | 44 to 48, 121 to 125, 218 to 222, 274 to 278, 599 to 603 |
| | >Os12t0610600-01 | 4 | LOC_Os12g41680 | NAC60 | 239 to 243, 323 to 327, 591 to 595, 656 to 660 |
| | >Os12t0630800-00 | 4 | LOC_Os12g43530 | NAC 140 | 93 to 97, 222 to 226, 373 to 377, 707 to 711 |
| | >Os12t0638300-01 | 4 | LOC_Os12g44110 | Peptide transporter. ligA | 104 to 108, 482 to 486, 551 to 555, 910 to 914, 914 to 918 |
| **TTGAC** | >Os12t0117100-01 | 6 | LOC_Os12g02500 | Alpha/beta hydrolase fold-1 domain. esterase | 5 to 9, 234 to 238, 290 to 294, 464 to 468, 845 to 849, 854 to 858 |
| | >Os12t0118900-01 | 5 | LOC_Os12g02630 | Cytochrome P450 family protein | 248 to 252, 275 to 279, 455 to 459, 579 to 583, 970 to 974 |
| | >Os12t0120400-01 | 4 | LOC_Os12g02750 | ATPase-like protein | 806 to 810, 811 to 815, 816 to 820, 844 to 848 |
| | >Os12t0124100-01 | 4 | LOC_Os12g03080 | rp3 protein | 280 to 284, 444 to 448, 561 to 565, 803 to 807 |
| | >Os12t0128000-01 | 4 | LOC_Os12g03440 | Cyclin-like F-box domain OsFBX436 - F-box domain | 94 to 98, 432 to 436, 631 to 635, 659 to 663 |
| | >Os12t0128000-01 | 4 | LOC_Os12g03480 | Arabinoxylan arabinofuranohydrolase isoenzyme AXAH-I. alpha-N-arabinofuranosidase | 184 to 188, 212 to 216, 425 to 429, 515 to 519 |
| | >Os12t0128800-02 | 4 | LOC_Os12g03480 | Arabinoxylan arabinofuranohydrolase isoenzyme AXAH-I. alpha-N-arabinofuranosidase | 115 to 119, 143 to 147, 356 to 360, 446 to 450 |
| | >Os12t0150200-01 | 4 | LOC_Os12g05440 | Cytochrome P450 CYP94C20 | 281 to 285, 290 to 294, 309 to 313, 978 to 982 |

| | | | | | |
|---|---|---|---|---|---|
| | >Os12t0153500-01 | 4 | LOC_Os12g05730 | Carbonic anhydrase, CAH1-like domain containing protein. bifunctional | 142 to 146, 183 to 187, 188 to 192, 311 to 315 |
| | >Os12t0158300-01 | 4 | LOC_Os12g06150 | Protein of unknown function DUF623, plant domain | 260 to 264, 457 to 461, 477 to 481, 709 to 713 |
| | >Os12t0169000-00 | 4 | LOC_Os12g07150 | Amidase family protein | 9 to 13, 53 to 57, 121 to 125, 467 to 471 |
| | >Os12t0185900-01 | 4 | LOC_Os12g08550 | Zinc knuckle family protein | 454 to 458, 508 to 512, 746 to 750, 769 to 773 |
| | >Os12t0189900-01 | 5 | LOC_Os12g08800 | Las1-like family protein | 117 to 121, 168 to 172, 178 to 182, 276 to 280, 836 to 840 |
| | >Os12t0209200-01 | 5 | LOC_Os12g10660 | CONSTANS-LIKE a. B-box zinc finger family protein | 15 to 19, 491 to 495, 693 to 697, 702 to 706, 721 to 725 |
| | >Os12t0222800-00 | 4 | LOC_Os12g12120 | Leucine-rich repeat, N-terminal domain | 135 to 139, 278 to 282, 308 to 312, 581 to 585 |
| | >Os12t0233200-00 | 4 | LOC_Os12g13110 | Pentatricopeptide | 156 to 160, 188 to 192, 338 to 342, 964 to 968 |
| | >Os12t0237900-01 | 4 | LOC_Os12g13550 | NB-ARC domain | 390 to 394, 398 to 402, 528 to 532, 942 to 946 |
| | >Os12t0274700-01 | 4 | LOC_Os12g17600 | Petunia ribulose 1,5-bisphosphate carboxylase small subunit mRNA | 9 to 13, 41 to 451, 90 to 194, 196 to 200 |
| | >Os12t0297500-02 | 4 | LOC_Os12g20150 | Pyruvate phosphate dikinase, PEP/pyruvate-binding domain. phosphoglucan, water dikinase, chloroplast precursor | 18 to 22, 422 to 426, 706 to 710, 886 to 890 |
| | >Os12t0411700-02 | 4 | LOC_Os12g22284 | ABC transporter-like domain | 248 to 252, 333 to 337, 373 to 377, 864 to 868 |
| | >Os12t0420400-01 | 4 | LOC_Os12g23200 | Photosystem I reaction center subunit XI | 20 to 24, 215 to 219, 655 to 659, 707 to 711 |
| | >Os12t0421000-01 | 5 | LOC_Os12g23280 | Barley stem rust resistance protein | 136 to 140, 510 to 514, 521 to 525, 565 to 569, 825 to 829 |
| | >Os12t0442800-01 | 4 | LOC_Os12g25630 | Sulfite oxidase | 78 to 82, 230 to 234, 459 to 463, 706 to 710 |
| | >Os12t0443000-00 | 4 | LOC_Os12g25660 | Cytochrome P450 family | 104 to 108, 501 to 505, 526 to 530, 787 to 791 |
| | >Os12t0443200-00 | 4 | LOC_Os12g25680 | Papain family cysteine protease domain | 126 to 130, 154 to 158, 882 to 886, 909 to 913 |

| | | | | | |
|---|---|---|---|---|---|
| | >Os12t0506500-01 | 4 | LOC_Os12g32190 | Calmodulin-binding, plant domain | 12 to 16, 342 to 346, 766 to 770, 853 to 857 |
| | >Os12t0511900-00 | 4 | LOC_Os12g32710 | NB-ARC domain | 643 to 647, 662 to 666, 897 to 901, 914 to 918 |
| | >Os12t0516300-02 | 4 | LOC_Os12g33160 | NB-ARC domain | 15 to 19, 37 to 41, 403 to 407, 739 to 743 |
| | >Os12t0541700-01 | 4 | LOC_Os12g35670 | Rapid alkalinization factor 2 | 63 to 67, 229 to 233, 303 to 307, 322 to 326 |
| | >Os12t0556200-02 | 4 | LOC_Os12g36910 | Calmodulin binding protein-like family protein | 278 to 282, 307 to 311, 536 to 540, 793 to 797 |
| | >Os12t0561750-00 | 4 | | ferroportin protein family | 148 to 152, 247 to 251, 318 to 322, 410 to 414 |
| | >Os12t0561900-01 | 5 | LOC_Os12g37510 | UDP-glucoronosyl and UDP-glucosyl transferase family. UDP-glucoronosyl | 17 to 21, 120 to 124, 732 to 736, 741 to 745, 760 to 764 |
| | >Os12t0568200-01 | 4 | LOC_Os12g38051 | Metallothionein-like protein type 1. metallothionein | 289 to 293, 341 to 345, 608 to 612, 924 to 928 |
| | >Os12t0573000-01 | 4 | LOC_Os12g38480 | Domain of unknown function DUF304l domain | 120 to 124, 359 to 363, 389 to 393, 778 to 782 |
| | >Os12t0573000-02 | 4 | LOC_Os12g38480 | Domain of unknown function DUF304l domain | 83 to 87, 322 to 326, 352 to 356, 741 to 745 |
| | >Os12t0582600-01 | 4 | LOC_Os12g39290 | TFIIS N-terminal domain | 538 to 542, 747 to 751, 813 to 817, 838 to 842 |
| | >Os12t0594600-01 | 4 | LOC_Os12g40300 | BTB/POZ fold domain | 266 to 270, 436 to 440, 468 to 472, 477 to 481 |
| | >Os12t0596800-01 | 4 | LOC_Os12g40490 | Zinc finger, LIM-type domain | 395 to 399, 455 to 459, 598 to 602, 861 to 865 |
| | >Os12t0597700-01 | 4 | LOC_Os12g40570 | WRKY94 | 382 to 386, 808 to 812, 821 to 825, 830 to 834 |
| | >Os12t0600200-03 | 4 | LOC_Os12g40790 | Zinc finger, C3HC4 type family protein. zinc finger family protein | 189 to 193, 699 to 703, 884 to 888, 963 to 967 |
| | >Os12t0604600-01 | 4 | LOC_Os12g41170 | GTP-binding protein-like; root hair defective 3 protein-like. SEY1 | 419 to 423, 559 to 563, 591 to 595, 748 to 752 |
| | >Os12t0610500-01 | 5 | LOC_Os12g41670 | SAM binding motif domain | 197 to 201, 212 to 216, 270 to 274, 569 to 573, 855 to 859 |
| | >Os12t0610600-01 | 4 | LOC_Os12g41680 | NAC60 | 238 to 242, 415 to 419, 590 to 594, 655 to 659 |
| | >Os12t0614500-00 | 4 | LOC_Os12g42010 | lipase class 3 family protein | 65 to 69, 74 to 78, 93 to 97, 529 to 533 |
| | >Os12t0614800-00 | 5 | LOC_Os12g42040 | Receptor-like Cytoplasmic Kinase 374 | 308 to 312, 440 to 444, 593 to 597, 632 to 636, 857 to 861 |
| | >Os12t0615400-01 | 4 | LOC_Os12g42090 | 37 kDa inner envelope membrane protein, | 175 to 179, 496 to 500, 505 to 509, 524 to 528 |

| | | | | | |
|---|---|---|---|---|---|
| | | | | chloroplast precursor (E37). methyltransferase domain | |
| | >Os12t0630800-00 | 6 | LOC_Os12g43530 | NAC140 | 65 to 69, 92b to 96, 221 to 225, 269 to 273, 372 to 376, 706 to 710 |
| **TTGACA** | >Os12t0117100-01 | 4 | LOC_Os12g02500 | Alpha/beta hydrolase fold-1 domain | 5 to 10, 290 to 295, 464 to 469, 845 to 850 |
| | >Os12t0556200-02 | 4 | LOC_Os12g36910 | Calmodulin binding protein-like family protein | 278 to 283, 307 to 312, 536 to 541, 793 to 798 |
| **TTGACC** | >Os12t0189900-01 | 4 | LOC_Os12g08800 | Las1-like family protein | 117 to 122, 168 to 173, 178 to 183, 276 to 281 |
| **TTGACT** | >Os12t0630800-00 | 4 | LOC_Os12g43530 | NAC 140 | 92 to 97, 221 to 226, 372 to 377, 706 to 711 |

**Supplementary table 2.** Interacting residues of WRKY13-*Aminotransferase* complex

**Hydrophobic Interactions**

| Residue | AA | Distance | Ligand Atom | Protein Atom |
|---|---|---|---|---|
| **108A** | GLN | 3.61 | 928 | 93 |
| **109A** | LYS | 3.82 | 616 | 103 |
| **109A** | LYS | 3.78 | 963 | 102 |

**Hydrogen Bonds**

| Residue | AA | Distance H-A | Distance D-A | Donor Angle | Protein donor? | Side chain | Donor Atom | Acceptor Atom |
|---|---|---|---|---|---|---|---|---|
| **105A** | LYS | 2.88 | 3.39 | 113.28 | ✔ | ✘ | 65 [Nam] | 890 [O2] |
| **106A** | TYR | 2.92 | 3.36 | 108.39 | ✘ | ✘ | 680 [Npl] | 85 [O2] |
| **106A** | TYR | 3.07 | 3.85 | 141.80 | ✔ | ✔ | 81 [O3] | 657 [Nar] |
| **108A** | GLN | 1.86 | 2.81 | 161.01 | ✔ | ✘ | 90 [Nam] | 940 [Nar] |
| **108A** | GLN | 2.29 | 3.01 | 129.10 | ✔ | ✔ | 96 [Nam] | 911 [O3] |
| **109A** | LYS | 2.01 | 2.55 | 112.24 | ✘ | ✔ | 1004 [Npl] | 105 [N3] |
| **109A** | LYS | 1.16 | 2.11 | 152.15 | ✔ | ✔ | 105 [N3] | 618 [O2] |
| **134A** | GLN | 2.84 | 3.54 | 130.35 | ✘ | ✔ | 606 [O3] | 298 [O2] |
| **150A** | SER | 2.15 | 3.06 | 155.17 | ✔ | ✔ | 425 [O3] | 626 [O3] |
| **150A** | SER | 2.39 | 3.06 | 125.17 | ✘ | ✔ | 626 [O3] | 425 [O3] |

**Salt Bridges**

| Residue | AA | Distance | Protein positive? | Ligand Group | Ligand Atoms |
|---|---|---|---|---|---|
| **104A** | ARG | 3.97 | ✔ | Phosphate | 867, 867, 866, 868, 869, 870 |
| **105A** | LYS | 5.16 | ✔ | Phosphate | 908, 908, 907, 909, 910, 911 |
| **112A** | LYS | 4.45 | ✔ | Phosphate | 585, 585, 586, 587, 588, 584 |
| **118A** | ARG | 3.38 | ✔ | Phosphate | 605, 605, 604, 606, 607, 608 |
| **132A** | ARG | 3.51 | ✔ | Phosphate | 647, 647, 646, 649, 650, 648 |
| **132A** | ARG | 5.29 | ✔ | Phosphate | 625, 625, 626, 627, 628, 624 |

**Supplementary table 3.** Interacting residues of WRKY13-*Ankyrin* complex

| Hydrophobic Interactions | | | | |
|---|---|---|---|---|
| **Residue** | **AA** | **Distance** | **Ligand Atom** | **Protein Atom** |
| **143A** | THR | 3.56 | 552 | 369 |
| **144A** | VAL | 3.92 | 552 | 376 |
| **145A** | LEU | 3.78 | 559 | 383 |
| **146A** | LEU | 3.52 | 978 | 393 |
| **149A** | TYR | 2.87 | 600 | 418 |
| **149A** | TYR | 3.45 | 612 | 419 |

| Hydrogen Bonds | | | | | | | |
|---|---|---|---|---|---|---|---|
| **Residue** | **AA** | **Distance H-A** | **Distance D-A** | **Donor Angle** | **Protein donor?** | **Side chain** | **Donor Atom** |
| **118A** | ARG | 2.90 | 3.35 | 108.96 | ✓ | ✓ | 176 [Ng+] |
| **120A** | TYR | 2.70 | 3.26 | 116.80 | ✗ | ✗ | 949 [O3] |
| **134A** | GLN | 1.41 | 2.32 | 150.82 | ✓ | ✓ | 299 [Nam] |
| **142A** | PRO | 3.37 | 3.75 | 105.62 | ✗ | ✗ | 544 [O3] |
| **143A** | THR | 3.25 | 3.94 | 129.60 | ✓ | ✓ | 370 [O3] |
| **145A** | LEU | 1.24 | 2.20 | 162.93 | ✗ | ✗ | 595 [Npl] |
| **145A** | LEU | 2.20 | 3.14 | 158.10 | ✓ | ✗ | 380 [Nam] |
| **147A** | VAL | 1.59 | 2.56 | 169.20 | ✗ | ✗ | 614 [N2] |
| **147A** | VAL | 2.92 | 3.77 | 144.96 | ✓ | ✗ | 396 [Nam] |
| **148A** | THR | 2.44 | 3.21 | 136.84 | ✓ | ✓ | 407 [O3] |
| **148A** | THR | 1.57 | 2.25 | 121.84 | ✗ | ✓ | 980 [N2] |
| **149A** | TYR | 2.84 | 3.28 | 110.09 | ✓ | ✓ | 417 [O3] |

| Salt Bridges | | | | | |
|---|---|---|---|---|---|
| **Residue** | **AA** | **Distance** | **Protein positive?** | **Ligand Group** | **Ligand Atoms** |
| **118A** | ARG | 3.79 | ✓ | Phosphate | 947, 947, 946, 948, 949, 950 |
| **132A** | ARG | 4.47 | ✓ | Phosphate | 926, 926, 929, 925, 927, 928 |
| **133A** | LYS | 3.70 | ✓ | Phosphate | 583, 583, 585, 586, 582, 584 |
| **137A** | ARG | 4.22 | ✓ | Phosphate | 542, 542, 545, 541, 543, 544 |
| **139A** | ARG | 4.06 | ✓ | Phosphate | 968, 968, 969, 970, 971, 967 |
| **139A** | ARG | 4.56 | ✓ | Phosphate | 987, 987, 986, 988, 989, 990 |

**Supplementary table 4.** Interacting residues of WRKY13-*ger*6 complex

**Hydrophobic Interactions**

| Residue | AA | Distance | Ligand Atom | Protein Atom |
|---|---|---|---|---|
| 143A | THR | 3.70 | 947 | 369 |
| 145A | LEU | 3.87 | 954 | 384 |
| 149A | TYR | 3.16 | 996 | 418 |
| 149A | TYR | 3.68 | 1008 | 419 |

**Hydrogen Bonds**

| Residue | AA | Distance H-A | Distance D-A | Donor Angle | Protein donor? | Sidechain | Donor Atom | Acceptor Atom |
|---|---|---|---|---|---|---|---|---|
| **99A** | ASP | 3.58 | 3.96 | 105.47 | ✓ | ✗ | 7 [Nam] | 979 [O2] |
| **118A** | ARG | 2.96 | 3.46 | 112.52 | ✓ | ✓ | 176 [Ng+] | 591 [O2] |
| **120A** | TYR | 2.43 | 2.98 | 115.11 | ✗ | ✓ | 569 [O3] | 190 [O3] |
| **132A** | ARG | 3.30 | 4.02 | 130.91 | ✓ | ✓ | 281 [Ng+] | 527 [O3] |
| **134A** | GLN | 1.58 | 2.51 | 155.00 | ✓ | ✓ | 299 [Nam] | 569 [O3] |
| **142A** | PRO | 3.40 | 3.80 | 107.11 | ✗ | ✗ | 939 [O3] | 365 [O2] |
| **143A** | THR | 3.18 | 3.85 | 128.38 | ✓ | ✓ | 370 [O3] | 939 [O3] |
| **143A** | THR | 3.50 | 4.08 | 120.02 | ✗ | ✓ | 919 [O3] | 370 [O3] |
| **145A** | LEU | 1.75 | 2.70 | 162.35 | ✗ | ✗ | 990 [Npl] | 387 [O2] |
| **145A** | LEU | 2.10 | 3.04 | 158.99 | ✓ | ✗ | 380 [Nam] | 966 [Nar] |
| **147A** | VAL | 1.71 | 2.41 | 125.02 | ✗ | ✗ | 1010 [N2] | 402 [O2] |
| **147A** | VAL | 3.17 | 3.88 | 130.02 | ✓ | ✗ | 396 [Nam] | 623 [O2] |
| **148A** | THR | 1.96 | 2.74 | 135.51 | ✓ | ✓ | 407 [O3] | 577 [Nar] |
| **148A** | THR | 2.09 | 2.97 | 147.52 | ✗ | ✓ | 602 [Npl] | 407 [O3] |
| **149A** | TYR | 3.59 | 4.04 | 112.23 | ✓ | ✓ | 417 [O3] | 978 [O3] |

**Salt Bridges**

| Residue | AA | Distance | Protein positive? | Ligand Group | Ligand Atoms |
|---|---|---|---|---|---|
| **118A** | ARG | 3.79 | ✓ | Phosphate | 567, 567, 569, 570, 566, 568 |
| **132A** | ARG | 4.14 | ✓ | Phosphate | 546, 546, 545, 547, 548, 549 |
| **133A** | LYS | 4.09 | ✓ | Phosphate | 977, 977, 976, 978, 979, 980 |
| **137A** | ARG | 4.11 | ✓ | Phosphate | 937, 937, 938, 939, 940, 936 |
| **139A** | ARG | 3.98 | ✓ | Phosphate | 589, 589, 588, 590, 591, 592 |
| **139A** | ARG | 4.32 | ✓ | Phosphate | 610, 610, 609, 611, 612, 613 |

**Supplementary table 5.** Interacting residues of WRKY13-*βGlycosidase*14 complex

| Hydrophobic Interactions | | | | |
|---|---|---|---|---|
| **Residue** | **AA** | **Distance** | **Ligand Atom** | **Protein Atom** |
| 144A | VAL | 3.73 | 533 | 375 |
| 147A | VAL | 3.93 | 540 | 399 |
| 149A | TYR | 2.94 | 561 | 418 |

| Hydrogen Bonds | | | | | | | | |
|---|---|---|---|---|---|---|---|---|
| **Residue** | **AA** | **Distance H-A** | **Distance D-A** | **Donor Angle** | **Protein donor?** | **Sidechain** | **Donor Atom** | **Acceptor Atom** |
| **98A** | SER | 2.52 | 2.99 | 110.24 | ✖ | ✖ | 543 [O3] | 6 [O2] |
| **120A** | TYR | 1.39 | 2.14 | 129.27 | ✖ | ✔ | 992 [O3] | 190 [O3] |
| **120A** | TYR | 3.72 | 4.03 | 103.04 | ✔ | ✔ | 190 [O3] | 991 [O2] |
| **134A** | GLN | 2.52 | 3.31 | 137.79 | ✔ | ✔ | 299 [Nam] | 992 [O3] |
| **145A** | LEU | 2.50 | 2.98 | 109.52 | ✔ | ✖ | 380 [Nam] | 525 [O3] |
| **147A** | VAL | 2.89 | 3.74 | 145.77 | ✔ | ✖ | 396 [Nam] | 552 [Nar] |
| **147A** | VAL | 2.79 | 3.71 | 155.66 | ✖ | ✖ | 576 [Npl] | 402 [O2] |
| **148A** | THR | 2.97 | 3.64 | 125.62 | ✖ | ✔ | 597 [Npl] | 407 [O3] |
| **148A** | THR | 2.77 | 3.58 | 141.70 | ✔ | ✔ | 407 [O3] | 1000 [Nar] |

| Salt Bridges | | | | | |
|---|---|---|---|---|---|
| **Residue** | **AA** | **Distance** | **Protein positive?** | **Ligand Group** | **Ligand Atoms** |
| **118A** | ARG | 3.65 | ✔ | Phosphate | 990, 990, 993, 989, 991, 992 |
| **118A** | ARG | 5.30 | ✔ | Phosphate | 1012, 1012, 1011, 1013, 1014, 1015 |
| **132A** | ARG | 3.71 | ✔ | Phosphate | 949, 949, 948, 950, 951, 952 |
| **133A** | LYS | 4.98 | ✔ | Phosphate | 542, 542, 541, 545, 543, 544 |
| **139A** | ARG | 4.58 | ✔ | Phosphate | 1012, 1012, 1011, 1013, 1014, 1015 |
| **139A** | ARG | 5.43 | ✔ | Phosphate | 1032, 1032, 1033, 1034, 1035, 1031 |

**Supplementary table 6.** Interacting residues of WRKY13-*PR*1b complex

**Hydrophobic Interactions**

| Residue | AA | Distance | Ligand Atom | Protein Atom |
|---|---|---|---|---|
| 145A | LEU | 3.44 | 863 | 384 |
| 145A | LEU | 3.19 | 876 | 382 |
| 146A | LEU | 3.58 | 715 | 392 |
| 147A | VAL | 2.89 | 875 | 399 |
| 148A | THR | 3.67 | 653 | 406 |
| 149A | TYR | 3.53 | 918 | 419 |

**Hydrogen Bonds**

| Residue | AA | Distance H-A | Distance D-A | Donor Angle | Protein donor? | Sidechain | Donor Atom | Acceptor Atom |
|---|---|---|---|---|---|---|---|---|
| **99A** | ASP | 3.59 | 4.06 | 112.23 | ✘ | ✔ | 886 [O3] | 11 [O3] |
| **99A** | ASP | 1.96 | 2.79 | 139.52 | ✔ | ✘ | 7 [Nam] | 866 [O2] |
| **134A** | GLN | 3.09 | 3.82 | 132.59 | ✔ | ✔ | 299 [Nam] | 664 [O3] |
| **142A** | PRO | 2.76 | 3.44 | 127.88 | ✘ | ✘ | 825 [O3] | 365 [O2] |
| **145A** | LEU | 3.00 | 3.79 | 137.45 | ✘ | ✘ | 737 [Npl] | 387 [O2] |
| **145A** | LEU | 2.05 | 3.02 | 166.82 | ✔ | ✘ | 380 [Nam] | 854 [Nar] |
| **147A** | VAL | 2.66 | 3.59 | 159.40 | ✔ | ✘ | 396 [Nam] | 898 [O2] |
| **148A** | THR | 2.08 | 2.61 | 112.23 | ✔ | ✔ | 407 [O3] | 672 [Nar] |
| **148A** | THR | 2.84 | 3.68 | 143.64 | ✘ | ✔ | 697 [Npl] | 407 [O3] |

**Salt Bridges**

| Residue | AA | Distance | Protein positive? | Ligand Group | Ligand Atoms |
|---|---|---|---|---|---|
| **118A** | ARG | 4.13 | ✔ | Phosphate | 662, 662, 665, 661, 663, 664 |
| **132A** | ARG | 4.60 | ✔ | Phosphate | 622, 622, 625, 621, 623, 624 |
| **132A** | ARG | 5.05 | ✔ | Phosphate | 642, 642, 641, 643, 644, 645 |
| **133A** | LYS | 4.53 | ✔ | Phosphate | 865, 865, 864, 866, 867, 868 |
| **137A** | ARG | 5.30 | ✔ | Phosphate | 844, 844, 843, 845, 846, 847 |
| **137A** | ARG | 4.68 | ✔ | Phosphate | 823, 823, 825, 826, 824 |
| **139A** | ARG | 3.26 | ✔ | Phosphate | 684, 684, 683, 685, 686, 687 |

**Supplementary table 7.** Interacting residues of WRKY13-*PR2* complex

| Hydrophobic Interactions | | | | |
|---|---|---|---|---|
| **Residue** | **AA** | **Distance** | **Ligand Atom** | **Protein Atom** |
| **144A** | VAL | 3.73 | 533 | 375 |
| **147A** | VAL | 3.93 | 540 | 399 |
| **149A** | TYR | 2.94 | 561 | 418 |

| Hydrogen Bonds | | | | | | | | |
|---|---|---|---|---|---|---|---|---|
| **Residue** | **AA** | **Distance H-A** | **Distance D-A** | **Donor Angle** | **Protein donor?** | **Sidechain** | **Donor Atom** | **Acceptor Atom** |
| **98A** | SER | 2.52 | 2.99 | 110.24 | ✗ | ✗ | 543 [O3] | 6 [O2] |
| **120A** | TYR | 1.39 | 2.14 | 129.27 | ✗ | ✓ | 992 [O3] | 190 [O3] |
| **120A** | TYR | 3.72 | 4.03 | 103.04 | ✓ | ✓ | 190 [O3] | 991 [O2] |
| **134A** | GLN | 2.52 | 3.31 | 137.79 | ✓ | ✓ | 299 [Nam] | 992 [O3] |
| **145A** | LEU | 2.50 | 2.98 | 109.52 | ✓ | ✗ | 380 [Nam] | 525 [O3] |
| **147A** | VAL | 2.89 | 3.74 | 145.77 | ✓ | ✗ | 396 [Nam] | 552 [Nar] |
| **147A** | VAL | 2.79 | 3.71 | 155.66 | ✗ | ✗ | 576 [Npl] | 402 [O2] |
| **148A** | THR | 2.97 | 3.64 | 125.62 | ✗ | ✓ | 597 [Npl] | 407 [O3] |
| **148A** | THR | 2.77 | 3.58 | 141.70 | ✓ | ✓ | 407 [O3] | 1000 [Nar] |

| Salt Bridges | | | | | |
|---|---|---|---|---|---|
| **Residue** | **AA** | **Distance** | **Protein positive?** | **Ligand Group** | **Ligand Atoms** |
| **118A** | ARG | 3.65 | ✓ | Phosphate | 990, 990, 993, 989, 991, 992 |
| **118A** | ARG | 5.30 | ✓ | Phosphate | 1012, 1012, 1011, 1013, 1014, 1015 |
| **132A** | ARG | 3.71 | ✓ | Phosphate | 949, 949, 948, 950, 951, 952 |
| **133A** | LYS | 4.98 | ✓ | Phosphate | 542, 542, 541, 545, 543, 544 |
| **139A** | ARG | 4.58 | ✓ | Phosphate | 1012, 1012, 1011, 1013, 1014, 1015 |
| **139A** | ARG | 5.43 | ✓ | Phosphate | 1032, 1032, 1033, 1034, 1035, 1031 |

**Supplementary table 8.** Interacting residues of WRKY13-*PR5* complex

| Hydrophobic Interactions | | | | |
|---|---|---|---|---|
| Residue | AA | Distance | Ligand Atom | Protein Atom |
| 105A | LYS | 3.76 | 903 | 68 |
| 120A | TYR | 2.84 | 646 | 191 |

| Hydrogen Bonds | | | | | | | | |
|---|---|---|---|---|---|---|---|---|
| Residue | AA | Distance H-A | Distance D-A | Donor Angle | Protein donor? | Sidechain | Donor Atom | Acceptor Atom |
| **106A** | TYR | 3.60 | 4.08 | 112.84 | ✘ | ✘ | 723 [Npl] | 85 [O2] |
| **106A** | TYR | 1.76 | 2.26 | 107.86 | ✘ | ✘ | 703 [Npl] | 85 [O2] |
| **106A** | TYR | 3.58 | 3.98 | 108.70 | ✔ | ✔ | 81 [O3] | 679 [Nar] |
| **108A** | GLN | 2.85 | 3.78 | 158.09 | ✔ | ✘ | 90 [Nam] | 946 [Npl] |
| **108A** | GLN | 2.23 | 2.93 | 127.32 | ✘ | ✘ | 946 [Npl] | 98 [O2] |
| **108A** | GLN | 2.82 | 3.49 | 125.86 | ✔ | ✔ | 96 [Nam] | 895 [O3] |
| **109A** | LYS | 2.50 | 3.12 | 119.11 | ✔ | ✔ | 105 [N3] | 614 [Nar] |
| **134A** | GLN | 2.36 | 3.30 | 158.09 | ✔ | ✔ | 299 [Nam] | 629 [O3] |
| **150A** | SER | 2.79 | 3.66 | 151.15 | ✔ | ✔ | 425 [O3] | 649 [O3] |

| Salt Bridges | | | | | |
|---|---|---|---|---|---|
| Residue | AA | Distance | Protein positive? | Ligand Group | Ligand Atoms |
| **104A** | ARG | 4.57 | ✔ | Phosphate | 851, 851, 850, 852, 853, 854 |
| **105A** | LYS | 5.36 | ✔ | Phosphate | 892, 892, 891, 893, 894, 895 |
| **118A** | ARG | 3.29 | ✔ | Phosphate | 626, 626, 625, 627, 628, 629 |
| **122A** | ARG | 5.20 | ✔ | Phosphate | 669, 669, 668, 670, 671, 672 |
| **132A** | ARG | 4.91 | ✔ | Phosphate | 669, 669, 668, 670, 671, 672 |
| **132A** | ARG | 4.39 | ✔ | Phosphate | 648, 648, 649, 650, 651, 647 |

| $\pi$-Cation Interactions | | | | | | |
|---|---|---|---|---|---|---|
| Residue | AA | Distance | Offset | Protein charged? | Ligand Group | Ligand Atoms |
| **109A** | LYS | 3.58 | 1.97 | ✔ | Aromatic | 612, 613, 614, 615, 622 |

**Supplementary table 9.** Interacting residues of WRKY13-*aro*DE complex

| Hydrophobic Interactions | | | | |
|---|---|---|---|---|
| **Residue** | **AA** | **Distance** | **Ligand Atom** | **Protein Atom** |
| 106A | TYR | 3.73 | 720 | 83 |
| 108A | GLN | 3.50 | 888 | 93 |
| 109A | LYS | 3.54 | 658 | 103 |
| 109A | LYS | 4.00 | 923 | 102 |

| Hydrogen Bonds | | | | | | | | |
|---|---|---|---|---|---|---|---|---|
| **Residue** | **AA** | **Distance H-A** | **Distance D-A** | **Donor Angle** | **Protein donor?** | **Side chain** | **Donor Atom** | **Acceptor Atom** |
| 105A | LYS | 3.35 | 4.06 | 130.29 | ✔ | ✘ | 65 [Nam] | 851 [O2] |
| 106A | TYR | 3.27 | 4.08 | 141.31 | ✘ | ✘ | 741 [Npl] | 85 [O2] |
| 106A | TYR | 2.79 | 3.50 | 133.45 | ✔ | ✔ | 81 [O3] | 699 [Nar] |
| 108A | GLN | 2.03 | 2.94 | 153.66 | ✔ | ✘ | 90 [Nam] | 900 [Nar] |
| 108A | GLN | 3.10 | 3.97 | 147.19 | ✔ | ✔ | 96 [Nam] | 874 [O3] |
| 109A | LYS | 2.70 | 3.31 | 120.12 | ✘ | ✔ | 964 [Npl] | 105 [N3] |
| 109A | LYS | 1.42 | 2.14 | 121.60 | ✔ | ✔ | 105 [N3] | 660 [O2] |
| 122A | ARG | 3.64 | 4.01 | 105.13 | ✔ | ✔ | 215 [Ng+] | 691 [O2] |
| 134A | GLN | 3.12 | 3.96 | 143.79 | ✔ | ✔ | 299 [Nam] | 650 [O3] |
| 150A | SER | 2.29 | 3.22 | 158.91 | ✔ | ✔ | 425 [O3] | 668 [O3] |

| Salt Bridges | | | | | |
|---|---|---|---|---|---|
| **Residue** | **AA** | **Distance** | **Protein positive?** | **Ligand Group** | **Ligand Atoms** |
| **104A** | ARG | 4.20 | ✔ | Phosphate | 828, 828, 829, 830, 831 |
| **105A** | LYS | 4.80 | ✔ | Phosphate | 868, 868, 867, 869, 870, 871 |
| **112A** | LYS | 4.76 | ✔ | Phosphate | 627, 627, 626, 628, 629, 630 |
| **118A** | ARG | 3.17 | ✔ | Phosphate | 647, 647, 649, 650, 646, 648 |
| **132A** | ARG | 3.91 | ✔ | Phosphate | 689, 689, 690, 691, 692, 688 |
| **132A** | ARG | 5.03 | ✔ | Phosphate | 667, 667, 666, 668, 669, 670 |

**Supplementary Table 10.** Interacting residues of WRKY13-*WRKY*12 complex

| Hydrophobic Interactions | | | | |
|---|---|---|---|---|
| Residue | AA | Distance | Ligand Atom | Protein Atom |
| **148A** | THR | 3.34 | 914 | 409 |
| **149A** | TYR | 2.62 | 640 | 419 |
| **149A** | TYR | 2.89 | 652 | 417 |

| Hydrogen Bonds | | | | | | | | |
|---|---|---|---|---|---|---|---|---|
| Residue | AA | Distance H-A | Distance D-A | Donor Angle | Protein donor? | Sidechain | Donor Atom | Acceptor Atom |
| **99A** | ASP | 3.01 | 3.81 | 143.85 | ✓ | ✓ | 14 [O3] | 623 [O2] |
| **106A** | TYR | 3.01 | 3.88 | 154.83 | ✓ | ✓ | 85 [O3] | 883 [O2] |
| **120A** | TYR | 1.77 | 2.52 | 131.75 | ✗ | ✓ | 906 [O3] | 194 [O3] |
| **120A** | TYR | 1.69 | 2.47 | 134.07 | ✗ | ✓ | 905 [O3] | 194 [O3] |
| **132A** | ARG | 1.40 | 2.15 | 127.90 | ✓ | ✓ | 282 [Ng+] | 892 [Nar] |
| **132A** | ARG | 3.38 | 3.78 | 106.44 | ✓ | ✓ | 283 [Ng+] | 892 [Nar] |
| **143A** | THR | 1.59 | 2.41 | 138.96 | ✓ | ✓ | 371 [O3] | 585 [O3] |
| **143A** | THR | 2.90 | 3.37 | 110.91 | ✗ | ✗ | 585 [O3] | 369 [O2] |
| **145A** | LEU | 3.01 | 3.76 | 133.62 | ✓ | ✗ | 380 [Nam] | 604 [O3] |
| **145A** | LEU | 3.15 | 3.55 | 106.55 | ✗ | ✗ | 604 [O3] | 383 [O2] |
| **147A** | VAL | 1.78 | 2.58 | 136.25 | ✗ | ✗ | 654 [N2] | 399 [O2] |
| **149A** | TYR | 1.77 | 2.63 | 143.40 | ✗ | ✗ | 673 [N2] | 413 [O2] |
| **149A** | TYR | 2.92 | 3.82 | 151.95 | ✓ | ✗ | 410 [Nam] | 654 [N2] |

| Salt Bridges | | | | | |
|---|---|---|---|---|---|
| Residue | AA | Distance | Protein positive? | Ligand Group | Ligand Atoms |
| **118A** | ARG | 3.30 | ✓ | Phosphate | 904, 904, 903, 905, 906, 907 |
| **118A** | ARG | 5.02 | ✓ | Phosphate | 923, 923, 922, 924, 925, 926 |
| **132A** | ARG | 4.13 | ✓ | Phosphate | 882, 882, 881, 883, 884, 885 |

**Supplementary Table 11.** Interacting residues of WRKY13-*TIFY*9 complex

**Hydrophobic Interactions**

| Residue | AA | Distance | Ligand Atom | Protein Atom |
|---|---|---|---|---|
| 143A | THR | 4.00 | 884 | 369 |
| 145A | LEU | 3.88 | 947 | 382 |
| 146A | LEU | 3.21 | 641 | 390 |
| 147A | VAL | 1.98 | 947 | 399 |

**Hydrogen Bonds**

| Residue | AA | Distance H-A | Distance D-A | Donor Angle | Protein donor? | Side chain | Donor Atom | Acceptor Atom |
|---|---|---|---|---|---|---|---|---|
| **120A** | TYR | 2.77 | 3.37 | 120.40 | ✘ | ✔ | 611 [O3] | 190 [O3] |
| **132A** | ARG | 3.33 | 3.99 | 126.48 | ✔ | ✔ | 281 [Ng+] | 569 [O3] |
| **134A** | GLN | 1.72 | 2.65 | 156.98 | ✔ | ✔ | 299 [Nam] | 611 [O3] |
| **143A** | THR | 3.47 | 4.07 | 122.34 | ✔ | ✔ | 370 [O3] | 876 [O3] |
| **145A** | LEU | 3.12 | 3.90 | 138.01 | ✘ | ✘ | 685 [Npl] | 387 [O2] |
| **145A** | LEU | 2.15 | 3.13 | 171.94 | ✔ | ✘ | 380 [Nam] | 924 [Nar] |
| **147A** | VAL | 1.59 | 2.48 | 148.21 | ✘ | ✘ | 968 [Npl] | 402 [O2] |
| **147A** | VAL | 3.02 | 3.83 | 140.47 | ✔ | ✘ | 396 [Nam] | 663 [O2] |
| **148A** | THR | 3.47 | 4.05 | 119.66 | ✘ | ✔ | 622 [Npl] | 407 [O3] |
| **148A** | THR | 1.70 | 2.39 | 125.52 | ✔ | ✔ | 407 [O3] | 619 [Nar] |

**Salt Bridges**

| Residue | AA | Distance | Protein positive? | Ligand Group | Ligand Atoms |
|---|---|---|---|---|---|
| **118A** | ARG | 3.85 | ✔ | Phosphate | 609, 609, 608, 610, 611, 612 |
| **118A** | ARG | 5.41 | ✔ | Phosphate | 630, 630, 633, 629, 631, 632 |
| **132A** | ARG | 4.32 | ✔ | Phosphate | 588, 588, 587, 589, 590, 591 |
| **133A** | LYS | 4.13 | ✔ | Phosphate | 936, 936, 935, 937, 938, 939 |
| **137A** | ARG | 4.47 | ✔ | Phosphate | 893, 893, 892, 894, 895, 896 |
| **139A** | ARG | 4.01 | ✔ | Phosphate | 630, 630, 633, 629, 631, 632 |

**Supplementary table 12.** Interacting residues of WRKY13-*WRKY*13 complex

**Hydrophobic Interactions**

| Residue | AA | Distance | Ligand Atom | Protein Atom |
|---|---|---|---|---|
| **106A** | TYR | 3.03 | 961 | 83 |
| **108A** | GLN | 3.74 | 643 | 93 |
| **109A** | LYS | 3.60 | 677 | 102 |
| **120A** | TYR | 3.92 | 906 | 191 |

**Hydrogen Bonds**

| Residue | AA | Distance H-A | Distance D-A | Donor Angle | Protein donor? | Side chain | Donor Atom | Acceptor Atom |
|---|---|---|---|---|---|---|---|---|
| **105A** | LYS | 3.52 | 4.08 | 118.83 | ✗ | ✗ | 607 [O3] | 73 [O2] |
| **106A** | TYR | 2.94 | 3.67 | 135.01 | ✓ | ✓ | 81 [O3] | 939 [Nar] |
| **106A** | TYR | 1.54 | 2.15 | 114.98 | ✗ | ✗ | 637 [Npl] | 85 [O2] |
| **106A** | TYR | 2.81 | 3.18 | 102.92 | ✗ | ✗ | 658 [Npl] | 85 [O2] |
| **108A** | GLN | 2.06 | 2.94 | 148.02 | ✓ | ✗ | 90 [Nam] | 655 [Nar] |
| **108A** | GLN | 3.33 | 4.00 | 127.21 | ✓ | ✓ | 96 [Nam] | 644 [O3] |
| **109A** | LYS | 3.34 | 3.77 | 108.59 | ✗ | ✓ | 921 [Npl] | 105 [N3] |
| **109A** | LYS | 2.25 | 2.66 | 104.06 | ✗ | ✓ | 719 [Npl] | 105 [N3] |
| **109A** | LYS | 1.39 | 2.24 | 135.48 | ✓ | ✓ | 105 [N3] | 901 [O2] |
| **122A** | ARG | 3.45 | 4.06 | 122.21 | ✓ | ✓ | 215 [Ng+] | 931 [O2] |
| **134A** | GLN | 2.94 | 3.73 | 138.47 | ✓ | ✓ | 299 [Nam] | 891 [O3] |
| **134A** | GLN | 2.20 | 3.05 | 145.67 | ✗ | ✓ | 889 [O3] | 298 [O2] |
| **150A** | SER | 1.79 | 2.74 | 166.28 | ✓ | ✓ | 425 [O3] | 909 [O3] |

**Salt Bridges**

| Residue | AA | Distance | Protein positive? | Ligand Group | Ligand Atoms |
|---|---|---|---|---|---|
| **104A** | ARG | 4.27 | ✓ | Phosphate | 585, 585, 584, 586, 587, 588 |
| **105A** | LYS | 4.86 | ✓ | Phosphate | 624, 624, 623, 625, 626, 627 |
| **112A** | LYS | 5.30 | ✓ | Phosphate | 867, 867, 866, 868, 869, 870 |
| **118A** | ARG | 3.65 | ✓ | Phosphate | 888, 888, 887, 889, 890, 891 |
| **132A** | ARG | 4.07 | ✓ | Phosphate | 929, 929, 930, 931, 932, 928 |
| **132A** | ARG | 4.49 | ✓ | Phosphate | 908, 908, 907, 909, 910, 911 |
